# Phylogenomic Discordance in the Eared Seals is best explained by Incomplete Lineage Sorting following Explosive Radiation in the Southern Hemisphere

**DOI:** 10.1101/2020.08.11.246108

**Authors:** Fernando Lopes, Larissa R. Oliveira, Amanda Kessler, Yago Beux, Enrique Crespo, Susana Cárdenas-Alayza, Patricia Majluf, Maritza Sepúlveda, Robert L. Brownell, Valentina Franco-Trecu, Diego Páez-Rosas, Jaime Chaves, Carolina Loch, Bruce C. Robertson, Karina Acevedo-Whitehouse, Fernando R. Elorriaga-Verplancken, Stephen P. Kirkman, Claire R. Peart, Jochen B. W. Wolf, Sandro L. Bonatto

## Abstract

The phylogeny and systematics of fur seals and sea lions (Otariidae) have long been studied with diverse data types, including an increasing amount of molecular data. However, only a few phylogenetic relationships have reached acceptance because of strong gene-tree species tree discordance. Divergence times estimates in the group also vary largely between studies. These uncertainties impeded the understanding of the biogeographical history of the group, such as when and how trans-equatorial dispersal and subsequent speciation events occurred. Here we used high-coverage genome-wide sequencing for 14 of the 15 species of Otariidae to elucidate the phylogeny of the family and its bearing on the taxonomy and biogeographical history. Despite extreme topological discordance among gene trees, we found a fully supported species tree that agrees with the few well-accepted relationships and establishes monophyly of the genus *Arctocephalus*. Our data support a relatively recent trans-hemispheric dispersal at the base of a southern clade, which rapidly diversified into six major lineages between 3 to 2.5 Ma. *Otaria* diverged first, followed by *Phocarctos* and then four major lineages within *Arctocephalus*. However, we found *Zalophus* to be non-monophyletic, with California *(Z. californianus)* and Steller sea lions *(Eumetopias jubatus)* grouping closer than the Galapagos sea lion (*Z. wollebaeki)* with evidence for introgression between the two genera. Overall, the high degree of genealogical discordance was best explained by incomplete lineage sorting resulting from quasi-simultaneous speciation within the southern clade with introgresssion playing a subordinate role in explaining the incongruence among and within prior phylogenetic studies of the family.

## Introduction

For some time, it was widely accepted that by increasing the volume of molecular data even simple phylogenetic methods would unravel the true phylogenetic history of species (Rokas et al. 2003; Faircloth et al. 2013; Hoban et al. 2013; McCormack and Faircloth 2013). However, studies using whole genome data have found that inference of the true species tree, if such a tree exists, may be extremely challenging for some parts of the tree of life (Nakhleh 2013). These difficulties stem from a high degree of genealogical discordance among genomic fragments (GF) trees estimated from partitioned genomic data (e.g., genes or independent genomic fragments) (Peter 2016; Harris and DeGiorgio 2016; Elworth et al. 2018; Jones 2019).

Theoretical and empirical studies have shown that genealogical incongruences have three leading causes: incorrect estimation of the gene trees (e.g., caused by insufficient phylogenetic information, incorrect model specification or intralocus recombination); incomplete lineage sorting (ILS), found when ancestral polymorphism is persistent between successive speciation events (see Maddison and Knowles 2006; Oliver 2013); and introgression between lineages (hybridization) (e.g., Rheindt et al. 2014; Figueiró et al. 2017; Zhang et al. 2017). While technical issues as in the first problem could, in theory, be resolved, the latter two reflect the biological reality of evolutionary independence among recombining, genomic fragments (Hudson 1983; Griffiths and Marjoram 1997). Most methods used to estimate species trees assume only ILS, ignoring the consequences of hybridization for phylogenetic reconstruction (Stamatakis 2014; Drummond and Bouckaert 2015). Despite recent progress in developing models that include introgression, such as the so-called multispecies network models (e.g., Leaché et al. 2014; Wen and Nakhleh 2018), they continue to present several limitations, in particular when dealing with more than a few species (Degnam 2018). Consequently, resolving relationships among species that radiated rapidly and putatively underwent both ILS and hybridization has proven challenging (Chakrabarty et al. 2017; Esselstyn et al. 2017; Reddy et al. 2017).

The difficulty in establishing phylogenetic consensus (see below) and evidence for current hybridization (Lancaster et al. 2006) make the otariids a compelling test case to assess the relative impact of ancestral polymorphism and introgression on phylogenetic reconstruction during rapid diversification. There are 15 extant species of fur seals and sea lions within the Otariidae (Berta et al. 2018) with some uncertainty regarding the taxonomic status of species such as *Arctocephalus philippii* and *A. townsendi* (see Committee on Taxonomy of the Society for Marine Mammalogy 2020 for details, see Repenning et al. 1971; Yonezawa et al. 2009; Berta and Churchill 2012; Churchill et al. 2014, Berta et al. 2018). The initial diversification of the main lineages of Otariidae occurred around 11 (Yonezawa et al. 2009) to 9 Ma (Berta et al. 2018, Nyakatura and Bininda-Emonds 2014, this study). On the other hand, species in the disputed genus *Arctocephalus* (see below) emerged during a near-simultaneous succession of cladogenetic events within less than 0.5 Ma (Berta et al. 2018; this study) corresponding to approximately 2.5 *Ne* generations (estimated from data in Suppl. Table 3 of Peart et al. 2020). During such a short period, lineage sorting is expected to be incomplete (Hudson et al. 2003; Rosenberg et al. 2003; Mugal et al. 2020) with putative events of hybridization occurring, which makes this group particularly suited to investigate the underpinnings of gene tree species tree discordance.

Otariids occur in the North Pacific Ocean and Southern Hemisphere and are found from tropical waters in the eastern Pacific to polar regions (Churchill et al. 2014; Berta et al. 2018). Although the systematics and phylogeny of the family have been extensively studied for over 100 years (Sclater 1897; Scheffer 1958; Wynen et al. 2001; Deméré et al. 2003; Árnason et al. 2006; Yonezawa et al. 2009; Berta and Churchill 2012; Nyakatura and Bininda-Emonds 2012; Churchill et al. 2014; Berta et al. 2018), several relationships, in particular those within *Arctocephalus*, the most diverse (eight species) otariid genus, remain unclear (Yonezawa et al. 2009; Berta and Churchill 2012). For example, older studies based on morphology suggested grouping the fur seals *(Callorhinus ursinus* and *Arctocephalus* spp.) in the Arctocephalinae, which are characterized by small body size and thick pelage, and the sea lions in the Otariinae, which are characterized by larger body size and reliance on blubber rather than fur for thermal insulation (Berta and Demeré 1986; see review in Berta et al. 2018). However, more recent studies that used a combination of a few mitochondrial or nuclear genes and morphological data did not support these subfamilies (e.g., Yonezawa et al. 2009; Berta and Churchill et al. 2012; Churchill et al. 2014; Nyakatura and Bininda-Emonds 2014; Berta et al. 2018). Most of these phylogenies grouped the Southern Hemisphere otariids (i.e., *Otaria, Neophoca, Phocarctos*, and *Arctocephalus*) in the so-called southern clade, which is considered the sister clade of the sea lions of the Northern Hemisphere (i.e., *Zalophus* and *Eumetopias*) (Yonezawa et al. 2009, Churchill et al. 2014).

Another major difference between studies concerns the monophyly of *Arctocephalus*. A combined phylogeny produced by analyzing published morphological and molecular data reported *Arctocephalus sensu lato* as paraphyletic (Berta and Churchill 2012), restricting the genus to the type species *Arctocephalus pusillus*, and assigning the remaining species to *Arctophoca.* Other authors proposed that the use of *Arctophoca* was premature because of the remaining uncertainties surrounding the phylogenetic relationships in the group (e.g., Nyakatura and Bininda-Emonds 2014). Subsequently, the Committee on Taxonomy of the Society for Marine Mammalogy, which initially supported the proposal of Berta and Churchill (2012), adopted the conservative use of *Arctocephalus sensu lato* for all southern fur seals pending further studies (Committee on Taxonomy 2020). In short, there seemed to be no two identical phylogenies for the family and no explanation for the high level of discordance between studies.

The divergence times and biogeography within the Otariidae also present uncertainties, given the disagreement between studies. The most recent biogeographical studies (e.g., Yonezawa et al. 2009; Churchill et al. 2014) agree on a North Pacific origin for Otariidae and support the hypothesis of one primary trans-equatorial dispersal event into the eastern South Pacific Ocean, that gave rise to the Southern Hemisphere clade (see Churchill et al. 2014). It has been estimated that this dispersal event and the diversification of the southern clade occurred at ~7 - 6 Ma (Yonezawa et al. 2009, Churchill et al. 2014, Berta et al. 2018). The more recent diversification within *Arctocephalus* may have occurred 4-3 Ma (Nyakatura and Bininda-Emonds 2012) or as recently as <1 Ma (Berta et al. 2018).

In this study, we used whole genome sequence data to investigate the phylogenetic relationships and estimate the divergence times of Otariidae species. We used several phylogenomic approaches, including multispecies coalescent models, to clarify most of the unresolved issues in the evolutionary history of Otariidae. We also investigated the main factors responsible for the high level of topological incongruences within the family, finding they were caused by rampant incomplete lineage sorting and some introgression events.

## Material and Methods

### Sample Collection and Genome Sequencing

Skin samples from nine otariid species (Table 1) were collected from live or fresh carcasses found ashore. Piglet ear notch pliers were used to extract ~0.5 cm^3^ skin samples. The samples were stored in ethanol 70% and cryo-preserved at −20 °C. Genomic DNA extractions were carried out with DNeasy Tissue Kit (Qiagen) following the manufacturer’s protocol.

We sequenced the whole genome of one individual from seven species of *Arctocephalus* and two other monospecific genera (*Phocarctos* and *Otaria*) (Table 1 and Supplementary Table S1 available on Dryad at https://doi:10.5061/dryad.pzgmsbchw). Genomic libraries were prepared with Illumina DNA PCR-free or TruSeq Nano kits with an insert size of 350 bp, and two libraries were sequenced (PE150) per lane on the Illumina HiSeq X platform. Raw genome reads from *Arctocephalus gazella, Zalophus wollebaeki, Zalophus californianus*, *Eumetopias jubatus*, and *Callorhinus ursinus* (Table 1 and Supplementary Table S1) were retrieved from the NCBI Sequencing Read Archive (https://www.ncbi.nlm.nih.gov/sra). We used the genome of the walrus, *Odobenus rosmarus* (ANOP00000000 - Foote et al. 2015) as the reference for mapping and as the outgroup for most analyses. Since we had already started several analyses before genome-wide data from *C. ursinus*, *E. jubatus*, and *Z. californianus* were available, we did not include them in some less critical but time-consuming analyses.

Our study included 14 of the 15 extant Otariidae species (all *Arctocephalus, Phocarctos*, *Otaria*, *Zalophus, Eumetopias*, and *Callorhinus*). *Neophoca cinerea* was not included in our study. However, its position as the sister species of *P. hookeri* is uncontentious (see Yonezawa et al. 2009; Berta and Churchill 2012; Nyakatura and Bininda-Emonds 2012; Berta et al. 2018).

Sequencing quality control was performed using FastQC (Andrews 2010). Reads were trimmed for vestigial adapters, mapped against the *O. rosmarus* genome and locally realigned using the bam_pipeline implemented on PALEOMIX 1.2.13.2 (Schubert et al. 2014). Reads with length-size <100 bp and Phred-score <30 were filtered out by AdapterRemoval v2 (Schubert et al. 2016); the remaining paired-end reads were mapped using BWA 0.7.17 (Li and Durbin 2009) and the-mem algorithm. Paired-end reads with mapping quality Phred-score < 20, unmapped reads and single-reads were discarded from the downstream pipeline and reads that were sequenced more than two or less than one standard deviations from the average of coverage of each genome (Supplementary Table S2) were not used in the analyses (Arnold et al. 2013; Gautier et al. 2013). PCR duplicates were detected and removed by Picard Tools 2.18.5 (broadinstitute.github.io/picard/), and miscalling indels were locally realigned by GATK 3.8 (McKenna et al. 2010).

### Consensus, Alignments and SNP Calling

Consensus sequences of all genomes were generated with ANGSD 0.921 (Korneliussen et al. 2014) using the parameters *doFasta 2, doCounts1*, and *explode 1.* Single-nucleotide polymorphisms (SNPs) were called following the filters: *uniqueOnly 1, remove_bads 1, only_proper_pairs 1, C 50, baq 1, setMinDepth* 140, *setMaxDepth 1400, setMinDepthInd 5, setMaxDepthInd 100, doCounts 1, GL 1, doMajorMinor 1, SNP_pval 1e-3, doGeno 32, doPost 1, doPlink2*. After the SNP calling, a PLINK variant panel was converted to VCF format with Plink 1.9 (Chang et al. 2015). The VCF file did not contain SNPs from the walrus genome. We removed all information of repetitive, coding, and transposons present in the General Feature Format File of *O. rosmarus* genome with BEDTools 2.27.0 maskfasta option (Quinlan and Hall 2010).

### Phylogenetic Information, Phylogenomic Analyses, and Genealogical Discordance Estimation

We first estimated relationships between species using the full sequence data set. A whole-genome maximum-likelihood (ML) tree was inferred with RAxML-NG-MPI (Kozlov et al. 2019) directly from the SNP panel using the HKY substitution model inferred with ModelTest-NG, 100 bootstrap replicates and *C. ursinus* as the outgroup. We also used the VCF2Dis script (github.com/BGI-shenzhen) to estimate the p-distance matrix from the VCF file, followed by a neighbor-joining tree with PHYLIP 3.697 (Felsenstein 1989). Additionally, we estimated ML trees for each alignment of the ten largest scaffolds with RAxML-HPC-PTHREADS 8.2 (Stamatakis 2014) using GTR+G (best-fit substitution model as estimated by ModelTest-NG for all the largest scaffolds, Darriba et al. 2019) and 100 bootstrap replicates.

Next, we estimated phylogenies using smaller segments partitioning scaffolds into sets of smaller nonoverlapping genomic fragments (GFs) of 10, 20, 50, 80, 100, and 200 kilobases (kb) in length. To reduce the effect of linkage disequilibrium between GFs, they were separated by 100 kb, regardless of window size, following Humble et al. (2018) demonstrating low levels of linkage disequilibrium (r^2^~0.05) at this physical distance in the Antarctic fur seal. Several filters were used: scaffolds smaller than the GF partition size were excluded; sites with more than 20% of missing data were removed with trimAl v1.4 (Capella-Gutierrez et al. 2009); alignments smaller than half of the original alignment size were also discarded. To reduce the effect of intra-fragment genetic recombination on the phylogenetic estimation, we used the software 3Seq on *full run mode* (Lam et al. 2017). We removed the alignments with evidence of recombination at a *p-value* < 0.01 after Bonferroni correction (Rice 1989). To test the effect in quantification of the genealogical discordance (see below) of both the spacing of GFs by 100 kb and of the 3Seq filtering for recombination, we generated additionally datasets (only 50 kb GFs): with no 3Seq filtering (i.e. with all GFs) and without the 100 kb spacing (i.e. contiguous GFs).

To assess the amount of genetic information content on GFs, we randomly sampled 10,000 GFs of 50 kb and used the AMAS tool (Borowiec 2016) to count the number of parsimony-informative sites in these alignments and the number of differences between two closely related fur seals (*A. australis* and *A. galapagoensis*). Finally, we reconstructed ML trees with RAxML-HPC-PTHREADS 8.2 for each GF that passed by the mentioned filters in all GF partitions (10 to 200 kb) using the same parameters as above.

To quantify the genealogical discordance throughout genomes, we counted the frequency of each topology with Newick Utilities 1.1 (Junier and Zdobnov 2010) for each set of GF trees by using the sub-programs nw_topology and nw_order in a pipeline. We also estimated the gene concordance factor (gCF) and the site concordance factor (sCF) (Minh et al. 2018) implemented in IQ-TREE 1.7 (Nguyen et al. 2015) as a complement to standard measures of branch support (in this case bootstrap) and to quantify the disagreement among loci and sites in our phylogenomic dataset. The gCF is the percentage of decisive GF trees showing a particular branch from a species tree, while sCF is the percentage of decisive alignment sites supporting a branch in the reference tree when individual gene alignments are relatively uninformative (Minh et al. 2018). The estimation of gCF and sCF followed three steps. First, in IQ-TREE, the species phylogeny used as reference was recovered based on all (10,806) GFs of 50 kb concatenated, the edge-linked proportional partition model and 1,000 replicates of ultrafast bootstraping. Second, using a maximum-likelihood approach and substitution models inferred for each locus, GF trees were estimated from each genomic fragment. Then, gCF and sCF were computed across all nodes of the generated species tree and GF trees. The outputs were visualized with the support of the R script available on http://www.robertlanfear.com/blog/files/concordance_factors.html

### Species Tree Estimation

Two methods were used to reconstruct the species tree from multiple GF trees. First, all GFs ML trees were used to estimate a maximum quartet support species tree with the multispecies coalescent model (MSC) of ASTRAL-III (Zhang et al. 2018) by applying the exact search method. Second, we estimated the species tree and divergence times with the Bayesian Inference method StarBEAST2 implemented in the BEAST 2.5.2 package (Rambaut and Drummond 2010; Bouckaert et al. 2014; Ogilvie et al. 2017). Since this Bayesian analysis is very time-consuming and the ASTRAL species trees of all GF datasets, except the 10 kb GF, were identical (see Results), we used 300 randomly selected GFs from the 50 kb dataset. The main priors used were: linked clock models, constant population sizes, the HKY substitution model with empirical base frequencies, an estimated six gamma categories site model, and the Yule Tree model. To estimate divergence times, we used a strict molecular clock as a prior with a lognormal distribution and a standard mammalian genomic mutation rate of 1×10^−8^ bp^−1^ gen^−1^ (Kumar and Subramanian 2001; Peart et al. 2020), with a large standard deviation of 0.4 (5% and 95% quantiles of 4 × 10^−9^ and 4 × 10^−8^ bp^−1^ gen^−1^, respectively) to account for other rates found in the literature. We assumed a generation time of 10 years based on generation time estimates published by the IUCN (IUCN, 2017) as compiled in Peart et al. (2020) for a subset of the species considered here. We also added two calibration points in the phylogeny. One was at the origin of the *Arctocephalus* spp. clade, based on the age of the oldest *Arctocephalus* fossil record *(Arctocephalus* sp. nov. - Varswater Formation of South Africa), which constrained the origin of this group to a lower bound of 2.7 Ma (Avery and Klein 2011), since the incomplete and imperfect nature of the fossil records only provides evidence for the minimum age of a clade (Benton and Ayala 2003). The second was the date of the root, which we set as a normal prior with a mean of 20 Ma (± 3.0) in the divergence between Otariidae and Odobenidae (Yonezawa et al. 2009; Nyakatura and Bininda-Emonds 2012). We ran a Bayesian Markov Chain Monte Carlo (MCMC) of 500,000,000 steps sampled each 20,000 with a burn-in of 10%. To test underestimation of the internal branches due to possible undetected hybridizations (Leaché et al. 2014, Elworth et al. 2019), we also estimated a StarBEAST2 species tree using only the GFs of 50 kb whose ML tree topology was identical to our main species tree (see results) using the same parameters as above. We checked the MCMC runs with Tracer 1.7 (Rambaut and Drummond 2007).

As an additional estimation of divergence times, the species tree topology (recovered by ASTRAL-III and StarBEAST2) was used as input in the Bayesian species tree estimation of the BP&P program (Ziheng 2015; Flouri et al. 2018). We used the same 300 GFs of 50 kb applied in the initial StarBEAST analysis, and the following parameters: an MCMC chain of 2,000,000 replicates with burn-in of 200,000, a theta prior of 0.01 and a tau prior of 0.02. The theta prior specifies the inverse-gamma prior, the number of differences per kb, and the tau specifies the divergence time parameter for the root. For this analysis, the divergence times were calibrated based on the age of the root as above (Yonezawa et al. 2009; Nyakatura and Bininda-Emonds 2012). All trees were visualized and edited for clarity on FigTree 1.4.4 (Rambaut 2017) or Dendroscope 3 (Huson and Scornavacca 2012).

### Simulation of Genomic Fragments Trees from Assuming a Known Species Tree

To test if the high level of topological discordances between trees from the GFs could be explained by ILS alone, we simulated 10,000 GF trees under a multispecies coalescent framework implemented in the function sim.coaltree.sp in the R phylogenetic package Phybase (Liu and Yu 2010). As input for the simulations, we used our species tree as estimated by StarBEAST2, which besides the topology, also estimated the branch lengths and effective population sizes *(dmv* parameter in the StarBEAST2 species tree) for all internal and terminal branches. Note that both the estimation of the species tree by StarBEAST2 and the GF trees simulated allowed the occurrence of ILS. We then tabulated the frequency of the tree topologies and calculated the linear Pearson’s correlation between the simulated and empirical frequency distribution (following Wang et al. 2018).

### Mitochondrial Genome Phylogeny

We obtained the mitochondrial genomes of the fur seals and sea lions by mapping all reads with PALEOMIX 1.2.13.2, using the parameters reported above for the nuclear genomes, against a mitochondrial genome (mtDNA) available on GenBank (*A. townsendi* - NC008420). In order to validate the recovered mtDNAs, we assembled and aligned the generated sequences with those published on GenBank. After the alignment step, the mitochondrial control region was excluded. An mtDNA Bayesian phylogenetic tree was estimated with BEAST 2.5.2 package with the parameters: Yule Tree Model prior; GTR substitution model with four gamma categories (estimated with ModelTest-NG); and the Uncorrelated Lognormal Clock Model with lognormal distribution with a mean substitution rate of 2% site^−1^ million year^−1^ (Nabholz et al. 2007) and a standard deviation of 0.8.

### Introgression Between Species

Within the Dsuite package, we used the program Dtrios (Malinski 2019) and jackknife blocks to infer *D* statistics (also called ABBA-BABA test). This analysis compares the distribution of ancestral (A) and derived (B) sites in a four-taxa asymmetric phylogeny (((P1, P2), P3), O) with P1 to P3 being ingroups and O being the outgroup. Under the null hypothesis that P1 and P2 descend from an ancestor that diverged at an earlier time from the ancestral population of P3, derived alleles B should be found equally often in P1 and P2. Consequently, GF trees following allelic ABBA or BABA relationships should be equally likely for incompletely sorted ancestral polymorphism. Gene flow between P2 and P3 will lead to an excess of ABBA patterns reflected in a positive *D*-statistic, gene flow between P1 and P3 to a surplus of BABA patterns reflected in a negative *D*-statistic (Durand et al. 2011). In the Dsuite package, P1 and P2 are ordered so that nABBA >= nBABA and, consequently, is never negative. Statistical significance for a deviation of the *D*-statistic from zero was assessed by calculating Z-scores and their associated p-values by the standard block-jackknife procedure (Durand et al. 2011), using p-value < 0.05 as an indication for a possible signal of introgression. To take into account the multiple testing problem, the p-values were adjusted by the Bonferroni correction (Malinski 2019). The Dtrios program orders each trio of taxa by assuming that the correct tree is the one where the BBAA pattern is more common than the discordant ABBA and BABA patterns, which are assumed to be introgressed loci.

We also estimated the *f_3_* and*f_4_*-statistics (Patterson et al. 2012) in threepop and fourpop modules, respectively, of *TreeMix* package (Pickrell and Pritchard 2012; Harris and DeGiorgio 2016). The *f_3_*-statistics explicitly tests whether a taxon of interest *C* is the result of admixture between two other taxa *A* and *B* considering the product of allelic differentials between populations (c-a)(c-b): negative values suggest that allele frequencies c are intermediate at many positions, which is consistent with a history of admixture while positive values are not evidence against admixture. *F*_4_-statistics use unrooted four-population phylogenies to visualize shared genetic drift among taxa. For a *f_4_* ((A,B),(C,D)) topology without invoking admixture the allele frequency difference between A and B (a-b) and between C and D (c-d) should be unrelated and hence results in *f_4_* =((a-b)(c-d))= 0. A significantly positive *f_4_* implies gene flow between A and C, or B and D. Otherwise, a significantly negative value implies gene flow between A and D, or B and C. Significant *f_4_* values may also be interpreted as a rejection of the given topology (Peter 2016; Zhenge and Janke 2018). The significance of*f_3_* and*f_4_*-statistics is based on the Z-score and was calculated over 872 jackknife blocks of 50,000 SNPs. Significantly positive (Z > 3) and significantly negative (Z < −3) values, after Bonferroni correction, reject the null hypothesis. We plotted the distribution of*f_4_*-values with the function f4stats from admixturegraph (Leppälä et al. 2017), an R package.

We also used the newly developed QuIBL approach (Quantifying Introgression via Branch Lengths - Edelman et al. 2019), a statistical framework to estimate the number of discordant loci in a set of GF trees that reflect introgression events or ILS alone. Unlike *D* and *f*-statistics, a QuIBL analysis does not rely on topology imbalances but instead uses the distribution of internal branch lengths and calculates the likelihood that the discordant GF tree for a given region is due to introgression rather than ILS (Edelman et al. 2019). To distinguish whether the regions with local topologies discordant from the species tree were more likely to introgression or ILS, we used a Bayesian information criterion (BIC) test with a strict cutoff of dBIC <−10 to accept the ILS+introgression model as a better fit for the data, as suggested by the authors (Edelman et al. 2019). For this analysis, we used the GF trees generated from the partition of 50 kb. Since the analysis with 15 taxa (14 Otariidae plus *Odobenus*) is time-consuming, we used every other topology (5,454 GF trees) from the full dataset of 10,908 topologies. We used *O. rosmarus* as outgroup and our species tree estimated above with QuIBL default parameters as recommended by the authors (Edelman et al. 2019).

## Results

Fourteen sequenced otariid genomes, including nine fur seals and five sea lion species (Table 1), were mapped on the walrus genome with an average coverage of 27.79X (± 12.07X) (see Supplementary Table S2). The largest scaffold was 231.63 million bases (Mb), and the ten largest scaffolds summed-up to around 1.5 Gb, ~62% of the reference genome (2.4 Gb). Repetitive regions in the reference genome were masked in the consensus genomes (~40% of the reference genome), resulting in a high-quality non-repetitive alignment of ~1.1 Gb for further analyses. After filtering (removing masked regions, missing data, genomic fragments (GFs) with less than 50% of the original information, and those with the signal of intra-locus recombination), we obtained between 14,075 (with 10 kb) and 5,701 (with 200 kb) GFs a minimum of 100 kb apart from each other for the GF trees analyses (Table 2).

The Bayesian species tree (estimated with StarBEAST2 using 300 GF of 50 kb) (Fig. 1), the ASTRAL-III species trees (from thousands of ML trees using GFs ranging from 20kb to 200 kb) (Supplementary Fig. S1), the ML trees of eight of the ten largest scaffolds (Supplementary Fig. S2), the ML whole-genome tree (Supplementary Fig. S3), and the NJ tree (estimated using the genetic distances among the whole genomes, Supplementary Fig. S4) all resulted in the same tree topology with high support for most or all branches, hereafter named as the Otariidae species tree.

**Figure 1.**
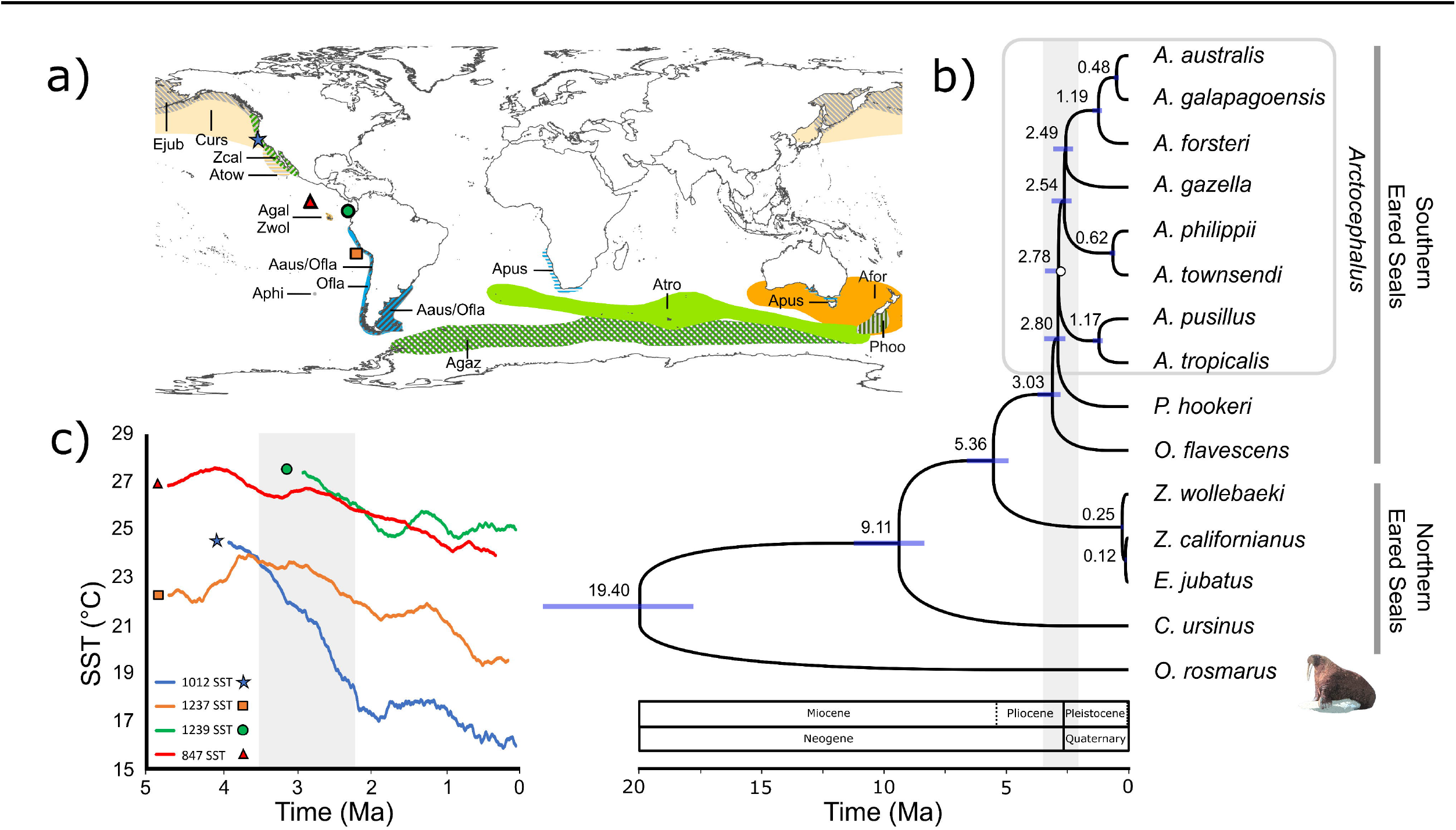
(a) Current distribution of fur seals and sea lions obtained from the IUCN Red List (IUCN 2020). *A. galapagoensis* (Agal) and *Z. wollebaeki* (Zwol) have very similar and small distributions and are represented with the same color and an arrow. *A. philippii* (Aphi) is endemic to the Juan Fernández Island, and its distribution is also represented with an arrow. The symbols represent the four sites of the past temperature data in “c.” (b) Time calibrated Bayesian species tree estimated with StarBEAST2 using 300 GFs of 50 kb. Blue bars represent the divergence time 95% confidence interval. The vertical gray bar represents the 95% confidence interval of the period of fast diversification of the southern clade. All nodes have the highest posterior density (HPD) = 1 except for the *Arctocephalus* node (HPD = 0.92), shown as an open circle in the phylogeny. (c) The Sea Surface Temperature (SST) temperature data from four sites in the Tropical and Subtropical Pacific eastern Pacific (Fedorov et al. 2013). The vertical gray bar represents the same time interval depicted in the species tree.

This species tree strongly supports the existence of a Southern Hemisphere clade (see Churchill et al. 2014), the monophyly of the genus *Arctocephalus* and its close relationship to *P. hookeri* and *O. flavescens*. The clade of *Z. californianus + E. jubatus* + *Z. wollebaeki*, Northern species with the southernmost range reaching the equator, was more distantly related to the Southern clade. *C. ursinus*, also a Northern hemispheric species, is sister to all other otariids (Fig. 1b). Surprisingly, *E. jubatus* and *Z. californianus* grouped as sister species, and *Z. wollebaeki* as sister to them. Within *Arctocephalus*, there were four main lineages: *A. pusillus + A. tropicalis; A. phillippii + A. townsendi; A. gazella* and the clade comprised of *A. forsteri + A. galapagoensis + A. australis* (Fig. 1b). Only three alternative topologies were found in these analyses, one in which *P. hookeri* and the *A. tropicalis + A. pusillus* clade switched position (ASTRAL-III with GFs 10 kb, Supplementary Fig. S1) and two in which *A. gazella* was found at two different positions within *Arctocephalus* (found in two ML scaffold trees) (Supplementary Fig. S2).

The phylogeny of the mitochondrial genomes (Supplementary Figs. S5 and S6) was similar to the species tree, with high posterior probabilities for most nodes and only two differences: (1) the switched position between *P. hookeri* and the *A. tropicalis + A. pusillus* clade, as in the ASTRAL species tree of 10 kb GFs, and 2) the sister relationship of *A. australis* with *A. forsteri*, instead of with *A. galapagoensis.* The time scales differ, the mtDNA phylogeny divergences were more recent, especially for *C. ursinus* and the northern clade, and the southern clade diversification would have started ~2 Ma and did not occur as rapidly as found in the nuclear genome species tree.

The species tree divergence times estimated with StarBEAST2 and BP&P were very similar (Fig. 1b and Supplementary Fig. S7). The divergence between walrus and the Otariidae was 19.4 Ma (95% confidence interval (CI) = 17.2 - 23.2 Ma), and within the Otariidae, *C. ursinus* diverged ~9.1 Ma (95% CI = 8.1 - 10.9 Ma) followed by the clade *Z. wollebaeki* + *Z. californianus* + *E. jubatus* at 5.4 Ma (95% CI = 4.8 - 6.4 Mya). After that, around the Pliocene to Pleistocene transition, six lineages diverged almost simultaneously (between ~3 and 2.5 Ma), originating in order: *O. flavescens, P. hookeri*, and the four main *Arctocephalus* lineages described above. Specifically, *Arctocephalus* diversification began ~2.8 Ma (95% CI = 2.5 - 3.3), the divergence times between *A. pusillus* + *A. tropicalis* and between *A. forsteri* + *A. australis + A. galapagoensis* were very similar at, ~1.2 Ma (95% CI = 1.0 - 1.4 and 1.0 - 1.5, respectively). The two groups that diverged more recently were *A. phillippii* + *A. townsendi* and *A. australis* + *A. galapagoensis*, at 0.6 and 0.5 Ma, respectively (95% CI = 0.5 - 0.7 and 0.4 - 0.6). The most recent divergence occurred between *Z. wollebaeki, Z. californianus*, and *E. jubatus* ~0.25 Ma (95% CI = 2.1 - 3.1). The main difference between StarBEAST2 and BP&P results was that in the latter, the divergence of *A. gazella* was almost simultaneous with that of *A. forsteri* (~1.2 Ma, Fig. 1 and Supplementary Figure S7).

Finally, we evaluated whether the almost simultaneous divergence time for the six lineages estimated in our species tree could be an artifact caused by the underestimation of divergence times (shortening of internal branch lengths) in methods that do not account for introgression (Elworth et al. 2019). In this context, we estimated a new StarBEAST2 calibrated species tree using only the 113 GFs of 50 kb whose ML tree topologies were identical to the species tree. The divergence times of this tree were almost identical to the 300 GFs species tree, in particular, the six nodes related to the explosive radiation (Supplementary Fig. S7), suggesting these very short divergence times were not artifacts of unaccounted hybridizations (see discussion).

### Genome Fragment Information Content and Phylogenetic Discordance

When the ML phylogeny of each GF was estimated separately, we found thousands of different topologies in each GF dataset (Supplementary Fig. S8); most occurred just once or a few times (that is, were estimated from one or a few GFs). The most frequent topology in the 10 kb dataset occurred in only 45 of the 14,012 GFs (i.e., in ~0.4% of the GF, Table 2). Although the frequency of the most common topology in each dataset proportionally increased with the size of the GFs (from 10 to 200kb), the most common topology only comprised ~3.8% of all topologies in even in the 200 kb data set (Table 2 and Supplementary Fig. S8). A very similar pattern was obtained for the alternative datasets with no filtering for recombination and without the 100 kb spacing between GFs (Supplementary Table S3). The Otariidae species tree was the most frequent topology in all datasets except for the 10 kb GF. Information content was high for all GF size classes. Considering the 50 kb GF as an example, the mean variable sites between the two closest species (*A. australis* and *A. galapagoensis)* was ~40, and the mean number of parsimony informative sites in the alignments ~200 (Supplementary Fig. S9) yielding enough variation to estimate reliable GF trees.

The IQ-TREE analysis recovered the same species tree topology with the highest branch support (100) for all nodes (Supplementary Fig. S10). However, the four nodes (nodes 16, 17, 19 and 20) that define the relationships between the six main lineages of the southern clade that arose almost simultaneously presented very low gene concordance (gCF: 19.9 - 32.2) and site concordance (sCF: 39.2 - 43.7) factors (Supplementary Fig. S9). In contrast, the other nodes showed much higher values for both statistics. Furthermore, less than 33% of the 10,806 GF trees supported the species tree for those nodes, but >76% supported the remaining nodes (Fig. 2).

**Figure 2.**
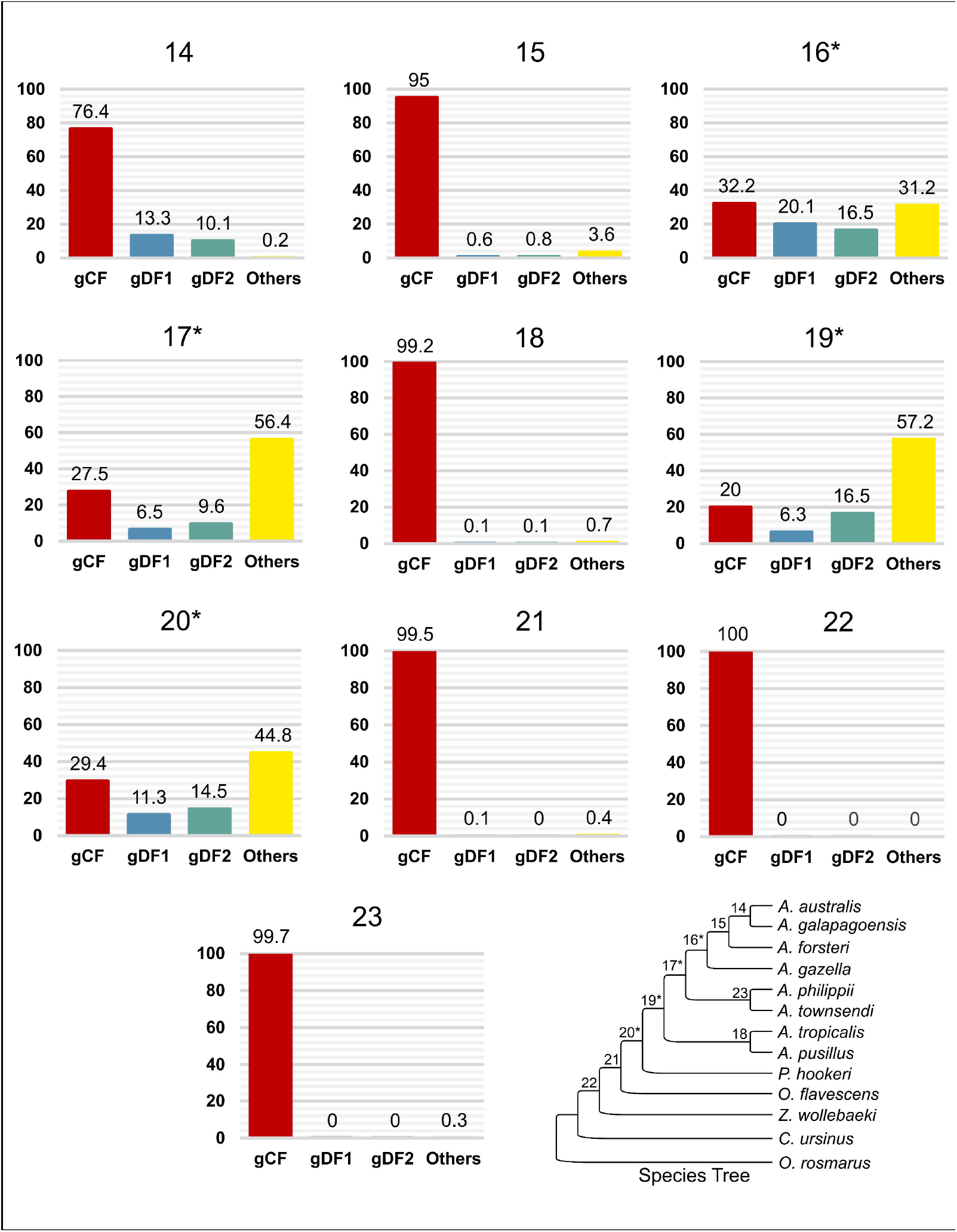
Gene concordance factor (gCF) for the nodes (14-23) that support (red bars) the species tree (bottom right) and the two most common alternative resolutions (gDF1 and gDF2, blue and green bars, respectively). The yellow bars are the relative frequencies of all other alternative resolutions. The nodes showing lower concordance factors (16, 17, 19 and 20) represent the lineages of fast radiation of eared seals in the Southern Hemisphere, with a remarkable number of alternative resolutions. These nodes are represented with an asterisk in the species tree.

### Hybridization vs. Incomplete Lineage Sorting

We used *D* (ABBA-BABA test), *f_3_*, and *f_4_*-statistics to investigate whether there is evidence of past events of hybridization (genomic introgression) between the species and if these events could explain the high level of topological discordance found in the southern clade. No evidence of introgression was found in the *f_3_*-statistics as all values were positive (not shown). For the ABBA-BABA test, a few *D*-statistics were significant *(p* < 0.05), but all turn out non-significant after Bonferroni correction (not shown). Otherwise, *f_4_*-statistics identified many significant (even after Bonferroni correction) sets of shared drift pathways between the species (Supplementary Figs. S11 and S12) that could be interpreted as signals of introgression or as supporting an alternative (to the species tree) phylogenetic relationship between the species considered (Peter 2016; Zhenge and Janke 2018). The strongest signals (*f_4_* > 0.01) supported introgression between *A. australis* and *A. forsteri.* The other significant *f_4_* values were very small (*f*_4_ < 0.001, Supplementary Figs. S11 and S12). Note that except for the tests that support introgression between *A. australis* and *A. forsteri* (Supplementary Fig. S11), all the other significant results could be interpreted as implying an alternative phylogenetic relationship between the six lineages that diverged almost simultaneously within the southern clade (see above and Fig. 1, and Supplementary Fig. S7), that is, where the internal branches were extremely small. Therefore, we next used two approaches to test if the high level of GF trees discordance (see above) and these *f_4_*-statistics results could be explained mostly by ILS, not introgression.

First, we used QuIBL analysis in distinguishing between ILS and introgression, which is thought to be more powerful than previous methods, such as *f_4_*-statistics or the *D* statistics (Edelman et al. 2019). This method uses the distribution of internal branch lengths to calculate the likelihood that a given genome fragment shows its GF topology due to introgression rather than ILS. QuIBL suggested that ILS could explain almost all significant *f_4_* results in those clades that emerged almost simultaneously (Supplementary Table S3). It identified only three significant events of hybridization with similar intensities: between *A. forsteri* and the ancestor of *A. australis* and *A. galapagoensis;* between the ancestor of *A. philippii* and *A. townsendi* and *A. gazella;* and between *Z. wollebaeki* and the ancestor of *Z. californianus* and *E. jubatus* (Fig. 3).

**Figure 3.**
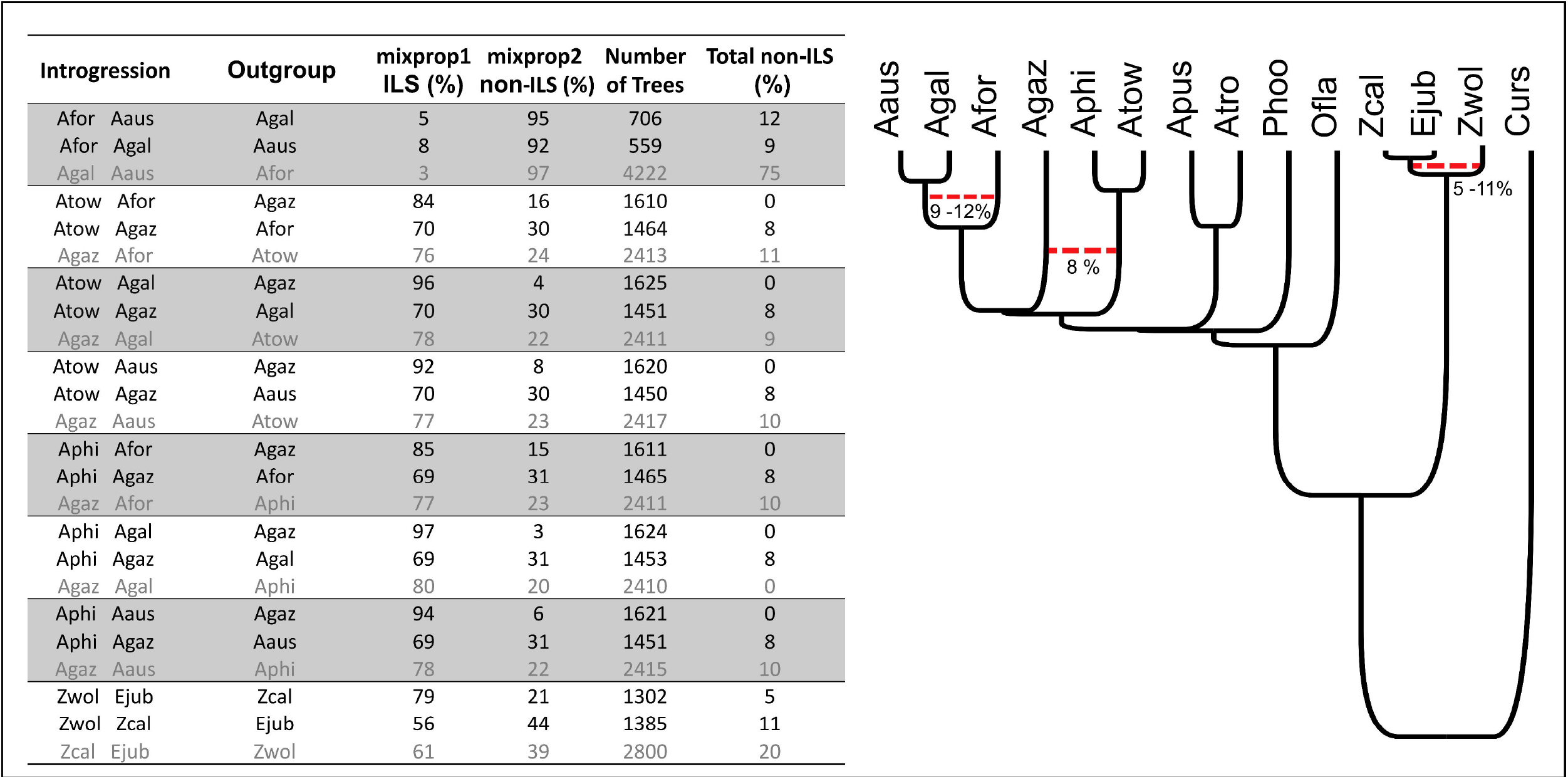
QuIBL significant results. The table at the left shows three alternative relationships for each species trio tested (the last of which, in grey type, is the species tree), the number of GF trees that significantly supported that relationship, and the proportion of these trees that could be explained by ILS or by alternative explanations (non-ILS, i.e., introgression or the phylogeny itself). Total non-ILS is the percentage of all GF trees that were introgressed between the pair of species that are not the outgroup (in the species tree this is explained by the phylogeny). To the right is the species tree depicting the introgression events supported.

Next, 10,000 GF trees were simulated using a multispecies coalescent model (that allows ILS but not introgression), whose parameters (the topology, divergence times, and effective population sizes) were those estimated by the StarBEAST2 species tree (Fig. 1).

The frequency distribution of the simulated GF tree topologies was similar to the observed distributions, in particular, for the 200 kb GFs partition (Supplementary Fig. S8). The simulation also presented the species tree as the most frequent topology (Table 2). The coefficient of correlation between the observed (50 and 200 kb data sets) and simulated distributions was 0.73 (Supplementary Fig. S13), which was high considering that the simulated topologies are true GF trees and are not affected by the uncertainties of the estimation as in the empirical dataset. These results suggest that the high level of GF tree discordance observed here could mostly be explained by ILS alone rather than by introgression events.

## Discussion

### Otariidae Phylogenomics

We present the first whole genome species tree of the Otariidae, which consistently recovered a phylogeny with high support using several different approaches. Our phylogeny also resolved uncertainties still prevalent to date in this group, such as the monophyly of *Arctocephalus*. The only species for which we could not obtain a genome sequence was *Neophoca cinerea.* Although some recent molecular studies support that *N. cinerea* is sister to *P. hookeri* (e.g., Yonezawa et al. 2009; Nyakatura and Bininda-Emonds 2012), some have suggested that it may be positioned elsewhere (Deméré and Berta 2003) including a sister position to all Otariidae except *C. ursinus* (Churchill et al. 2014). Future integration of the *Neophoca* genome to the data presented here thus constitutes a critical step to fully resolve the phylogeny of the Otariidae.

Our results strongly support *C. ursinus* as a sister species to all other Otariidae, which was also supported by other phylogenetic studies (Wynen et al. 2001; Árnason et al. 2006; Yonezawa et al. 2009; Nyakatura and Bininda-Emonds 2012; Berta and Churchill 2012; Churchill et al. 2014), thoroughly refuting the validity of the subfamilies Arctocephalinae and Otariinae. It is noteworthy that, considering our whole-genome phylogenies and other recent studies, *Callorhinus* diverged ca. 4 million years before the diversification of the rest of the family (Yonezawa et al. 2009; Boessenecker 2011; Nyakatura and Bininda-Emonds 2012; Berta et al. 2018).

Our study results offer robust support for the existence of the Northern Sea Lion clade, proposed by Churchill et al. (2014), consisting of *Zalophus* and *Eumetopias* (see Fig. 1). This Northern clade has been recovered in several previous studies (see Berta et al. 2018; Churchill et al. 2014; Berta and Churchill 2012; Yonezawa et al. 2009; Higdon et al. 2007; Árnason et al. 2006; Wynen et al. 2001), that also supported the monophyly of *Zalophus* (Wolf et al. 2007; Yonezawa et al. 2009; Churchill et al. 2014; Berta and Churchill 2012; Berta et al. 2018). Our analysis, however, recovered an unexpected but fully supported and close relationship between *E. jubatus* and *Z. californianus* with *Z. wollebaeki* as sister to them. It should be noted that most previous studies that support the monophyly of *Zalophus* used a few fragments of mtDNA (Yonezawa et al. 2009; Nyakatura and Bininda-Emonds 2012; Berta and Churchill 2012; Churchill et al. 2014, including the extinct Japanese sea lion - Wolf et al. 2007). Interestingly, the only study that used exclusively nuclear markers (AFLP data) found the same non-monophyletic relationship as found here (Dasmahapatra et al. 2009). If this relationship is supported by further studies, a taxonomy change would be necessary, such as synonymizing *Zalophus* with *Eumetopias*. The introgression we found between these species (see below) may help to explain their very recent divergence times and the very short internal branch separating *Z. wollebaeki* from the other two species (Fig. 1b). Together, our results motivate in-depth genomic-scale studies of this clade revisiting previous small-scale genetic studies of these species (Wolf et al. 2008; Schramm et al. 2009).

Previous authors had reached no consensus regarding the relationships between *Arctocephalus* spp., *P. hookeri, O. flavescens*, and *N. cinerea*, which has been called the southern clade (Churchill et al. 2014), except for a few subgroups within *Arctocephalus* (Berta et al. 2018). Our dated species trees showed that most of the speciation within the southern clade was almost simultaneous (Fig. 1b), which could explain the high number of different phylogenetic relationships found for this group to date. Our analyses based on genome-wide data provide strong support for this clade, with the South American sea lion (*O. flavescens*) being the first species to diverge around 3 Mya, followed by the New Zealand sea lion *(P. hookeri)* and a monophyletic *Arctocephalus*, both at ~2.8 Mya. The genomic data and the many different phylogenetic approaches we used support monophyly of *Arctocephalus* and did not support the use of *Arctophoca* as first suggested by Berta and Churchill (2012).

Within *Arctocephalus*, four main lineages originated in fast succession between ~2.8 and 2.5 Ma. The first to diverge was *A. pusillus* + *A. tropicalis*. This position within *Arctocephalus* for this clade was also found in several other recent studies (e.g., Berta et al. 2018), although some studies found it to have diverged before *Phocarctos* (e.g., Yonezawa et al. 2009). The divergence time between the two species was ~1.2 Ma. The next clade to diverge was the clade with *A. phillippii* + *A. townsendi*, with those species diverging more recently at ~0.6 Ma. These results were expected since they were reported as sister species in all previous molecular phylogenies (Yonezawa et al. 2009; Nyakatura and Bininda-Emonds 2012) and are morphologically very similar, with some authors still considering *A. townsendi* a subspecies of *A. phillippii* (e.g., Committee on Taxonomy 2020 following Berta and Churchill 2012). Considering the divergence time between these two species, which is similar or older than that between *A. australis* and *A. galapagoensis*, and their geographic isolation, we agree with most of the recent literature on their taxonomic status as full species (Repenning et al. 1971; Higdon et al. 2007; Yonezawa et al. 2009; Nyakatura et al. 2012; Aurioles-Gamboa 2015; Berta et al. 2018). The grouping of *A. gazella* with the *A. forsteri + A. australis + A. galapagoensis* clade was found in some recent studies (e.g., Yonezawa et al. 2009; Churchill et al. 2014), although *A. gazella* was found in a polytomy with other *Arctocephalus* species in most cases (e.g., Yonezawa et al. 2009; Berta et al. 2018) or in other positions. Finally, the *A. forsteri + A. australis + A. galapagoensis* clade was also highly expected, as these species were always closely related in previous phylogenetic studies and, until around 1970s, these three taxa were considered conspecific (Reppening et al. 1971, Brunner 2004). Nevertheless, there is a question over the species status of the New Zealand fur seal (*A. forsteri*), that was also considered a subspecies under *A. australis* (see Berta and Churchill 2012), and indeed, we found evidence of a low level of introgression between New Zealand fur seal and the South American fur seal (see below). However, we support *A. forsteri* as a full species based on the same arguments mentioned above for *A. phillippii* and *A. townsendi*, also considering that they diverged from *A. australis + A. galapagoensis* more than 1 Ma.

The placement of *P. hookeri* and *A. tropicalis* + *A. pusillus* were switched in the phylogeny obtained from mitochondrial genomes (Supplementary Figs. S5 and S6). This helps to explain why most of the previous molecular studies recovered a non-monophyletic *Arctocephalus*, as mtDNA constituted most or all the sequence data used in these studies (e.g., Árnason et al. 2006; Higdon et al. 2007; Wolf et al. 2007; Yonezawa et al. 2009; Churchill et al. 2014; Berta et al. 2018). The position of *A. forsteri* as sister species of *A. australis* in our mtDNA tree instead of *A. galapagoensis*, as in our nuclear genome species tree, is also observed in other mtDNA phylogenies (e.g., Wynen et al. 2001; Yonezawa et al. 2009). However, studies with mtDNA sequences from multiple individuals from *A. forsteri* and *A. australis* found several lineages in each species that are intermixed (e.g., Yonezawa et al. 2009). This complex picture, in particular the intermixing of lineages, could be explained by ILS since the grouping of *A. australis* and *A. forsteri* occurred in ~0.8% of the simulated trees. However, introgression could also have played a role in the history of this group, as the QulBL analysis and the *f_4_*-statistics (Fig. 3 and Supplementary Fig. S11) indicated significant admixture between *A. australis* and *A. forsteri* (see below). Intermixing of individuals of *A. australis* and *A. galapagoensis* has likewise been reported for mtDNA (Wolf et al. 2007), further emphasizing that the *A. forsteri/australis/galapagoensis* clade warrants further study.

### Divergence Times and Biogeographical Inferences

Our results agree with most previous divergence time estimates and fossil dating that supported a North Pacific origin for Otariidae and the split from Odobenidae at ~19 Ma, in the lower Miocene (Fig. 1 and Supplementary Fig. S7 - Demeré et al. 2003; Árnason et al. 2006; Yonezawa et al. 2009; Nyakatura and Bininda-Emonds 2012; Churchill et al. 2014; Berta et al. 2018), when early Odobenidae and Otariidae co-occurred (Boessenecker and Churchill 2015). The divergence of *Callorhinus* at ~9 Ma is also similar to most previous estimates (e.g., Yonezawa et al. 2009; Nyakatura and Bininda-Emonds 2012), although Berta et al. (2018) suggested a much older divergence at ~16 Ma. A comparison of our results with previous divergence times (Yonezawa et al. 2009; Nyakatura and Bininda-Emonds 2012; Berta et al. 2018) is not straightforward given the differences in topologies. Some significant points, however, can be made. First, no previous study detected the explosive radiation at the beginning of the diversification of the southern clade around the Pliocene-Pleistocene boundary. Second, our estimates of divergence between the northern (*C. ursinus, Zalophus* spp. and *E. jubatus*) and the southern clades (*O. flavescens, P. hookeri*, and *Arctocephalus* spp.) at ~5.3 Ma, and the initial diversification within the northern (~0.25 Ma) and the southern (3-2.5 Ma) clades are younger than most previous estimates (Yonezawa et al. 2009; Nyakatura and Bininda-Emonds 2014; Churchill et al. 2014; Berta et al. 2018). As an extreme example, Berta et al. (2018) estimated the divergence between *Otaria, Phocarctos*, and *Arctocephalus* at >6 Ma. On the other hand, Berta et al. (2018) estimated that the diversifications within *Arctocephalus* (except for the *A. pusillus* + *A. tropicalis* clade) are very recent, <1 Ma. It should be noted that, although we detected possible evidence for three introgression events with only a moderate extent (~10% of genomic introgression, Fig. 3 and below), we may have still underestimated some divergence times since the methods used here did not consider introgression. This may be the case for the very recent speciation times between the three species of the northern clade that could be underestimated due to past introgression events (Fig. 3).

Our phylogenomic results broadly agree with a scenario of a relatively recent trans-equatorial dispersal towards the Southern Hemisphere, likely along the Pacific coast of South America (see Yonezawa et al. 2009; Churchill et al. 2014, Berta et al. 2018). For a better understanding of this biogeographical history, we used data such as the age and phylogenetic position of fossils. In the Southern Hemisphere, most of the otariid fossils date back to the Pleistocene, with *Hydrarctos* known from sediments of the end of the Pliocene. *Hydrarctos* (Muizon 1978; Muizon and DeVries 1985; Avery and Klein 2011) is the oldest known otariid fossil from South America and comes from the Pisco Formation of Peru. However, its phylogenetic position is uncertain. It has been more consistently placed within the southern clade due to its morphological similarity with *Arctocephalus* (Muizon 1978), but was also positioned outside the southern clade as the sister taxon to all extant otariids (Churchill et al. 2014; Berta et al. 2018). Our divergence time estimates do not support the position of *Hydrarctos* inside the southern clade since the diversification of the latter group started at ~3 Mya, and the youngest date of the fossil is ~3.9 Ma (it may be as old as ~6.6 Ma, see Muizon 1978; Muizon and DeVries 1985; Churchill et al. 2014). Therefore, *Hydrarctos* is likely a sister clade to the southern clade or, assuming the oldest dates, may represent an independent (and extinct) trans-equatorial dispersal towards the west coast of South America that preceded the one that gave rise to the extant southern clade. *Arctocephalus* sp. nov., a fossil that belongs to the Varswater Formation of South Africa, was dated between ~2.7-5 Ma (Avery and Klein 2011), and we used its most recent date as the minimum age for *Arctocephalus* in our StarBEAST2 calibrated species tree. Our point estimate in the origin of *Arctocephalus* was ~2.8 Ma (95% CI ranging from 2.4 to 3.3 Ma, Fig. 1b), close to the minimum limit. The divergence times of the species tree reestimated using only the calibrated point at the root (20 Ma) with BP&P (Supplementary Fig. S7b) resulted in very similar values, therefore supporting our estimates of diversification of the southern clade dates between ~3 Mya and 2.5 Ma. However, the occurrence of *Arctocephalus* in South Africa at this time means that it had already been established in South America before its eastern dispersal to Africa.

Based on these results, dispersal to the Southern Hemisphere could have occurred anytime between ~5 Ma, the split of the southern clade from the northern *Zalophus* group, and ~3 Ma, the beginning of the explosive radiation within the southern clade. However, if *Hydrarctos* is considered a member of the southern clade with a minimum age of ~4 Ma, the southern dispersal could have occurred more than 1 myr before the burst of diversification of the extant species. At the moment, it is not practical to speculate as to the specific moment of the trans-equatorial dispersal within this large interval (between ~3 and 5 Ma). There are, however, environmental conditions within this timeframe that may have facilitated trans-equatorial dispersal such as lower sea temperatures in the equatorial zone (Churchill et al. 2014) and its concomitant higher ocean primary productivity. The period between the early Pliocene (~4 Ma) and the mid-Pliocene (~3.5 Ma) was characterized by warm temperatures (Fig. 1c, Fedorov et al. 2013) and low-productivity waters that likely impeded the trans-hemispheric dispersal at that time (O’Dea et al. 2012; Fedorov et al. 2013; Churchill et al. 2014). A trans-hemispheric dispersal more recent than estimated in previous studies (>5Ma, e.g., Yonezawa et al. 2009), and closer to the time of the southern clade diversification (~3 Ma), better explains the absence of otariids in the North Atlantic waters, since the total closure of the Central American Seaway finished ~3 Ma (O’Dea et al. 2012).

Conversely, the rapid diversification of the southern clade over the Southern Hemisphere, occurring during a relatively short time interval (between ~3.0-2.5 Ma), may be more firmly linked to major climatic events. Around 3 Ma, a sharp global cooling started (Fig. 1c), associated with the beginning of the Northern Hemisphere Glaciation (Fedorov et al. 2013; Marlow et al. 2000). The environmental changes caused by the concomitant global cooling during the Plio-Pleistocene transition and the total closure of the Panama Isthmus would have provided a suitable niche for otariids, driven by the increase of primary productivity in the Southern Pacific Ocean (O’Dea et al. 2012; Churchill et al. 2014). These changes may have opened the way for long-distance dispersal events within the Southern Hemisphere, with the establishment of new colonies and local adaptation to new niches, facilitating rapid speciation.

### Genealogical Discordances, ILS and Introgression

We found a high degree of topological discordance between the trees estimated from GFs along genomes (including the single locus mtDNA), with many topologies appearing in only one GF, even in the GFs of 200 kb (Table 2, Supplementary Fig. S8). These results explain the high degree of discordances among the phylogenies estimated by all previous studies and why it has been challenging to find a robust classification for the Otariidae based on a few genes. This high GF tree discordance could not be attributed to a lack of information in the GFs in general since most internal nodes, both older and recent, had high support values (e.g., sCF values in Supplementary Figs. S10). Most of the discordance was concentrated in the four nodes that gave rise to the six main lineages in the southern clade (Fig. 2 and Supplementary Fig. S10) and their extremely short internal branches. The explosive radiation (i.e., fast successive speciation events) at the origin of the southern clade was accompanied by short internal branches increasing the occurrence of ILS (see the small gCF values at theses nodes in Fig. 2) (Suh et al. 2015).

Topological discordance between genomic regions is not unusual and is being observed with increasing frequency in recent phylogenomic studies (e.g., Martin et al. 2013; Li et al. 2016; Pease et al. 2016; Árnason et al., 2018; Sun et al. 2020). The sources of topological discordances are mainly attributed to ILS, as suggested above, and hybridization (Bravo et al. 2019). Recent genomic studies have shown that introgression, mainly inferred with *D*-statistics or related statistics (e.g., ABBA-BABA, *f_3_*, and*f_4_)*, is widespread in the history of several groups (e.g., Pease et al. 2016; Figueiró et al. 2017; Masello et al. 2019). Here, we suggest that the several rapid successive events of speciation violated the assumptions of the bifurcating species tree and led to substantially false-positive signals of introgression in *f_4_* analysis, since the extremely short internal nodes do not allow this method to distinguish the true tree from alternative topologies (Durand et al. 2011; Eriksson and Manica 2012; Malinsky et al. 2018). In similar cases, we suggest replacing *f_4_-* statistics with methods that seem more robust to such artifacts, such as the recently developed QuIBL approach (Edelman et al. 2019). Instead, for a limited number of cases, prominent in the genus *Arctocephalus*, introgression seems to have contributed to the incongruencies. Considering present-day lack of firm reproductive barriers between several *Arctocephalus* species (Churchill et al. 2014), introgression during cladogenesis or shortly thereafter seems indeed plausible.

There have been recent implementations in the phylogenetic algorithms to infer divergence times that included hybridization in the multispecies coalescent models to estimate species networks (Zhang et al. 2017; Wen et al. 2018; Wen and Nakhleh 2018; Jones 2019). We have tried to recover a species network using four of these methods: the SpeciesNetwork (Zhang et al. 2018) and DENIM (Jones 2019), both implemented in StarBEAST2, and the MCMC_GT and MCMC_SEQ from PhyloNet (Wen et al. 2018). For these analyses, we used 100 GFs of 1 kb and 5 kb selected among those with more variation from the 300 GFs of 50 kb used in the StarBEAST2 analyses (Fig. 1b and Supplementary Fig. S7a). Unfortunately, either the Bayesian estimations did not stabilize even after long runs (>1 billion steps, DENIM), or we recovered several topologies that differed markedly from all other main topologies recovered with other methods (SpeciesNetwork and PhyloNet). Similar inconsistencies have also been found in other studies (e.g., Chen et al. 2019) and could be related to the high complexity of the models with a higher number of taxa (since these methods are mostly recommended for use with less than six taxa). The virtual polytomy between six lineages and the consequent high levels of ILS in our dataset may also have contributed to the non-stabilization of the analyses.

The relationships within the clade comprising *A. australis, A. galapagoensis*, and *A. forsteri* seem to reflect a complex scenario since we found evidence for both introgression and ILS between these species. Furthermore, previous studies based on mtDNA have found the absence of reciprocal monophyly among species (Wynen et al. 2001; Wolf et al. 2007; Yonezawa et al. 2009) and the possible existence of at least one cryptic species (King 1954; Repenning et al. 1971; Wynen et al. 2001; Oliveira et al. 2008; Yonezawa et al. 2009; Oliveira and Brownell 2014). Therefore, this clade needs a more in-depth study, analyzing samples from several populations.

## Conclusions

1. We used entire genome sequencing for 14 (missing *Neophoca cinerea*) of the extant 15 species of Otariidae to determine the phylogeny of this family and its bearing on its taxonomy and biogeographical history. Despite extreme topological discordance among GF trees, we found a fully supported species tree that agrees with the few well-accepted relationships found in previous studies.
2. Overall, the substantial degree of genealogical discordance was mostly accounted for by incomplete lineage sorting of ancestral genetic variation, though with a contribution of introgression in some clades.
3. A relatively recent trans-equatorial dispersal 3 to 2.5 Ma at the base of the southern clade rapidly diversified into six major lineages. *Otaria* was the first to diverge, followed by *Phocarctos* and then four lineages within *Arctocephalus*. This dispersal most likely occurred along the Pacific coast.
4. We found *Zalophus* and *Eumetopias*, from the northern clade, to be paraphyletic, with California sea lions (*Z. californianus)* more closely related to Steller sea lions (*Eumetopias jubatus*) than to Galapagos sea lions (*Z. wollebaeki*). However, the internal branch separating *Z. wollebaeki* from the other two species is very short, their divergence times are very recent, and we detected a signal of introgression between these two groups. It is necessary to conduct a more in-depth study of this clade, with genomic information from many individuals throughout the species’ distributions.
5. Quasi-simultaneous speciation within the southern clade led to extensive incomplete lineage sorting throughout the genomes, resulting in a high level of genealogical discordance, which explains the incongruence among and within prior phylogenetic studies of the family.
6. We suggest the use of recently developed methods of QuIBL when rapid successive events of speciation are detected to quantify events of genomic introgression. In similar cases, *f_4_*-statistics can violate the assumptions of the bifurcating species tree, leading to substantially false-positive signals of introgression.
7. Resolving a long-standing controversy, we found that the genus *Arctocephalus* is monophyletic, which makes the genus *Arctophoca* a junior synonym of *Arctocephalus.*

## Supplementary Material

- Raw sequencing data is available at NCBI BioProject PRJNA576431
- Data available from the Dryad Digital Repository: doi:10.5061/dryad.pzgmsbchw

## Acknowledgments

We are very thankful for the support of all people and institutions that contributed to this manuscript. We are also very grateful for the support of GEMARS, NOAA-SWFSC and Kelly Robertson which contributed with samples; Laboratório de Biologia Genômica e Molecular-PUCRS staff and students for support, general suggestions, and discussions; Jochen B. W. Wolf Group and Ludwig-Maximilians-Universität München which kindly hosted F.L. in his sandwich Ph.D.; two anonymous reviewers, and the associate editor and editor-in-chief for their helpful comment; and Nathaly Miranda who reviewed many versions of this manuscript.

---

**Table 1.** Whole genome shotgun sequences produced for this study (in bold) and obtained from GenBank digital repository.

**Table 2.** The ten most frequent topologies for the southern clade estimated with RAxML in each GF dataset and the absolute frequencies of occurrence in the different sets of windows sizes.

