## Supplementary Files and Tables for "Phylogenomic Discordance in the Eared Seals is best explained by Incomplete Lineage Sorting following Explosive Radiation in the Southern Hemisphere"

**Table S1.** Information on Otariidae samples used in this study.

| Species | Origin | Geographical coordinates | GenBank Access |
|---|---|---|---|
| *Arctocephalus australis* | Provincia de Chubut (Argentina) | 45°06'S, 65°24'W | SRX6989525 |
| *Arctocephalus galapagoensis* | Isla Santiago, Galapagos (Ecuador) | 0°14'S, 90°51'W | SRX7011050 |
| *Arctocephalus forsteri* | Otago Harbour (New Zealand) | 45°49'S, 170°39'E | SRX7011168 |
| *Arctocephalus philippii* | Juan Fernández Islands (Chile) | 33°46'S, 80°46'W | SRX7048039 |
| *Arctocephalus townsendi* | Malibu, California (USA) | 34°01'N, 118°01W | SRX7050505 |
| *Arctocephalus pusillus* | Kleinsee (South Africa) | 29°33'S, 17°00'E | SRX7050503 |
| *Arctocephalus tropicalis* | Coast of Rio Grande do Sul (Brazil) | 29°59'S, 50°08'W | SRX7050511 |
| *Phocarctos hookeri* | Otago Peninsula (New Zealand) | 45°50'S, 170°43'E | SRX7050502 |
| *Otaria flavescens* | Punta San Juan (Peru) | 15°22'S, 75°11'W | SRX7081214 |
| *Arctocephalus gazella* | N.A (Antarctic continent) | N.A. | SRX1338463-82 |
| *Eumetopias jubatus* | N.A. | N.A. | SRX3524690 |
| *Zalophus californianus* | San Miguel Island (USA) | 34°01'N, 120°26'W | SRX9590141 |
| *Zalophus wollebaeki* | Isla Mosquera, Galapagos (Ecuador) | 0°22'S, 90°35'W | SRX9590053 |
| *Callorhinus ursinus* | N.A. | N.A. | SRX4182256 |

**Table S2.** PALEOMIX statistics for each species mapped against the *O. rosmarus* genome (2.4 Gb genome).

| | Library | Coverage | Proportion Ref. Genome Mapped | Fraction of mapped reads | Fraction of unmapped reads | Fraction of PCR duplicates |
|---|---|---|---|---|---|---|
| *A. australis* | TruSeq Nano | 21.34 | 0.85 | 0.91 | 0.09 | 0.06 |
| *A. galapagoensis* | TruSeq Nano | 15.42 | 0.79 | 0.90 | 0.10 | 0.12 |
| *A. forsteri* | TruSeq PCR Free | 26.05 | 0.86 | 0.91 | 0.09 | 0.05 |
| *A. gazella* | GenBank | 32.44 | 0.83 | 0.89 | 0.11 | 0.07 |
| *A. philippii* | TruSeq PCR Free | 17.44 | 0.85 | 0.90 | 0.10 | 0.06 |
| *A. townsendi* | TruSeq PCR Free | 28.20 | 0.84 | 0.90 | 0.10 | 0.07 |
| *A. pusillus* | TruSeq PCR Free | 25.28 | 0.84 | 0.91 | 0.09 | 0.08 |

| | | | | | | |
|---|---|---|---|---|---|---|
| *A. tropicalis* | TruSeq PCR Free | 28.63 | 0.82 | 0.91 | 0.09 | 0.10 |
| *P. hookeri* | TruSeq PCR Free | 17.49 | 0.85 | 0.90 | 0.10 | 0.05 |
| *O. flavescens* | TruSeq PCR Free | 22.95 | 0.86 | 0.91 | 0.09 | 0.05 |
| *E. jubatus* | GenBank | 61.65 | 0.81 | 0.91 | 0.09 | 0.10 |
| *Z. californianus* | TruSeq PCR Free | 43.34 | 0.74 | 0.92 | 0.08 | 0.19 |
| *Z. wollebaeki* | GenBank | 21.15 | 0.66 | 0.91 | 0.09 | 0.28 |
| *C. ursinus* | GenBank | 27.67 | 0.70 | 0.84 | 0.16 | 0.17 |

**Table S3.** Absolute and relative frequencies of the top ten topologies of 50 kb. a) The same dataset used in the Table 2, without filtering for recombination. b) Contiguous 50 kb without filtering for recombination; and c) Contiguous 50 kb filtered for recombination. The most frequent topology (the species tree topology) is shown in bold.

a) Frequency of topologies in the original dataset of GFs of 50 kb (100 kb spaced) without filtering for recombination with 3Seq (10,806 GFs).

| Abs. Freq | % | Topology |
|---|---|---|
| **354** | **1.6%** | **((((((((((ARaus,ARgal),ARfor),ARgaz),(ARphi,ARtow)),(ARpus,ARtro)),PHhoo),OTfla)** |
| 274 | 1.3% | ((((((((((ARaus,ARgal),ARfor),ARgaz),(ARphi,ARtow)),PHhoo),(ARpus,ARtro)),OTfla) |
| 218 | 1.0% | (((((((((ARaus,ARgal),ARfor),(ARgaz,(ARphi,ARtow))),(ARpus,ARtro)),PHhoo),OTfla) |
| 210 | 1.0% | ((((((((((ARaus,ARgal),ARfor),ARgaz),(ARphi,ARtow)),(ARpus,ARtro)),OTfla),PHhoo) |
| 202 | 0.9% | (((((((((ARaus,ARgal),ARfor),ARgaz),(ARphi,ARtow)),PHhoo),((ARpus,ARtro),OTfla)) |
| 192 | 0.9% | ((((((((((ARaus,ARgal),ARfor),(ARphi,ARtow)),ARgaz),(ARpus,ARtro)),PHhoo),OTfla) |
| 182 | 0.8% | (((((((((ARaus,ARgal),ARfor),(ARgaz,(ARphi,ARtow))),PHhoo),(ARpus,ARtro)),OTfla) |
| 168 | 0.8% | (((((((((ARaus,ARgal),ARfor),ARgaz),(ARphi,ARtow)),(ARpus,ARtro)),(OTfla,PHhoo)) |
| 156 | 0.7% | (((((((((ARaus,ARgal),ARfor),(ARphi,ARtow)),(ARgaz,(ARpus,ARtro))),PHhoo),OTfla) |
| 152 | 0.7% | ((((((((((ARaus,ARgal),ARfor),ARgaz),(ARphi,ARtow)),PHhoo),OTfla),(ARpus,ARtro)) |

b) Frequency of topologies in the dataset of contiguous GFs of 50 kb (not spaced to avoid LD) without filtering for recombination with 3Seq (31,941 GFs).

| Abs. Freq | % | Topology |
|---|---|---|
| **251** | **0.8%** | **((((((((((Aaus,Agal),Afor),Agaz),(Aphi,Atow)),(Apus,Atro)),Phoo),Ofla),((Ejub,Zcal),Zwol)),Curs),Oros)** |
| 216 | 0.7% | ((((((((((Aaus,Agal),Afor),Agaz),(Aphi,Atow)),Phoo),(Apus,Atro)),Ofla),((Ejub,Zcal),Zwol)),Curs),Oros) |
| 183 | 0.6% | (((((((((Aaus,Agal),Afor),Agaz),(Aphi,Atow)),Phoo),((Apus,Atro),Ofla)),((Ejub,Zcal),Zwol)),Curs),Oros) |
| 145 | 0.5% | (((((((((Aaus,Agal),Afor),(Agaz,(Aphi,Atow))),Phoo),(Apus,Atro)),Ofla),((Ejub,Zcal),Zwol)),Curs),Oros) |
| 144 | 0.5% | (((((((((Aaus,Agal),Afor),Agaz),(Aphi,Atow)),(Apus,Atro)),(Ofla,Phoo)),((Ejub,Zcal),Zwol)),Curs),Oros) |
| 143 | 0.4% | (((((((((Aaus,Agal),Afor),(Agaz,(Aphi,Atow))),(Apus,Atro)),Phoo),Ofla),((Ejub,Zcal),Zwol)),Curs),Oros) |
| 129 | 0.4% | ((((((((((Aaus,Agal),Afor),Agaz),(Aphi,Atow)),(Apus,Atro)),Ofla),Phoo),((Ejub,Zcal),Zwol)),Curs),Oros) |
| 128 | 0.4% | ((((((((((Aaus,Agal),Afor),(Aphi,Atow)),Agaz),(Apus,Atro)),Phoo),Ofla),((Ejub,Zcal),Zwol)),Curs),Oros) |
| 122 | 0.4% | (((((((((Aaus,Agal),Afor),(Aphi,Atow)),(Agaz,(Apus,Atro))),Phoo),Ofla),((Ejub,Zcal),Zwol)),Curs),Oros) |
| 113 | 0.4% | ((((((((((Aaus,Agal),Afor),Agaz),(Aphi,Atow)),(Apus,Atro)),Phoo),Ofla),(Ejub,(Zcal,Zwol))),Curs),Oros) |

c) Frequency of topologies in the dataset of contiguous GFs of 50 kb (not spaced to avoid LD) filtered for recombination with 3Seq (18,228 GFs).

| Abs. Freq | % | Topology |
|---|---|---|
| **103** | **0.6%** | **((((((((((Aaus,Agal),Afor),Agaz),(Aphi,Atow)),(Apus,Atro)),Phoo),Ofla),((Ejub,Zcal),Zwol)),Curs),Oros)** |
| 91 | 0.5% | ((((((((((Aaus,Agal),Afor),Agaz),(Aphi,Atow)),Phoo),(Apus,Atro)),Ofla),((Ejub,Zcal),Zwol)),Curs),Oros) |

| | | |
|---|---|---|
| 79 | 0.4% | (((((((((Aaus,Agal),Afor),Agaz),(Aphi,Atow)),(Apus,Atro)),(Ofla,Phoo)),((Ejub,Zcal),Zwol)),Curs),Oros) |
| 74 | 0.4% | (((((((((Aaus,Agal),Afor),(Agaz,(Aphi,Atow))),(Apus,Atro)),Phoo),Ofla),((Ejub,Zcal),Zwol)),Curs),Oros) |
| 71 | 0.4% | (((((((((Aaus,Agal),Afor),Agaz),(Aphi,Atow)),Phoo),((Apus,Atro),Ofla)),((Ejub,Zcal),Zwol)),Curs),Oros) |
| 63 | 0.3% | ((((((((((Aaus,Agal),Afor),(Aphi,Atow)),Agaz),(Apus,Atro)),Phoo),Ofla),((Ejub,Zcal),Zwol)),Curs),Oros) |
| 61 | 0.3% | (((((((((Aaus,Agal),Afor),(Agaz,(Aphi,Atow))),Phoo),(Apus,Atro)),Ofla),((Ejub,Zcal),Zwol)),Curs),Oros) |
| 56 | 0.3% | ((((((((((Aaus,Agal),Afor),Agaz),(Aphi,Atow)),(Apus,Atro)),Phoo),Ofla),((Ejub,Zwol),Zcal)),Curs),Oros) |
| 55 | 0.3% | (((((((((Aaus,Agal),Afor),(Aphi,Atow)),Agaz),(Apus,Atro)),(Ofla,Phoo)),((Ejub,Zcal),Zwol)),Curs),Oros) |
| 53 | 0.3% | (((((((((Aaus,Agal),Afor),Agaz),(Apus,Atro)),((Aphi,Atow),Phoo)),Ofla),((Ejub,Zcal),Zwol)),Curs),Oros) |

**Table S4.** All QuIBL results. The table shows three alternative relationships for each species trio tested, the number of gene trees that supported that relationship, and the proportion of these trees that could be explained by ILS or by alternative explanations (non-ILS, i.e., introgression or the phylogeny itself). Total non-ILS is the percentage of all gene trees that were introgressed between the pair of species that are not the outgroup (in the species tree this is explained by the phylogeny).

| triplet | outgroup | C1 | C2 | mixprop1 | mixprop2 | lambda2Dist | lambda1Dist | BIC2Dist ILS + Introg | BIC1Dist ILS only | DBIC | count | % total non-ILS | % total ILS |
|---|---|---|---|---|---|---|---|---|---|---|---|---|---|
| **Apus_Atro_Ofla** | **Apus** | 0 | 6.37 | 0.90 | 0.10 | 0.0005 | 0.0006 | -124 | -125 | 0 | 10 | 0.0% | 0.0% |
| **Apus_Atro_Ofla** | **Atro** | 0 | 2.37 | 0.63 | 0.37 | 0.0005 | 0.0007 | -72 | -74 | 2 | 6 | 0.0% | 0.0% |
| **Apus_Atro_Ofla** | **Ofla** | 0 | 1.41 | 0.00 | 1.00 | 0.0005 | 0.0009 | -70017 | -65725 | -4292 | 5471 | 99.7% | 0.0% |
| **Apus_Atro_Atow** | **Apus** | 0 | 4.02 | 0.90 | 0.10 | 0.0004 | 0.0004 | -106 | -109 | 3 | 8 | 0.0% | 0.0% |
| **Apus_Atro_Atow** | **Atro** | 0 | 2.79 | 0.84 | 0.16 | 0.0005 | 0.0005 | -99 | -103 | 3 | 8 | 0.0% | 0.0% |
| **Apus_Atro_Atow** | **Atow** | 0 | 1.28 | 0.00 | 1.00 | 0.0005 | 0.0008 | -71032 | -66730 | -4302 | 5471 | 99.7% | 0.0% |
| **Apus_Atro_Aphi** | **Apus** | 0 | 3.70 | 0.93 | 0.07 | 0.0004 | 0.0004 | -117 | -121 | 4 | 9 | 0.0% | 0.0% |
| **Apus_Atro_Aphi** | **Atro** | 0 | 4.09 | 0.88 | 0.12 | 0.0003 | 0.0003 | -93 | -96 | 2 | 7 | 0.0% | 0.0% |
| **Apus_Atro_Aphi** | **Aphi** | 0 | 1.28 | 0.00 | 1.00 | 0.0005 | 0.0008 | -71014 | -66717 | -4297 | 5471 | 99.7% | 0.0% |
| **Apus_Atro_Agaz** | **Apus** | 0 | 3.82 | 0.93 | 0.07 | 0.0002 | 0.0002 | -140 | -144 | 4 | 10 | 0.0% | 0.0% |
| **Apus_Atro_Agaz** | **Atro** | 0 | 3.51 | 0.78 | 0.22 | 0.0005 | 0.0006 | -124 | -126 | 1 | 10 | 0.0% | 0.0% |
| **Apus_Atro_Agaz** | **Agaz** | 0 | 1.22 | 0.00 | 1.00 | 0.0005 | 0.0008 | -71283 | -67091 | -4192 | 5467 | 99.6% | 0.0% |
| **Apus_Atro_Afor** | **Apus** | 0 | 2.94 | 0.83 | 0.17 | 0.0005 | 0.0006 | -137 | -140 | 3 | 11 | 0.0% | 0.0% |
| **Apus_Atro_Afor** | **Atro** | 0 | 4.08 | 0.86 | 0.14 | 0.0003 | 0.0003 | -80 | -82 | 2 | 6 | 0.0% | 0.0% |
| **Apus_Atro_Afor** | **Afor** | 0 | 1.25 | 0.00 | 1.00 | 0.0005 | 0.0008 | -71162 | -66941 | -4221 | 5470 | 99.7% | 0.0% |
| **Apus_Atro_Agal** | **Apus** | 0 | 3.97 | 0.92 | 0.08 | 0.0004 | 0.0004 | -119 | -122 | 3 | 9 | 0.0% | 0.0% |
| **Apus_Atro_Agal** | **Atro** | 0 | 2.79 | 0.80 | 0.20 | 0.0005 | 0.0006 | -86 | -89 | 2 | 7 | 0.0% | 0.0% |
| **Apus_Atro_Agal** | **Agal** | 0 | 1.25 | 0.00 | 1.00 | 0.0005 | 0.0008 | -71150 | -66949 | -4202 | 5471 | 99.7% | 0.0% |
| **Apus_Atro_Aaus** | **Apus** | 0 | 3.70 | 0.94 | 0.06 | 0.0004 | 0.0004 | -130 | -134 | 4 | 10 | 0.0% | 0.0% |
| **Apus_Atro_Aaus** | **Atro** | 0 | 4.08 | 0.86 | 0.14 | 0.0003 | 0.0003 | -80 | -82 | 2 | 6 | 0.0% | 0.0% |
| **Apus_Atro_Aaus** | **Aaus** | 0 | 1.25 | 0.00 | 1.00 | 0.0005 | 0.0008 | -71174 | -66960 | -4214 | 5471 | 99.7% | 0.0% |
| **Apus_Atro_Phoo** | **Apus** | 0 | 9.16 | 0.86 | 0.14 | 0.0005 | 0.0010 | -84 | -81 | -3 | 7 | 0.0% | 0.0% |
| **Apus_Atro_Phoo** | **Atro** | 0 | 0.66 | 0.48 | 0.52 | 0.0004 | 0.0004 | -76 | -79 | 3 | 6 | 0.0% | 0.0% |
| **Apus_Atro_Phoo** | **Phoo** | 0 | 1.37 | 0.00 | 1.00 | 0.0005 | 0.0009 | -70494 | -66140 | -4354 | 5474 | 99.8% | 0.0% |
| **Apus_Atro_Zwol** | **Apus** | 0 | 0 | 0 | 0 | 0 | 0 | 0 | 0 | 0 | 0 | 0.0% | 0.0% |

| | | | | | | | | | | | | | |
|---|---|---|---|---|---|---|---|---|---|---|---|---|---|
| **Apus_Atro_Zwol** | **Atro** | 0 | 0 | 0 | 0 | 0 | 0 | 0 | 0 | 0 | 0 | 0.0% | 0.0% |
| **Apus_Atro_Zwol** | **Zwol** | 0 | 3.14 | 0.00 | 1.00 | 0.0007 | 0.0022 | -63582 | -56030 | -7552 | 5487 | 100.0% | 0.0% |
| **Apus_Atro_Zcal** | **Apus** | 0 | 0 | 0 | 0 | 0 | 0 | 0 | 0 | 0 | 0 | 0.0% | 0.0% |
| **Apus_Atro_Zcal** | **Atro** | 0 | 0 | 0 | 0 | 0 | 0 | 0 | 0 | 0 | 0 | 0.0% | 0.0% |
| **Apus_Atro_Zcal** | **Zcal** | 0 | 3.14 | 0.00 | 1.00 | 0.0007 | 0.0022 | -63582 | -56030 | -7552 | 5487 | 100.0% | 0.0% |
| **Apus_Atro_Ejub** | **Apus** | 0 | 0 | 0 | 0 | 0 | 0 | 0 | 0 | 0 | 0 | 0.0% | 0.0% |
| **Apus_Atro_Ejub** | **Atro** | 0 | 0 | 0 | 0 | 0 | 0 | 0 | 0 | 0 | 0 | 0.0% | 0.0% |
| **Apus_Atro_Ejub** | **Ejub** | 0 | 3.14 | 0.00 | 1.00 | 0.0007 | 0.0022 | -63582 | -56030 | -7552 | 5487 | 100.0% | 0.0% |
| **Apus_Atro_Curs** | **Apus** | 0 | 0 | 0 | 0 | 0 | 0 | 0 | 0 | 0 | 0 | 0.0% | 0.0% |
| **Apus_Atro_Curs** | **Atro** | 0 | 0 | 0 | 0 | 0 | 0 | 0 | 0 | 0 | 0 | 0.0% | 0.0% |
| **Apus_Atro_Curs** | **Curs** | 0 | 3.90 | 0.00 | 1.00 | 0.0010 | 0.0042 | -58117 | -48970 | -9147 | 5487 | 100.0% | 0.0% |
| **Apus_Atro_Oros** | **Apus** | 0 | 0 | 0 | 0 | 0 | 0 | 0 | 0 | 0 | 0 | 0.0% | 0.0% |
| **Apus_Atro_Oros** | **Atro** | 0 | 0 | 0 | 0 | 0 | 0 | 0 | 0 | 0 | 0 | 0.0% | 0.0% |
| **Apus_Atro_Oros** | **Oros** | 0 | 5.61 | 0.00 | 1.00 | 0.0022 | 0.0133 | -48611 | -36410 | -12201 | 5487 | 100.0% | 0.0% |
| **Apus_Ofla_Atow** | **Apus** | 0 | 5.87 | 1.00 | 0.00 | 0.0002 | 0.0002 | -23196 | -23210 | 14 | 1533 | 0.0% | 27.9% |
| **Apus_Ofla_Atow** | **Ofla** | 0 | 6.41 | 1.00 | 0.00 | 0.0002 | 0.0002 | -39858 | -39867 | 9 | 2711 | 0.0% | 0.0% |
| **Apus_Ofla_Atow** | **Atow** | 0 | 5.02 | 0.99 | 0.01 | 0.0002 | 0.0002 | -19171 | -19182 | 11 | 1243 | 0.0% | 22.5% |
| **Apus_Ofla_Aphi** | **Apus** | 0 | 6.15 | 1.00 | 0.00 | 0.0002 | 0.0002 | -23215 | -23229 | 14 | 1534 | 0.0% | 27.9% |
| **Apus_Ofla_Aphi** | **Ofla** | 0 | 6.07 | 1.00 | 0.00 | 0.0002 | 0.0002 | -39861 | -39874 | 13 | 2708 | 0.0% | 49.2% |
| **Apus_Ofla_Aphi** | **Aphi** | 0 | 5.01 | 0.99 | 0.01 | 0.0002 | 0.0002 | -19198 | -19209 | 11 | 1245 | 0.0% | 22.5% |
| **Apus_Ofla_Agaz** | **Apus** | 0 | 3.98 | 1.00 | 0.00 | 0.0002 | 0.0002 | -18103 | -18117 | 14 | 1186 | 0.0% | 21.5% |
| **Apus_Ofla_Agaz** | **Ofla** | 0 | 6.78 | 1.00 | 0.00 | 0.0003 | 0.0003 | -45025 | -45030 | 5 | 3100 | 0.0% | 0.0% |
| **Apus_Ofla_Agaz** | **Agaz** | 0 | 5.35 | 0.99 | 0.01 | 0.0002 | 0.0002 | -18574 | -18583 | 9 | 1201 | 0.0% | 0.0% |
| **Apus_Ofla_Afor** | **Apus** | 0 | 4.37 | 0.99 | 0.01 | 0.0002 | 0.0002 | -15797 | -15807 | 11 | 1015 | 0.0% | 18.3% |
| **Apus_Ofla_Afor** | **Ofla** | 0 | 6.47 | 1.00 | 0.00 | 0.0002 | 0.0002 | -45748 | -45764 | 16 | 3115 | 0.0% | 56.7% |
| **Apus_Ofla_Afor** | **Afor** | 0 | 5.08 | 1.00 | 0.00 | 0.0002 | 0.0002 | -21028 | -21041 | 13 | 1357 | 0.0% | 24.6% |
| **Apus_Ofla_Agal** | **Apus** | 0 | 4.33 | 0.99 | 0.01 | 0.0002 | 0.0002 | -15680 | -15690 | 10 | 1006 | 0.0% | 18.1% |
| **Apus_Ofla_Agal** | **Ofla** | 0 | 6.40 | 1.00 | 0.00 | 0.0002 | 0.0002 | -45904 | -45919 | 15 | 3125 | 0.0% | 56.9% |
| **Apus_Ofla_Agal** | **Agal** | 0 | 5.38 | 0.99 | 0.01 | 0.0002 | 0.0002 | -20988 | -20998 | 10 | 1356 | 0.0% | 0.0% |
| **Apus_Ofla_Aaus** | **Apus** | 0 | 4.33 | 0.99 | 0.01 | 0.0002 | 0.0002 | -15821 | -15831 | 10 | 1015 | 0.0% | 18.3% |
| **Apus_Ofla_Aaus** | **Ofla** | 0 | 6.44 | 1.00 | 0.00 | 0.0002 | 0.0002 | -45757 | -45773 | 16 | 3117 | 0.0% | 56.8% |
| **Apus_Ofla_Aaus** | **Aaus** | 0 | 5.05 | 0.99 | 0.01 | 0.0002 | 0.0002 | -20982 | -20995 | 13 | 1355 | 0.0% | 24.6% |

| | | | | | | | | | | | | | |
|---|---|---|---|---|---|---|---|---|---|---|---|---|---|
| **Apus_Ofla_Phoo** | **Apus** | 0 | 4.58 | 0.99 | 0.01 | 0.0002 | 0.0002 | -21951 | -21962 | 11 | 1437 | 0.0% | 25.9% |
| **Apus_Ofla_Phoo** | **Ofla** | 0 | 5.82 | 1.00 | 0.00 | 0.0002 | 0.0002 | -34184 | -34199 | 15 | 2287 | 0.0% | 41.6% |
| **Apus_Ofla_Phoo** | **Phoo** | 0 | 4.64 | 0.99 | 0.01 | 0.0002 | 0.0002 | -26930 | -26941 | 11 | 1763 | 0.0% | 31.9% |
| **Apus_Ofla_Zwol** | **Apus** | 0 | 0.60 | 0.04 | 0.96 | 0.0005 | 0.0007 | -161 | -162 | 1 | 13 | 0.0% | 0.0% |
| **Apus_Ofla_Zwol** | **Ofla** | 0 | 0.58 | 0.00 | 1.00 | 0.0002 | 0.0002 | -43 | -43 | 0 | 3 | 0.0% | 0.0% |
| **Apus_Ofla_Zwol** | **Zwol** | 0 | 2.43 | 0.00 | 1.00 | 0.0005 | 0.0013 | -67332 | -61508 | -5824 | 5471 | 99.7% | 0.0% |
| **Apus_Ofla_Zcal** | **Apus** | 0 | 0.60 | 0.04 | 0.96 | 0.0005 | 0.0007 | -161 | -162 | 1 | 13 | 0.0% | 0.0% |
| **Apus_Ofla_Zcal** | **Ofla** | 0 | 0.58 | 0.00 | 1.00 | 0.0002 | 0.0002 | -43 | -43 | 0 | 3 | 0.0% | 0.0% |
| **Apus_Ofla_Zcal** | **Zcal** | 0 | 2.43 | 0.00 | 1.00 | 0.0005 | 0.0013 | -67332 | -61508 | -5824 | 5471 | 99.7% | 0.0% |
| **Apus_Ofla_Ejub** | **Apus** | 0 | 0.60 | 0.04 | 0.96 | 0.0005 | 0.0007 | -161 | -162 | 1 | 13 | 0.0% | 0.0% |
| **Apus_Ofla_Ejub** | **Ofla** | 0 | 0.58 | 0.00 | 1.00 | 0.0002 | 0.0002 | -43 | -43 | 0 | 3 | 0.0% | 0.0% |
| **Apus_Ofla_Ejub** | **Ejub** | 0 | 2.43 | 0.00 | 1.00 | 0.0005 | 0.0013 | -67332 | -61508 | -5824 | 5471 | 99.7% | 0.0% |
| **Apus_Ofla_Curs** | **Apus** | 0 | 1.38 | 0.00 | 1.00 | 0.0008 | 0.0018 | -21 | -21 | -1 | 2 | 0.0% | 0.0% |
| **Apus_Ofla_Curs** | **Ofla** | 0 | 0 | 0 | 0 | 0 | 0 | 0 | 0 | 0 | 0 | 0.0% | 0.0% |
| **Apus_Ofla_Curs** | **Curs** | 0 | 3.79 | 0.00 | 1.00 | 0.0008 | 0.0033 | -60367 | -51572 | -8796 | 5485 | 100.0% | 0.0% |
| **Apus_Ofla_Oros** | **Apus** | 0 | 0 | 0 | 0 | 0 | 0 | 0 | 0 | 0 | 0 | 0.0% | 0.0% |
| **Apus_Ofla_Oros** | **Ofla** | 0 | 0 | 0 | 0 | 0 | 0 | 0 | 0 | 0 | 0 | 0.0% | 0.0% |
| **Apus_Ofla_Oros** | **Oros** | 0 | 5.74 | 0.00 | 1.00 | 0.0020 | 0.0124 | -49563 | -37180 | -12383 | 5487 | 100.0% | 0.0% |
| **Apus_Atow_Aphi** | **Apus** | 0 | 1.97 | 0.00 | 1.00 | 0.0005 | 0.0011 | -69335 | -63347 | -5988 | 5480 | 99.9% | 0.0% |
| **Apus_Atow_Aphi** | **Atow** | 0 | 0 | 0 | 0.00 | 0 | 0 | 0 | 0 | 0.00 | 1 | 0.0% | 0.0% |
| **Apus_Atow_Aphi** | **Aphi** | 0 | 8.17 | 0.83 | 0.17 | 0.0005 | 0.0013 | -66 | -66 | 0 | 6 | 0.0% | 0.0% |
| **Apus_Atow_Agaz** | **Apus** | 0 | 8.13 | 1.00 | 0.00 | 0.0003 | 0.0003 | -40215 | -40231 | 16 | 2770 | 0.0% | 50.5% |
| **Apus_Atow_Agaz** | **Atow** | 0 | 7.29 | 1.00 | 0.00 | 0.0002 | 0.0002 | -24939 | -24952 | 13 | 1664 | 0.0% | 30.3% |
| **Apus_Atow_Agaz** | **Agaz** | 0 | 5.23 | 0.99 | 0.01 | 0.0002 | 0.0002 | -16296 | -16308 | 12 | 1053 | 0.0% | 19.1% |
| **Apus_Atow_Afor** | **Apus** | 0 | 6.86 | 1.00 | 0.00 | 0.0002 | 0.0002 | -42942 | -42958 | 16 | 2945 | 0.0% | 53.7% |
| **Apus_Atow_Afor** | **Atow** | 0 | 5.28 | 1.00 | 0.00 | 0.0002 | 0.0002 | -22594 | -22608 | 14 | 1475 | 0.0% | 26.8% |
| **Apus_Atow_Afor** | **Afor** | 0 | 5.28 | 0.99 | 0.01 | 0.0002 | 0.0002 | -16585 | -16593 | 7 | 1067 | 0.0% | 0.0% |
| **Apus_Atow_Agal** | **Apus** | 0 | 8.57 | 1.00 | 0.00 | 0.0003 | 0.0003 | -42943 | -42959 | 16 | 2951 | 0.0% | 53.8% |
| **Apus_Atow_Agal** | **Atow** | 0 | 5.29 | 1.00 | 0.00 | 0.0002 | 0.0002 | -22598 | -22612 | 14 | 1475 | 0.0% | 26.8% |
| **Apus_Atow_Agal** | **Agal** | 0 | 5.24 | 0.99 | 0.01 | 0.0002 | 0.0002 | -16473 | -16482 | 9 | 1061 | 0.0% | 0.0% |
| **Apus_Atow_Aaus** | **Apus** | 0 | 8.14 | 1.00 | 0.00 | 0.0003 | 0.0003 | -43001 | -43016 | 16 | 2954 | 0.0% | 53.8% |
| **Apus_Atow_Aaus** | **Atow** | 0 | 5.26 | 1.00 | 0.00 | 0.0002 | 0.0002 | -22506 | -22520 | 14 | 1470 | 0.0% | 26.7% |

| | | | | | | | | | | | | | |
|---|---|---|---|---|---|---|---|---|---|---|---|---|---|
| **Apus_Atow_Aaus** | **Aaus** | 0 | 5.24 | 0.99 | 0.01 | 0.0002 | 0.0002 | -16538 | -16546 | 8 | 1063 | 0.0% | 0.0% |
| **Apus_Atow_Phoo** | **Apus** | 0 | 6.03 | 1.00 | 0.00 | 0.0002 | 0.0002 | -30166 | -30181 | 15 | 1990 | 0.0% | 36.3% |
| **Apus_Atow_Phoo** | **Atow** | 0 | 5.27 | 0.99 | 0.01 | 0.0001 | 0.0001 | -18622 | -18631 | 9 | 1190 | 0.0% | 0.0% |
| **Apus_Atow_Phoo** | **Phoo** | 0 | 4.73 | 0.99 | 0.01 | 0.0002 | 0.0002 | -34841 | -34850 | 10 | 2307 | 0.0% | 0.0% |
| **Apus_Atow_Zwol** | **Apus** | 0 | 1.09 | 0.50 | 0.50 | 0.0004 | 0.0004 | -64 | -67 | 2 | 5 | 0.0% | 0.0% |
| **Apus_Atow_Zwol** | **Atow** | 0 | 0.57 | 0.00 | 1.00 | 0.0004 | 0.0004 | -39 | -40 | 0 | 3 | 0.0% | 0.0% |
| **Apus_Atow_Zwol** | **Zwol** | 0 | 2.53 | 0.00 | 1.00 | 0.0005 | 0.0014 | -66885 | -60978 | -5908 | 5479 | 99.9% | 0.0% |
| **Apus_Atow_Zcal** | **Apus** | 0 | 1.09 | 0.50 | 0.50 | 0.0004 | 0.0004 | -64 | -67 | 2 | 5 | 0.0% | 0.0% |
| **Apus_Atow_Zcal** | **Atow** | 0 | 0.57 | 0.00 | 1.00 | 0.0004 | 0.0004 | -39 | -40 | 0 | 3 | 0.0% | 0.0% |
| **Apus_Atow_Zcal** | **Zcal** | 0 | 2.53 | 0.00 | 1.00 | 0.0005 | 0.0014 | -66885 | -60978 | -5908 | 5479 | 99.9% | 0.0% |
| **Apus_Atow_Ejub** | **Apus** | 0 | 1.09 | 0.50 | 0.50 | 0.0004 | 0.0004 | -64 | -67 | 2 | 5 | 0.0% | 0.0% |
| **Apus_Atow_Ejub** | **Atow** | 0 | 0.57 | 0.00 | 1.00 | 0.0004 | 0.0004 | -39 | -40 | 0 | 3 | 0.0% | 0.0% |
| **Apus_Atow_Ejub** | **Ejub** | 0 | 2.53 | 0.00 | 1.00 | 0.0005 | 0.0014 | -66885 | -60978 | -5908 | 5479 | 99.9% | 0.0% |
| **Apus_Atow_Curs** | **Apus** | 0 | 0 | 0 | 0 | 0 | 0 | 0 | 0 | 0 | 1 | 0.0% | 0.0% |
| **Apus_Atow_Curs** | **Atow** | 0 | 0 | 0 | 0 | 0 | 0 | 0 | 0 | 0 | 0 | 0.0% | 0.0% |
| **Apus_Atow_Curs** | **Curs** | 0 | 3.72 | 0.00 | 1.00 | 0.0009 | 0.0034 | -60050 | -51326 | -8724 | 5486 | 100.0% | 0.0% |
| **Apus_Atow_Oros** | **Apus** | 0 | 0 | 0 | 0 | 0 | 0 | 0 | 0 | 0 | 0 | 0.0% | 0.0% |
| **Apus_Atow_Oros** | **Atow** | 0 | 0 | 0 | 0 | 0 | 0 | 0 | 0 | 0 | 0 | 0.0% | 0.0% |
| **Apus_Atow_Oros** | **Oros** | 0 | 5.67 | 0.00 | 1.00 | 0.0021 | 0.0125 | -49406 | -37110 | -12296 | 5487 | 100.0% | 0.0% |
| **Apus_Aphi_Agaz** | **Apus** | 0 | 8.12 | 1.00 | 0.00 | 0.0003 | 0.0003 | -40193 | -40208 | 16 | 2769 | 0.0% | 50.5% |
| **Apus_Aphi_Agaz** | **Aphi** | 0 | 7.44 | 1.00 | 0.00 | 0.0002 | 0.0002 | -24968 | -24979 | 11 | 1667 | 0.0% | 30.3% |
| **Apus_Aphi_Agaz** | **Agaz** | 0 | 5.27 | 0.99 | 0.01 | 0.0002 | 0.0002 | -16326 | -16336 | 11 | 1051 | 0.0% | 19.0% |
| **Apus_Aphi_Afor** | **Apus** | 0 | 6.84 | 1.00 | 0.00 | 0.0003 | 0.0003 | -42938 | -42954 | 16 | 2946 | 0.0% | 53.7% |
| **Apus_Aphi_Afor** | **Aphi** | 0 | 5.22 | 1.00 | 0.00 | 0.0002 | 0.0002 | -22589 | -22603 | 14 | 1477 | 0.0% | 26.9% |
| **Apus_Aphi_Afor** | **Afor** | 0 | 4.89 | 0.99 | 0.01 | 0.0002 | 0.0002 | -16581 | -16591 | 10 | 1064 | 0.0% | 19.2% |
| **Apus_Aphi_Agal** | **Apus** | 0 | 8.55 | 1.00 | 0.00 | 0.0003 | 0.0003 | -42933 | -42949 | 16 | 2951 | 0.0% | 53.8% |
| **Apus_Aphi_Agal** | **Aphi** | 0 | 5.24 | 1.00 | 0.00 | 0.0002 | 0.0002 | -22589 | -22603 | 14 | 1476 | 0.0% | 26.8% |
| **Apus_Aphi_Agal** | **Agal** | 0 | 5.31 | 0.99 | 0.01 | 0.0002 | 0.0002 | -16522 | -16530 | 8 | 1060 | 0.0% | 0.0% |
| **Apus_Aphi_Aaus** | **Apus** | 0 | 8.11 | 1.00 | 0.00 | 0.0003 | 0.0003 | -42996 | -43012 | 16 | 2955 | 0.0% | 53.9% |
| **Apus_Aphi_Aaus** | **Aphi** | 0 | 5.19 | 1.00 | 0.00 | 0.0002 | 0.0002 | -22486 | -22500 | 14 | 1472 | 0.0% | 26.8% |
| **Apus_Aphi_Aaus** | **Aaus** | 0 | 4.88 | 0.99 | 0.01 | 0.0002 | 0.0002 | -16518 | -16528 | 10 | 1060 | 0.0% | 0.0% |
| **Apus_Aphi_Phoo** | **Apus** | 0 | 6.03 | 1.00 | 0.00 | 0.0002 | 0.0002 | -30178 | -30193 | 15 | 1991 | 0.0% | 36.3% |

| | | | | | | | | | | | | | |
|---|---|---|---|---|---|---|---|---|---|---|---|---|---|
| **Apus_Aphi_Phoo** | **Aphi** | 0 | 6.15 | 0.99 | 0.01 | 0.0002 | 0.0002 | -18627 | -18627 | 0 | 1194 | 0.0% | 0.0% |
| **Apus_Aphi_Phoo** | **Phoo** | 0 | 4.74 | 0.99 | 0.01 | 0.0002 | 0.0002 | -34774 | -34784 | 10 | 2302 | 0.0% | 0.0% |
| **Apus_Aphi_Zwol** | **Apus** | 0 | 1.09 | 0.50 | 0.50 | 0.0004 | 0.0004 | -64 | -67 | 2 | 5 | 0.0% | 0.0% |
| **Apus_Aphi_Zwol** | **Aphi** | 0 | 0.57 | 0.00 | 1.00 | 0.0004 | 0.0004 | -39 | -40 | 0 | 3 | 0.0% | 0.0% |
| **Apus_Aphi_Zwol** | **Zwol** | 0 | 2.56 | 0.00 | 1.00 | 0.0005 | 0.0014 | -66909 | -60988 | -5920 | 5479 | 99.9% | 0.0% |
| **Apus_Aphi_Zcal** | **Apus** | 0 | 1.09 | 0.50 | 0.50 | 0.0004 | 0.0004 | -64 | -67 | 2 | 5 | 0.0% | 0.0% |
| **Apus_Aphi_Zcal** | **Aphi** | 0 | 0.57 | 0.00 | 1.00 | 0.0004 | 0.0004 | -39 | -40 | 0 | 3 | 0.0% | 0.0% |
| **Apus_Aphi_Zcal** | **Zcal** | 0 | 2.56 | 0.00 | 1.00 | 0.0005 | 0.0014 | -66909 | -60988 | -5920 | 5479 | 99.9% | 0.0% |
| **Apus_Aphi_Ejub** | **Apus** | 0 | 1.09 | 0.50 | 0.50 | 0.0004 | 0.0004 | -64 | -67 | 2 | 5 | 0.0% | 0.0% |
| **Apus_Aphi_Ejub** | **Aphi** | 0 | 0.57 | 0.00 | 1.00 | 0.0004 | 0.0004 | -39 | -40 | 0 | 3 | 0.0% | 0.0% |
| **Apus_Aphi_Ejub** | **Ejub** | 0 | 2.56 | 0.00 | 1.00 | 0.0005 | 0.0014 | -66909 | -60988 | -5920 | 5479 | 99.9% | 0.0% |
| **Apus_Aphi_Curs** | **Apus** | 0 | 0 | 0 | 0 | 0 | 0 | 0 | 0 | 0 | 1 | 0.0% | 0.0% |
| **Apus_Aphi_Curs** | **Aphi** | 0 | 0 | 0 | 0 | 0 | 0 | 0 | 0 | 0 | 0 | 0.0% | 0.0% |
| **Apus_Aphi_Curs** | **Curs** | 0 | 3.73 | 0.00 | 1.00 | 0.0009 | 0.0034 | -60063 | -51330 | -8733 | 5486 | 100.0% | 0.0% |
| **Apus_Aphi_Oros** | **Apus** | 0 | 0 | 0 | 0 | 0 | 0 | 0 | 0 | 0 | 0 | 0.0% | 0.0% |
| **Apus_Aphi_Oros** | **Aphi** | 0 | 0 | 0 | 0 | 0 | 0 | 0 | 0 | 0 | 0 | 0.0% | 0.0% |
| **Apus_Aphi_Oros** | **Oros** | 0 | 5.68 | 0.00 | 1.00 | 0.0021 | 0.0125 | -49412 | -37111 | -12300 | 5487 | 100.0% | 0.0% |
| **Apus_Agaz_Afor** | **Apus** | 0 | 4.33 | 1.00 | 0.00 | 0.0003 | 0.0003 | -45709 | -45726 | 18 | 3196 | 0.0% | 58.2% |
| **Apus_Agaz_Afor** | **Agaz** | 0 | 4.16 | 0.99 | 0.01 | 0.0002 | 0.0002 | -15545 | -15559 | 14 | 1018 | 0.0% | 18.5% |
| **Apus_Agaz_Afor** | **Afor** | 0 | 7.68 | 1.00 | 0.00 | 0.0002 | 0.0002 | -19070 | -19072 | 2 | 1273 | 0.0% | 0.0% |
| **Apus_Agaz_Agal** | **Apus** | 0 | 4.91 | 1.00 | 0.00 | 0.0003 | 0.0003 | -45748 | -45766 | 18 | 3201 | 0.0% | 58.3% |
| **Apus_Agaz_Agal** | **Agaz** | 0 | 4.08 | 0.99 | 0.01 | 0.0002 | 0.0002 | -15690 | -15702 | 12 | 1023 | 0.0% | 18.5% |
| **Apus_Agaz_Agal** | **Agal** | 0 | 7.50 | 1.00 | 0.00 | 0.0002 | 0.0002 | -18941 | -18944 | 3 | 1263 | 0.0% | 0.0% |
| **Apus_Agaz_Aaus** | **Apus** | 0 | 4.31 | 1.00 | 0.00 | 0.0003 | 0.0003 | -45746 | -45764 | 18 | 3201 | 0.0% | 58.3% |
| **Apus_Agaz_Aaus** | **Agaz** | 0 | 4.19 | 1.00 | 0.00 | 0.0002 | 0.0002 | -15568 | -15582 | 14 | 1019 | 0.0% | 18.5% |
| **Apus_Agaz_Aaus** | **Aaus** | 0 | 7.71 | 1.00 | 0.00 | 0.0002 | 0.0002 | -18993 | -18995 | 2 | 1267 | 0.0% | 0.0% |
| **Apus_Agaz_Phoo** | **Apus** | 0 | 6.88 | 1.00 | 0.00 | 0.0002 | 0.0002 | -30427 | -30436 | 9 | 2023 | 0.0% | 0.0% |
| **Apus_Agaz_Phoo** | **Agaz** | 0 | 6.46 | 0.99 | 0.01 | 0.0001 | 0.0001 | -14562 | -14564 | 2 | 930 | 0.0% | 0.0% |
| **Apus_Agaz_Phoo** | **Phoo** | 0 | 5.75 | 1.00 | 0.00 | 0.0002 | 0.0002 | -37415 | -37429 | 14 | 2534 | 0.0% | 46.1% |
| **Apus_Agaz_Zwol** | **Apus** | 0 | 0.94 | 0.05 | 0.95 | 0.0005 | 0.0005 | -50 | -51 | 1 | 4 | 0.0% | 0.0% |
| **Apus_Agaz_Zwol** | **Agaz** | 0 | 1.02 | 0.59 | 0.41 | 0.0003 | 0.0003 | -52 | -55 | 2 | 4 | 0.0% | 0.0% |
| **Apus_Agaz_Zwol** | **Zwol** | 0 | 2.49 | 0.00 | 1.00 | 0.0005 | 0.0014 | -66583 | -60737 | -5846 | 5479 | 99.9% | 0.0% |

| | | | | | | | | | | | | | |
|---|---|---|---|---|---|---|---|---|---|---|---|---|---|
| **Apus_Agaz_Zcal** | **Apus** | 0 | 0.94 | 0.05 | 0.95 | 0.0005 | 0.0005 | -50 | -51 | 1 | 4 | 0.0% | 0.0% |
| **Apus_Agaz_Zcal** | **Agaz** | 0 | 1.02 | 0.59 | 0.41 | 0.0003 | 0.0003 | -52 | -55 | 2 | 4 | 0.0% | 0.0% |
| **Apus_Agaz_Zcal** | **Zcal** | 0 | 2.49 | 0.00 | 1.00 | 0.0005 | 0.0014 | -66583 | -60737 | -5846 | 5479 | 99.9% | 0.0% |
| **Apus_Agaz_Ejub** | **Apus** | 0 | 0.94 | 0.05 | 0.95 | 0.0005 | 0.0005 | -50 | -51 | 1 | 4 | 0.0% | 0.0% |
| **Apus_Agaz_Ejub** | **Agaz** | 0 | 1.02 | 0.59 | 0.41 | 0.0003 | 0.0003 | -52 | -55 | 2 | 4 | 0.0% | 0.0% |
| **Apus_Agaz_Ejub** | **Ejub** | 0 | 2.49 | 0.00 | 1.00 | 0.0005 | 0.0014 | -66583 | -60737 | -5846 | 5479 | 99.9% | 0.0% |
| **Apus_Agaz_Curs** | **Apus** | 0 | 0 | 0 | 0 | 0 | 0 | 0 | 0 | 0 | 1 | 0.0% | 0.0% |
| **Apus_Agaz_Curs** | **Agaz** | 0 | 0 | 0 | 0 | 0 | 0 | 0 | 0 | 0 | 1 | 0.0% | 0.0% |
| **Apus_Agaz_Curs** | **Curs** | 0 | 3.67 | 0.00 | 1.00 | 0.0009 | 0.0035 | -59904 | -51214 | -8690 | 5485 | 100.0% | 0.0% |
| **Apus_Agaz_Oros** | **Apus** | 0 | 0 | 0 | 0 | 0 | 0 | 0 | 0 | 0 | 0 | 0.0% | 0.0% |
| **Apus_Agaz_Oros** | **Agaz** | 0 | 0 | 0 | 0 | 0 | 0 | 0 | 0 | 0 | 0 | 0.0% | 0.0% |
| **Apus_Agaz_Oros** | **Oros** | 0 | 5.66 | 0.00 | 1.00 | 0.0021 | 0.0125 | -49353 | -37083 | -12271 | 5487 | 100.0% | 0.0% |
| **Apus_Afor_Agal** | **Apus** | 0 | 1.18 | 0.01 | 0.99 | 0.0005 | 0.0008 | -69213 | -66096 | -3117 | 5383 | 96.9% | 0.0% |
| **Apus_Afor_Agal** | **Afor** | 0 | 8.30 | 0.98 | 0.02 | 0.0002 | 0.0002 | -832 | -834 | 3 | 57 | 0.0% | 0.0% |
| **Apus_Afor_Agal** | **Agal** | 0 | 11.28 | 0.98 | 0.02 | 0.0003 | 0.0003 | -668 | -665 | -4 | 47 | 0.0% | 0.0% |
| **Apus_Afor_Aaus** | **Apus** | 0 | 1.18 | 0.00 | 1.00 | 0.0005 | 0.0008 | -69285 | -66082 | -3203 | 5393 | 98.0% | 0.0% |
| **Apus_Afor_Aaus** | **Afor** | 0 | 4.88 | 0.97 | 0.03 | 0.0002 | 0.0002 | -722 | -728 | 6 | 49 | 0.0% | 0.0% |
| **Apus_Afor_Aaus** | **Aaus** | 0 | 2.62 | 0.98 | 0.02 | 0.0002 | 0.0002 | -672 | -680 | 8 | 45 | 0.0% | 0.0% |
| **Apus_Afor_Phoo** | **Apus** | 0 | 5.54 | 1.00 | 0.00 | 0.0002 | 0.0002 | -31628 | -31642 | 14 | 2095 | 0.0% | 38.1% |
| **Apus_Afor_Phoo** | **Afor** | 0 | 4.17 | 0.99 | 0.01 | 0.0001 | 0.0001 | -15226 | -15237 | 11 | 957 | 0.0% | 17.2% |
| **Apus_Afor_Phoo** | **Phoo** | 0 | 4.79 | 0.99 | 0.01 | 0.0002 | 0.0002 | -36632 | -36644 | 12 | 2435 | 0.0% | 44.1% |
| **Apus_Afor_Zwol** | **Apus** | 0 | 0.94 | 0.05 | 0.95 | 0.0005 | 0.0005 | -50 | -51 | 1 | 4 | 0.0% | 0.0% |
| **Apus_Afor_Zwol** | **Afor** | 0 | 0.64 | 0.00 | 1.00 | 0.0004 | 0.0004 | -54 | -54 | 0 | 4 | 0.0% | 0.0% |
| **Apus_Afor_Zwol** | **Zwol** | 0 | 2.59 | 0.00 | 1.00 | 0.0005 | 0.0014 | -66961 | -60851 | -6111 | 5479 | 99.9% | 0.0% |
| **Apus_Afor_Zcal** | **Apus** | 0 | 0.94 | 0.05 | 0.95 | 0.0005 | 0.0005 | -50 | -51 | 1 | 4 | 0.0% | 0.0% |
| **Apus_Afor_Zcal** | **Afor** | 0 | 0.64 | 0.00 | 1.00 | 0.0004 | 0.0004 | -54 | -54 | 0 | 4 | 0.0% | 0.0% |
| **Apus_Afor_Zcal** | **Zcal** | 0 | 2.59 | 0.00 | 1.00 | 0.0005 | 0.0014 | -66961 | -60851 | -6111 | 5479 | 99.9% | 0.0% |
| **Apus_Afor_Ejub** | **Apus** | 0 | 0.94 | 0.05 | 0.95 | 0.0005 | 0.0005 | -50 | -51 | 1 | 4 | 0.0% | 0.0% |
| **Apus_Afor_Ejub** | **Afor** | 0 | 0.64 | 0.00 | 1.00 | 0.0004 | 0.0004 | -54 | -54 | 0 | 4 | 0.0% | 0.0% |
| **Apus_Afor_Ejub** | **Ejub** | 0 | 2.59 | 0.00 | 1.00 | 0.0005 | 0.0014 | -66961 | -60851 | -6111 | 5479 | 99.9% | 0.0% |
| **Apus_Afor_Curs** | **Apus** | 0 | 13.37 | 0.50 | 0.50 | 0.0005 | 0.0036 | -22 | -18 | -4 | 2 | 0.0% | 0.0% |
| **Apus_Afor_Curs** | **Afor** | 0 | 0 | 0 | 0 | 0 | 0 | 0 | 0 | 0 | 0 | 0.0% | 0.0% |

| | | | | | | | | | | | | | |
|---|---|---|---|---|---|---|---|---|---|---|---|---|---|
| **Apus_Afor_Curs** | **Curs** | 0 | 3.79 | 0.00 | 1.00 | 0.0008 | 0.0034 | -60101 | -51262 | -8839 | 5485 | 100.0% | 0.0% |
| **Apus_Afor_Oros** | **Apus** | 0 | 0 | 0 | 0 | 0 | 0 | 0 | 0 | 0 | 0 | 0.0% | 0.0% |
| **Apus_Afor_Oros** | **Afor** | 0 | 0 | 0 | 0 | 0 | 0 | 0 | 0 | 0 | 0 | 0.0% | 0.0% |
| **Apus_Afor_Oros** | **Oros** | 0 | 5.70 | 0.00 | 1.00 | 0.0021 | 0.0125 | -49423 | -37096 | -12328 | 5487 | 100.0% | 0.0% |
| **Apus_Agal_Aaus** | **Apus** | 0 | 1.88 | 0.01 | 0.99 | 0.0005 | 0.0011 | -67442 | -63223 | -4218 | 5437 | 98.4% | 0.0% |
| **Apus_Agal_Aaus** | **Agal** | 0 | 10.23 | 0.95 | 0.05 | 0.0005 | 0.0005 | -296 | -291 | -5 | 22 | 0.0% | 0.0% |
| **Apus_Agal_Aaus** | **Aaus** | 0 | 7.51 | 0.97 | 0.03 | 0.0003 | 0.0003 | -401 | -403 | 2 | 28 | 0.0% | 0.0% |
| **Apus_Agal_Phoo** | **Apus** | 0 | 5.56 | 1.00 | 0.00 | 0.0002 | 0.0002 | -31540 | -31554 | 14 | 2088 | 0.0% | 37.9% |
| **Apus_Agal_Phoo** | **Agal** | 0 | 4.21 | 0.99 | 0.01 | 0.0001 | 0.0001 | -15134 | -15146 | 12 | 954 | 0.0% | 17.2% |
| **Apus_Agal_Phoo** | **Phoo** | 0 | 4.79 | 0.99 | 0.01 | 0.0002 | 0.0002 | -36787 | -36798 | 10 | 2445 | 0.0% | 44.2% |
| **Apus_Agal_Zwol** | **Apus** | 0 | 0.94 | 0.05 | 0.95 | 0.0005 | 0.0005 | -50 | -51 | 1 | 4 | 0.0% | 0.0% |
| **Apus_Agal_Zwol** | **Agal** | 0 | 0.64 | 0.00 | 1.00 | 0.0004 | 0.0004 | -54 | -54 | 0 | 4 | 0.0% | 0.0% |
| **Apus_Agal_Zwol** | **Zwol** | 0 | 2.60 | 0.00 | 1.00 | 0.0005 | 0.0014 | -66961 | -60852 | -6110 | 5479 | 99.9% | 0.0% |
| **Apus_Agal_Zcal** | **Apus** | 0 | 0.94 | 0.05 | 0.95 | 0.0005 | 0.0005 | -50 | -51 | 1 | 4 | 0.0% | 0.0% |
| **Apus_Agal_Zcal** | **Agal** | 0 | 0.64 | 0.00 | 1.00 | 0.0004 | 0.0004 | -54 | -54 | 0 | 4 | 0.0% | 0.0% |
| **Apus_Agal_Zcal** | **Zcal** | 0 | 2.60 | 0.00 | 1.00 | 0.0005 | 0.0014 | -66961 | -60852 | -6110 | 5479 | 99.9% | 0.0% |
| **Apus_Agal_Ejub** | **Apus** | 0 | 0.94 | 0.05 | 0.95 | 0.0005 | 0.0005 | -50 | -51 | 1 | 4 | 0.0% | 0.0% |
| **Apus_Agal_Ejub** | **Agal** | 0 | 0.64 | 0.00 | 1.00 | 0.0004 | 0.0004 | -54 | -54 | 0 | 4 | 0.0% | 0.0% |
| **Apus_Agal_Ejub** | **Ejub** | 0 | 2.60 | 0.00 | 1.00 | 0.0005 | 0.0014 | -66961 | -60852 | -6110 | 5479 | 99.9% | 0.0% |
| **Apus_Agal_Curs** | **Apus** | 0 | 10.97 | 0.50 | 0.50 | 0.0005 | 0.0030 | -22 | -19 | -3 | 2 | 0.0% | 0.0% |
| **Apus_Agal_Curs** | **Agal** | 0 | 0 | 0 | 0 | 0 | 0 | 0 | 0 | 0 | 0 | 0.0% | 0.0% |
| **Apus_Agal_Curs** | **Curs** | 0 | 3.79 | 0.00 | 1.00 | 0.0008 | 0.0034 | -60100 | -51262 | -8837 | 5485 | 100.0% | 0.0% |
| **Apus_Agal_Oros** | **Apus** | 0 | 0 | 0 | 0 | 0 | 0 | 0 | 0 | 0 | 0 | 0.0% | 0.0% |
| **Apus_Agal_Oros** | **Agal** | 0 | 0 | 0 | 0 | 0 | 0 | 0 | 0 | 0 | 0 | 0.0% | 0.0% |
| **Apus_Agal_Oros** | **Oros** | 0 | 5.70 | 0.00 | 1.00 | 0.0021 | 0.0125 | -49422 | -37096 | -12327 | 5487 | 100.0% | 0.0% |
| **Apus_Aaus_Phoo** | **Apus** | 0 | 5.57 | 1.00 | 0.00 | 0.0002 | 0.0002 | -31609 | -31623 | 14 | 2092 | 0.0% | 38.0% |
| **Apus_Aaus_Phoo** | **Aaus** | 0 | 4.25 | 0.99 | 0.01 | 0.0001 | 0.0001 | -15199 | -15209 | 10 | 954 | 0.0% | 17.2% |
| **Apus_Aaus_Phoo** | **Phoo** | 0 | 4.79 | 0.99 | 0.01 | 0.0002 | 0.0002 | -36732 | -36743 | 11 | 2441 | 0.0% | 44.2% |
| **Apus_Aaus_Zwol** | **Apus** | 0 | 0.94 | 0.05 | 0.95 | 0.0005 | 0.0005 | -50 | -51 | 1 | 4 | 0.0% | 0.0% |
| **Apus_Aaus_Zwol** | **Aaus** | 0 | 0.64 | 0.00 | 1.00 | 0.0004 | 0.0004 | -54 | -54 | 0 | 4 | 0.0% | 0.0% |
| **Apus_Aaus_Zwol** | **Zwol** | 0 | 2.60 | 0.00 | 1.00 | 0.0005 | 0.0014 | -66945 | -60848 | -6098 | 5479 | 99.9% | 0.0% |
| **Apus_Aaus_Zcal** | **Apus** | 0 | 0.94 | 0.05 | 0.95 | 0.0005 | 0.0005 | -50 | -51 | 1 | 4 | 0.0% | 0.0% |

| | | | | | | | | | | | | | |
|---|---|---|---|---|---|---|---|---|---|---|---|---|---|
| **Apus_Aaus_Zcal** | **Aaus** | 0 | 0.64 | 0.00 | 1.00 | 0.0004 | 0.0004 | -54 | -54 | 0 | 4 | 0.0% | 0.0% |
| **Apus_Aaus_Zcal** | **Zcal** | 0 | 2.60 | 0.00 | 1.00 | 0.0005 | 0.0014 | -66945 | -60848 | -6098 | 5479 | 99.9% | 0.0% |
| **Apus_Aaus_Ejub** | **Apus** | 0 | 0.94 | 0.05 | 0.95 | 0.0005 | 0.0005 | -50 | -51 | 1 | 4 | 0.0% | 0.0% |
| **Apus_Aaus_Ejub** | **Aaus** | 0 | 0.64 | 0.00 | 1.00 | 0.0004 | 0.0004 | -54 | -54 | 0 | 4 | 0.0% | 0.0% |
| **Apus_Aaus_Ejub** | **Ejub** | 0 | 2.60 | 0.00 | 1.00 | 0.0005 | 0.0014 | -66945 | -60848 | -6098 | 5479 | 99.9% | 0.0% |
| **Apus_Aaus_Curs** | **Apus** | 0 | 10.97 | 0.50 | 0.50 | 0.0005 | 0.0030 | -22 | -19 | -3 | 2 | 0.0% | 0.0% |
| **Apus_Aaus_Curs** | **Aaus** | 0 | 0 | 0 | 0 | 0 | 0 | 0 | 0 | 0 | 0 | 0.0% | 0.0% |
| **Apus_Aaus_Curs** | **Curs** | 0 | 3.78 | 0.00 | 1.00 | 0.0008 | 0.0034 | -60093 | -51261 | -8832 | 5485 | 100.0% | 0.0% |
| **Apus_Aaus_Oros** | **Apus** | 0 | 0 | 0 | 0 | 0 | 0 | 0 | 0 | 0 | 0 | 0.0% | 0.0% |
| **Apus_Aaus_Oros** | **Aaus** | 0 | 0 | 0 | 0 | 0 | 0 | 0 | 0 | 0 | 0 | 0.0% | 0.0% |
| **Apus_Aaus_Oros** | **Oros** | 0 | 5.70 | 0.00 | 1.00 | 0.0021 | 0.0125 | -49421 | -37095 | -12326 | 5487 | 100.0% | 0.0% |
| **Apus_Phoo_Zwol** | **Apus** | 0 | 1.47 | 0.37 | 0.63 | 0.0005 | 0.0006 | -86 | -88 | 2 | 7 | 0.0% | 0.0% |
| **Apus_Phoo_Zwol** | **Phoo** | 0 | 0.74 | 0.50 | 0.50 | 0.0003 | 0.0003 | -107 | -111 | 3 | 8 | 0.0% | 0.0% |
| **Apus_Phoo_Zwol** | **Zwol** | 0 | 2.48 | 0.00 | 1.00 | 0.0005 | 0.0014 | -67165 | -61274 | -5891 | 5472 | 99.7% | 0.0% |
| **Apus_Phoo_Zcal** | **Apus** | 0 | 1.47 | 0.37 | 0.63 | 0.0005 | 0.0006 | -86 | -88 | 2 | 7 | 0.0% | 0.0% |
| **Apus_Phoo_Zcal** | **Phoo** | 0 | 0.74 | 0.50 | 0.50 | 0.0003 | 0.0003 | -107 | -111 | 3 | 8 | 0.0% | 0.0% |
| **Apus_Phoo_Zcal** | **Zcal** | 0 | 2.48 | 0.00 | 1.00 | 0.0005 | 0.0014 | -67165 | -61274 | -5891 | 5472 | 99.7% | 0.0% |
| **Apus_Phoo_Ejub** | **Apus** | 0 | 1.47 | 0.37 | 0.63 | 0.0005 | 0.0006 | -86 | -88 | 2 | 7 | 0.0% | 0.0% |
| **Apus_Phoo_Ejub** | **Phoo** | 0 | 0.74 | 0.50 | 0.50 | 0.0003 | 0.0003 | -107 | -111 | 3 | 8 | 0.0% | 0.0% |
| **Apus_Phoo_Ejub** | **Ejub** | 0 | 2.48 | 0.00 | 1.00 | 0.0005 | 0.0014 | -67165 | -61274 | -5891 | 5472 | 99.7% | 0.0% |
| **Apus_Phoo_Curs** | **Apus** | 0 | 0 | 0 | 0 | 0 | 0 | 0 | 0 | 0 | 1 | 0.0% | 0.0% |
| **Apus_Phoo_Curs** | **Phoo** | 0 | 0 | 0 | 0 | 0 | 0 | 0 | 0 | 0 | 1 | 0.0% | 0.0% |
| **Apus_Phoo_Curs** | **Curs** | 0 | 3.78 | 0.00 | 1.00 | 0.0008 | 0.0034 | -60265 | -51474 | -8791 | 5485 | 100.0% | 0.0% |
| **Apus_Phoo_Oros** | **Apus** | 0 | 0 | 0 | 0 | 0 | 0 | 0 | 0 | 0 | 0 | 0.0% | 0.0% |
| **Apus_Phoo_Oros** | **Phoo** | 0 | 0 | 0 | 0 | 0 | 0 | 0 | 0 | 0 | 0 | 0.0% | 0.0% |
| **Apus_Phoo_Oros** | **Oros** | 0 | 5.71 | 0.00 | 1.00 | 0.0021 | 0.0124 | -49507 | -37153 | -12353 | 5487 | 100.0% | 0.0% |
| **Apus_Zwol_Zcal** | **Apus** | 0 | 3.97 | 0.00 | 1.00 | 0.0006 | 0.0026 | -63343 | -54296 | -9047 | 5487 | 100.0% | 0.0% |
| **Apus_Zwol_Zcal** | **Zwol** | 0 | 0 | 0 | 0 | 0 | 0 | 0 | 0 | 0 | 0 | 0.0% | 0.0% |
| **Apus_Zwol_Zcal** | **Zcal** | 0 | 0 | 0 | 0 | 0 | 0 | 0 | 0 | 0 | 0 | 0.0% | 0.0% |
| **Apus_Zwol_Ejub** | **Apus** | 0 | 3.98 | 0.00 | 1.00 | 0.0006 | 0.0026 | -63346 | -54308 | -9038 | 5487 | 100.0% | 0.0% |
| **Apus_Zwol_Ejub** | **Zwol** | 0 | 0 | 0 | 0 | 0 | 0 | 0 | 0 | 0 | 0 | 0.0% | 0.0% |
| **Apus_Zwol_Ejub** | **Ejub** | 0 | 0 | 0 | 0 | 0 | 0 | 0 | 0 | 0 | 0 | 0.0% | 0.0% |

| | | | | | | | | | | | | | |
|---|---|---|---|---|---|---|---|---|---|---|---|---|---|
| **Apus_Zwol_Curs** | **Apus** | 0 | 0 | 0 | 0 | 0 | 0 | 0 | 0 | 0 | 1 | 0.0% | 0.0% |
| **Apus_Zwol_Curs** | **Zwol** | 0 | 1.56 | 0.04 | 0.96 | 0.0005 | 0.0006 | -37 | -37 | 0 | 3 | 0.0% | 0.0% |
| **Apus_Zwol_Curs** | **Curs** | 0 | 3.13 | 0.00 | 1.00 | 0.0006 | 0.0020 | -64468 | -57101 | -7366 | 5483 | 99.9% | 0.0% |
| **Apus_Zwol_Oros** | **Apus** | 0 | 0 | 0 | 0 | 0 | 0 | 0 | 0 | 0 | 0 | 0.0% | 0.0% |
| **Apus_Zwol_Oros** | **Zwol** | 0 | 0 | 0 | 0 | 0 | 0 | 0 | 0 | 0 | 0 | 0.0% | 0.0% |
| **Apus_Zwol_Oros** | **Oros** | 0 | 5.89 | 0.00 | 1.00 | 0.0018 | 0.0111 | -51054 | -38420 | -12634 | 5487 | 100.0% | 0.0% |
| **Apus_Zcal_Ejub** | **Apus** | 0 | 3.97 | 0.00 | 1.00 | 0.0006 | 0.0027 | -63112 | -54047 | -9065 | 5487 | 100.0% | 0.0% |
| **Apus_Zcal_Ejub** | **Zcal** | 0 | 0 | 0 | 0 | 0 | 0 | 0 | 0 | 0 | 0 | 0.0% | 0.0% |
| **Apus_Zcal_Ejub** | **Ejub** | 0 | 0 | 0 | 0 | 0 | 0 | 0 | 0 | 0 | 0 | 0.0% | 0.0% |
| **Apus_Zcal_Curs** | **Apus** | 0 | 0 | 0 | 0 | 0 | 0 | 0 | 0 | 0 | 1 | 0.0% | 0.0% |
| **Apus_Zcal_Curs** | **Zcal** | 0 | 1.56 | 0.04 | 0.96 | 0.0005 | 0.0006 | -37 | -37 | 0 | 3 | 0.0% | 0.0% |
| **Apus_Zcal_Curs** | **Curs** | 0 | 3.13 | 0.00 | 1.00 | 0.0006 | 0.0020 | -64468 | -57101 | -7366 | 5483 | 99.9% | 0.0% |
| **Apus_Zcal_Oros** | **Apus** | 0 | 0 | 0 | 0 | 0 | 0 | 0 | 0 | 0 | 0 | 0.0% | 0.0% |
| **Apus_Zcal_Oros** | **Zcal** | 0 | 0 | 0 | 0 | 0 | 0 | 0 | 0 | 0 | 0 | 0.0% | 0.0% |
| **Apus_Zcal_Oros** | **Oros** | 0 | 5.89 | 0.00 | 1.00 | 0.0018 | 0.0111 | -51054 | -38420 | -12634 | 5487 | 100.0% | 0.0% |
| **Apus_Ejub_Curs** | **Apus** | 0 | 0 | 0 | 0 | 0 | 0 | 0 | 0 | 0 | 1 | 0.0% | 0.0% |
| **Apus_Ejub_Curs** | **Ejub** | 0 | 1.56 | 0.04 | 0.96 | 0.0005 | 0.0006 | -37 | -37 | 0 | 3 | 0.0% | 0.0% |
| **Apus_Ejub_Curs** | **Curs** | 0 | 3.13 | 0.00 | 1.00 | 0.0006 | 0.0020 | -64468 | -57101 | -7366 | 5483 | 99.9% | 0.0% |
| **Apus_Ejub_Oros** | **Apus** | 0 | 0 | 0 | 0 | 0 | 0 | 0 | 0 | 0 | 0 | 0.0% | 0.0% |
| **Apus_Ejub_Oros** | **Ejub** | 0 | 0 | 0 | 0 | 0 | 0 | 0 | 0 | 0 | 0 | 0.0% | 0.0% |
| **Apus_Ejub_Oros** | **Oros** | 0 | 5.89 | 0.00 | 1.00 | 0.0018 | 0.0111 | -51054 | -38420 | -12634 | 5487 | 100.0% | 0.0% |
| **Apus_Curs_Oros** | **Apus** | 0 | 0 | 0 | 0 | 0 | 0 | 0 | 0 | 0 | 0 | 0.0% | 0.0% |
| **Apus_Curs_Oros** | **Curs** | 0 | 0 | 0 | 0 | 0 | 0 | 0 | 0 | 0 | 0 | 0.0% | 0.0% |
| **Apus_Curs_Oros** | **Oros** | 0 | 6.17 | 0.00 | 1.00 | 0.0014 | 0.0091 | -53696 | -40616 | -13080 | 5487 | 100.0% | 0.0% |
| **Atro_Ofla_Atow** | **Atro** | 0 | 6.13 | 1.00 | 0.00 | 0.0002 | 0.0002 | -23174 | -23188 | 13 | 1532 | 0.0% | 27.9% |
| **Atro_Ofla_Atow** | **Ofla** | 0 | 6.13 | 1.00 | 0.00 | 0.0002 | 0.0002 | -39814 | -39824 | 11 | 2708 | 0.0% | 49.2% |
| **Atro_Ofla_Atow** | **Atow** | 0 | 4.96 | 0.99 | 0.01 | 0.0002 | 0.0002 | -19203 | -19216 | 13 | 1247 | 0.0% | 22.6% |
| **Atro_Ofla_Aphi** | **Atro** | 0 | 6.15 | 1.00 | 0.00 | 0.0002 | 0.0002 | -23182 | -23196 | 13 | 1532 | 0.0% | 27.9% |
| **Atro_Ofla_Aphi** | **Ofla** | 0 | 6.17 | 1.00 | 0.00 | 0.0002 | 0.0002 | -39821 | -39832 | 11 | 2706 | 0.0% | 49.2% |
| **Atro_Ofla_Aphi** | **Aphi** | 0 | 4.97 | 1.00 | 0.00 | 0.0002 | 0.0002 | -19238 | -19251 | 13 | 1249 | 0.0% | 22.7% |
| **Atro_Ofla_Agaz** | **Atro** | 0 | 3.97 | 0.99 | 0.01 | 0.0002 | 0.0002 | -18065 | -18078 | 14 | 1185 | 0.0% | 21.5% |
| **Atro_Ofla_Agaz** | **Ofla** | 0 | 6.34 | 1.00 | 0.00 | 0.0003 | 0.0003 | -45033 | -45043 | 11 | 3100 | 0.0% | 56.4% |

| | | | | | | | | | | | | | |
|---|---|---|---|---|---|---|---|---|---|---|---|---|---|
| **Atro_Ofla_Agaz** | **Agaz** | 0 | 5.24 | 0.99 | 0.01 | 0.0002 | 0.0002 | -18536 | -18547 | 11 | 1202 | 0.0% | 21.8% |
| **Atro_Ofla_Afor** | **Atro** | 0 | 4.39 | 0.99 | 0.01 | 0.0002 | 0.0002 | -15771 | -15781 | 10 | 1012 | 0.0% | 0.0% |
| **Atro_Ofla_Afor** | **Ofla** | 0 | 6.37 | 1.00 | 0.00 | 0.0002 | 0.0002 | -45713 | -45728 | 16 | 3114 | 0.0% | 56.7% |
| **Atro_Ofla_Afor** | **Afor** | 0 | 5.08 | 1.00 | 0.00 | 0.0002 | 0.0002 | -21092 | -21106 | 14 | 1361 | 0.0% | 24.7% |
| **Atro_Ofla_Agal** | **Atro** | 0 | 4.37 | 0.99 | 0.01 | 0.0002 | 0.0002 | -15636 | -15645 | 10 | 1004 | 0.0% | 0.0% |
| **Atro_Ofla_Agal** | **Ofla** | 0 | 6.26 | 1.00 | 0.00 | 0.0002 | 0.0002 | -45870 | -45885 | 16 | 3123 | 0.0% | 56.9% |
| **Atro_Ofla_Agal** | **Agal** | 0 | 5.44 | 0.99 | 0.01 | 0.0002 | 0.0002 | -21016 | -21028 | 12 | 1360 | 0.0% | 24.7% |
| **Atro_Ofla_Aaus** | **Atro** | 0 | 4.38 | 0.99 | 0.01 | 0.0002 | 0.0002 | -15766 | -15777 | 10 | 1012 | 0.0% | 18.2% |
| **Atro_Ofla_Aaus** | **Ofla** | 0 | 5.72 | 1.00 | 0.00 | 0.0002 | 0.0002 | -45723 | -45739 | 16 | 3116 | 0.0% | 56.8% |
| **Atro_Ofla_Aaus** | **Aaus** | 0 | 5.49 | 1.00 | 0.00 | 0.0002 | 0.0002 | -21020 | -21033 | 13 | 1359 | 0.0% | 24.7% |
| **Atro_Ofla_Phoo** | **Atro** | 0 | 4.53 | 0.99 | 0.01 | 0.0002 | 0.0002 | -21962 | -21972 | 10 | 1436 | 0.0% | 0.0% |
| **Atro_Ofla_Phoo** | **Ofla** | 0 | 5.79 | 1.00 | 0.00 | 0.0002 | 0.0002 | -34167 | -34181 | 14 | 2287 | 0.0% | 41.6% |
| **Atro_Ofla_Phoo** | **Phoo** | 0 | 4.64 | 0.99 | 0.01 | 0.0002 | 0.0002 | -26948 | -26959 | 11 | 1764 | 0.0% | 31.9% |
| **Atro_Ofla_Zwol** | **Atro** | 0 | 0.60 | 0.04 | 0.96 | 0.0005 | 0.0007 | -161 | -162 | 1 | 13 | 0.0% | 0.0% |
| **Atro_Ofla_Zwol** | **Ofla** | 0 | 0.58 | 0.00 | 1.00 | 0.0002 | 0.0002 | -43 | -43 | 0 | 3 | 0.0% | 0.0% |
| **Atro_Ofla_Zwol** | **Zwol** | 0 | 2.43 | 0.00 | 1.00 | 0.0005 | 0.0013 | -67316 | -61505 | -5811 | 5471 | 99.7% | 0.0% |
| **Atro_Ofla_Zcal** | **Atro** | 0 | 0.60 | 0.04 | 0.96 | 0.0005 | 0.0007 | -161 | -162 | 1 | 13 | 0.0% | 0.0% |
| **Atro_Ofla_Zcal** | **Ofla** | 0 | 0.58 | 0.00 | 1.00 | 0.0002 | 0.0002 | -43 | -43 | 0 | 3 | 0.0% | 0.0% |
| **Atro_Ofla_Zcal** | **Zcal** | 0 | 2.43 | 0.00 | 1.00 | 0.0005 | 0.0013 | -67316 | -61505 | -5811 | 5471 | 99.7% | 0.0% |
| **Atro_Ofla_Ejub** | **Atro** | 0 | 0.60 | 0.04 | 0.96 | 0.0005 | 0.0007 | -161 | -162 | 1 | 13 | 0.0% | 0.0% |
| **Atro_Ofla_Ejub** | **Ofla** | 0 | 0.58 | 0.00 | 1.00 | 0.0002 | 0.0002 | -43 | -43 | 0 | 3 | 0.0% | 0.0% |
| **Atro_Ofla_Ejub** | **Ejub** | 0 | 2.43 | 0.00 | 1.00 | 0.0005 | 0.0013 | -67316 | -61505 | -5811 | 5471 | 99.7% | 0.0% |
| **Atro_Ofla_Curs** | **Atro** | 0 | 1.38 | 0.00 | 1.00 | 0.0008 | 0.0018 | -21 | -21 | -1 | 2 | 0.0% | 0.0% |
| **Atro_Ofla_Curs** | **Ofla** | 0 | 0 | 0 | 0 | 0 | 0 | 0 | 0 | 0 | 0 | 0.0% | 0.0% |
| **Atro_Ofla_Curs** | **Curs** | 0 | 3.78 | 0.00 | 1.00 | 0.0008 | 0.0033 | -60359 | -51570 | -8789 | 5485 | 100.0% | 0.0% |
| **Atro_Ofla_Oros** | **Atro** | 0 | 0 | 0 | 0 | 0 | 0 | 0 | 0 | 0 | 0 | 0.0% | 0.0% |
| **Atro_Ofla_Oros** | **Ofla** | 0 | 0 | 0 | 0 | 0 | 0 | 0 | 0 | 0 | 0 | 0.0% | 0.0% |
| **Atro_Ofla_Oros** | **Oros** | 0 | 5.74 | 0.00 | 1.00 | 0.0020 | 0.0124 | -49559 | -37179 | -12380 | 5487 | 100.0% | 0.0% |
| **Atro_Atow_Aphi** | **Atro** | 0 | 1.97 | 0.00 | 1.00 | 0.0005 | 0.0011 | -69333 | -63348 | -5986 | 5480 | 99.9% | 0.0% |
| **Atro_Atow_Aphi** | **Atow** | 0 | 0.65 | 0.00 | 1.00 | 0.0005 | 0.0005 | -26 | -26 | 0 | 2 | 0.0% | 0.0% |
| **Atro_Atow_Aphi** | **Aphi** | 0 | 8.83 | 0.80 | 0.20 | 0.0005 | 0.0012 | -59 | -56 | -3 | 5 | 0.0% | 0.0% |
| **Atro_Atow_Agaz** | **Atro** | 0 | 6.79 | 1.00 | 0.00 | 0.0003 | 0.0003 | -40200 | -40216 | 16 | 2770 | 0.0% | 50.5% |

| | | | | | | | | | | | | | |
|---|---|---|---|---|---|---|---|---|---|---|---|---|---|
| **Atro_Atow_Agaz** | **Atow** | 0 | 7.32 | 1.00 | 0.00 | 0.0002 | 0.0002 | -24954 | -24968 | 14 | 1664 | 0.0% | 30.3% |
| **Atro_Atow_Agaz** | **Agaz** | 0 | 5.21 | 0.99 | 0.01 | 0.0002 | 0.0002 | -16288 | -16301 | 12 | 1053 | 0.0% | 19.1% |
| **Atro_Atow_Afor** | **Atro** | 0 | 6.86 | 1.00 | 0.00 | 0.0003 | 0.0003 | -42877 | -42893 | 16 | 2941 | 0.0% | 53.6% |
| **Atro_Atow_Afor** | **Atow** | 0 | 5.21 | 1.00 | 0.00 | 0.0002 | 0.0002 | -22601 | -22615 | 14 | 1478 | 0.0% | 26.9% |
| **Atro_Atow_Afor** | **Afor** | 0 | 5.25 | 0.99 | 0.01 | 0.0002 | 0.0002 | -16624 | -16632 | 8 | 1068 | 0.0% | 0.0% |
| **Atro_Atow_Agal** | **Atro** | 0 | 8.54 | 1.00 | 0.00 | 0.0003 | 0.0003 | -42881 | -42897 | 16 | 2948 | 0.0% | 53.7% |
| **Atro_Atow_Agal** | **Atow** | 0 | 5.27 | 1.00 | 0.00 | 0.0002 | 0.0002 | -22604 | -22618 | 14 | 1476 | 0.0% | 26.8% |
| **Atro_Atow_Agal** | **Agal** | 0 | 5.24 | 0.99 | 0.01 | 0.0002 | 0.0002 | -16505 | -16514 | 9 | 1063 | 0.0% | 0.0% |
| **Atro_Atow_Aaus** | **Atro** | 0 | 8.13 | 1.00 | 0.00 | 0.0003 | 0.0003 | -42936 | -42952 | 16 | 2950 | 0.0% | 53.8% |
| **Atro_Atow_Aaus** | **Atow** | 0 | 5.24 | 1.00 | 0.00 | 0.0002 | 0.0002 | -22538 | -22552 | 14 | 1473 | 0.0% | 26.8% |
| **Atro_Atow_Aaus** | **Aaus** | 0 | 4.87 | 0.99 | 0.01 | 0.0002 | 0.0002 | -16575 | -16586 | 10 | 1064 | 0.0% | 19.2% |
| **Atro_Atow_Phoo** | **Atro** | 0 | 5.76 | 1.00 | 0.00 | 0.0002 | 0.0002 | -30195 | -30211 | 15 | 1992 | 0.0% | 36.3% |
| **Atro_Atow_Phoo** | **Atow** | 0 | 6.14 | 0.99 | 0.01 | 0.0001 | 0.0001 | -18581 | -18585 | 4 | 1191 | 0.0% | 0.0% |
| **Atro_Atow_Phoo** | **Phoo** | 0 | 4.71 | 0.99 | 0.01 | 0.0002 | 0.0002 | -34799 | -34809 | 10 | 2304 | 0.0% | 0.0% |
| **Atro_Atow_Zwol** | **Atro** | 0 | 1.09 | 0.50 | 0.50 | 0.0004 | 0.0004 | -64 | -67 | 2 | 5 | 0.0% | 0.0% |
| **Atro_Atow_Zwol** | **Atow** | 0 | 0.57 | 0.00 | 1.00 | 0.0004 | 0.0004 | -39 | -40 | 0 | 3 | 0.0% | 0.0% |
| **Atro_Atow_Zwol** | **Zwol** | 0 | 2.53 | 0.00 | 1.00 | 0.0005 | 0.0014 | -66893 | -60980 | -5914 | 5479 | 99.9% | 0.0% |
| **Atro_Atow_Zcal** | **Atro** | 0 | 1.09 | 0.50 | 0.50 | 0.0004 | 0.0004 | -64 | -67 | 2 | 5 | 0.0% | 0.0% |
| **Atro_Atow_Zcal** | **Atow** | 0 | 0.57 | 0.00 | 1.00 | 0.0004 | 0.0004 | -39 | -40 | 0 | 3 | 0.0% | 0.0% |
| **Atro_Atow_Zcal** | **Zcal** | 0 | 2.53 | 0.00 | 1.00 | 0.0005 | 0.0014 | -66893 | -60980 | -5914 | 5479 | 99.9% | 0.0% |
| **Atro_Atow_Ejub** | **Atro** | 0 | 1.09 | 0.50 | 0.50 | 0.0004 | 0.0004 | -64 | -67 | 2 | 5 | 0.0% | 0.0% |
| **Atro_Atow_Ejub** | **Atow** | 0 | 0.57 | 0.00 | 1.00 | 0.0004 | 0.0004 | -39 | -40 | 0 | 3 | 0.0% | 0.0% |
| **Atro_Atow_Ejub** | **Ejub** | 0 | 2.53 | 0.00 | 1.00 | 0.0005 | 0.0014 | -66893 | -60980 | -5914 | 5479 | 99.9% | 0.0% |
| **Atro_Atow_Curs** | **Atro** | 0 | 0 | 0 | 0 | 0 | 0 | 0 | 0 | 0 | 1 | 0.0% | 0.0% |
| **Atro_Atow_Curs** | **Atow** | 0 | 0 | 0 | 0 | 0 | 0 | 0 | 0 | 0 | 0 | 0.0% | 0.0% |
| **Atro_Atow_Curs** | **Curs** | 0 | 3.72 | 0.00 | 1.00 | 0.0009 | 0.0034 | -60059 | -51326 | -8733 | 5486 | 100.0% | 0.0% |
| **Atro_Atow_Oros** | **Atro** | 0 | 0 | 0 | 0 | 0 | 0 | 0 | 0 | 0 | 0 | 0.0% | 0.0% |
| **Atro_Atow_Oros** | **Atow** | 0 | 0 | 0 | 0 | 0 | 0 | 0 | 0 | 0 | 0 | 0.0% | 0.0% |
| **Atro_Atow_Oros** | **Oros** | 0 | 5.68 | 0.00 | 1.00 | 0.0021 | 0.0125 | -49410 | -37110 | -12300 | 5487 | 100.0% | 0.0% |
| **Atro_Aphi_Agaz** | **Atro** | 0 | 6.97 | 1.00 | 0.00 | 0.0003 | 0.0003 | -40174 | -40190 | 16 | 2768 | 0.0% | 50.4% |
| **Atro_Aphi_Agaz** | **Aphi** | 0 | 7.32 | 1.00 | 0.00 | 0.0002 | 0.0002 | -24982 | -24996 | 14 | 1666 | 0.0% | 30.3% |
| **Atro_Aphi_Agaz** | **Agaz** | 0 | 5.24 | 0.99 | 0.01 | 0.0002 | 0.0002 | -16343 | -16354 | 11 | 1053 | 0.0% | 19.1% |

| | | | | | | | | | | | | | |
|---|---|---|---|---|---|---|---|---|---|---|---|---|---|
| **Atro_Aphi_Afor** | **Atro** | 0 | 6.84 | 1.00 | 0.00 | 0.0003 | 0.0003 | -42862 | -42878 | 16 | 2941 | 0.0% | 53.6% |
| **Atro_Aphi_Afor** | **Aphi** | 0 | 5.18 | 1.00 | 0.00 | 0.0002 | 0.0002 | -22603 | -22618 | 14 | 1480 | 0.0% | 26.9% |
| **Atro_Aphi_Afor** | **Afor** | 0 | 4.89 | 0.99 | 0.01 | 0.0002 | 0.0002 | -16613 | -16623 | 10 | 1066 | 0.0% | 19.3% |
| **Atro_Aphi_Agal** | **Atro** | 0 | 8.54 | 1.00 | 0.00 | 0.0003 | 0.0003 | -42870 | -42885 | 15 | 2947 | 0.0% | 53.7% |
| **Atro_Aphi_Agal** | **Aphi** | 0 | 5.27 | 1.00 | 0.00 | 0.0002 | 0.0002 | -22617 | -22631 | 14 | 1477 | 0.0% | 26.9% |
| **Atro_Aphi_Agal** | **Agal** | 0 | 4.88 | 0.99 | 0.01 | 0.0002 | 0.0002 | -16562 | -16572 | 10 | 1063 | 0.0% | 19.2% |
| **Atro_Aphi_Aaus** | **Atro** | 0 | 8.11 | 1.00 | 0.00 | 0.0003 | 0.0003 | -42936 | -42952 | 16 | 2951 | 0.0% | 53.8% |
| **Atro_Aphi_Aaus** | **Aphi** | 0 | 5.18 | 1.00 | 0.00 | 0.0002 | 0.0002 | -22509 | -22523 | 14 | 1474 | 0.0% | 26.8% |
| **Atro_Aphi_Aaus** | **Aaus** | 0 | 4.89 | 0.99 | 0.01 | 0.0002 | 0.0002 | -16550 | -16560 | 10 | 1062 | 0.0% | 0.0% |
| **Atro_Aphi_Phoo** | **Atro** | 0 | 6.62 | 1.00 | 0.00 | 0.0002 | 0.0002 | -30201 | -30216 | 15 | 1992 | 0.0% | 36.3% |
| **Atro_Aphi_Phoo** | **Aphi** | 0 | 7.87 | 0.99 | 0.01 | 0.0002 | 0.0002 | -18588 | -18578 | -10 | 1194 | 0.2% | 0.0% |
| **Atro_Aphi_Phoo** | **Phoo** | 0 | 4.71 | 0.99 | 0.01 | 0.0002 | 0.0002 | -34751 | -34761 | 10 | 2301 | 0.0% | 0.0% |
| **Atro_Aphi_Zwol** | **Atro** | 0 | 1.09 | 0.50 | 0.50 | 0.0004 | 0.0004 | -64 | -67 | 2 | 5 | 0.0% | 0.0% |
| **Atro_Aphi_Zwol** | **Aphi** | 0 | 0.57 | 0.00 | 1.00 | 0.0004 | 0.0004 | -39 | -40 | 0 | 3 | 0.0% | 0.0% |
| **Atro_Aphi_Zwol** | **Zwol** | 0 | 2.56 | 0.00 | 1.00 | 0.0005 | 0.0014 | -66907 | -60987 | -5920 | 5479 | 99.9% | 0.0% |
| **Atro_Aphi_Zcal** | **Atro** | 0 | 1.09 | 0.50 | 0.50 | 0.0004 | 0.0004 | -64 | -67 | 2 | 5 | 0.0% | 0.0% |
| **Atro_Aphi_Zcal** | **Aphi** | 0 | 0.57 | 0.00 | 1.00 | 0.0004 | 0.0004 | -39 | -40 | 0 | 3 | 0.0% | 0.0% |
| **Atro_Aphi_Zcal** | **Zcal** | 0 | 2.56 | 0.00 | 1.00 | 0.0005 | 0.0014 | -66907 | -60987 | -5920 | 5479 | 99.9% | 0.0% |
| **Atro_Aphi_Ejub** | **Atro** | 0 | 1.09 | 0.50 | 0.50 | 0.0004 | 0.0004 | -64 | -67 | 2 | 5 | 0.0% | 0.0% |
| **Atro_Aphi_Ejub** | **Aphi** | 0 | 0.57 | 0.00 | 1.00 | 0.0004 | 0.0004 | -39 | -40 | 0 | 3 | 0.0% | 0.0% |
| **Atro_Aphi_Ejub** | **Ejub** | 0 | 2.56 | 0.00 | 1.00 | 0.0005 | 0.0014 | -66907 | -60987 | -5920 | 5479 | 99.9% | 0.0% |
| **Atro_Aphi_Curs** | **Atro** | 0 | 0 | 0 | 0 | 0 | 0 | 0 | 0 | 0 | 1 | 0.0% | 0.0% |
| **Atro_Aphi_Curs** | **Aphi** | 0 | 0 | 0 | 0 | 0 | 0 | 0 | 0 | 0 | 0 | 0.0% | 0.0% |
| **Atro_Aphi_Curs** | **Curs** | 0 | 3.73 | 0.00 | 1.00 | 0.0009 | 0.0034 | -60066 | -51329 | -8737 | 5486 | 100.0% | 0.0% |
| **Atro_Aphi_Oros** | **Atro** | 0 | 0 | 0 | 0 | 0 | 0 | 0 | 0 | 0 | 0 | 0.0% | 0.0% |
| **Atro_Aphi_Oros** | **Aphi** | 0 | 0 | 0 | 0 | 0 | 0 | 0 | 0 | 0 | 0 | 0.0% | 0.0% |
| **Atro_Aphi_Oros** | **Oros** | 0 | 5.68 | 0.00 | 1.00 | 0.0021 | 0.0125 | -49413 | -37111 | -12302 | 5487 | 100.0% | 0.0% |
| **Atro_Agaz_Afor** | **Atro** | 0 | 4.93 | 1.00 | 0.00 | 0.0003 | 0.0003 | -45658 | -45676 | 17 | 3193 | 0.0% | 58.2% |
| **Atro_Agaz_Afor** | **Agaz** | 0 | 4.08 | 1.00 | 0.00 | 0.0002 | 0.0002 | -15547 | -15561 | 14 | 1021 | 0.0% | 18.5% |
| **Atro_Agaz_Afor** | **Afor** | 0 | 7.54 | 1.00 | 0.00 | 0.0002 | 0.0002 | -19098 | -19106 | 8 | 1273 | 0.0% | 0.0% |
| **Atro_Agaz_Agal** | **Atro** | 0 | 5.72 | 1.00 | 0.00 | 0.0003 | 0.0003 | -45702 | -45719 | 17 | 3199 | 0.0% | 58.3% |
| **Atro_Agaz_Agal** | **Agaz** | 0 | 4.06 | 0.99 | 0.01 | 0.0002 | 0.0002 | -15691 | -15704 | 13 | 1024 | 0.0% | 18.5% |

| | | | | | | | | | | | | |
|---|---|---|---|---|---|---|---|---|---|---|---|---|
| **Atro_Agaz_Agal** | **Agal** | 0 | 7.56 | 1.00 | 0.00 | 0.0002 | 0.0002 | -18971 | -18977 | 6 | 1264 | 0.0% | 0.0% |
| **Atro_Agaz_Aaus** | **Atro** | 0 | 4.69 | 1.00 | 0.00 | 0.0003 | 0.0003 | -45695 | -45714 | 18 | 3198 | 0.0% | 58.2% |
| **Atro_Agaz_Aaus** | **Agaz** | 0 | 4.15 | 1.00 | 0.00 | 0.0002 | 0.0002 | -15595 | -15609 | 14 | 1022 | 0.0% | 18.6% |
| **Atro_Agaz_Aaus** | **Aaus** | 0 | 7.58 | 1.00 | 0.00 | 0.0002 | 0.0002 | -19021 | -19029 | 8 | 1267 | 0.0% | 0.0% |
| **Atro_Agaz_Phoo** | **Atro** | 0 | 6.87 | 1.00 | 0.00 | 0.0002 | 0.0002 | -30420 | -30429 | 9 | 2023 | 0.0% | 0.0% |
| **Atro_Agaz_Phoo** | **Agaz** | 0 | 36.42 | 1.00 | 0.00 | 0.0002 | 0.0002 | -14586 | -14496 | -90 | 930 | 0.0% | 0.0% |
| **Atro_Agaz_Phoo** | **Phoo** | 0 | 5.78 | 1.00 | 0.00 | 0.0002 | 0.0002 | -37441 | -37455 | 15 | 2534 | 0.0% | 46.1% |
| **Atro_Agaz_Zwol** | **Atro** | 0 | 0.94 | 0.05 | 0.95 | 0.0005 | 0.0005 | -50 | -51 | 1 | 4 | 0.0% | 0.0% |
| **Atro_Agaz_Zwol** | **Agaz** | 0 | 1.02 | 0.59 | 0.41 | 0.0003 | 0.0003 | -52 | -55 | 2 | 4 | 0.0% | 0.0% |
| **Atro_Agaz_Zwol** | **Zwol** | 0 | 2.50 | 0.00 | 1.00 | 0.0005 | 0.0014 | -66603 | -60742 | -5861 | 5479 | 99.9% | 0.0% |
| **Atro_Agaz_Zcal** | **Atro** | 0 | 0.94 | 0.05 | 0.95 | 0.0005 | 0.0005 | -50 | -51 | 1 | 4 | 0.0% | 0.0% |
| **Atro_Agaz_Zcal** | **Agaz** | 0 | 1.02 | 0.59 | 0.41 | 0.0003 | 0.0003 | -52 | -55 | 2 | 4 | 0.0% | 0.0% |
| **Atro_Agaz_Zcal** | **Zcal** | 0 | 2.50 | 0.00 | 1.00 | 0.0005 | 0.0014 | -66603 | -60742 | -5861 | 5479 | 99.9% | 0.0% |
| **Atro_Agaz_Ejub** | **Atro** | 0 | 0.94 | 0.05 | 0.95 | 0.0005 | 0.0005 | -50 | -51 | 1 | 4 | 0.0% | 0.0% |
| **Atro_Agaz_Ejub** | **Agaz** | 0 | 1.02 | 0.59 | 0.41 | 0.0003 | 0.0003 | -52 | -55 | 2 | 4 | 0.0% | 0.0% |
| **Atro_Agaz_Ejub** | **Ejub** | 0 | 2.50 | 0.00 | 1.00 | 0.0005 | 0.0014 | -66603 | -60742 | -5861 | 5479 | 99.9% | 0.0% |
| **Atro_Agaz_Curs** | **Atro** | 0 | 0 | 0 | 0 | 0 | 0 | 0 | 0 | 0 | 1 | 0.0% | 0.0% |
| **Atro_Agaz_Curs** | **Agaz** | 0 | 0 | 0 | 0 | 0 | 0 | 0 | 0 | 0 | 1 | 0.0% | 0.0% |
| **Atro_Agaz_Curs** | **Curs** | 0 | 3.68 | 0.00 | 1.00 | 0.0009 | 0.0034 | -59919 | -51217 | -8702 | 5485 | 100.0% | 0.0% |
| **Atro_Agaz_Oros** | **Atro** | 0 | 0 | 0 | 0 | 0 | 0 | 0 | 0 | 0 | 0 | 0.0% | 0.0% |
| **Atro_Agaz_Oros** | **Agaz** | 0 | 0 | 0 | 0 | 0 | 0 | 0 | 0 | 0 | 0 | 0.0% | 0.0% |
| **Atro_Agaz_Oros** | **Oros** | 0 | 5.66 | 0.00 | 1.00 | 0.0021 | 0.0125 | -49359 | -37083 | -12276 | 5487 | 100.0% | 0.0% |
| **Atro_Afor_Agal** | **Atro** | 0 | 1.18 | 0.01 | 0.99 | 0.0005 | 0.0008 | -69204 | -66085 | -3120 | 5382 | 96.9% | 0.0% |
| **Atro_Afor_Agal** | **Afor** | 0 | 5.03 | 0.99 | 0.01 | 0.0002 | 0.0002 | -828 | -835 | 8 | 56 | 0.0% | 0.0% |
| **Atro_Afor_Agal** | **Agal** | 0 | 7.19 | 0.96 | 0.04 | 0.0003 | 0.0003 | -680 | -679 | -2 | 49 | 0.0% | 0.0% |
| **Atro_Afor_Aaus** | **Atro** | 0 | 1.18 | 0.00 | 1.00 | 0.0005 | 0.0008 | -69259 | -66058 | -3201 | 5391 | 98.0% | 0.0% |
| **Atro_Afor_Aaus** | **Afor** | 0 | 4.88 | 0.97 | 0.03 | 0.0002 | 0.0002 | -722 | -728 | 6 | 49 | 0.0% | 0.0% |
| **Atro_Afor_Aaus** | **Aaus** | 0 | 0.32 | 0.84 | 0.16 | 0.0002 | 0.0002 | -684 | -692 | 8 | 47 | 0.0% | 0.0% |
| **Atro_Afor_Phoo** | **Atro** | 0 | 5.56 | 1.00 | 0.00 | 0.0002 | 0.0002 | -31616 | -31631 | 15 | 2093 | 0.0% | 38.1% |
| **Atro_Afor_Phoo** | **Afor** | 0 | 4.24 | 0.99 | 0.01 | 0.0001 | 0.0001 | -15194 | -15206 | 12 | 957 | 0.0% | 17.3% |
| **Atro_Afor_Phoo** | **Phoo** | 0 | 4.78 | 0.99 | 0.01 | 0.0002 | 0.0002 | -36648 | -36660 | 12 | 2437 | 0.0% | 44.1% |
| **Atro_Afor_Zwol** | **Atro** | 0 | 0.94 | 0.05 | 0.95 | 0.0005 | 0.0005 | -50 | -51 | 1 | 4 | 0.0% | 0.0% |

| | | | | | | | | | | | | | |
|---|---|---|---|---|---|---|---|---|---|---|---|---|---|
| **Atro_Afor_Zwol** | **Afor** | 0 | 0.64 | 0.00 | 1.00 | 0.0004 | 0.0004 | -54 | -54 | 0 | 4 | 0.0% | 0.0% |
| **Atro_Afor_Zwol** | **Zwol** | 0 | 2.59 | 0.00 | 1.00 | 0.0005 | 0.0014 | -66949 | -60845 | -6104 | 5479 | 99.9% | 0.0% |
| **Atro_Afor_Zcal** | **Atro** | 0 | 0.94 | 0.05 | 0.95 | 0.0005 | 0.0005 | -50 | -51 | 1 | 4 | 0.0% | 0.0% |
| **Atro_Afor_Zcal** | **Afor** | 0 | 0.64 | 0.00 | 1.00 | 0.0004 | 0.0004 | -54 | -54 | 0 | 4 | 0.0% | 0.0% |
| **Atro_Afor_Zcal** | **Zcal** | 0 | 2.59 | 0.00 | 1.00 | 0.0005 | 0.0014 | -66949 | -60845 | -6104 | 5479 | 99.9% | 0.0% |
| **Atro_Afor_Ejub** | **Atro** | 0 | 0.94 | 0.05 | 0.95 | 0.0005 | 0.0005 | -50 | -51 | 1 | 4 | 0.0% | 0.0% |
| **Atro_Afor_Ejub** | **Afor** | 0 | 0.64 | 0.00 | 1.00 | 0.0004 | 0.0004 | -54 | -54 | 0 | 4 | 0.0% | 0.0% |
| **Atro_Afor_Ejub** | **Ejub** | 0 | 2.59 | 0.00 | 1.00 | 0.0005 | 0.0014 | -66949 | -60845 | -6104 | 5479 | 99.9% | 0.0% |
| **Atro_Afor_Curs** | **Atro** | 0 | 13.37 | 0.50 | 0.50 | 0.0005 | 0.0036 | -22 | -18 | -4 | 2 | 0.0% | 0.0% |
| **Atro_Afor_Curs** | **Afor** | 0 | 0 | 0 | 0 | 0 | 0 | 0 | 0 | 0 | 0 | 0.0% | 0.0% |
| **Atro_Afor_Curs** | **Curs** | 0 | 3.79 | 0.00 | 1.00 | 0.0008 | 0.0034 | -60100 | -51259 | -8840 | 5485 | 100.0% | 0.0% |
| **Atro_Afor_Oros** | **Atro** | 0 | 0 | 0 | 0 | 0 | 0 | 0 | 0 | 0 | 0 | 0.0% | 0.0% |
| **Atro_Afor_Oros** | **Afor** | 0 | 0 | 0 | 0 | 0 | 0 | 0 | 0 | 0 | 0 | 0.0% | 0.0% |
| **Atro_Afor_Oros** | **Oros** | 0 | 5.70 | 0.00 | 1.00 | 0.0021 | 0.0125 | -49423 | -37095 | -12328 | 5487 | 100.0% | 0.0% |
| **Atro_Agal_Aaus** | **Atro** | 0 | 1.88 | 0.01 | 0.99 | 0.0005 | 0.0011 | -67441 | -63226 | -4215 | 5437 | 98.4% | 0.0% |
| **Atro_Agal_Aaus** | **Agal** | 0 | 10.05 | 0.96 | 0.04 | 0.0005 | 0.0005 | -308 | -304 | -4 | 23 | 0.0% | 0.0% |
| **Atro_Agal_Aaus** | **Aaus** | 0 | 1.97 | 0.91 | 0.09 | 0.0002 | 0.0002 | -397 | -403 | 6 | 27 | 0.0% | 0.0% |
| **Atro_Agal_Phoo** | **Atro** | 0 | 5.56 | 1.00 | 0.00 | 0.0002 | 0.0002 | -31537 | -31551 | 14 | 2088 | 0.0% | 38.0% |
| **Atro_Agal_Phoo** | **Agal** | 0 | 33.13 | 1.00 | 0.00 | 0.0001 | 0.0001 | -15167 | -15078 | -90 | 954 | 0.0% | 0.0% |
| **Atro_Agal_Phoo** | **Phoo** | 0 | 4.80 | 0.99 | 0.01 | 0.0002 | 0.0002 | -36791 | -36802 | 11 | 2445 | 0.0% | 44.3% |
| **Atro_Agal_Zwol** | **Atro** | 0 | 0.94 | 0.05 | 0.95 | 0.0005 | 0.0005 | -50 | -51 | 1 | 4 | 0.0% | 0.0% |
| **Atro_Agal_Zwol** | **Agal** | 0 | 0.64 | 0.00 | 1.00 | 0.0004 | 0.0004 | -54 | -54 | 0 | 4 | 0.0% | 0.0% |
| **Atro_Agal_Zwol** | **Zwol** | 0 | 2.60 | 0.00 | 1.00 | 0.0005 | 0.0014 | -66968 | -60853 | -6116 | 5479 | 99.9% | 0.0% |
| **Atro_Agal_Zcal** | **Atro** | 0 | 0.94 | 0.05 | 0.95 | 0.0005 | 0.0005 | -50 | -51 | 1 | 4 | 0.0% | 0.0% |
| **Atro_Agal_Zcal** | **Agal** | 0 | 0.64 | 0.00 | 1.00 | 0.0004 | 0.0004 | -54 | -54 | 0 | 4 | 0.0% | 0.0% |
| **Atro_Agal_Zcal** | **Zcal** | 0 | 2.60 | 0.00 | 1.00 | 0.0005 | 0.0014 | -66968 | -60853 | -6116 | 5479 | 99.9% | 0.0% |
| **Atro_Agal_Ejub** | **Atro** | 0 | 0.94 | 0.05 | 0.95 | 0.0005 | 0.0005 | -50 | -51 | 1 | 4 | 0.0% | 0.0% |
| **Atro_Agal_Ejub** | **Agal** | 0 | 0.64 | 0.00 | 1.00 | 0.0004 | 0.0004 | -54 | -54 | 0 | 4 | 0.0% | 0.0% |
| **Atro_Agal_Ejub** | **Ejub** | 0 | 2.60 | 0.00 | 1.00 | 0.0005 | 0.0014 | -66968 | -60853 | -6116 | 5479 | 99.9% | 0.0% |
| **Atro_Agal_Curs** | **Atro** | 0 | 10.97 | 0.50 | 0.50 | 0.0005 | 0.0030 | -22 | -19 | -3 | 2 | 0.0% | 0.0% |
| **Atro_Agal_Curs** | **Agal** | 0 | 0 | 0 | 0 | 0 | 0 | 0 | 0 | 0 | 0 | 0.0% | 0.0% |
| **Atro_Agal_Curs** | **Curs** | 0 | 3.79 | 0.00 | 1.00 | 0.0008 | 0.0034 | -60110 | -51263 | -8847 | 5485 | 100.0% | 0.0% |

| | | | | | | | | | | | | | |
|---|---|---|---|---|---|---|---|---|---|---|---|---|---|
| **Atro_Agal_Oros** | **Atro** | 0 | 0 | 0 | 0 | 0 | 0 | 0 | 0 | 0 | 0 | 0.0% | 0.0% |
| **Atro_Agal_Oros** | **Agal** | 0 | 0 | 0 | 0 | 0 | 0 | 0 | 0 | 0 | 0 | 0.0% | 0.0% |
| **Atro_Agal_Oros** | **Oros** | 0 | 5.70 | 0.00 | 1.00 | 0.0021 | 0.0125 | -49427 | -37096 | -12331 | 5487 | 100.0% | 0.0% |
| **Atro_Aaus_Phoo** | **Atro** | 0 | 5.58 | 1.00 | 0.00 | 0.0002 | 0.0002 | -31599 | -31613 | 14 | 2091 | 0.0% | 38.0% |
| **Atro_Aaus_Phoo** | **Aaus** | 0 | 4.23 | 0.99 | 0.01 | 0.0001 | 0.0001 | -15123 | -15136 | 13 | 953 | 0.0% | 17.2% |
| **Atro_Aaus_Phoo** | **Phoo** | 0 | 4.79 | 0.99 | 0.01 | 0.0002 | 0.0002 | -36754 | -36765 | 11 | 2443 | 0.0% | 44.2% |
| **Atro_Aaus_Zwol** | **Atro** | 0 | 0.94 | 0.05 | 0.95 | 0.0005 | 0.0005 | -50 | -51 | 1 | 4 | 0.0% | 0.0% |
| **Atro_Aaus_Zwol** | **Aaus** | 0 | 0.64 | 0.00 | 1.00 | 0.0004 | 0.0004 | -54 | -54 | 0 | 4 | 0.0% | 0.0% |
| **Atro_Aaus_Zwol** | **Zwol** | 0 | 2.60 | 0.00 | 1.00 | 0.0005 | 0.0014 | -66942 | -60845 | -6097 | 5479 | 99.9% | 0.0% |
| **Atro_Aaus_Zcal** | **Atro** | 0 | 0.94 | 0.05 | 0.95 | 0.0005 | 0.0005 | -50 | -51 | 1 | 4 | 0.0% | 0.0% |
| **Atro_Aaus_Zcal** | **Aaus** | 0 | 0.64 | 0.00 | 1.00 | 0.0004 | 0.0004 | -54 | -54 | 0 | 4 | 0.0% | 0.0% |
| **Atro_Aaus_Zcal** | **Zcal** | 0 | 2.60 | 0.00 | 1.00 | 0.0005 | 0.0014 | -66942 | -60845 | -6097 | 5479 | 99.9% | 0.0% |
| **Atro_Aaus_Ejub** | **Atro** | 0 | 0.94 | 0.05 | 0.95 | 0.0005 | 0.0005 | -50 | -51 | 1 | 4 | 0.0% | 0.0% |
| **Atro_Aaus_Ejub** | **Aaus** | 0 | 0.64 | 0.00 | 1.00 | 0.0004 | 0.0004 | -54 | -54 | 0 | 4 | 0.0% | 0.0% |
| **Atro_Aaus_Ejub** | **Ejub** | 0 | 2.60 | 0.00 | 1.00 | 0.0005 | 0.0014 | -66942 | -60845 | -6097 | 5479 | 99.9% | 0.0% |
| **Atro_Aaus_Curs** | **Atro** | 0 | 10.97 | 0.50 | 0.50 | 0.0005 | 0.0030 | -22 | -19 | -3 | 2 | 0.0% | 0.0% |
| **Atro_Aaus_Curs** | **Aaus** | 0 | 0 | 0 | 0 | 0 | 0 | 0 | 0 | 0 | 0 | 0.0% | 0.0% |
| **Atro_Aaus_Curs** | **Curs** | 0 | 3.79 | 0.00 | 1.00 | 0.0008 | 0.0034 | -60096 | -51259 | -8837 | 5485 | 100.0% | 0.0% |
| **Atro_Aaus_Oros** | **Atro** | 0 | 0 | 0 | 0 | 0 | 0 | 0 | 0 | 0 | 0 | 0.0% | 0.0% |
| **Atro_Aaus_Oros** | **Aaus** | 0 | 0 | 0 | 0 | 0 | 0 | 0 | 0 | 0 | 0 | 0.0% | 0.0% |
| **Atro_Aaus_Oros** | **Oros** | 0 | 5.70 | 0.00 | 1.00 | 0.0021 | 0.0125 | -49423 | -37095 | -12328 | 5487 | 100.0% | 0.0% |
| **Atro_Phoo_Zwol** | **Atro** | 0 | 1.47 | 0.37 | 0.63 | 0.0005 | 0.0006 | -86 | -88 | 2 | 7 | 0.0% | 0.0% |
| **Atro_Phoo_Zwol** | **Phoo** | 0 | 0.74 | 0.50 | 0.50 | 0.0003 | 0.0003 | -107 | -111 | 3 | 8 | 0.0% | 0.0% |
| **Atro_Phoo_Zwol** | **Zwol** | 0 | 2.48 | 0.00 | 1.00 | 0.0005 | 0.0014 | -67154 | -61268 | -5886 | 5472 | 99.7% | 0.0% |
| **Atro_Phoo_Zcal** | **Atro** | 0 | 1.47 | 0.37 | 0.63 | 0.0005 | 0.0006 | -86 | -88 | 2 | 7 | 0.0% | 0.0% |
| **Atro_Phoo_Zcal** | **Phoo** | 0 | 0.74 | 0.50 | 0.50 | 0.0003 | 0.0003 | -107 | -111 | 3 | 8 | 0.0% | 0.0% |
| **Atro_Phoo_Zcal** | **Zcal** | 0 | 2.48 | 0.00 | 1.00 | 0.0005 | 0.0014 | -67154 | -61268 | -5886 | 5472 | 99.7% | 0.0% |
| **Atro_Phoo_Ejub** | **Atro** | 0 | 1.47 | 0.37 | 0.63 | 0.0005 | 0.0006 | -86 | -88 | 2 | 7 | 0.0% | 0.0% |
| **Atro_Phoo_Ejub** | **Phoo** | 0 | 0.74 | 0.50 | 0.50 | 0.0003 | 0.0003 | -107 | -111 | 3 | 8 | 0.0% | 0.0% |
| **Atro_Phoo_Ejub** | **Ejub** | 0 | 2.48 | 0.00 | 1.00 | 0.0005 | 0.0014 | -67154 | -61268 | -5886 | 5472 | 99.7% | 0.0% |
| **Atro_Phoo_Curs** | **Atro** | 0 | 0 | 0 | 0 | 0 | 0 | 0 | 0 | 0 | 1 | 0.0% | 0.0% |
| **Atro_Phoo_Curs** | **Phoo** | 0 | 0 | 0 | 0 | 0 | 0 | 0 | 0 | 0 | 1 | 0.0% | 0.0% |

| | | | | | | | | | | | | | |
|---|---|---|---|---|---|---|---|---|---|---|---|---|---|
| **Atro_Phoo_Curs** | **Curs** | 0 | 3.77 | 0.00 | 1.00 | 0.0008 | 0.0034 | -60258 | -51471 | -8787 | 5485 | 100.0% | 0.0% |
| **Atro_Phoo_Oros** | **Atro** | 0 | 0 | 0 | 0 | 0 | 0 | 0 | 0 | 0 | 0 | 0.0% | 0.0% |
| **Atro_Phoo_Oros** | **Phoo** | 0 | 0 | 0 | 0 | 0 | 0 | 0 | 0 | 0 | 0 | 0.0% | 0.0% |
| **Atro_Phoo_Oros** | **Oros** | 0 | 5.71 | 0.00 | 1.00 | 0.0021 | 0.0124 | -49505 | -37153 | -12352 | 5487 | 100.0% | 0.0% |
| **Atro_Zwol_Zcal** | **Atro** | 0 | 3.97 | 0.00 | 1.00 | 0.0006 | 0.0026 | -63343 | -54296 | -9047 | 5487 | 100.0% | 0.0% |
| **Atro_Zwol_Zcal** | **Zwol** | 0 | 0 | 0 | 0 | 0 | 0 | 0 | 0 | 0 | 0 | 0.0% | 0.0% |
| **Atro_Zwol_Zcal** | **Zcal** | 0 | 0 | 0 | 0 | 0 | 0 | 0 | 0 | 0 | 0 | 0.0% | 0.0% |
| **Atro_Zwol_Ejub** | **Atro** | 0 | 3.98 | 0.00 | 1.00 | 0.0006 | 0.0026 | -63346 | -54308 | -9038 | 5487 | 100.0% | 0.0% |
| **Atro_Zwol_Ejub** | **Zwol** | 0 | 0 | 0 | 0 | 0 | 0 | 0 | 0 | 0 | 0 | 0.0% | 0.0% |
| **Atro_Zwol_Ejub** | **Ejub** | 0 | 0 | 0 | 0 | 0 | 0 | 0 | 0 | 0 | 0 | 0.0% | 0.0% |
| **Atro_Zwol_Curs** | **Atro** | 0 | 0 | 0 | 0 | 0 | 0 | 0 | 0 | 0 | 1 | 0.0% | 0.0% |
| **Atro_Zwol_Curs** | **Zwol** | 0 | 1.56 | 0.04 | 0.96 | 0.0005 | 0.0006 | -37 | -37 | 0 | 3 | 0.0% | 0.0% |
| **Atro_Zwol_Curs** | **Curs** | 0 | 3.13 | 0.00 | 1.00 | 0.0006 | 0.0020 | -64468 | -57101 | -7366 | 5483 | 99.9% | 0.0% |
| **Atro_Zwol_Oros** | **Atro** | 0 | 0 | 0 | 0 | 0 | 0 | 0 | 0 | 0 | 0 | 0.0% | 0.0% |
| **Atro_Zwol_Oros** | **Zwol** | 0 | 0 | 0 | 0 | 0 | 0 | 0 | 0 | 0 | 0 | 0.0% | 0.0% |
| **Atro_Zwol_Oros** | **Oros** | 0 | 5.89 | 0.00 | 1.00 | 0.0018 | 0.0111 | -51054 | -38420 | -12634 | 5487 | 100.0% | 0.0% |
| **Atro_Zcal_Ejub** | **Atro** | 0 | 3.97 | 0.00 | 1.00 | 0.0006 | 0.0027 | -63112 | -54047 | -9065 | 5487 | 100.0% | 0.0% |
| **Atro_Zcal_Ejub** | **Zcal** | 0 | 0 | 0 | 0 | 0 | 0 | 0 | 0 | 0 | 0 | 0.0% | 0.0% |
| **Atro_Zcal_Ejub** | **Ejub** | 0 | 0 | 0 | 0 | 0 | 0 | 0 | 0 | 0 | 0 | 0.0% | 0.0% |
| **Atro_Zcal_Curs** | **Atro** | 0 | 0 | 0 | 0 | 0 | 0 | 0 | 0 | 0 | 1 | 0.0% | 0.0% |
| **Atro_Zcal_Curs** | **Zcal** | 0 | 1.56 | 0.04 | 0.96 | 0.0005 | 0.0006 | -37 | -37 | 0 | 3 | 0.0% | 0.0% |
| **Atro_Zcal_Curs** | **Curs** | 0 | 3.13 | 0.00 | 1.00 | 0.0006 | 0.0020 | -64468 | -57101 | -7366 | 5483 | 99.9% | 0.0% |
| **Atro_Zcal_Oros** | **Atro** | 0 | 0 | 0 | 0 | 0 | 0 | 0 | 0 | 0 | 0 | 0.0% | 0.0% |
| **Atro_Zcal_Oros** | **Zcal** | 0 | 0 | 0 | 0 | 0 | 0 | 0 | 0 | 0 | 0 | 0.0% | 0.0% |
| **Atro_Zcal_Oros** | **Oros** | 0 | 5.89 | 0.00 | 1.00 | 0.0018 | 0.0111 | -51054 | -38420 | -12634 | 5487 | 100.0% | 0.0% |
| **Atro_Ejub_Curs** | **Atro** | 0 | 0 | 0 | 0 | 0 | 0 | 0 | 0 | 0 | 1 | 0.0% | 0.0% |
| **Atro_Ejub_Curs** | **Ejub** | 0 | 1.56 | 0.04 | 0.96 | 0.0005 | 0.0006 | -37 | -37 | 0 | 3 | 0.0% | 0.0% |
| **Atro_Ejub_Curs** | **Curs** | 0 | 3.13 | 0.00 | 1.00 | 0.0006 | 0.0020 | -64468 | -57101 | -7366 | 5483 | 99.9% | 0.0% |
| **Atro_Ejub_Oros** | **Atro** | 0 | 0 | 0 | 0 | 0 | 0 | 0 | 0 | 0 | 0 | 0.0% | 0.0% |
| **Atro_Ejub_Oros** | **Ejub** | 0 | 0 | 0 | 0 | 0 | 0 | 0 | 0 | 0 | 0 | 0.0% | 0.0% |
| **Atro_Ejub_Oros** | **Oros** | 0 | 5.89 | 0.00 | 1.00 | 0.0018 | 0.0111 | -51054 | -38420 | -12634 | 5487 | 100.0% | 0.0% |

| | | | | | | | | | | | | | |
|---|---|---|---|---|---|---|---|---|---|---|---|---|---|
| **Atro_Curs_Oros** | **Atro** | 0 | 0 | 0 | 0 | 0 | 0 | 0 | 0 | 0 | 0 | 0.0% | 0.0% |
| **Atro_Curs_Oros** | **Curs** | 0 | 0 | 0 | 0 | 0 | 0 | 0 | 0 | 0 | 0 | 0.0% | 0.0% |
| **Atro_Curs_Oros** | **Oros** | 0 | 6.17 | 0.00 | 1.00 | 0.0014 | 0.0091 | -53696 | -40616 | -13080 | 5487 | 100.0% | 0.0% |
| **Ofla_Atow_Aphi** | **Ofla** | 0 | 2.05 | 0.00 | 1.00 | 0.0005 | 0.0012 | -68474 | -62787 | -5687 | 5482 | 99.9% | 0.0% |
| **Ofla_Atow_Aphi** | **Atow** | 0 | 0.66 | 0.00 | 1.00 | 0.0005 | 0.0005 | -26 | -26 | 0 | 2 | 0.0% | 0.0% |
| **Ofla_Atow_Aphi** | **Aphi** | 0 | 2.81 | 0.74 | 0.26 | 0.0005 | 0.0006 | -37 | -38 | 1 | 3 | 0.0% | 0.0% |
| **Ofla_Atow_Agaz** | **Ofla** | 0 | 6.89 | 1.00 | 0.00 | 0.0003 | 0.0003 | -49560 | -49576 | 16 | 3513 | 0.0% | 64.0% |
| **Ofla_Atow_Agaz** | **Atow** | 0 | 7.21 | 0.99 | 0.01 | 0.0002 | 0.0002 | -12996 | -13002 | 6 | 850 | 0.0% | 0.0% |
| **Ofla_Atow_Agaz** | **Agaz** | 0 | 6.11 | 1.00 | 0.00 | 0.0002 | 0.0002 | -16945 | -16959 | 14 | 1124 | 0.0% | 20.5% |
| **Ofla_Atow_Afor** | **Ofla** | 0 | 7.13 | 1.00 | 0.00 | 0.0003 | 0.0003 | -51591 | -51607 | 16 | 3656 | 0.0% | 66.6% |
| **Ofla_Atow_Afor** | **Atow** | 0 | 4.53 | 0.99 | 0.01 | 0.0001 | 0.0001 | -9645 | -9655 | 10 | 617 | 0.0% | 0.0% |
| **Ofla_Atow_Afor** | **Afor** | 0 | 6.09 | 1.00 | 0.00 | 0.0002 | 0.0002 | -18294 | -18307 | 13 | 1214 | 0.0% | 22.1% |
| **Ofla_Atow_Agal** | **Ofla** | 0 | 6.73 | 1.00 | 0.00 | 0.0003 | 0.0003 | -51650 | -51666 | 16 | 3664 | 0.0% | 66.8% |
| **Ofla_Atow_Agal** | **Atow** | 0 | 4.83 | 0.99 | 0.01 | 0.0001 | 0.0001 | -9675 | -9685 | 10 | 614 | 0.0% | 0.0% |
| **Ofla_Atow_Agal** | **Agal** | 0 | 6.23 | 1.00 | 0.00 | 0.0002 | 0.0002 | -18218 | -18230 | 12 | 1209 | 0.0% | 22.0% |
| **Ofla_Atow_Aaus** | **Ofla** | 0 | 6.70 | 1.00 | 0.00 | 0.0003 | 0.0003 | -51514 | -51531 | 16 | 3656 | 0.0% | 66.6% |
| **Ofla_Atow_Aaus** | **Atow** | 0 | 4.55 | 0.99 | 0.01 | 0.0001 | 0.0001 | -9784 | -9794 | 10 | 622 | 0.0% | 11.2% |
| **Ofla_Atow_Aaus** | **Aaus** | 0 | 6.09 | 1.00 | 0.00 | 0.0002 | 0.0002 | -18219 | -18232 | 13 | 1209 | 0.0% | 22.0% |
| **Ofla_Atow_Phoo** | **Ofla** | 0 | 6.87 | 1.00 | 0.00 | 0.0002 | 0.0002 | -38388 | -38402 | 14 | 2610 | 0.0% | 47.5% |
| **Ofla_Atow_Phoo** | **Atow** | 0 | 4.37 | 0.99 | 0.01 | 0.0002 | 0.0002 | -17308 | -17319 | 11 | 1128 | 0.0% | 20.4% |
| **Ofla_Atow_Phoo** | **Phoo** | 0 | 5.78 | 1.00 | 0.00 | 0.0002 | 0.0002 | -26412 | -26424 | 12 | 1749 | 0.0% | 31.8% |
| **Ofla_Atow_Zwol** | **Ofla** | 0 | 0.92 | 0.00 | 1.00 | 0.0002 | 0.0002 | -73 | -72 | -1 | 5 | 0.0% | 0.0% |
| **Ofla_Atow_Zwol** | **Atow** | 0 | 1.17 | 0.35 | 0.65 | 0.0005 | 0.0007 | -144 | -148 | 4 | 12 | 0.0% | 0.0% |
| **Ofla_Atow_Zwol** | **Zwol** | 0 | 2.45 | 0.00 | 1.00 | 0.0005 | 0.0013 | -67312 | -61366 | -5947 | 5470 | 99.7% | 0.0% |
| **Ofla_Atow_Zcal** | **Ofla** | 0 | 0.92 | 0.00 | 1.00 | 0.0002 | 0.0002 | -73 | -72 | -1 | 5 | 0.0% | 0.0% |
| **Ofla_Atow_Zcal** | **Atow** | 0 | 1.17 | 0.35 | 0.65 | 0.0005 | 0.0007 | -144 | -148 | 4 | 12 | 0.0% | 0.0% |
| **Ofla_Atow_Zcal** | **Zcal** | 0 | 2.45 | 0.00 | 1.00 | 0.0005 | 0.0013 | -67312 | -61366 | -5947 | 5470 | 99.7% | 0.0% |
| **Ofla_Atow_Ejub** | **Ofla** | 0 | 0.92 | 0.00 | 1.00 | 0.0002 | 0.0002 | -73 | -72 | -1 | 5 | 0.0% | 0.0% |
| **Ofla_Atow_Ejub** | **Atow** | 0 | 1.17 | 0.35 | 0.65 | 0.0005 | 0.0007 | -144 | -148 | 4 | 12 | 0.0% | 0.0% |
| **Ofla_Atow_Ejub** | **Ejub** | 0 | 2.45 | 0.00 | 1.00 | 0.0005 | 0.0013 | -67312 | -61366 | -5947 | 5470 | 99.7% | 0.0% |
| **Ofla_Atow_Curs** | **Ofla** | 0 | 0 | 0 | 0 | 0 | 0 | 0 | 0 | 0 | 0 | 0.0% | 0.0% |
| **Ofla_Atow_Curs** | **Atow** | 0 | 4.74 | 0.49 | 0.51 | 0.0005 | 0.0015 | -22 | -21 | -1 | 2 | 0.0% | 0.0% |

| | | | | | | | | | | | | | |
|---|---|---|---|---|---|---|---|---|---|---|---|---|---|
| **Ofla_Atow_Curs** | **Curs** | 0 | 3.80 | 0.00 | 1.00 | 0.0008 | 0.0034 | -60341 | -51520 | -8821 | 5485 | 100.0% | 0.0% |
| **Ofla_Atow_Oros** | **Ofla** | 0 | 0 | 0 | 0 | 0 | 0 | 0 | 0 | 0 | 0 | 0.0% | 0.0% |
| **Ofla_Atow_Oros** | **Atow** | 0 | 0 | 0 | 0 | 0 | 0 | 0 | 0 | 0 | 0 | 0.0% | 0.0% |
| **Ofla_Atow_Oros** | **Oros** | 0 | 5.76 | 0.00 | 1.00 | 0.0020 | 0.0124 | -49567 | -37166 | -12401 | 5487 | 100.0% | 0.0% |
| **Ofla_Aphi_Agaz** | **Ofla** | 0 | 6.90 | 1.00 | 0.00 | 0.0003 | 0.0003 | -49553 | -49569 | 16 | 3512 | 0.0% | 64.0% |
| **Ofla_Aphi_Agaz** | **Aphi** | 0 | 7.21 | 0.99 | 0.01 | 0.0002 | 0.0002 | -13011 | -13017 | 6 | 851 | 0.0% | 0.0% |
| **Ofla_Aphi_Agaz** | **Agaz** | 0 | 6.13 | 1.00 | 0.00 | 0.0002 | 0.0002 | -16951 | -16965 | 14 | 1124 | 0.0% | 20.5% |
| **Ofla_Aphi_Afor** | **Ofla** | 0 | 6.78 | 1.00 | 0.00 | 0.0003 | 0.0003 | -51600 | -51616 | 16 | 3656 | 0.0% | 66.6% |
| **Ofla_Aphi_Afor** | **Aphi** | 0 | 4.57 | 0.99 | 0.01 | 0.0001 | 0.0001 | -9669 | -9679 | 10 | 618 | 0.0% | 0.0% |
| **Ofla_Aphi_Afor** | **Afor** | 0 | 6.11 | 1.00 | 0.00 | 0.0002 | 0.0002 | -18289 | -18303 | 14 | 1213 | 0.0% | 22.1% |
| **Ofla_Aphi_Agal** | **Ofla** | 0 | 6.73 | 1.00 | 0.00 | 0.0003 | 0.0003 | -51639 | -51656 | 16 | 3663 | 0.0% | 66.8% |
| **Ofla_Aphi_Agal** | **Aphi** | 0 | 4.83 | 0.99 | 0.01 | 0.0001 | 0.0001 | -9690 | -9700 | 10 | 615 | 0.0% | 0.0% |
| **Ofla_Aphi_Agal** | **Agal** | 0 | 6.09 | 1.00 | 0.00 | 0.0002 | 0.0002 | -18223 | -18236 | 13 | 1209 | 0.0% | 22.0% |
| **Ofla_Aphi_Aaus** | **Ofla** | 0 | 6.72 | 1.00 | 0.00 | 0.0003 | 0.0003 | -51546 | -51562 | 16 | 3657 | 0.0% | 66.6% |
| **Ofla_Aphi_Aaus** | **Aphi** | 0 | 4.79 | 0.99 | 0.01 | 0.0001 | 0.0001 | -9791 | -9801 | 10 | 622 | 0.0% | 11.2% |
| **Ofla_Aphi_Aaus** | **Aaus** | 0 | 6.11 | 1.00 | 0.00 | 0.0002 | 0.0002 | -18214 | -18228 | 14 | 1208 | 0.0% | 22.0% |
| **Ofla_Aphi_Phoo** | **Ofla** | 0 | 6.92 | 1.00 | 0.00 | 0.0002 | 0.0002 | -38429 | -38444 | 15 | 2610 | 0.0% | 47.5% |
| **Ofla_Aphi_Phoo** | **Aphi** | 0 | 4.36 | 0.99 | 0.01 | 0.0002 | 0.0002 | -17305 | -17317 | 12 | 1128 | 0.0% | 20.4% |
| **Ofla_Aphi_Phoo** | **Phoo** | 0 | 5.79 | 1.00 | 0.00 | 0.0002 | 0.0002 | -26417 | -26430 | 13 | 1749 | 0.0% | 31.8% |
| **Ofla_Aphi_Zwol** | **Ofla** | 0 | 0.92 | 0.00 | 1.00 | 0.0002 | 0.0002 | -73 | -72 | -1 | 5 | 0.0% | 0.0% |
| **Ofla_Aphi_Zwol** | **Aphi** | 0 | 1.17 | 0.35 | 0.65 | 0.0005 | 0.0007 | -144 | -148 | 4 | 12 | 0.0% | 0.0% |
| **Ofla_Aphi_Zwol** | **Zwol** | 0 | 2.45 | 0.00 | 1.00 | 0.0005 | 0.0013 | -67315 | -61367 | -5948 | 5470 | 99.7% | 0.0% |
| **Ofla_Aphi_Zcal** | **Ofla** | 0 | 0.92 | 0.00 | 1.00 | 0.0002 | 0.0002 | -73 | -72 | -1 | 5 | 0.0% | 0.0% |
| **Ofla_Aphi_Zcal** | **Aphi** | 0 | 1.17 | 0.35 | 0.65 | 0.0005 | 0.0007 | -144 | -148 | 4 | 12 | 0.0% | 0.0% |
| **Ofla_Aphi_Zcal** | **Zcal** | 0 | 2.45 | 0.00 | 1.00 | 0.0005 | 0.0013 | -67315 | -61367 | -5948 | 5470 | 99.7% | 0.0% |
| **Ofla_Aphi_Ejub** | **Ofla** | 0 | 0.92 | 0.00 | 1.00 | 0.0002 | 0.0002 | -73 | -72 | -1 | 5 | 0.0% | 0.0% |
| **Ofla_Aphi_Ejub** | **Aphi** | 0 | 1.17 | 0.35 | 0.65 | 0.0005 | 0.0007 | -144 | -148 | 4 | 12 | 0.0% | 0.0% |
| **Ofla_Aphi_Ejub** | **Ejub** | 0 | 2.45 | 0.00 | 1.00 | 0.0005 | 0.0013 | -67315 | -61367 | -5948 | 5470 | 99.7% | 0.0% |
| **Ofla_Aphi_Curs** | **Ofla** | 0 | 0 | 0 | 0 | 0 | 0 | 0 | 0 | 0 | 0 | 0.0% | 0.0% |
| **Ofla_Aphi_Curs** | **Aphi** | 0 | 4.74 | 0.49 | 0.51 | 0.0005 | 0.0015 | -22 | -21 | -1 | 2 | 0.0% | 0.0% |
| **Ofla_Aphi_Curs** | **Curs** | 0 | 3.80 | 0.00 | 1.00 | 0.0008 | 0.0034 | -60344 | -51521 | -8823 | 5485 | 100.0% | 0.0% |
| **Ofla_Aphi_Oros** | **Ofla** | 0 | 0 | 0 | 0 | 0 | 0 | 0 | 0 | 0 | 0 | 0.0% | 0.0% |

| | | | | | | | | | | | | | |
|---|---|---|---|---|---|---|---|---|---|---|---|---|---|
| **Ofla_Aphi_Oros** | **Aphi** | 0 | 0 | 0 | 0 | 0 | 0 | 0 | 0 | 0 | 0 | 0.0% | 0.0% |
| **Ofla_Aphi_Oros** | **Oros** | 0 | 5.76 | 0.00 | 1.00 | 0.0020 | 0.0124 | -49568 | -37166 | -12402 | 5487 | 100.0% | 0.0% |
| **Ofla_Agaz_Afor** | **Ofla** | 0 | 6.09 | 1.00 | 0.00 | 0.0003 | 0.0003 | -56263 | -56279 | 17 | 4043 | 0.0% | 73.7% |
| **Ofla_Agaz_Afor** | **Agaz** | 0 | 4.39 | 0.99 | 0.01 | 0.0002 | 0.0002 | -8740 | -8751 | 11 | 563 | 0.0% | 10.2% |
| **Ofla_Agaz_Afor** | **Afor** | 0 | 6.79 | 1.00 | 0.00 | 0.0002 | 0.0002 | -13361 | -13371 | 10 | 881 | 0.0% | 16.0% |
| **Ofla_Agaz_Agal** | **Ofla** | 0 | 6.34 | 1.00 | 0.00 | 0.0004 | 0.0004 | -56220 | -56236 | 17 | 4044 | 0.0% | 73.7% |
| **Ofla_Agaz_Agal** | **Agaz** | 0 | 4.45 | 0.99 | 0.01 | 0.0001 | 0.0001 | -8792 | -8802 | 11 | 562 | 0.0% | 10.1% |
| **Ofla_Agaz_Agal** | **Agal** | 0 | 4.61 | 1.00 | 0.00 | 0.0002 | 0.0002 | -13357 | -13370 | 13 | 881 | 0.0% | 16.0% |
| **Ofla_Agaz_Aaus** | **Ofla** | 0 | 6.36 | 1.00 | 0.00 | 0.0004 | 0.0004 | -56268 | -56285 | 17 | 4046 | 0.0% | 73.7% |
| **Ofla_Agaz_Aaus** | **Agaz** | 0 | 4.38 | 0.99 | 0.01 | 0.0001 | 0.0001 | -8846 | -8858 | 11 | 566 | 0.0% | 10.2% |
| **Ofla_Agaz_Aaus** | **Aaus** | 0 | 6.80 | 1.00 | 0.00 | 0.0002 | 0.0002 | -13272 | -13282 | 10 | 875 | 0.0% | 0.0% |
| **Ofla_Agaz_Phoo** | **Ofla** | 0 | 6.09 | 1.00 | 0.00 | 0.0002 | 0.0002 | -41987 | -42003 | 16 | 2861 | 0.0% | 52.1% |
| **Ofla_Agaz_Phoo** | **Agaz** | 0 | 7.19 | 0.99 | 0.01 | 0.0002 | 0.0002 | -18101 | -18100 | -1 | 1174 | 0.0% | 0.0% |
| **Ofla_Agaz_Phoo** | **Phoo** | 0 | 5.22 | 1.00 | 0.00 | 0.0002 | 0.0002 | -22038 | -22051 | 14 | 1452 | 0.0% | 26.4% |
| **Ofla_Agaz_Zwol** | **Ofla** | 0 | 0.92 | 0.00 | 1.00 | 0.0002 | 0.0002 | -73 | -72 | -1 | 5 | 0.0% | 0.0% |
| **Ofla_Agaz_Zwol** | **Agaz** | 0 | 0.51 | 0.00 | 1.00 | 0.0005 | 0.0006 | -161 | -162 | 1 | 13 | 0.0% | 0.0% |
| **Ofla_Agaz_Zwol** | **Zwol** | 0 | 2.43 | 0.00 | 1.00 | 0.0005 | 0.0013 | -67345 | -61456 | -5888 | 5469 | 99.7% | 0.0% |
| **Ofla_Agaz_Zcal** | **Ofla** | 0 | 0.92 | 0.00 | 1.00 | 0.0002 | 0.0002 | -73 | -72 | -1 | 5 | 0.0% | 0.0% |
| **Ofla_Agaz_Zcal** | **Agaz** | 0 | 0.51 | 0.00 | 1.00 | 0.0005 | 0.0006 | -161 | -162 | 1 | 13 | 0.0% | 0.0% |
| **Ofla_Agaz_Zcal** | **Zcal** | 0 | 2.43 | 0.00 | 1.00 | 0.0005 | 0.0013 | -67345 | -61456 | -5888 | 5469 | 99.7% | 0.0% |
| **Ofla_Agaz_Ejub** | **Ofla** | 0 | 0.92 | 0.00 | 1.00 | 0.0002 | 0.0002 | -73 | -72 | -1 | 5 | 0.0% | 0.0% |
| **Ofla_Agaz_Ejub** | **Agaz** | 0 | 0.51 | 0.00 | 1.00 | 0.0005 | 0.0006 | -161 | -162 | 1 | 13 | 0.0% | 0.0% |
| **Ofla_Agaz_Ejub** | **Ejub** | 0 | 2.43 | 0.00 | 1.00 | 0.0005 | 0.0013 | -67345 | -61456 | -5888 | 5469 | 99.7% | 0.0% |
| **Ofla_Agaz_Curs** | **Ofla** | 0 | 0 | 0 | 0 | 0 | 0 | 0 | 0 | 0 | 1 | 0.0% | 0.0% |
| **Ofla_Agaz_Curs** | **Agaz** | 0 | 0 | 0 | 0 | 0 | 0 | 0 | 0 | 0 | 1 | 0.0% | 0.0% |
| **Ofla_Agaz_Curs** | **Curs** | 0 | 3.79 | 0.00 | 1.00 | 0.0008 | 0.0033 | -60384 | -51562 | -8823 | 5485 | 100.0% | 0.0% |
| **Ofla_Agaz_Oros** | **Ofla** | 0 | 0 | 0 | 0 | 0 | 0 | 0 | 0 | 0 | 0 | 0.0% | 0.0% |
| **Ofla_Agaz_Oros** | **Agaz** | 0 | 0 | 0 | 0 | 0 | 0 | 0 | 0 | 0 | 0 | 0.0% | 0.0% |
| **Ofla_Agaz_Oros** | **Oros** | 0 | 5.76 | 0.00 | 1.00 | 0.0020 | 0.0124 | -49572 | -37177 | -12395 | 5487 | 100.0% | 0.0% |
| **Ofla_Afor_Agal** | **Ofla** | 0 | 1.35 | 0.01 | 0.99 | 0.0005 | 0.0009 | -68531 | -65312 | -3218 | 5425 | 97.8% | 0.0% |
| **Ofla_Afor_Agal** | **Afor** | 0 | 2.08 | 0.91 | 0.09 | 0.0002 | 0.0002 | -486 | -492 | 6 | 32 | 0.0% | 0.0% |
| **Ofla_Afor_Agal** | **Agal** | 0 | 5.59 | 0.94 | 0.06 | 0.0003 | 0.0003 | -414 | -415 | 1 | 30 | 0.0% | 0.0% |

| | | | | | | | | | | | | | |
|---|---|---|---|---|---|---|---|---|---|---|---|---|---|
| **Ofla_Afor_Aaus** | **Ofla** | 0 | 1.38 | 0.01 | 0.99 | 0.0005 | 0.0009 | -68539 | -65309 | -3231 | 5435 | 98.0% | 0.0% |
| **Ofla_Afor_Aaus** | **Afor** | 0 | 2.55 | 0.95 | 0.05 | 0.0001 | 0.0001 | -463 | -470 | 7 | 30 | 0.0% | 0.0% |
| **Ofla_Afor_Aaus** | **Aaus** | 0 | 8.09 | 0.95 | 0.05 | 0.0003 | 0.0003 | -309 | -309 | -1 | 22 | 0.0% | 0.0% |
| **Ofla_Afor_Phoo** | **Ofla** | 0 | 5.77 | 1.00 | 0.00 | 0.0002 | 0.0002 | -43585 | -43600 | 15 | 2969 | 0.0% | 54.0% |
| **Ofla_Afor_Phoo** | **Afor** | 0 | 5.02 | 0.99 | 0.01 | 0.0002 | 0.0002 | -19331 | -19334 | 3 | 1251 | 0.0% | 0.0% |
| **Ofla_Afor_Phoo** | **Phoo** | 0 | 4.26 | 0.99 | 0.01 | 0.0002 | 0.0002 | -19620 | -19633 | 13 | 1267 | 0.0% | 22.9% |
| **Ofla_Afor_Zwol** | **Ofla** | 0 | 0.92 | 0.00 | 1.00 | 0.0002 | 0.0002 | -73 | -72 | -1 | 5 | 0.0% | 0.0% |
| **Ofla_Afor_Zwol** | **Afor** | 0 | 0.47 | 0.00 | 1.00 | 0.0005 | 0.0006 | -192 | -191 | 0 | 15 | 0.0% | 0.0% |
| **Ofla_Afor_Zwol** | **Zwol** | 0 | 2.41 | 0.00 | 1.00 | 0.0005 | 0.0013 | -67411 | -61545 | -5866 | 5467 | 99.6% | 0.0% |
| **Ofla_Afor_Zcal** | **Ofla** | 0 | 0.92 | 0.00 | 1.00 | 0.0002 | 0.0002 | -73 | -72 | -1 | 5 | 0.0% | 0.0% |
| **Ofla_Afor_Zcal** | **Afor** | 0 | 0.47 | 0.00 | 1.00 | 0.0005 | 0.0006 | -192 | -191 | 0 | 15 | 0.0% | 0.0% |
| **Ofla_Afor_Zcal** | **Zcal** | 0 | 2.41 | 0.00 | 1.00 | 0.0005 | 0.0013 | -67411 | -61545 | -5866 | 5467 | 99.6% | 0.0% |
| **Ofla_Afor_Ejub** | **Ofla** | 0 | 0.92 | 0.00 | 1.00 | 0.0002 | 0.0002 | -73 | -72 | -1 | 5 | 0.0% | 0.0% |
| **Ofla_Afor_Ejub** | **Afor** | 0 | 0.47 | 0.00 | 1.00 | 0.0005 | 0.0006 | -192 | -191 | 0 | 15 | 0.0% | 0.0% |
| **Ofla_Afor_Ejub** | **Ejub** | 0 | 2.41 | 0.00 | 1.00 | 0.0005 | 0.0013 | -67411 | -61545 | -5866 | 5467 | 99.6% | 0.0% |
| **Ofla_Afor_Curs** | **Ofla** | 0 | 0 | 0 | 0 | 0 | 0 | 0 | 0 | 0 | 1 | 0.0% | 0.0% |
| **Ofla_Afor_Curs** | **Afor** | 0 | 0 | 0 | 0 | 0 | 0 | 0 | 0 | 0 | 1 | 0.0% | 0.0% |
| **Ofla_Afor_Curs** | **Curs** | 0 | 3.79 | 0.00 | 1.00 | 0.0008 | 0.0033 | -60411 | -51608 | -8803 | 5485 | 100.0% | 0.0% |
| **Ofla_Afor_Oros** | **Ofla** | 0 | 0 | 0 | 0 | 0 | 0 | 0 | 0 | 0 | 0 | 0.0% | 0.0% |
| **Ofla_Afor_Oros** | **Afor** | 0 | 0 | 0 | 0 | 0 | 0 | 0 | 0 | 0 | 0 | 0.0% | 0.0% |
| **Ofla_Afor_Oros** | **Oros** | 0 | 5.74 | 0.00 | 1.00 | 0.0020 | 0.0124 | -49573 | -37189 | -12384 | 5487 | 100.0% | 0.0% |
| **Ofla_Agal_Aaus** | **Ofla** | 0 | 1.90 | 0.00 | 1.00 | 0.0005 | 0.0012 | -66723 | -62409 | -4314 | 5452 | 99.0% | 0.0% |
| **Ofla_Agal_Aaus** | **Agal** | 0 | 3.16 | 0.95 | 0.05 | 0.0003 | 0.0003 | -297 | -303 | 6 | 21 | 0.0% | 0.0% |
| **Ofla_Agal_Aaus** | **Aaus** | 0 | 1.00 | 0.76 | 0.24 | 0.0002 | 0.0002 | -203 | -208 | 5 | 14 | 0.0% | 0.0% |
| **Ofla_Agal_Phoo** | **Ofla** | 0 | 5.57 | 1.00 | 0.00 | 0.0002 | 0.0002 | -43573 | -43589 | 15 | 2967 | 0.0% | 54.0% |
| **Ofla_Agal_Phoo** | **Agal** | 0 | 5.00 | 0.99 | 0.01 | 0.0002 | 0.0002 | -19277 | -19281 | 3 | 1250 | 0.0% | 0.0% |
| **Ofla_Agal_Phoo** | **Phoo** | 0 | 4.32 | 0.99 | 0.01 | 0.0002 | 0.0002 | -19702 | -19715 | 13 | 1270 | 0.0% | 23.0% |
| **Ofla_Agal_Zwol** | **Ofla** | 0 | 0.92 | 0.00 | 1.00 | 0.0002 | 0.0002 | -73 | -72 | -1 | 5 | 0.0% | 0.0% |
| **Ofla_Agal_Zwol** | **Agal** | 0 | 0.47 | 0.00 | 1.00 | 0.0005 | 0.0006 | -192 | -191 | 0 | 15 | 0.0% | 0.0% |
| **Ofla_Agal_Zwol** | **Zwol** | 0 | 2.41 | 0.00 | 1.00 | 0.0005 | 0.0013 | -67431 | -61553 | -5878 | 5467 | 99.6% | 0.0% |
| **Ofla_Agal_Zcal** | **Ofla** | 0 | 0.92 | 0.00 | 1.00 | 0.0002 | 0.0002 | -73 | -72 | -1 | 5 | 0.0% | 0.0% |
| **Ofla_Agal_Zcal** | **Agal** | 0 | 0.47 | 0.00 | 1.00 | 0.0005 | 0.0006 | -192 | -191 | 0 | 15 | 0.0% | 0.0% |

| | | | | | | | | | | | | | |
|---|---|---|---|---|---|---|---|---|---|---|---|---|---|
| **Ofla_Agal_Zcal** | **Zcal** | 0 | 2.41 | 0.00 | 1.00 | 0.0005 | 0.0013 | -67431 | -61553 | -5878 | 5467 | 99.6% | 0.0% |
| **Ofla_Agal_Ejub** | **Ofla** | 0 | 0.92 | 0.00 | 1.00 | 0.0002 | 0.0002 | -73 | -72 | -1 | 5 | 0.0% | 0.0% |
| **Ofla_Agal_Ejub** | **Agal** | 0 | 0.47 | 0.00 | 1.00 | 0.0005 | 0.0006 | -192 | -191 | 0 | 15 | 0.0% | 0.0% |
| **Ofla_Agal_Ejub** | **Ejub** | 0 | 2.41 | 0.00 | 1.00 | 0.0005 | 0.0013 | -67431 | -61553 | -5878 | 5467 | 99.6% | 0.0% |
| **Ofla_Agal_Curs** | **Ofla** | 0 | 0 | 0 | 0 | 0 | 0 | 0 | 0 | 0 | 1 | 0.0% | 0.0% |
| **Ofla_Agal_Curs** | **Agal** | 0 | 0 | 0 | 0 | 0 | 0 | 0 | 0 | 0 | 1 | 0.0% | 0.0% |
| **Ofla_Agal_Curs** | **Curs** | 0 | 3.80 | 0.00 | 1.00 | 0.0008 | 0.0033 | -60423 | -51611 | -8812 | 5485 | 100.0% | 0.0% |
| **Ofla_Agal_Oros** | **Ofla** | 0 | 0 | 0 | 0 | 0 | 0 | 0 | 0 | 0 | 0 | 0.0% | 0.0% |
| **Ofla_Agal_Oros** | **Agal** | 0 | 0 | 0 | 0 | 0 | 0 | 0 | 0 | 0 | 0 | 0.0% | 0.0% |
| **Ofla_Agal_Oros** | **Oros** | 0 | 5.75 | 0.00 | 1.00 | 0.0020 | 0.0124 | -49578 | -37190 | -12388 | 5487 | 100.0% | 0.0% |
| **Ofla_Aaus_Phoo** | **Ofla** | 0 | 5.54 | 1.00 | 0.00 | 0.0002 | 0.0002 | -43495 | -43510 | 16 | 2964 | 0.0% | 54.0% |
| **Ofla_Aaus_Phoo** | **Aaus** | 0 | 5.01 | 0.99 | 0.01 | 0.0002 | 0.0002 | -19268 | -19271 | 3 | 1249 | 0.0% | 0.0% |
| **Ofla_Aaus_Phoo** | **Phoo** | 0 | 4.29 | 0.99 | 0.01 | 0.0002 | 0.0002 | -19748 | -19761 | 13 | 1274 | 0.0% | 23.0% |
| **Ofla_Aaus_Zwol** | **Ofla** | 0 | 0.92 | 0.00 | 1.00 | 0.0002 | 0.0002 | -73 | -72 | -1 | 5 | 0.0% | 0.0% |
| **Ofla_Aaus_Zwol** | **Aaus** | 0 | 0.47 | 0.00 | 1.00 | 0.0005 | 0.0006 | -192 | -191 | 0 | 15 | 0.0% | 0.0% |
| **Ofla_Aaus_Zwol** | **Zwol** | 0 | 2.41 | 0.00 | 1.00 | 0.0005 | 0.0013 | -67427 | -61549 | -5878 | 5467 | 99.6% | 0.0% |
| **Ofla_Aaus_Zcal** | **Ofla** | 0 | 0.92 | 0.00 | 1.00 | 0.0002 | 0.0002 | -73 | -72 | -1 | 5 | 0.0% | 0.0% |
| **Ofla_Aaus_Zcal** | **Aaus** | 0 | 0.47 | 0.00 | 1.00 | 0.0005 | 0.0006 | -192 | -191 | 0 | 15 | 0.0% | 0.0% |
| **Ofla_Aaus_Zcal** | **Zcal** | 0 | 2.41 | 0.00 | 1.00 | 0.0005 | 0.0013 | -67427 | -61549 | -5878 | 5467 | 99.6% | 0.0% |
| **Ofla_Aaus_Ejub** | **Ofla** | 0 | 0.92 | 0.00 | 1.00 | 0.0002 | 0.0002 | -73 | -72 | -1 | 5 | 0.0% | 0.0% |
| **Ofla_Aaus_Ejub** | **Aaus** | 0 | 0.47 | 0.00 | 1.00 | 0.0005 | 0.0006 | -192 | -191 | 0 | 15 | 0.0% | 0.0% |
| **Ofla_Aaus_Ejub** | **Ejub** | 0 | 2.41 | 0.00 | 1.00 | 0.0005 | 0.0013 | -67427 | -61549 | -5878 | 5467 | 99.6% | 0.0% |
| **Ofla_Aaus_Curs** | **Ofla** | 0 | 0 | 0 | 0 | 0 | 0 | 0 | 0 | 0 | 1 | 0.0% | 0.0% |
| **Ofla_Aaus_Curs** | **Aaus** | 0 | 0 | 0 | 0 | 0 | 0 | 0 | 0 | 0 | 1 | 0.0% | 0.0% |
| **Ofla_Aaus_Curs** | **Curs** | 0 | 3.80 | 0.00 | 1.00 | 0.0008 | 0.0033 | -60420 | -51610 | -8810 | 5485 | 100.0% | 0.0% |
| **Ofla_Aaus_Oros** | **Ofla** | 0 | 0 | 0 | 0 | 0 | 0 | 0 | 0 | 0 | 0 | 0.0% | 0.0% |
| **Ofla_Aaus_Oros** | **Aaus** | 0 | 0 | 0 | 0 | 0 | 0 | 0 | 0 | 0 | 0 | 0.0% | 0.0% |
| **Ofla_Aaus_Oros** | **Oros** | 0 | 5.75 | 0.00 | 1.00 | 0.0020 | 0.0124 | -49576 | -37190 | -12386 | 5487 | 100.0% | 0.0% |
| **Ofla_Phoo_Zwol** | **Ofla** | 0 | 0.30 | 0.00 | 1.00 | 0.0005 | 0.0005 | -77 | -79 | 2 | 6 | 0.0% | 0.0% |
| **Ofla_Phoo_Zwol** | **Phoo** | 0 | 1.19 | 0.80 | 0.20 | 0.0005 | 0.0006 | -184 | -190 | 6 | 15 | 0.0% | 0.0% |
| **Ofla_Phoo_Zwol** | **Zwol** | 0 | 2.41 | 0.00 | 1.00 | 0.0005 | 0.0013 | -67404 | -61529 | -5875 | 5466 | 99.6% | 0.0% |
| **Ofla_Phoo_Zcal** | **Ofla** | 0 | 0.30 | 0.00 | 1.00 | 0.0005 | 0.0005 | -77 | -79 | 2 | 6 | 0.0% | 0.0% |

| | | | | | | | | | | | | | |
|---|---|---|---|---|---|---|---|---|---|---|---|---|---|
| **Ofla_Phoo_Zcal** | **Phoo** | 0 | 1.19 | 0.80 | 0.20 | 0.0005 | 0.0006 | -184 | -190 | 6 | 15 | 0.0% | 0.0% |
| **Ofla_Phoo_Zcal** | **Zcal** | 0 | 2.41 | 0.00 | 1.00 | 0.0005 | 0.0013 | -67404 | -61529 | -5875 | 5466 | 99.6% | 0.0% |
| **Ofla_Phoo_Ejub** | **Ofla** | 0 | 0.30 | 0.00 | 1.00 | 0.0005 | 0.0005 | -77 | -79 | 2 | 6 | 0.0% | 0.0% |
| **Ofla_Phoo_Ejub** | **Phoo** | 0 | 1.19 | 0.80 | 0.20 | 0.0005 | 0.0006 | -184 | -190 | 6 | 15 | 0.0% | 0.0% |
| **Ofla_Phoo_Ejub** | **Ejub** | 0 | 2.41 | 0.00 | 1.00 | 0.0005 | 0.0013 | -67404 | -61529 | -5875 | 5466 | 99.6% | 0.0% |
| **Ofla_Phoo_Curs** | **Ofla** | 0 | 0 | 0 | 0 | 0 | 0 | 0 | 0 | 0 | 1 | 0.0% | 0.0% |
| **Ofla_Phoo_Curs** | **Phoo** | 0 | 0 | 0 | 0 | 0 | 0 | 0 | 0 | 0 | 1 | 0.0% | 0.0% |
| **Ofla_Phoo_Curs** | **Curs** | 0 | 3.80 | 0.00 | 1.00 | 0.0008 | 0.0033 | -60433 | -51607 | -8826 | 5485 | 100.0% | 0.0% |
| **Ofla_Phoo_Oros** | **Ofla** | 0 | 0 | 0 | 0 | 0 | 0 | 0 | 0 | 0 | 0 | 0.0% | 0.0% |
| **Ofla_Phoo_Oros** | **Phoo** | 0 | 0 | 0 | 0 | 0 | 0 | 0 | 0 | 0 | 0 | 0.0% | 0.0% |
| **Ofla_Phoo_Oros** | **Oros** | 0 | 5.75 | 0.00 | 1.00 | 0.0020 | 0.0124 | -49577 | -37189 | -12388 | 5487 | 100.0% | 0.0% |
| **Ofla_Zwol_Zcal** | **Ofla** | 0 | 3.98 | 0.00 | 1.00 | 0.0006 | 0.0026 | -63363 | -54302 | -9061 | 5487 | 100.0% | 0.0% |
| **Ofla_Zwol_Zcal** | **Zwol** | 0 | 0 | 0 | 0 | 0 | 0 | 0 | 0 | 0 | 0 | 0.0% | 0.0% |
| **Ofla_Zwol_Zcal** | **Zcal** | 0 | 0 | 0 | 0 | 0 | 0 | 0 | 0 | 0 | 0 | 0.0% | 0.0% |
| **Ofla_Zwol_Ejub** | **Ofla** | 0 | 3.99 | 0.00 | 1.00 | 0.0006 | 0.0026 | -63364 | -54314 | -9050 | 5487 | 100.0% | 0.0% |
| **Ofla_Zwol_Ejub** | **Zwol** | 0 | 0 | 0 | 0 | 0 | 0 | 0 | 0 | 0 | 0 | 0.0% | 0.0% |
| **Ofla_Zwol_Ejub** | **Ejub** | 0 | 0 | 0 | 0 | 0 | 0 | 0 | 0 | 0 | 0 | 0.0% | 0.0% |
| **Ofla_Zwol_Curs** | **Ofla** | 0 | 0 | 0 | 0 | 0 | 0 | 0 | 0 | 0 | 1 | 0.0% | 0.0% |
| **Ofla_Zwol_Curs** | **Zwol** | 0 | 3.92 | 0.65 | 0.35 | 0.0005 | 0.0010 | -33 | -34 | 1 | 3 | 0.0% | 0.0% |
| **Ofla_Zwol_Curs** | **Curs** | 0 | 3.13 | 0.00 | 1.00 | 0.0006 | 0.0020 | -64457 | -57096 | -7362 | 5483 | 99.9% | 0.0% |
| **Ofla_Zwol_Oros** | **Ofla** | 0 | 0 | 0 | 0 | 0 | 0 | 0 | 0 | 0 | 0 | 0.0% | 0.0% |
| **Ofla_Zwol_Oros** | **Zwol** | 0 | 0 | 0 | 0 | 0 | 0 | 0 | 0 | 0 | 0 | 0.0% | 0.0% |
| **Ofla_Zwol_Oros** | **Oros** | 0 | 5.89 | 0.00 | 1.00 | 0.0018 | 0.0111 | -51049 | -38418 | -12631 | 5487 | 100.0% | 0.0% |
| **Ofla_Zcal_Ejub** | **Ofla** | 0 | 3.98 | 0.00 | 1.00 | 0.0006 | 0.0027 | -63132 | -54053 | -9078 | 5487 | 100.0% | 0.0% |
| **Ofla_Zcal_Ejub** | **Zcal** | 0 | 0 | 0 | 0 | 0 | 0 | 0 | 0 | 0 | 0 | 0.0% | 0.0% |
| **Ofla_Zcal_Ejub** | **Ejub** | 0 | 0 | 0 | 0 | 0 | 0 | 0 | 0 | 0 | 0 | 0.0% | 0.0% |
| **Ofla_Zcal_Curs** | **Ofla** | 0 | 0 | 0 | 0 | 0 | 0 | 0 | 0 | 0 | 1 | 0.0% | 0.0% |
| **Ofla_Zcal_Curs** | **Zcal** | 0 | 3.92 | 0.65 | 0.35 | 0.0005 | 0.0010 | -33 | -34 | 1 | 3 | 0.0% | 0.0% |
| **Ofla_Zcal_Curs** | **Curs** | 0 | 3.13 | 0.00 | 1.00 | 0.0006 | 0.0020 | -64457 | -57096 | -7362 | 5483 | 99.9% | 0.0% |
| **Ofla_Zcal_Oros** | **Ofla** | 0 | 0 | 0 | 0 | 0 | 0 | 0 | 0 | 0 | 0 | 0.0% | 0.0% |
| **Ofla_Zcal_Oros** | **Zcal** | 0 | 0 | 0 | 0 | 0 | 0 | 0 | 0 | 0 | 0 | 0.0% | 0.0% |
| **Ofla_Zcal_Oros** | **Oros** | 0 | 5.89 | 0.00 | 1.00 | 0.0018 | 0.0111 | -51049 | -38418 | -12631 | 5487 | 100.0% | 0.0% |

| | | | | | | | | | | | | |
|---|---|---|---|---|---|---|---|---|---|---|---|---|
| **Ofla_Ejub_Curs** | **Ofla** | 0 | 0 | 0 | 0 | 0 | 0 | 0 | 0 | 0 | 1 | 0.0% | 0.0% |
| **Ofla_Ejub_Curs** | **Ejub** | 0 | 3.92 | 0.65 | 0.35 | 0.0005 | 0.0010 | -33 | -34 | 1 | 3 | 0.0% | 0.0% |
| **Ofla_Ejub_Curs** | **Curs** | 0 | 3.13 | 0.00 | 1.00 | 0.0006 | 0.0020 | -64457 | -57096 | -7362 | 5483 | 99.9% | 0.0% |
| **Ofla_Ejub_Oros** | **Ofla** | 0 | 0 | 0 | 0 | 0 | 0 | 0 | 0 | 0 | 0 | 0.0% | 0.0% |
| **Ofla_Ejub_Oros** | **Ejub** | 0 | 0 | 0 | 0 | 0 | 0 | 0 | 0 | 0 | 0 | 0.0% | 0.0% |
| **Ofla_Ejub_Oros** | **Oros** | 0 | 5.89 | 0.00 | 1.00 | 0.0018 | 0.0111 | -51049 | -38418 | -12631 | 5487 | 100.0% | 0.0% |
| **Ofla_Curs_Oros** | **Ofla** | 0 | 0 | 0 | 0 | 0 | 0 | 0 | 0 | 0 | 0 | 0.0% | 0.0% |
| **Ofla_Curs_Oros** | **Curs** | 0 | 0 | 0 | 0 | 0 | 0 | 0 | 0 | 0 | 0 | 0.0% | 0.0% |
| **Ofla_Curs_Oros** | **Oros** | 0 | 6.17 | 0.00 | 1.00 | 0.0014 | 0.0091 | -53691 | -40616 | -13076 | 5487 | 100.0% | 0.0% |
| **Atow_Aphi_Agaz** | **Atow** | 0 | 1.04 | 0.00 | 1.00 | 0.0003 | 0.0003 | -70 | -70 | 0 | 5 | 0.0% | 0.0% |
| **Atow_Aphi_Agaz** | **Aphi** | 0 | 3.51 | 0.84 | 0.16 | 0.0005 | 0.0006 | -60 | -63 | 2 | 5 | 0.0% | 0.0% |
| **Atow_Aphi_Agaz** | **Agaz** | 0 | 1.82 | 0.00 | 1.00 | 0.0005 | 0.0010 | -70181 | -64322 | -5859 | 5477 | 99.8% | 0.0% |
| **Atow_Aphi_Afor** | **Atow** | 0 | 1.95 | 0.36 | 0.64 | 0.0005 | 0.0009 | -46 | -47 | 2 | 4 | 0.0% | 0.0% |
| **Atow_Aphi_Afor** | **Aphi** | 0 | 6.07 | 0.75 | 0.25 | 0.0005 | 0.0011 | -46 | -45 | 0 | 4 | 0.0% | 0.0% |
| **Atow_Aphi_Afor** | **Afor** | 0 | 1.80 | 0.00 | 1.00 | 0.0005 | 0.0010 | -70190 | -64395 | -5795 | 5479 | 99.9% | 0.0% |
| **Atow_Aphi_Agal** | **Atow** | 0 | 1.95 | 0.35 | 0.65 | 0.0005 | 0.0007 | -36 | -36 | 0 | 3 | 0.0% | 0.0% |
| **Atow_Aphi_Agal** | **Aphi** | 0 | 0.74 | 0.64 | 0.36 | 0.0006 | 0.0007 | -59 | -62 | 3 | 5 | 0.0% | 0.0% |
| **Atow_Aphi_Agal** | **Agal** | 0 | 1.79 | 0.00 | 1.00 | 0.0005 | 0.0010 | -70182 | -64420 | -5761 | 5479 | 99.9% | 0.0% |
| **Atow_Aphi_Aaus** | **Atow** | 0 | 2.09 | 0.24 | 0.76 | 0.0005 | 0.0009 | -47 | -46 | 0 | 4 | 0.0% | 0.0% |
| **Atow_Aphi_Aaus** | **Aphi** | 0 | 8.60 | 0.80 | 0.20 | 0.0005 | 0.0011 | -59 | -56 | -3 | 5 | 0.0% | 0.0% |
| **Atow_Aphi_Aaus** | **Aaus** | 0 | 1.79 | 0.00 | 1.00 | 0.0005 | 0.0010 | -70175 | -64404 | -5771 | 5478 | 99.8% | 0.0% |
| **Atow_Aphi_Phoo** | **Atow** | 0 | 2.02 | 0.81 | 0.19 | 0.0002 | 0.0002 | -43 | -45 | 2 | 3 | 0.0% | 0.0% |
| **Atow_Aphi_Phoo** | **Aphi** | 0 | 8.28 | 0.80 | 0.20 | 0.0005 | 0.0011 | -59 | -56 | -3 | 5 | 0.0% | 0.0% |
| **Atow_Aphi_Phoo** | **Phoo** | 0 | 1.98 | 0.00 | 1.00 | 0.0005 | 0.0011 | -69189 | -63210 | -5979 | 5479 | 99.9% | 0.0% |
| **Atow_Aphi_Zwol** | **Atow** | 0 | 0 | 0 | 0 | 0 | 0 | 0 | 0 | 0 | 0 | 0.0% | 0.0% |
| **Atow_Aphi_Zwol** | **Aphi** | 0 | 0 | 0 | 0 | 0 | 0 | 0 | 0 | 0 | 0 | 0.0% | 0.0% |
| **Atow_Aphi_Zwol** | **Zwol** | 0 | 3.45 | 0.00 | 1.00 | 0.0007 | 0.0025 | -62746 | -54600 | -8146 | 5487 | 100.0% | 0.0% |
| **Atow_Aphi_Zcal** | **Atow** | 0 | 0 | 0 | 0 | 0 | 0 | 0 | 0 | 0 | 0 | 0.0% | 0.0% |
| **Atow_Aphi_Zcal** | **Aphi** | 0 | 0 | 0 | 0 | 0 | 0 | 0 | 0 | 0 | 0 | 0.0% | 0.0% |
| **Atow_Aphi_Zcal** | **Zcal** | 0 | 3.45 | 0.00 | 1.00 | 0.0007 | 0.0025 | -62746 | -54600 | -8146 | 5487 | 100.0% | 0.0% |
| **Atow_Aphi_Ejub** | **Atow** | 0 | 0 | 0 | 0 | 0 | 0 | 0 | 0 | 0 | 0 | 0.0% | 0.0% |
| **Atow_Aphi_Ejub** | **Aphi** | 0 | 0 | 0 | 0 | 0 | 0 | 0 | 0 | 0 | 0 | 0.0% | 0.0% |

| | | | | | | | | | | | | | |
|---|---|---|---|---|---|---|---|---|---|---|---|---|---|
| **Atow_Aphi_Ejub** | **Ejub** | 0 | 3.45 | 0.00 | 1.00 | 0.0007 | 0.0025 | -62746 | -54600 | -8146 | 5487 | 100.0% | 0.0% |
| **Atow_Aphi_Curs** | **Atow** | 0 | 0 | 0 | 0 | 0 | 0 | 0 | 0 | 0 | 0 | 0.0% | 0.0% |
| **Atow_Aphi_Curs** | **Aphi** | 0 | 0 | 0 | 0 | 0 | 0 | 0 | 0 | 0 | 0 | 0.0% | 0.0% |
| **Atow_Aphi_Curs** | **Curs** | 0 | 4.05 | 0.00 | 1.00 | 0.0010 | 0.0046 | -57636 | -48196 | -9440 | 5487 | 100.0% | 0.0% |
| **Atow_Aphi_Oros** | **Atow** | 0 | 0 | 0 | 0 | 0 | 0 | 0 | 0 | 0 | 0 | 0.0% | 0.0% |
| **Atow_Aphi_Oros** | **Aphi** | 0 | 0 | 0 | 0 | 0 | 0 | 0 | 0 | 0 | 0 | 0.0% | 0.0% |
| **Atow_Aphi_Oros** | **Oros** | 0 | 5.64 | 0.00 | 1.00 | 0.0023 | 0.0136 | -48411 | -36158 | -12253 | 5487 | 100.0% | 0.0% |
| **Atow_Agaz_Afor** | **Atow** | 0 | 1.10 | 0.76 | 0.24 | 0.0002 | 0.0002 | -35349 | -35321 | -28 | 2413 | 10.6% | 0.0% |
| **Atow_Agaz_Afor** | **Agaz** | 0 | 1.22 | 0.84 | 0.16 | 0.0002 | 0.0002 | -24104 | -24119 | 15 | 1610 | 0.0% | 24.5% |
| **Atow_Agaz_Afor** | **Afor** | 0 | 0.86 | 0.70 | 0.30 | 0.0002 | 0.0002 | -21915 | -21881 | -34 | 1464 | 8.1% | 0.0% |
| **Atow_Agaz_Agal** | **Atow** | 0 | 1.17 | 0.78 | 0.22 | 0.0002 | 0.0002 | -35278 | -35266 | -12 | 2411 | 9.5% | 0.0% |
| **Atow_Agaz_Agal** | **Agaz** | 0 | 2.42 | 0.96 | 0.04 | 0.0002 | 0.0002 | -24293 | -24332 | 39 | 1625 | 0.0% | 28.4% |
| **Atow_Agaz_Agal** | **Agal** | 0 | 0.95 | 0.70 | 0.30 | 0.0002 | 0.0002 | -21757 | -21734 | -23 | 1451 | 7.9% | 0.0% |
| **Atow_Agaz_Aaus** | **Atow** | 0 | 1.11 | 0.77 | 0.23 | 0.0002 | 0.0002 | -35380 | -35352 | -28 | 2417 | 10.3% | 0.0% |
| **Atow_Agaz_Aaus** | **Agaz** | 0 | 1.70 | 0.92 | 0.08 | 0.0002 | 0.0002 | -24165 | -24204 | 39 | 1620 | 0.0% | 27.2% |
| **Atow_Agaz_Aaus** | **Aaus** | 0 | 0.87 | 0.70 | 0.30 | 0.0002 | 0.0002 | -21721 | -21687 | -35 | 1450 | 8.0% | 0.0% |
| **Atow_Agaz_Phoo** | **Atow** | 0 | 6.87 | 1.00 | 0.00 | 0.0002 | 0.0002 | -21083 | -21093 | 10 | 1389 | 0.0% | 0.0% |
| **Atow_Agaz_Phoo** | **Agaz** | 0 | 30.04 | 1.00 | 0.00 | 0.0002 | 0.0002 | -17140 | -17111 | -29 | 1114 | 0.0% | 0.0% |
| **Atow_Agaz_Phoo** | **Phoo** | 0 | 6.67 | 1.00 | 0.00 | 0.0003 | 0.0003 | -43031 | -43047 | 16 | 2984 | 0.0% | 54.4% |
| **Atow_Agaz_Zwol** | **Atow** | 0 | 0 | 0 | 0 | 0 | 0 | 0 | 0 | 0 | 1 | 0.0% | 0.0% |
| **Atow_Agaz_Zwol** | **Agaz** | 0 | 2.01 | 0.82 | 0.18 | 0.0002 | 0.0002 | -42 | -44 | 2 | 3 | 0.0% | 0.0% |
| **Atow_Agaz_Zwol** | **Zwol** | 0 | 2.40 | 0.00 | 1.00 | 0.0006 | 0.0015 | -66028 | -60281 | -5747 | 5483 | 99.9% | 0.0% |
| **Atow_Agaz_Zcal** | **Atow** | 0 | 0 | 0 | 0 | 0 | 0 | 0 | 0 | 0 | 1 | 0.0% | 0.0% |
| **Atow_Agaz_Zcal** | **Agaz** | 0 | 2.01 | 0.82 | 0.18 | 0.0002 | 0.0002 | -42 | -44 | 2 | 3 | 0.0% | 0.0% |
| **Atow_Agaz_Zcal** | **Zcal** | 0 | 2.40 | 0.00 | 1.00 | 0.0006 | 0.0015 | -66028 | -60281 | -5747 | 5483 | 99.9% | 0.0% |
| **Atow_Agaz_Ejub** | **Atow** | 0 | 0 | 0 | 0 | 0 | 0 | 0 | 0 | 0 | 1 | 0.0% | 0.0% |
| **Atow_Agaz_Ejub** | **Agaz** | 0 | 2.01 | 0.82 | 0.18 | 0.0002 | 0.0002 | -42 | -44 | 2 | 3 | 0.0% | 0.0% |
| **Atow_Agaz_Ejub** | **Ejub** | 0 | 2.40 | 0.00 | 1.00 | 0.0006 | 0.0015 | -66028 | -60281 | -5747 | 5483 | 99.9% | 0.0% |
| **Atow_Agaz_Curs** | **Atow** | 0 | 0 | 0 | 0 | 0 | 0 | 0 | 0 | 0 | 1 | 0.0% | 0.0% |
| **Atow_Agaz_Curs** | **Agaz** | 0 | 0 | 0 | 0 | 0 | 0 | 0 | 0 | 0 | 1 | 0.0% | 0.0% |
| **Atow_Agaz_Curs** | **Curs** | 0 | 3.61 | 0.00 | 1.00 | 0.0009 | 0.0035 | -59585 | -50999 | -8586 | 5485 | 100.0% | 0.0% |
| **Atow_Agaz_Oros** | **Atow** | 0 | 0 | 0 | 0 | 0 | 0 | 0 | 0 | 0 | 0 | 0.0% | 0.0% |

| | | | | | | | | | | | | |
|---|---|---|---|---|---|---|---|---|---|---|---|---|
| **Atow_Agaz_Oros** | **Agaz** | 0 | 0 | 0 | 0 | 0 | 0 | 0 | 0 | 0 | 0 | 0.0% | 0.0% |
| **Atow_Agaz_Oros** | **Oros** | 0 | 5.62 | 0.00 | 1.00 | 0.0021 | 0.0126 | -49224 | -37023 | - 12201 | 5487 | 100.0% | 0.0% |
| **Atow_Afor_Agal** | **Atow** | 0 | 1.01 | 0.02 | 0.98 | 0.0005 | 0.0007 | -69411 | -66633 | -2778 | 5332 | 95.6% | 0.0% |
| **Atow_Afor_Agal** | **Afor** | 0 | 4.64 | 0.99 | 0.01 | 0.0003 | 0.0003 | -1246 | -1255 | 8 | 90 | 0.0% | 0.0% |
| **Atow_Afor_Agal** | **Agal** | 0 | 0.57 | 0.81 | 0.19 | 0.0003 | 0.0003 | -920 | -928 | 8 | 65 | 0.0% | 0.0% |
| **Atow_Afor_Aaus** | **Atow** | 0 | 1.02 | 0.01 | 0.99 | 0.0005 | 0.0007 | -69453 | -66616 | -2837 | 5343 | 96.2% | 0.0% |
| **Atow_Afor_Aaus** | **Afor** | 0 | 1.17 | 0.88 | 0.12 | 0.0003 | 0.0003 | -1205 | -1214 | 8 | 86 | 0.0% | 0.0% |
| **Atow_Afor_Aaus** | **Aaus** | 0 | 0.62 | 0.68 | 0.32 | 0.0002 | 0.0002 | -845 | -852 | 7 | 58 | 0.0% | 0.0% |
| **Atow_Afor_Phoo** | **Atow** | 0 | 6.53 | 1.00 | 0.00 | 0.0002 | 0.0002 | -20719 | -20728 | 9 | 1363 | 0.0% | 0.0% |
| **Atow_Afor_Phoo** | **Afor** | 0 | 6.58 | 1.00 | 0.00 | 0.0002 | 0.0002 | -16334 | -16347 | 13 | 1054 | 0.0% | 19.2% |
| **Atow_Afor_Phoo** | **Phoo** | 0 | 5.31 | 1.00 | 0.00 | 0.0003 | 0.0003 | -44433 | -44450 | 16 | 3070 | 0.0% | 55.9% |
| **Atow_Afor_Zwol** | **Atow** | 0 | 0 | 0 | 0 | 0 | 0 | 0 | 0 | 0 | 1 | 0.0% | 0.0% |
| **Atow_Afor_Zwol** | **Afor** | 0 | 0.46 | 0.00 | 1.00 | 0.0003 | 0.0003 | -42 | -42 | 1 | 3 | 0.0% | 0.0% |
| **Atow_Afor_Zwol** | **Zwol** | 0 | 2.43 | 0.00 | 1.00 | 0.0006 | 0.0015 | -66002 | -60248 | -5753 | 5483 | 99.9% | 0.0% |
| **Atow_Afor_Zcal** | **Atow** | 0 | 0 | 0 | 0 | 0 | 0 | 0 | 0 | 0 | 1 | 0.0% | 0.0% |
| **Atow_Afor_Zcal** | **Afor** | 0 | 0.46 | 0.00 | 1.00 | 0.0003 | 0.0003 | -42 | -42 | 1 | 3 | 0.0% | 0.0% |
| **Atow_Afor_Zcal** | **Zcal** | 0 | 2.43 | 0.00 | 1.00 | 0.0006 | 0.0015 | -66002 | -60248 | -5753 | 5483 | 99.9% | 0.0% |
| **Atow_Afor_Ejub** | **Atow** | 0 | 0 | 0 | 0 | 0 | 0 | 0 | 0 | 0 | 1 | 0.0% | 0.0% |
| **Atow_Afor_Ejub** | **Afor** | 0 | 0.46 | 0.00 | 1.00 | 0.0003 | 0.0003 | -42 | -42 | 1 | 3 | 0.0% | 0.0% |
| **Atow_Afor_Ejub** | **Ejub** | 0 | 2.43 | 0.00 | 1.00 | 0.0006 | 0.0015 | -66002 | -60248 | -5753 | 5483 | 99.9% | 0.0% |
| **Atow_Afor_Curs** | **Atow** | 0 | 12.23 | 0.50 | 0.50 | 0.0005 | 0.0033 | -22 | -18 | -4 | 2 | 0.0% | 0.0% |
| **Atow_Afor_Curs** | **Afor** | 0 | 0 | 0 | 0 | 0 | 0 | 0 | 0 | 0 | 0 | 0.0% | 0.0% |
| **Atow_Afor_Curs** | **Curs** | 0 | 3.62 | 0.00 | 1.00 | 0.0009 | 0.0035 | -59567 | -50985 | -8582 | 5485 | 100.0% | 0.0% |
| **Atow_Afor_Oros** | **Atow** | 0 | 0 | 0 | 0 | 0 | 0 | 0 | 0 | 0 | 0 | 0.0% | 0.0% |
| **Atow_Afor_Oros** | **Afor** | 0 | 0 | 0 | 0 | 0 | 0 | 0 | 0 | 0 | 0 | 0.0% | 0.0% |
| **Atow_Afor_Oros** | **Oros** | 0 | 5.62 | 0.00 | 1.00 | 0.0021 | 0.0126 | -49214 | -37019 | - 12195 | 5487 | 100.0% | 0.0% |
| **Atow_Agal_Aaus** | **Atow** | 0 | 1.73 | 0.01 | 0.99 | 0.0005 | 0.0010 | -67863 | -63744 | -4120 | 5408 | 97.9% | 0.0% |
| **Atow_Agal_Aaus** | **Agal** | 0 | 0.60 | 0.62 | 0.38 | 0.0005 | 0.0005 | -539 | -545 | 6 | 41 | 0.0% | 0.0% |
| **Atow_Agal_Aaus** | **Aaus** | 0 | 4.10 | 0.99 | 0.01 | 0.0005 | 0.0005 | -498 | -505 | 7 | 38 | 0.0% | 0.0% |
| **Atow_Agal_Phoo** | **Atow** | 0 | 5.30 | 1.00 | 0.00 | 0.0002 | 0.0002 | -20748 | -20761 | 14 | 1362 | 0.0% | 24.7% |
| **Atow_Agal_Phoo** | **Agal** | 0 | 28.28 | 1.00 | 0.00 | 0.0002 | 0.0002 | -16274 | -16248 | -25 | 1049 | 0.0% | 0.0% |
| **Atow_Agal_Phoo** | **Phoo** | 0 | 5.32 | 1.00 | 0.00 | 0.0003 | 0.0003 | -44436 | -44452 | 16 | 3076 | 0.0% | 56.0% |

| | | | | | | | | | | | | | |
|---|---|---|---|---|---|---|---|---|---|---|---|---|---|
| **Atow_Agal_Zwol** | **Atow** | 0 | 0 | 0 | 0 | 0 | 0 | 0 | 0 | 0 | 1 | 0.0% | 0.0% |
| **Atow_Agal_Zwol** | **Agal** | 0 | 0.46 | 0.00 | 1.00 | 0.0003 | 0.0003 | -42 | -42 | 1 | 3 | 0.0% | 0.0% |
| **Atow_Agal_Zwol** | **Zwol** | 0 | 2.42 | 0.00 | 1.00 | 0.0006 | 0.0015 | -65974 | -60233 | -5742 | 5483 | 99.9% | 0.0% |
| **Atow_Agal_Zcal** | **Atow** | 0 | 0 | 0 | 0 | 0 | 0 | 0 | 0 | 0 | 1 | 0.0% | 0.0% |
| **Atow_Agal_Zcal** | **Agal** | 0 | 0.46 | 0.00 | 1.00 | 0.0003 | 0.0003 | -42 | -42 | 1 | 3 | 0.0% | 0.0% |
| **Atow_Agal_Zcal** | **Zcal** | 0 | 2.42 | 0.00 | 1.00 | 0.0006 | 0.0015 | -65974 | -60233 | -5742 | 5483 | 99.9% | 0.0% |
| **Atow_Agal_Ejub** | **Atow** | 0 | 0 | 0 | 0 | 0 | 0 | 0 | 0 | 0 | 1 | 0.0% | 0.0% |
| **Atow_Agal_Ejub** | **Agal** | 0 | 0.46 | 0.00 | 1.00 | 0.0003 | 0.0003 | -42 | -42 | 1 | 3 | 0.0% | 0.0% |
| **Atow_Agal_Ejub** | **Ejub** | 0 | 2.42 | 0.00 | 1.00 | 0.0006 | 0.0015 | -65974 | -60233 | -5742 | 5483 | 99.9% | 0.0% |
| **Atow_Agal_Curs** | **Atow** | 0 | 9.83 | 0.50 | 0.50 | 0.0005 | 0.0027 | -22 | -19 | -3 | 2 | 0.0% | 0.0% |
| **Atow_Agal_Curs** | **Agal** | 0 | 0 | 0 | 0 | 0 | 0 | 0 | 0 | 0 | 0 | 0.0% | 0.0% |
| **Atow_Agal_Curs** | **Curs** | 0 | 3.62 | 0.00 | 1.00 | 0.0009 | 0.0035 | -59560 | -50978 | -8582 | 5485 | 100.0% | 0.0% |
| **Atow_Agal_Oros** | **Atow** | 0 | 0 | 0 | 0 | 0 | 0 | 0 | 0 | 0 | 0 | 0.0% | 0.0% |
| **Atow_Agal_Oros** | **Agal** | 0 | 0 | 0 | 0 | 0 | 0 | 0 | 0 | 0 | 0 | 0.0% | 0.0% |
| **Atow_Agal_Oros** | **Oros** | 0 | 5.62 | 0.00 | 1.00 | 0.0021 | 0.0126 | -49213 | -37017 | -<br>12196 | 5487 | 100.0% | 0.0% |
| **Atow_Aaus_Phoo** | **Atow** | 0 | 6.64 | 1.00 | 0.00 | 0.0002 | 0.0002 | -20841 | -20849 | 8 | 1368 | 0.0% | 0.0% |
| **Atow_Aaus_Phoo** | **Aaus** | 0 | 6.60 | 1.00 | 0.00 | 0.0002 | 0.0002 | -16215 | -16229 | 14 | 1046 | 0.0% | 19.0% |
| **Atow_Aaus_Phoo** | **Phoo** | 0 | 5.32 | 1.00 | 0.00 | 0.0003 | 0.0003 | -44396 | -44412 | 16 | 3073 | 0.0% | 56.0% |
| **Atow_Aaus_Zwol** | **Atow** | 0 | 0 | 0 | 0 | 0 | 0 | 0 | 0 | 0 | 1 | 0.0% | 0.0% |
| **Atow_Aaus_Zwol** | **Aaus** | 0 | 0.46 | 0.00 | 1.00 | 0.0003 | 0.0003 | -42 | -42 | 1 | 3 | 0.0% | 0.0% |
| **Atow_Aaus_Zwol** | **Zwol** | 0 | 2.42 | 0.00 | 1.00 | 0.0006 | 0.0015 | -65969 | -60231 | -5738 | 5483 | 99.9% | 0.0% |
| **Atow_Aaus_Zcal** | **Atow** | 0 | 0 | 0 | 0 | 0 | 0 | 0 | 0 | 0 | 1 | 0.0% | 0.0% |
| **Atow_Aaus_Zcal** | **Aaus** | 0 | 0.46 | 0.00 | 1.00 | 0.0003 | 0.0003 | -42 | -42 | 1 | 3 | 0.0% | 0.0% |
| **Atow_Aaus_Zcal** | **Zcal** | 0 | 2.42 | 0.00 | 1.00 | 0.0006 | 0.0015 | -65969 | -60231 | -5738 | 5483 | 99.9% | 0.0% |
| **Atow_Aaus_Ejub** | **Atow** | 0 | 0 | 0 | 0 | 0 | 0 | 0 | 0 | 0 | 1 | 0.0% | 0.0% |
| **Atow_Aaus_Ejub** | **Aaus** | 0 | 0.46 | 0.00 | 1.00 | 0.0003 | 0.0003 | -42 | -42 | 1 | 3 | 0.0% | 0.0% |
| **Atow_Aaus_Ejub** | **Ejub** | 0 | 2.42 | 0.00 | 1.00 | 0.0006 | 0.0015 | -65969 | -60231 | -5738 | 5483 | 99.9% | 0.0% |
| **Atow_Aaus_Curs** | **Atow** | 0 | 9.83 | 0.50 | 0.50 | 0.0005 | 0.0027 | -22 | -19 | -3 | 2 | 0.0% | 0.0% |
| **Atow_Aaus_Curs** | **Aaus** | 0 | 0 | 0 | 0 | 0 | 0 | 0 | 0 | 0 | 0 | 0.0% | 0.0% |
| **Atow_Aaus_Curs** | **Curs** | 0 | 3.62 | 0.00 | 1.00 | 0.0009 | 0.0035 | -59555 | -50978 | -8577 | 5485 | 100.0% | 0.0% |
| **Atow_Aaus_Oros** | **Atow** | 0 | 0 | 0 | 0 | 0 | 0 | 0 | 0 | 0 | 0 | 0.0% | 0.0% |
| **Atow_Aaus_Oros** | **Aaus** | 0 | 0 | 0 | 0 | 0 | 0 | 0 | 0 | 0 | 0 | 0.0% | 0.0% |

| | | | | | | | | | | | | | |
|---|---|---|---|---|---|---|---|---|---|---|---|---|---|
| **Atow_Aaus_Oros** | **Oros** | 0 | 5.62 | 0.00 | 1.00 | 0.0021 | 0.0126 | -49213 | -37017 | -12197 | 5487 | 100.0% | 0.0% |
| **Atow_Phoo_Zwol** | **Atow** | 0 | 1.33 | 0.23 | 0.77 | 0.0005 | 0.0006 | -61 | -62 | 1 | 5 | 0.0% | 0.0% |
| **Atow_Phoo_Zwol** | **Phoo** | 0 | 0.85 | 0.69 | 0.31 | 0.0003 | 0.0003 | -109 | -113 | 4 | 8 | 0.0% | 0.0% |
| **Atow_Phoo_Zwol** | **Zwol** | 0 | 2.54 | 0.00 | 1.00 | 0.0005 | 0.0014 | -66897 | -61015 | -5882 | 5474 | 99.8% | 0.0% |
| **Atow_Phoo_Zcal** | **Atow** | 0 | 1.33 | 0.23 | 0.77 | 0.0005 | 0.0006 | -61 | -62 | 1 | 5 | 0.0% | 0.0% |
| **Atow_Phoo_Zcal** | **Phoo** | 0 | 0.85 | 0.69 | 0.31 | 0.0003 | 0.0003 | -109 | -113 | 4 | 8 | 0.0% | 0.0% |
| **Atow_Phoo_Zcal** | **Zcal** | 0 | 2.54 | 0.00 | 1.00 | 0.0005 | 0.0014 | -66897 | -61015 | -5882 | 5474 | 99.8% | 0.0% |
| **Atow_Phoo_Ejub** | **Atow** | 0 | 1.33 | 0.23 | 0.77 | 0.0005 | 0.0006 | -61 | -62 | 1 | 5 | 0.0% | 0.0% |
| **Atow_Phoo_Ejub** | **Phoo** | 0 | 0.85 | 0.69 | 0.31 | 0.0003 | 0.0003 | -109 | -113 | 4 | 8 | 0.0% | 0.0% |
| **Atow_Phoo_Ejub** | **Ejub** | 0 | 2.54 | 0.00 | 1.00 | 0.0005 | 0.0014 | -66897 | -61015 | -5882 | 5474 | 99.8% | 0.0% |
| **Atow_Phoo_Curs** | **Atow** | 0 | 0 | 0 | 0 | 0 | 0 | 0 | 0 | 0 | 1 | 0.0% | 0.0% |
| **Atow_Phoo_Curs** | **Phoo** | 0 | 0 | 0 | 0 | 0 | 0 | 0 | 0 | 0 | 1 | 0.0% | 0.0% |
| **Atow_Phoo_Curs** | **Curs** | 0 | 3.74 | 0.00 | 1.00 | 0.0008 | 0.0034 | -60085 | -51357 | -8728 | 5485 | 100.0% | 0.0% |
| **Atow_Phoo_Oros** | **Atow** | 0 | 0 | 0 | 0 | 0 | 0 | 0 | 0 | 0 | 0 | 0.0% | 0.0% |
| **Atow_Phoo_Oros** | **Phoo** | 0 | 0 | 0 | 0 | 0 | 0 | 0 | 0 | 0 | 0 | 0.0% | 0.0% |
| **Atow_Phoo_Oros** | **Oros** | 0 | 5.69 | 0.00 | 1.00 | 0.0021 | 0.0125 | -49430 | -37122 | -12309 | 5487 | 100.0% | 0.0% |
| **Atow_Zwol_Zcal** | **Atow** | 0 | 3.97 | 0.00 | 1.00 | 0.0006 | 0.0026 | -63346 | -54297 | -9049 | 5487 | 100.0% | 0.0% |
| **Atow_Zwol_Zcal** | **Zwol** | 0 | 0 | 0 | 0 | 0 | 0 | 0 | 0 | 0 | 0 | 0.0% | 0.0% |
| **Atow_Zwol_Zcal** | **Zcal** | 0 | 0 | 0 | 0 | 0 | 0 | 0 | 0 | 0 | 0 | 0.0% | 0.0% |
| **Atow_Zwol_Ejub** | **Atow** | 0 | 3.98 | 0.00 | 1.00 | 0.0006 | 0.0026 | -63347 | -54308 | -9038 | 5487 | 100.0% | 0.0% |
| **Atow_Zwol_Ejub** | **Zwol** | 0 | 0 | 0 | 0 | 0 | 0 | 0 | 0 | 0 | 0 | 0.0% | 0.0% |
| **Atow_Zwol_Ejub** | **Ejub** | 0 | 0 | 0 | 0 | 0 | 0 | 0 | 0 | 0 | 0 | 0.0% | 0.0% |
| **Atow_Zwol_Curs** | **Atow** | 0 | 0 | 0 | 0 | 0 | 0 | 0 | 0 | 0 | 1 | 0.0% | 0.0% |
| **Atow_Zwol_Curs** | **Zwol** | 0 | 1.56 | 0.04 | 0.96 | 0.0005 | 0.0008 | -35 | -35 | 1 | 3 | 0.0% | 0.0% |
| **Atow_Zwol_Curs** | **Curs** | 0 | 3.13 | 0.00 | 1.00 | 0.0006 | 0.0020 | -64468 | -57100 | -7368 | 5483 | 99.9% | 0.0% |
| **Atow_Zwol_Oros** | **Atow** | 0 | 0 | 0 | 0 | 0 | 0 | 0 | 0 | 0 | 0 | 0.0% | 0.0% |
| **Atow_Zwol_Oros** | **Zwol** | 0 | 0 | 0 | 0 | 0 | 0 | 0 | 0 | 0 | 0 | 0.0% | 0.0% |
| **Atow_Zwol_Oros** | **Oros** | 0 | 5.89 | 0.00 | 1.00 | 0.0018 | 0.0111 | -51053 | -38420 | -12634 | 5487 | 100.0% | 0.0% |
| **Atow_Zcal_Ejub** | **Atow** | 0 | 3.97 | 0.00 | 1.00 | 0.0006 | 0.0027 | -63115 | -54048 | -9067 | 5487 | 100.0% | 0.0% |
| **Atow_Zcal_Ejub** | **Zcal** | 0 | 0 | 0 | 0 | 0 | 0 | 0 | 0 | 0 | 0 | 0.0% | 0.0% |
| **Atow_Zcal_Ejub** | **Ejub** | 0 | 0 | 0 | 0 | 0 | 0 | 0 | 0 | 0 | 0 | 0.0% | 0.0% |
| **Atow_Zcal_Curs** | **Atow** | 0 | 0 | 0 | 0 | 0 | 0 | 0 | 0 | 0 | 1 | 0.0% | 0.0% |

| | | | | | | | | | | | | | |
|---|---|---|---|---|---|---|---|---|---|---|---|---|---|
| **Atow_Zcal_Curs** | **Zcal** | 0 | 1.56 | 0.04 | 0.96 | 0.0005 | 0.0008 | -35 | -35 | 1 | 3 | 0.0% | 0.0% |
| **Atow_Zcal_Curs** | **Curs** | 0 | 3.13 | 0.00 | 1.00 | 0.0006 | 0.0020 | -64468 | -57100 | -7368 | 5483 | 99.9% | 0.0% |
| **Atow_Zcal_Oros** | **Atow** | 0 | 0 | 0 | 0 | 0 | 0 | 0 | 0 | 0 | 0 | 0.0% | 0.0% |
| **Atow_Zcal_Oros** | **Zcal** | 0 | 0 | 0 | 0 | 0 | 0 | 0 | 0 | 0 | 0 | 0.0% | 0.0% |
| **Atow_Zcal_Oros** | **Oros** | 0 | 5.89 | 0.00 | 1.00 | 0.0018 | 0.0111 | -51053 | -38420 | - 12634 | 5487 | 100.0% | 0.0% |
| **Atow_Ejub_Curs** | **Atow** | 0 | 0 | 0 | 0 | 0 | 0 | 0 | 0 | 0 | 1 | 0.0% | 0.0% |
| **Atow_Ejub_Curs** | **Ejub** | 0 | 1.56 | 0.04 | 0.96 | 0.0005 | 0.0008 | -35 | -35 | 1 | 3 | 0.0% | 0.0% |
| **Atow_Ejub_Curs** | **Curs** | 0 | 3.13 | 0.00 | 1.00 | 0.0006 | 0.0020 | -64468 | -57100 | -7368 | 5483 | 99.9% | 0.0% |
| **Atow_Ejub_Oros** | **Atow** | 0 | 0 | 0 | 0 | 0 | 0 | 0 | 0 | 0 | 0 | 0.0% | 0.0% |
| **Atow_Ejub_Oros** | **Ejub** | 0 | 0 | 0 | 0 | 0 | 0 | 0 | 0 | 0 | 0 | 0.0% | 0.0% |
| **Atow_Ejub_Oros** | **Oros** | 0 | 5.89 | 0.00 | 1.00 | 0.0018 | 0.0111 | -51053 | -38420 | - 12634 | 5487 | 100.0% | 0.0% |
| **Atow_Curs_Oros** | **Atow** | 0 | 0 | 0 | 0 | 0 | 0 | 0 | 0 | 0 | 0 | 0.0% | 0.0% |
| **Atow_Curs_Oros** | **Curs** | 0 | 0 | 0 | 0 | 0 | 0 | 0 | 0 | 0 | 0 | 0.0% | 0.0% |
| **Atow_Curs_Oros** | **Oros** | 0 | 6.17 | 0.00 | 1.00 | 0.0014 | 0.0091 | -53695 | -40616 | - 13079 | 5487 | 100.0% | 0.0% |
| **Aphi_Agaz_Afor** | **Aphi** | 0 | 1.12 | 0.77 | 0.23 | 0.0002 | 0.0002 | -35311 | -35288 | -23 | 2411 | 10.3% | 0.0% |
| **Aphi_Agaz_Afor** | **Agaz** | 0 | 1.25 | 0.85 | 0.15 | 0.0002 | 0.0002 | -24129 | -24144 | 15 | 1611 | 0.0% | 24.8% |
| **Aphi_Agaz_Afor** | **Afor** | 0 | 0.86 | 0.69 | 0.31 | 0.0002 | 0.0002 | -21950 | -21913 | -37 | 1465 | 8.4% | 0.0% |
| **Aphi_Agaz_Agal** | **Aphi** | 0 | 1.21 | 0.80 | 0.20 | 0.0002 | 0.0002 | -35238 | -35234 | -4 | 2410 | 0.0% | 0.0% |
| **Aphi_Agaz_Agal** | **Agaz** | 0 | 2.67 | 0.97 | 0.03 | 0.0002 | 0.0002 | -24273 | -24307 | 35 | 1624 | 0.0% | 28.7% |
| **Aphi_Agaz_Agal** | **Agal** | 0 | 0.88 | 0.69 | 0.31 | 0.0002 | 0.0002 | -21802 | -21766 | -36 | 1453 | 8.1% | 0.0% |
| **Aphi_Agaz_Aaus** | **Aphi** | 0 | 1.13 | 0.78 | 0.22 | 0.0002 | 0.0002 | -35341 | -35318 | -23 | 2415 | 9.9% | 0.0% |
| **Aphi_Agaz_Aaus** | **Agaz** | 0 | 2.15 | 0.94 | 0.06 | 0.0002 | 0.0002 | -24197 | -24239 | 42 | 1621 | 0.0% | 27.9% |
| **Aphi_Agaz_Aaus** | **Aaus** | 0 | 0.87 | 0.69 | 0.31 | 0.0002 | 0.0002 | -21755 | -21718 | -37 | 1451 | 8.2% | 0.0% |
| **Aphi_Agaz_Phoo** | **Aphi** | 0 | 5.04 | 1.00 | 0.00 | 0.0002 | 0.0002 | -21103 | -21117 | 14 | 1391 | 0.0% | 25.3% |
| **Aphi_Agaz_Phoo** | **Agaz** | 0 | 5.74 | 1.00 | 0.00 | 0.0002 | 0.0002 | -17147 | -17161 | 14 | 1114 | 0.0% | 20.3% |
| **Aphi_Agaz_Phoo** | **Phoo** | 0 | 6.67 | 1.00 | 0.00 | 0.0003 | 0.0003 | -43003 | -43019 | 16 | 2982 | 0.0% | 54.3% |
| **Aphi_Agaz_Zwol** | **Aphi** | 0 | 0 | 0 | 0 | 0 | 0 | 0 | 0 | 0 | 1 | 0.0% | 0.0% |
| **Aphi_Agaz_Zwol** | **Agaz** | 0 | 2.01 | 0.82 | 0.18 | 0.0002 | 0.0002 | -42 | -44 | 2 | 3 | 0.0% | 0.0% |
| **Aphi_Agaz_Zwol** | **Zwol** | 0 | 2.40 | 0.00 | 1.00 | 0.0006 | 0.0015 | -66028 | -60283 | -5745 | 5483 | 99.9% | 0.0% |
| **Aphi_Agaz_Zcal** | **Aphi** | 0 | 0 | 0 | 0 | 0 | 0 | 0 | 0 | 0 | 1 | 0.0% | 0.0% |
| **Aphi_Agaz_Zcal** | **Agaz** | 0 | 2.01 | 0.82 | 0.18 | 0.0002 | 0.0002 | -42 | -44 | 2 | 3 | 0.0% | 0.0% |
| **Aphi_Agaz_Zcal** | **Zcal** | 0 | 2.40 | 0.00 | 1.00 | 0.0006 | 0.0015 | -66028 | -60283 | -5745 | 5483 | 99.9% | 0.0% |

| | | | | | | | | | | | | |
|---|---|---|---|---|---|---|---|---|---|---|---|---|
| **Aphi_Agaz_Ejub** | **Aphi** | 0 | 0 | 0 | 0 | 0 | 0 | 0 | 0 | 0 | 1 | 0.0% | 0.0% |
| **Aphi_Agaz_Ejub** | **Agaz** | 0 | 2.01 | 0.82 | 0.18 | 0.0002 | 0.0002 | -42 | -44 | 2 | 3 | 0.0% | 0.0% |
| **Aphi_Agaz_Ejub** | **Ejub** | 0 | 2.40 | 0.00 | 1.00 | 0.0006 | 0.0015 | -66028 | -60283 | -5745 | 5483 | 99.9% | 0.0% |
| **Aphi_Agaz_Curs** | **Aphi** | 0 | 0 | 0 | 0 | 0 | 0 | 0 | 0 | 0 | 1 | 0.0% | 0.0% |
| **Aphi_Agaz_Curs** | **Agaz** | 0 | 0 | 0 | 0 | 0 | 0 | 0 | 0 | 0 | 1 | 0.0% | 0.0% |
| **Aphi_Agaz_Curs** | **Curs** | 0 | 3.61 | 0.00 | 1.00 | 0.0009 | 0.0035 | -59584 | -51000 | -8584 | 5485 | 100.0% | 0.0% |
| **Aphi_Agaz_Oros** | **Aphi** | 0 | 0 | 0 | 0 | 0 | 0 | 0 | 0 | 0 | 0 | 0.0% | 0.0% |
| **Aphi_Agaz_Oros** | **Agaz** | 0 | 0 | 0 | 0 | 0 | 0 | 0 | 0 | 0 | 0 | 0.0% | 0.0% |
| **Aphi_Agaz_Oros** | **Oros** | 0 | 5.62 | 0.00 | 1.00 | 0.0021 | 0.0126 | -49223 | -37023 | -12200 | 5487 | 100.0% | 0.0% |
| **Aphi_Afor_Agal** | **Aphi** | 0 | 1.01 | 0.02 | 0.98 | 0.0005 | 0.0007 | -69411 | -66636 | -2775 | 5332 | 95.6% | 0.0% |
| **Aphi_Afor_Agal** | **Afor** | 0 | 4.47 | 0.99 | 0.01 | 0.0003 | 0.0003 | -1242 | -1251 | 9 | 89 | 0.0% | 0.0% |
| **Aphi_Afor_Agal** | **Agal** | 0 | 1.39 | 0.95 | 0.05 | 0.0003 | 0.0003 | -950 | -958 | 8 | 66 | 0.0% | 0.0% |
| **Aphi_Afor_Aaus** | **Aphi** | 0 | 1.02 | 0.01 | 0.99 | 0.0005 | 0.0007 | -69443 | -66612 | -2831 | 5343 | 96.2% | 0.0% |
| **Aphi_Afor_Aaus** | **Afor** | 0 | 1.19 | 0.87 | 0.13 | 0.0003 | 0.0003 | -1209 | -1218 | 8 | 86 | 0.0% | 0.0% |
| **Aphi_Afor_Aaus** | **Aaus** | 0 | 0.58 | 0.59 | 0.41 | 0.0002 | 0.0002 | -841 | -847 | 6 | 58 | 0.0% | 0.0% |
| **Aphi_Afor_Phoo** | **Aphi** | 0 | 6.46 | 1.00 | 0.00 | 0.0002 | 0.0002 | -20704 | -20712 | 8 | 1364 | 0.0% | 0.0% |
| **Aphi_Afor_Phoo** | **Afor** | 0 | 6.65 | 1.00 | 0.00 | 0.0002 | 0.0002 | -16341 | -16355 | 14 | 1053 | 0.0% | 19.2% |
| **Aphi_Afor_Phoo** | **Phoo** | 0 | 5.38 | 1.00 | 0.00 | 0.0003 | 0.0003 | -44417 | -44433 | 16 | 3070 | 0.0% | 55.9% |
| **Aphi_Afor_Zwol** | **Aphi** | 0 | 0 | 0 | 0 | 0 | 0 | 0 | 0 | 0 | 1 | 0.0% | 0.0% |
| **Aphi_Afor_Zwol** | **Afor** | 0 | 0.46 | 0.00 | 1.00 | 0.0003 | 0.0003 | -42 | -42 | 1 | 3 | 0.0% | 0.0% |
| **Aphi_Afor_Zwol** | **Zwol** | 0 | 2.43 | 0.00 | 1.00 | 0.0006 | 0.0015 | -66000 | -60249 | -5750 | 5483 | 99.9% | 0.0% |
| **Aphi_Afor_Zcal** | **Aphi** | 0 | 0 | 0 | 0 | 0 | 0 | 0 | 0 | 0 | 1 | 0.0% | 0.0% |
| **Aphi_Afor_Zcal** | **Afor** | 0 | 0.46 | 0.00 | 1.00 | 0.0003 | 0.0003 | -42 | -42 | 1 | 3 | 0.0% | 0.0% |
| **Aphi_Afor_Zcal** | **Zcal** | 0 | 2.43 | 0.00 | 1.00 | 0.0006 | 0.0015 | -66000 | -60249 | -5750 | 5483 | 99.9% | 0.0% |
| **Aphi_Afor_Ejub** | **Aphi** | 0 | 0 | 0 | 0 | 0 | 0 | 0 | 0 | 0 | 1 | 0.0% | 0.0% |
| **Aphi_Afor_Ejub** | **Afor** | 0 | 0.46 | 0.00 | 1.00 | 0.0003 | 0.0003 | -42 | -42 | 1 | 3 | 0.0% | 0.0% |
| **Aphi_Afor_Ejub** | **Ejub** | 0 | 2.43 | 0.00 | 1.00 | 0.0006 | 0.0015 | -66000 | -60249 | -5750 | 5483 | 99.9% | 0.0% |
| **Aphi_Afor_Curs** | **Aphi** | 0 | 12.23 | 0.50 | 0.50 | 0.0005 | 0.0033 | -22 | -18 | -4 | 2 | 0.0% | 0.0% |
| **Aphi_Afor_Curs** | **Afor** | 0 | 0 | 0 | 0 | 0 | 0 | 0 | 0 | 0 | 0 | 0.0% | 0.0% |
| **Aphi_Afor_Curs** | **Curs** | 0 | 3.62 | 0.00 | 1.00 | 0.0009 | 0.0035 | -59565 | -50986 | -8579 | 5485 | 100.0% | 0.0% |
| **Aphi_Afor_Oros** | **Aphi** | 0 | 0 | 0 | 0 | 0 | 0 | 0 | 0 | 0 | 0 | 0.0% | 0.0% |
| **Aphi_Afor_Oros** | **Afor** | 0 | 0 | 0 | 0 | 0 | 0 | 0 | 0 | 0 | 0 | 0.0% | 0.0% |

| | | | | | | | | | | | | | |
|---|---|---|---|---|---|---|---|---|---|---|---|---|---|
| **Aphi_Afor_Oros** | **Oros** | 0 | 5.62 | 0.00 | 1.00 | 0.0021 | 0.0126 | -49213 | -37019 | -12194 | 5487 | 100.0% | 0.0% |
| **Aphi_Agal_Aaus** | **Aphi** | 0 | 1.73 | 0.01 | 0.99 | 0.0005 | 0.0010 | -67866 | -63745 | -4121 | 5408 | 98.0% | 0.0% |
| **Aphi_Agal_Aaus** | **Agal** | 0 | 0.74 | 0.42 | 0.58 | 0.0004 | 0.0004 | -572 | -575 | 3 | 42 | 0.0% | 0.0% |
| **Aphi_Agal_Aaus** | **Aaus** | 0 | 1.88 | 0.94 | 0.06 | 0.0004 | 0.0004 | -492 | -499 | 7 | 37 | 0.0% | 0.0% |
| **Aphi_Agal_Phoo** | **Aphi** | 0 | 5.28 | 1.00 | 0.00 | 0.0002 | 0.0002 | -20754 | -20767 | 13 | 1363 | 0.0% | 24.7% |
| **Aphi_Agal_Phoo** | **Agal** | 0 | 9.60 | 1.00 | 0.00 | 0.0002 | 0.0002 | -16306 | -16318 | 12 | 1050 | 0.0% | 19.1% |
| **Aphi_Agal_Phoo** | **Phoo** | 0 | 5.32 | 1.00 | 0.00 | 0.0003 | 0.0003 | -44403 | -44419 | 16 | 3074 | 0.0% | 56.0% |
| **Aphi_Agal_Zwol** | **Aphi** | 0 | 0 | 0 | 0 | 0 | 0 | 0 | 0 | 0 | 1 | 0.0% | 0.0% |
| **Aphi_Agal_Zwol** | **Agal** | 0 | 0.46 | 0.00 | 1.00 | 0.0003 | 0.0003 | -42 | -42 | 1 | 3 | 0.0% | 0.0% |
| **Aphi_Agal_Zwol** | **Zwol** | 0 | 2.42 | 0.00 | 1.00 | 0.0006 | 0.0015 | -65973 | -60234 | -5739 | 5483 | 99.9% | 0.0% |
| **Aphi_Agal_Zcal** | **Aphi** | 0 | 0 | 0 | 0 | 0 | 0 | 0 | 0 | 0 | 1 | 0.0% | 0.0% |
| **Aphi_Agal_Zcal** | **Agal** | 0 | 0.46 | 0.00 | 1.00 | 0.0003 | 0.0003 | -42 | -42 | 1 | 3 | 0.0% | 0.0% |
| **Aphi_Agal_Zcal** | **Zcal** | 0 | 2.42 | 0.00 | 1.00 | 0.0006 | 0.0015 | -65973 | -60234 | -5739 | 5483 | 99.9% | 0.0% |
| **Aphi_Agal_Ejub** | **Aphi** | 0 | 0 | 0 | 0 | 0 | 0 | 0 | 0 | 0 | 1 | 0.0% | 0.0% |
| **Aphi_Agal_Ejub** | **Agal** | 0 | 0.46 | 0.00 | 1.00 | 0.0003 | 0.0003 | -42 | -42 | 1 | 3 | 0.0% | 0.0% |
| **Aphi_Agal_Ejub** | **Ejub** | 0 | 2.42 | 0.00 | 1.00 | 0.0006 | 0.0015 | -65973 | -60234 | -5739 | 5483 | 99.9% | 0.0% |
| **Aphi_Agal_Curs** | **Aphi** | 0 | 9.83 | 0.50 | 0.50 | 0.0005 | 0.0027 | -22 | -19 | -3 | 2 | 0.0% | 0.0% |
| **Aphi_Agal_Curs** | **Agal** | 0 | 0 | 0 | 0 | 0 | 0 | 0 | 0 | 0 | 0 | 0.0% | 0.0% |
| **Aphi_Agal_Curs** | **Curs** | 0 | 3.62 | 0.00 | 1.00 | 0.0009 | 0.0035 | -59558 | -50979 | -8579 | 5485 | 100.0% | 0.0% |
| **Aphi_Agal_Oros** | **Aphi** | 0 | 0 | 0 | 0 | 0 | 0 | 0 | 0 | 0 | 0 | 0.0% | 0.0% |
| **Aphi_Agal_Oros** | **Agal** | 0 | 0 | 0 | 0 | 0 | 0 | 0 | 0 | 0 | 0 | 0.0% | 0.0% |
| **Aphi_Agal_Oros** | **Oros** | 0 | 5.62 | 0.00 | 1.00 | 0.0021 | 0.0126 | -49212 | -37017 | -12195 | 5487 | 100.0% | 0.0% |
| **Aphi_Aaus_Phoo** | **Aphi** | 0 | 6.53 | 0.99 | 0.01 | 0.0002 | 0.0002 | -20813 | -20820 | 7 | 1369 | 0.0% | 0.0% |
| **Aphi_Aaus_Phoo** | **Aaus** | 0 | 6.76 | 1.00 | 0.00 | 0.0002 | 0.0002 | -16226 | -16240 | 14 | 1046 | 0.0% | 19.0% |
| **Aphi_Aaus_Phoo** | **Phoo** | 0 | 5.22 | 1.00 | 0.00 | 0.0003 | 0.0003 | -44364 | -44380 | 16 | 3072 | 0.0% | 55.9% |
| **Aphi_Aaus_Zwol** | **Aphi** | 0 | 0 | 0 | 0 | 0 | 0 | 0 | 0 | 0 | 1 | 0.0% | 0.0% |
| **Aphi_Aaus_Zwol** | **Aaus** | 0 | 0.46 | 0.00 | 1.00 | 0.0003 | 0.0003 | -42 | -42 | 1 | 3 | 0.0% | 0.0% |
| **Aphi_Aaus_Zwol** | **Zwol** | 0 | 2.42 | 0.00 | 1.00 | 0.0006 | 0.0015 | -65970 | -60233 | -5737 | 5483 | 99.9% | 0.0% |
| **Aphi_Aaus_Zcal** | **Aphi** | 0 | 0 | 0 | 0 | 0 | 0 | 0 | 0 | 0 | 1 | 0.0% | 0.0% |
| **Aphi_Aaus_Zcal** | **Aaus** | 0 | 0.46 | 0.00 | 1.00 | 0.0003 | 0.0003 | -42 | -42 | 1 | 3 | 0.0% | 0.0% |
| **Aphi_Aaus_Zcal** | **Zcal** | 0 | 2.42 | 0.00 | 1.00 | 0.0006 | 0.0015 | -65970 | -60233 | -5737 | 5483 | 99.9% | 0.0% |
| **Aphi_Aaus_Ejub** | **Aphi** | 0 | 0 | 0 | 0 | 0 | 0 | 0 | 0 | 0 | 1 | 0.0% | 0.0% |

| | | | | | | | | | | | | | |
|---|---|---|---|---|---|---|---|---|---|---|---|---|---|
| **Aphi_Aaus_Ejub** | **Aaus** | 0 | 0.46 | 0.00 | 1.00 | 0.0003 | 0.0003 | -42 | -42 | 1 | 3 | 0.0% | 0.0% |
| **Aphi_Aaus_Ejub** | **Ejub** | 0 | 2.42 | 0.00 | 1.00 | 0.0006 | 0.0015 | -65970 | -60233 | -5737 | 5483 | 99.9% | 0.0% |
| **Aphi_Aaus_Curs** | **Aphi** | 0 | 9.83 | 0.50 | 0.50 | 0.0005 | 0.0027 | -22 | -19 | -3 | 2 | 0.0% | 0.0% |
| **Aphi_Aaus_Curs** | **Aaus** | 0 | 0 | 0 | 0 | 0 | 0 | 0 | 0 | 0 | 0 | 0.0% | 0.0% |
| **Aphi_Aaus_Curs** | **Curs** | 0 | 3.62 | 0.00 | 1.00 | 0.0009 | 0.0035 | -59554 | -50979 | -8576 | 5485 | 100.0% | 0.0% |
| **Aphi_Aaus_Oros** | **Aphi** | 0 | 0 | 0 | 0 | 0 | 0 | 0 | 0 | 0 | 0 | 0.0% | 0.0% |
| **Aphi_Aaus_Oros** | **Aaus** | 0 | 0 | 0 | 0 | 0 | 0 | 0 | 0 | 0 | 0 | 0.0% | 0.0% |
| **Aphi_Aaus_Oros** | **Oros** | 0 | 5.62 | 0.00 | 1.00 | 0.0021 | 0.0126 | -49213 | -37017 | -12196 | 5487 | 100.0% | 0.0% |
| **Aphi_Phoo_Zwol** | **Aphi** | 0 | 1.33 | 0.23 | 0.77 | 0.0005 | 0.0006 | -61 | -62 | 1 | 5 | 0.0% | 0.0% |
| **Aphi_Phoo_Zwol** | **Phoo** | 0 | 0.85 | 0.69 | 0.31 | 0.0003 | 0.0003 | -109 | -113 | 4 | 8 | 0.0% | 0.0% |
| **Aphi_Phoo_Zwol** | **Zwol** | 0 | 2.54 | 0.00 | 1.00 | 0.0005 | 0.0014 | -66914 | -61022 | -5892 | 5474 | 99.8% | 0.0% |
| **Aphi_Phoo_Zcal** | **Aphi** | 0 | 1.33 | 0.23 | 0.77 | 0.0005 | 0.0006 | -61 | -62 | 1 | 5 | 0.0% | 0.0% |
| **Aphi_Phoo_Zcal** | **Phoo** | 0 | 0.85 | 0.69 | 0.31 | 0.0003 | 0.0003 | -109 | -113 | 4 | 8 | 0.0% | 0.0% |
| **Aphi_Phoo_Zcal** | **Zcal** | 0 | 2.54 | 0.00 | 1.00 | 0.0005 | 0.0014 | -66914 | -61022 | -5892 | 5474 | 99.8% | 0.0% |
| **Aphi_Phoo_Ejub** | **Aphi** | 0 | 1.33 | 0.23 | 0.77 | 0.0005 | 0.0006 | -61 | -62 | 1 | 5 | 0.0% | 0.0% |
| **Aphi_Phoo_Ejub** | **Phoo** | 0 | 0.85 | 0.69 | 0.31 | 0.0003 | 0.0003 | -109 | -113 | 4 | 8 | 0.0% | 0.0% |
| **Aphi_Phoo_Ejub** | **Ejub** | 0 | 2.54 | 0.00 | 1.00 | 0.0005 | 0.0014 | -66914 | -61022 | -5892 | 5474 | 99.8% | 0.0% |
| **Aphi_Phoo_Curs** | **Aphi** | 0 | 0 | 0 | 0 | 0 | 0 | 0 | 0 | 0 | 1 | 0.0% | 0.0% |
| **Aphi_Phoo_Curs** | **Phoo** | 0 | 0 | 0 | 0 | 0 | 0 | 0 | 0 | 0 | 1 | 0.0% | 0.0% |
| **Aphi_Phoo_Curs** | **Curs** | 0 | 3.75 | 0.00 | 1.00 | 0.0008 | 0.0034 | -60095 | -51360 | -8735 | 5485 | 100.0% | 0.0% |
| **Aphi_Phoo_Oros** | **Aphi** | 0 | 0 | 0 | 0 | 0 | 0 | 0 | 0 | 0 | 0 | 0.0% | 0.0% |
| **Aphi_Phoo_Oros** | **Phoo** | 0 | 0 | 0 | 0 | 0 | 0 | 0 | 0 | 0 | 0 | 0.0% | 0.0% |
| **Aphi_Phoo_Oros** | **Oros** | 0 | 5.70 | 0.00 | 1.00 | 0.0021 | 0.0125 | -49434 | -37122 | -12312 | 5487 | 100.0% | 0.0% |
| **Aphi_Zwol_Zcal** | **Aphi** | 0 | 3.97 | 0.00 | 1.00 | 0.0006 | 0.0026 | -63346 | -54297 | -9049 | 5487 | 100.0% | 0.0% |
| **Aphi_Zwol_Zcal** | **Zwol** | 0 | 0 | 0 | 0 | 0 | 0 | 0 | 0 | 0 | 0 | 0.0% | 0.0% |
| **Aphi_Zwol_Zcal** | **Zcal** | 0 | 0 | 0 | 0 | 0 | 0 | 0 | 0 | 0 | 0 | 0.0% | 0.0% |
| **Aphi_Zwol_Ejub** | **Aphi** | 0 | 3.98 | 0.00 | 1.00 | 0.0006 | 0.0026 | -63347 | -54308 | -9038 | 5487 | 100.0% | 0.0% |
| **Aphi_Zwol_Ejub** | **Zwol** | 0 | 0 | 0 | 0 | 0 | 0 | 0 | 0 | 0 | 0 | 0.0% | 0.0% |
| **Aphi_Zwol_Ejub** | **Ejub** | 0 | 0 | 0 | 0 | 0 | 0 | 0 | 0 | 0 | 0 | 0.0% | 0.0% |
| **Aphi_Zwol_Curs** | **Aphi** | 0 | 0 | 0 | 0 | 0 | 0 | 0 | 0 | 0 | 1 | 0.0% | 0.0% |
| **Aphi_Zwol_Curs** | **Zwol** | 0 | 1.56 | 0.04 | 0.96 | 0.0005 | 0.0008 | -35 | -35 | 1 | 3 | 0.0% | 0.0% |
| **Aphi_Zwol_Curs** | **Curs** | 0 | 3.13 | 0.00 | 1.00 | 0.0006 | 0.0020 | -64468 | -57100 | -7368 | 5483 | 99.9% | 0.0% |

| | | | | | | | | | | | | | |
|---|---|---|---|---|---|---|---|---|---|---|---|---|---|
| **Aphi_Zwol_Oros** | **Aphi** | 0 | 0 | 0 | 0 | 0 | 0 | 0 | 0 | 0 | 0 | 0.0% | 0.0% |
| **Aphi_Zwol_Oros** | **Zwol** | 0 | 0 | 0 | 0 | 0 | 0 | 0 | 0 | 0 | 0 | 0.0% | 0.0% |
| **Aphi_Zwol_Oros** | **Oros** | 0 | 5.89 | 0.00 | 1.00 | 0.0018 | 0.0111 | -51053 | -38420 | -12634 | 5487 | 100.0% | 0.0% |
| **Aphi_Zcal_Ejub** | **Aphi** | 0 | 3.97 | 0.00 | 1.00 | 0.0006 | 0.0027 | -63115 | -54048 | -9067 | 5487 | 100.0% | 0.0% |
| **Aphi_Zcal_Ejub** | **Zcal** | 0 | 0 | 0 | 0 | 0 | 0 | 0 | 0 | 0 | 0 | 0.0% | 0.0% |
| **Aphi_Zcal_Ejub** | **Ejub** | 0 | 0 | 0 | 0 | 0 | 0 | 0 | 0 | 0 | 0 | 0.0% | 0.0% |
| **Aphi_Zcal_Curs** | **Aphi** | 0 | 0 | 0 | 0 | 0 | 0 | 0 | 0 | 0 | 1 | 0.0% | 0.0% |
| **Aphi_Zcal_Curs** | **Zcal** | 0 | 1.56 | 0.04 | 0.96 | 0.0005 | 0.0008 | -35 | -35 | 1 | 3 | 0.0% | 0.0% |
| **Aphi_Zcal_Curs** | **Curs** | 0 | 3.13 | 0.00 | 1.00 | 0.0006 | 0.0020 | -64468 | -57100 | -7368 | 5483 | 99.9% | 0.0% |
| **Aphi_Zcal_Oros** | **Aphi** | 0 | 0 | 0 | 0 | 0 | 0 | 0 | 0 | 0 | 0 | 0.0% | 0.0% |
| **Aphi_Zcal_Oros** | **Zcal** | 0 | 0 | 0 | 0 | 0 | 0 | 0 | 0 | 0 | 0 | 0.0% | 0.0% |
| **Aphi_Zcal_Oros** | **Oros** | 0 | 5.89 | 0.00 | 1.00 | 0.0018 | 0.0111 | -51053 | -38420 | -12634 | 5487 | 100.0% | 0.0% |
| **Aphi_Ejub_Curs** | **Aphi** | 0 | 0 | 0 | 0 | 0 | 0 | 0 | 0 | 0 | 1 | 0.0% | 0.0% |
| **Aphi_Ejub_Curs** | **Ejub** | 0 | 1.56 | 0.04 | 0.96 | 0.0005 | 0.0008 | -35 | -35 | 1 | 3 | 0.0% | 0.0% |
| **Aphi_Ejub_Curs** | **Curs** | 0 | 3.13 | 0.00 | 1.00 | 0.0006 | 0.0020 | -64468 | -57100 | -7368 | 5483 | 99.9% | 0.0% |
| **Aphi_Ejub_Oros** | **Aphi** | 0 | 0 | 0 | 0 | 0 | 0 | 0 | 0 | 0 | 0 | 0.0% | 0.0% |
| **Aphi_Ejub_Oros** | **Ejub** | 0 | 0 | 0 | 0 | 0 | 0 | 0 | 0 | 0 | 0 | 0.0% | 0.0% |
| **Aphi_Ejub_Oros** | **Oros** | 0 | 5.89 | 0.00 | 1.00 | 0.0018 | 0.0111 | -51053 | -38420 | -12634 | 5487 | 100.0% | 0.0% |
| **Aphi_Curs_Oros** | **Aphi** | 0 | 0 | 0 | 0 | 0 | 0 | 0 | 0 | 0 | 0 | 0.0% | 0.0% |
| **Aphi_Curs_Oros** | **Curs** | 0 | 0 | 0 | 0 | 0 | 0 | 0 | 0 | 0 | 0 | 0.0% | 0.0% |
| **Aphi_Curs_Oros** | **Oros** | 0 | 6.17 | 0.00 | 1.00 | 0.0014 | 0.0091 | -53695 | -40616 | -13079 | 5487 | 100.0% | 0.0% |
| **Agaz_Afor_Agal** | **Agaz** | 0 | 0.90 | 0.02 | 0.98 | 0.0005 | 0.0007 | -69519 | -67006 | -2513 | 5306 | 95.0% | 0.0% |
| **Agaz_Afor_Agal** | **Afor** | 0 | 0.46 | 0.32 | 0.68 | 0.0003 | 0.0003 | -1401 | -1401 | 0 | 98 | 0.0% | 0.0% |
| **Agaz_Afor_Agal** | **Agal** | 0 | 1.40 | 0.85 | 0.15 | 0.0002 | 0.0002 | -1228 | -1236 | 7 | 83 | 0.0% | 0.0% |
| **Agaz_Afor_Aaus** | **Agaz** | 0 | 0.90 | 0.01 | 0.99 | 0.0005 | 0.0007 | -69644 | -67076 | -2568 | 5324 | 96.1% | 0.0% |
| **Agaz_Afor_Aaus** | **Afor** | 0 | 0.78 | 0.62 | 0.38 | 0.0002 | 0.0002 | -1369 | -1373 | 4 | 94 | 0.0% | 0.0% |
| **Agaz_Afor_Aaus** | **Aaus** | 0 | 1.46 | 0.86 | 0.14 | 0.0002 | 0.0002 | -1025 | -1032 | 7 | 69 | 0.0% | 0.0% |
| **Agaz_Afor_Phoo** | **Agaz** | 0 | 20.50 | 1.00 | 0.00 | 0.0002 | 0.0002 | -15815 | -15779 | -37 | 1046 | 0.0% | 0.0% |
| **Agaz_Afor_Phoo** | **Afor** | 0 | 5.04 | 1.00 | 0.00 | 0.0002 | 0.0002 | -15620 | -15634 | 14 | 1028 | 0.0% | 18.7% |
| **Agaz_Afor_Phoo** | **Phoo** | 0 | 6.52 | 1.00 | 0.00 | 0.0003 | 0.0003 | -48539 | -48555 | 16 | 3413 | 0.0% | 62.2% |
| **Agaz_Afor_Zwol** | **Agaz** | 0 | 0 | 0 | 0 | 0 | 0 | 0 | 0 | 0 | 1 | 0.0% | 0.0% |
| **Agaz_Afor_Zwol** | **Afor** | 0 | 1.44 | 0.57 | 0.43 | 0.0004 | 0.0004 | -25 | -26 | 1 | 2 | 0.0% | 0.0% |

| | | | | | | | | | | | | | |
|---|---|---|---|---|---|---|---|---|---|---|---|---|---|
| **Agaz_Afor_Zwol** | **Zwol** | 0 | 2.40 | 0.00 | 1.00 | 0.0006 | 0.0016 | -65730 | -59926 | -5805 | 5484 | 99.9% | 0.0% |
| **Agaz_Afor_Zcal** | **Agaz** | 0 | 0 | 0 | 0 | 0 | 0 | 0 | 0 | 0 | 1 | 0.0% | 0.0% |
| **Agaz_Afor_Zcal** | **Afor** | 0 | 1.44 | 0.57 | 0.43 | 0.0004 | 0.0004 | -25 | -26 | 1 | 2 | 0.0% | 0.0% |
| **Agaz_Afor_Zcal** | **Zcal** | 0 | 2.40 | 0.00 | 1.00 | 0.0006 | 0.0016 | -65730 | -59926 | -5805 | 5484 | 99.9% | 0.0% |
| **Agaz_Afor_Ejub** | **Agaz** | 0 | 0 | 0 | 0 | 0 | 0 | 0 | 0 | 0 | 1 | 0.0% | 0.0% |
| **Agaz_Afor_Ejub** | **Afor** | 0 | 1.44 | 0.57 | 0.43 | 0.0004 | 0.0004 | -25 | -26 | 1 | 2 | 0.0% | 0.0% |
| **Agaz_Afor_Ejub** | **Ejub** | 0 | 2.40 | 0.00 | 1.00 | 0.0006 | 0.0016 | -65730 | -59926 | -5805 | 5484 | 99.9% | 0.0% |
| **Agaz_Afor_Curs** | **Agaz** | 0 | 7.08 | 0.50 | 0.50 | 0.0005 | 0.0021 | -22 | -20 | -2 | 2 | 0.0% | 0.0% |
| **Agaz_Afor_Curs** | **Afor** | 0 | 0 | 0 | 0 | 0 | 0 | 0 | 0 | 0 | 0 | 0.0% | 0.0% |
| **Agaz_Afor_Curs** | **Curs** | 0 | 3.62 | 0.00 | 1.00 | 0.0009 | 0.0036 | -59420 | -50841 | -8579 | 5485 | 100.0% | 0.0% |
| **Agaz_Afor_Oros** | **Agaz** | 0 | 0 | 0 | 0 | 0 | 0 | 0 | 0 | 0 | 0 | 0.0% | 0.0% |
| **Agaz_Afor_Oros** | **Afor** | 0 | 0 | 0 | 0 | 0 | 0 | 0 | 0 | 0 | 0 | 0.0% | 0.0% |
| **Agaz_Afor_Oros** | **Oros** | 0 | 5.61 | 0.00 | 1.00 | 0.0021 | 0.0126 | -49165 | -36978 | -12187 | 5487 | 100.0% | 0.0% |
| **Agaz_Agal_Aaus** | **Agaz** | 0 | 1.64 | 0.01 | 0.99 | 0.0005 | 0.0010 | -68162 | -64176 | -3986 | 5401 | 97.7% | 0.0% |
| **Agaz_Agal_Aaus** | **Agal** | 0 | 5.34 | 0.99 | 0.01 | 0.0002 | 0.0002 | -692 | -699 | 7 | 47 | 0.0% | 0.0% |
| **Agaz_Agal_Aaus** | **Aaus** | 0 | 0.43 | 0.74 | 0.26 | 0.0003 | 0.0003 | -547 | -554 | 7 | 39 | 0.0% | 0.0% |
| **Agaz_Agal_Phoo** | **Agaz** | 0 | 12.61 | 1.00 | 0.00 | 0.0002 | 0.0002 | -15823 | -15829 | 6 | 1043 | 0.0% | 0.0% |
| **Agaz_Agal_Phoo** | **Agal** | 0 | 9.53 | 1.00 | 0.00 | 0.0002 | 0.0002 | -15549 | -15560 | 12 | 1021 | 0.0% | 18.6% |
| **Agaz_Agal_Phoo** | **Phoo** | 0 | 6.50 | 1.00 | 0.00 | 0.0003 | 0.0003 | -48661 | -48677 | 16 | 3423 | 0.0% | 62.4% |
| **Agaz_Agal_Zwol** | **Agaz** | 0 | 0 | 0 | 0 | 0 | 0 | 0 | 0 | 0 | 1 | 0.0% | 0.0% |
| **Agaz_Agal_Zwol** | **Agal** | 0 | 1.44 | 0.57 | 0.43 | 0.0004 | 0.0004 | -25 | -26 | 1 | 2 | 0.0% | 0.0% |
| **Agaz_Agal_Zwol** | **Zwol** | 0 | 2.40 | 0.00 | 1.00 | 0.0006 | 0.0016 | -65713 | -59912 | -5800 | 5484 | 99.9% | 0.0% |
| **Agaz_Agal_Zcal** | **Agaz** | 0 | 0 | 0 | 0 | 0 | 0 | 0 | 0 | 0 | 1 | 0.0% | 0.0% |
| **Agaz_Agal_Zcal** | **Agal** | 0 | 1.44 | 0.57 | 0.43 | 0.0004 | 0.0004 | -25 | -26 | 1 | 2 | 0.0% | 0.0% |
| **Agaz_Agal_Zcal** | **Zcal** | 0 | 2.40 | 0.00 | 1.00 | 0.0006 | 0.0016 | -65713 | -59912 | -5800 | 5484 | 99.9% | 0.0% |
| **Agaz_Agal_Ejub** | **Agaz** | 0 | 0 | 0 | 0 | 0 | 0 | 0 | 0 | 0 | 1 | 0.0% | 0.0% |
| **Agaz_Agal_Ejub** | **Agal** | 0 | 1.44 | 0.57 | 0.43 | 0.0004 | 0.0004 | -25 | -26 | 1 | 2 | 0.0% | 0.0% |
| **Agaz_Agal_Ejub** | **Ejub** | 0 | 2.40 | 0.00 | 1.00 | 0.0006 | 0.0016 | -65713 | -59912 | -5800 | 5484 | 99.9% | 0.0% |
| **Agaz_Agal_Curs** | **Agaz** | 0 | 4.68 | 0.48 | 0.52 | 0.0005 | 0.0015 | -22 | -21 | 0 | 2 | 0.0% | 0.0% |
| **Agaz_Agal_Curs** | **Agal** | 0 | 0 | 0 | 0 | 0 | 0 | 0 | 0 | 0 | 0 | 0.0% | 0.0% |
| **Agaz_Agal_Curs** | **Curs** | 0 | 3.62 | 0.00 | 1.00 | 0.0009 | 0.0036 | -59401 | -50836 | -8566 | 5485 | 100.0% | 0.0% |
| **Agaz_Agal_Oros** | **Agaz** | 0 | 0 | 0 | 0 | 0 | 0 | 0 | 0 | 0 | 0 | 0.0% | 0.0% |

| | | | | | | | | | | | | | |
|---|---|---|---|---|---|---|---|---|---|---|---|---|---|
| **Agaz_Agal_Oros** | **Agal** | 0 | 0 | 0 | 0 | 0 | 0 | 0 | 0 | 0 | 0 | 0.0% | 0.0% |
| **Agaz_Agal_Oros** | **Oros** | 0 | 5.61 | 0.00 | 1.00 | 0.0021 | 0.0126 | -49162 | -36976 | -12186 | 5487 | 100.0% | 0.0% |
| **Agaz_Aaus_Phoo** | **Agaz** | 0 | 26.93 | 1.00 | 0.00 | 0.0002 | 0.0002 | -15875 | -15853 | -23 | 1047 | 0.0% | 0.0% |
| **Agaz_Aaus_Phoo** | **Aaus** | 0 | 9.50 | 1.00 | 0.00 | 0.0002 | 0.0002 | -15510 | -15523 | 13 | 1019 | 0.0% | 18.6% |
| **Agaz_Aaus_Phoo** | **Phoo** | 0 | 6.33 | 1.00 | 0.00 | 0.0003 | 0.0003 | -48637 | -48653 | 16 | 3421 | 0.0% | 62.3% |
| **Agaz_Aaus_Zwol** | **Agaz** | 0 | 0 | 0 | 0 | 0 | 0 | 0 | 0 | 0 | 1 | 0.0% | 0.0% |
| **Agaz_Aaus_Zwol** | **Aaus** | 0 | 1.44 | 0.57 | 0.43 | 0.0004 | 0.0004 | -25 | -26 | 1 | 2 | 0.0% | 0.0% |
| **Agaz_Aaus_Zwol** | **Zwol** | 0 | 2.40 | 0.00 | 1.00 | 0.0006 | 0.0016 | -65719 | -59914 | -5805 | 5484 | 99.9% | 0.0% |
| **Agaz_Aaus_Zcal** | **Agaz** | 0 | 0 | 0 | 0 | 0 | 0 | 0 | 0 | 0 | 1 | 0.0% | 0.0% |
| **Agaz_Aaus_Zcal** | **Aaus** | 0 | 1.44 | 0.57 | 0.43 | 0.0004 | 0.0004 | -25 | -26 | 1 | 2 | 0.0% | 0.0% |
| **Agaz_Aaus_Zcal** | **Zcal** | 0 | 2.40 | 0.00 | 1.00 | 0.0006 | 0.0016 | -65719 | -59914 | -5805 | 5484 | 99.9% | 0.0% |
| **Agaz_Aaus_Ejub** | **Agaz** | 0 | 0 | 0 | 0 | 0 | 0 | 0 | 0 | 0 | 1 | 0.0% | 0.0% |
| **Agaz_Aaus_Ejub** | **Aaus** | 0 | 1.44 | 0.57 | 0.43 | 0.0004 | 0.0004 | -25 | -26 | 1 | 2 | 0.0% | 0.0% |
| **Agaz_Aaus_Ejub** | **Ejub** | 0 | 2.40 | 0.00 | 1.00 | 0.0006 | 0.0016 | -65719 | -59914 | -5805 | 5484 | 99.9% | 0.0% |
| **Agaz_Aaus_Curs** | **Agaz** | 0 | 4.68 | 0.48 | 0.52 | 0.0005 | 0.0015 | -22 | -21 | 0 | 2 | 0.0% | 0.0% |
| **Agaz_Aaus_Curs** | **Aaus** | 0 | 0 | 0 | 0 | 0 | 0 | 0 | 0 | 0 | 0 | 0.0% | 0.0% |
| **Agaz_Aaus_Curs** | **Curs** | 0 | 3.61 | 0.00 | 1.00 | 0.0009 | 0.0036 | -59404 | -50836 | -8568 | 5485 | 100.0% | 0.0% |
| **Agaz_Aaus_Oros** | **Agaz** | 0 | 0 | 0 | 0 | 0 | 0 | 0 | 0 | 0 | 0 | 0.0% | 0.0% |
| **Agaz_Aaus_Oros** | **Aaus** | 0 | 0 | 0 | 0 | 0 | 0 | 0 | 0 | 0 | 0 | 0.0% | 0.0% |
| **Agaz_Aaus_Oros** | **Oros** | 0 | 5.61 | 0.00 | 1.00 | 0.0021 | 0.0126 | -49164 | -36977 | -12187 | 5487 | 100.0% | 0.0% |
| **Agaz_Phoo_Zwol** | **Agaz** | 0 | 1.33 | 0.23 | 0.77 | 0.0005 | 0.0006 | -61 | -62 | 1 | 5 | 0.0% | 0.0% |
| **Agaz_Phoo_Zwol** | **Phoo** | 0 | 0.65 | 0.41 | 0.59 | 0.0004 | 0.0004 | -79 | -81 | 3 | 6 | 0.0% | 0.0% |
| **Agaz_Phoo_Zwol** | **Zwol** | 0 | 2.56 | 0.00 | 1.00 | 0.0005 | 0.0014 | -66901 | -60944 | -5957 | 5476 | 99.8% | 0.0% |
| **Agaz_Phoo_Zcal** | **Agaz** | 0 | 1.33 | 0.23 | 0.77 | 0.0005 | 0.0006 | -61 | -62 | 1 | 5 | 0.0% | 0.0% |
| **Agaz_Phoo_Zcal** | **Phoo** | 0 | 0.65 | 0.41 | 0.59 | 0.0004 | 0.0004 | -79 | -81 | 3 | 6 | 0.0% | 0.0% |
| **Agaz_Phoo_Zcal** | **Zcal** | 0 | 2.56 | 0.00 | 1.00 | 0.0005 | 0.0014 | -66901 | -60944 | -5957 | 5476 | 99.8% | 0.0% |
| **Agaz_Phoo_Ejub** | **Agaz** | 0 | 1.33 | 0.23 | 0.77 | 0.0005 | 0.0006 | -61 | -62 | 1 | 5 | 0.0% | 0.0% |
| **Agaz_Phoo_Ejub** | **Phoo** | 0 | 0.65 | 0.41 | 0.59 | 0.0004 | 0.0004 | -79 | -81 | 3 | 6 | 0.0% | 0.0% |
| **Agaz_Phoo_Ejub** | **Ejub** | 0 | 2.56 | 0.00 | 1.00 | 0.0005 | 0.0014 | -66901 | -60944 | -5957 | 5476 | 99.8% | 0.0% |
| **Agaz_Phoo_Curs** | **Agaz** | 0 | 0 | 0 | 0 | 0 | 0 | 0 | 0 | 0 | 1 | 0.0% | 0.0% |
| **Agaz_Phoo_Curs** | **Phoo** | 0 | 0 | 0 | 0 | 0 | 0 | 0 | 0 | 0 | 0 | 0.0% | 0.0% |
| **Agaz_Phoo_Curs** | **Curs** | 0 | 3.76 | 0.00 | 1.00 | 0.0008 | 0.0034 | -60090 | -51330 | -8760 | 5486 | 100.0% | 0.0% |

| | | | | | | | | | | | | | |
|---|---|---|---|---|---|---|---|---|---|---|---|---|---|
| **Agaz_Phoo_Oros** | **Agaz** | 0 | 0 | 0 | 0 | 0 | 0 | 0 | 0 | 0 | 0 | 0.0% | 0.0% |
| **Agaz_Phoo_Oros** | **Phoo** | 0 | 0 | 0 | 0 | 0 | 0 | 0 | 0 | 0 | 0 | 0.0% | 0.0% |
| **Agaz_Phoo_Oros** | **Oros** | 0 | 5.70 | 0.00 | 1.00 | 0.0021 | 0.0125 | -49438 | -37111 | -12327 | 5487 | 100.0% | 0.0% |
| **Agaz_Zwol_Zcal** | **Agaz** | 0 | 3.97 | 0.00 | 1.00 | 0.0006 | 0.0026 | -63347 | -54297 | -9051 | 5487 | 100.0% | 0.0% |
| **Agaz_Zwol_Zcal** | **Zwol** | 0 | 0 | 0 | 0 | 0 | 0 | 0 | 0 | 0 | 0 | 0.0% | 0.0% |
| **Agaz_Zwol_Zcal** | **Zcal** | 0 | 0 | 0 | 0 | 0 | 0 | 0 | 0 | 0 | 0 | 0.0% | 0.0% |
| **Agaz_Zwol_Ejub** | **Agaz** | 0 | 3.98 | 0.00 | 1.00 | 0.0006 | 0.0026 | -63348 | -54308 | -9039 | 5487 | 100.0% | 0.0% |
| **Agaz_Zwol_Ejub** | **Zwol** | 0 | 0 | 0 | 0 | 0 | 0 | 0 | 0 | 0 | 0 | 0.0% | 0.0% |
| **Agaz_Zwol_Ejub** | **Ejub** | 0 | 0 | 0 | 0 | 0 | 0 | 0 | 0 | 0 | 0 | 0.0% | 0.0% |
| **Agaz_Zwol_Curs** | **Agaz** | 0 | 0 | 0 | 0 | 0 | 0 | 0 | 0 | 0 | 1 | 0.0% | 0.0% |
| **Agaz_Zwol_Curs** | **Zwol** | 0 | 7.89 | 0.67 | 0.33 | 0.0005 | 0.0017 | -33 | -31 | -1 | 3 | 0.0% | 0.0% |
| **Agaz_Zwol_Curs** | **Curs** | 0 | 3.13 | 0.00 | 1.00 | 0.0006 | 0.0020 | -64465 | -57100 | -7365 | 5483 | 99.9% | 0.0% |
| **Agaz_Zwol_Oros** | **Agaz** | 0 | 0 | 0 | 0 | 0 | 0 | 0 | 0 | 0 | 0 | 0.0% | 0.0% |
| **Agaz_Zwol_Oros** | **Zwol** | 0 | 0 | 0 | 0 | 0 | 0 | 0 | 0 | 0 | 0 | 0.0% | 0.0% |
| **Agaz_Zwol_Oros** | **Oros** | 0 | 5.89 | 0.00 | 1.00 | 0.0018 | 0.0111 | -51053 | -38420 | -12633 | 5487 | 100.0% | 0.0% |
| **Agaz_Zcal_Ejub** | **Agaz** | 0 | 3.97 | 0.00 | 1.00 | 0.0006 | 0.0027 | -63115 | -54048 | -9068 | 5487 | 100.0% | 0.0% |
| **Agaz_Zcal_Ejub** | **Zcal** | 0 | 0 | 0 | 0 | 0 | 0 | 0 | 0 | 0 | 0 | 0.0% | 0.0% |
| **Agaz_Zcal_Ejub** | **Ejub** | 0 | 0 | 0 | 0 | 0 | 0 | 0 | 0 | 0 | 0 | 0.0% | 0.0% |
| **Agaz_Zcal_Curs** | **Agaz** | 0 | 0 | 0 | 0 | 0 | 0 | 0 | 0 | 0 | 1 | 0.0% | 0.0% |
| **Agaz_Zcal_Curs** | **Zcal** | 0 | 7.89 | 0.67 | 0.33 | 0.0005 | 0.0017 | -33 | -31 | -1 | 3 | 0.0% | 0.0% |
| **Agaz_Zcal_Curs** | **Curs** | 0 | 3.13 | 0.00 | 1.00 | 0.0006 | 0.0020 | -64465 | -57100 | -7365 | 5483 | 99.9% | 0.0% |
| **Agaz_Zcal_Oros** | **Agaz** | 0 | 0 | 0 | 0 | 0 | 0 | 0 | 0 | 0 | 0 | 0.0% | 0.0% |
| **Agaz_Zcal_Oros** | **Zcal** | 0 | 0 | 0 | 0 | 0 | 0 | 0 | 0 | 0 | 0 | 0.0% | 0.0% |
| **Agaz_Zcal_Oros** | **Oros** | 0 | 5.89 | 0.00 | 1.00 | 0.0018 | 0.0111 | -51053 | -38420 | -12633 | 5487 | 100.0% | 0.0% |
| **Agaz_Ejub_Curs** | **Agaz** | 0 | 0 | 0 | 0 | 0 | 0 | 0 | 0 | 0 | 1 | 0.0% | 0.0% |
| **Agaz_Ejub_Curs** | **Ejub** | 0 | 7.89 | 0.67 | 0.33 | 0.0005 | 0.0017 | -33 | -31 | -1 | 3 | 0.0% | 0.0% |
| **Agaz_Ejub_Curs** | **Curs** | 0 | 3.13 | 0.00 | 1.00 | 0.0006 | 0.0020 | -64465 | -57100 | -7365 | 5483 | 99.9% | 0.0% |
| **Agaz_Ejub_Oros** | **Agaz** | 0 | 0 | 0 | 0 | 0 | 0 | 0 | 0 | 0 | 0 | 0.0% | 0.0% |
| **Agaz_Ejub_Oros** | **Ejub** | 0 | 0 | 0 | 0 | 0 | 0 | 0 | 0 | 0 | 0 | 0.0% | 0.0% |
| **Agaz_Ejub_Oros** | **Oros** | 0 | 5.89 | 0.00 | 1.00 | 0.0018 | 0.0111 | -51053 | -38420 | -12633 | 5487 | 100.0% | 0.0% |
| **Agaz_Curs_Oros** | **Agaz** | 0 | 0 | 0 | 0 | 0 | 0 | 0 | 0 | 0 | 0 | 0.0% | 0.0% |
| **Agaz_Curs_Oros** | **Curs** | 0 | 0 | 0 | 0 | 0 | 0 | 0 | 0 | 0 | 0 | 0.0% | 0.0% |

| | | | | | | | | | | | | | |
|---|---|---|---|---|---|---|---|---|---|---|---|---|---|
| **Agaz_Curs_Oros** | **Oros** | 0 | 6.17 | 0.00 | 1.00 | 0.0014 | 0.0091 | -53692 | -40616 | -13076 | 5487 | 100.0% | 0.0% |
| **Afor_Agal_Aaus** | **Afor** | 0 | 0.58 | 0.03 | 0.97 | 0.0004 | 0.0004 | -58212 | -56757 | -1455 | 4222 | 74.9% | 0.0% |
| **Afor_Agal_Aaus** | **Agal** | 0 | 0.54 | 0.05 | 0.95 | 0.0003 | 0.0003 | -10091 | -9903 | -187 | 706 | 12.2% | 0.0% |
| **Afor_Agal_Aaus** | **Aaus** | 0 | 0.62 | 0.08 | 0.92 | 0.0003 | 0.0003 | -8122 | -7968 | -154 | 559 | 9.4% | 0.0% |
| **Afor_Agal_Phoo** | **Afor** | 0 | 4.17 | 0.96 | 0.04 | 0.0002 | 0.0002 | -662 | -668 | 7 | 46 | 0.0% | 0.0% |
| **Afor_Agal_Phoo** | **Agal** | 0 | 7.36 | 0.94 | 0.06 | 0.0004 | 0.0004 | -657 | -648 | -9 | 47 | 0.0% | 0.0% |
| **Afor_Agal_Phoo** | **Phoo** | 0 | 1.20 | 0.01 | 0.99 | 0.0005 | 0.0008 | -69115 | -66049 | -3066 | 5394 | 97.0% | 0.0% |
| **Afor_Agal_Zwol** | **Afor** | 0 | 0 | 0 | 0 | 0 | 0 | 0 | 0 | 0 | 0 | 0.0% | 0.0% |
| **Afor_Agal_Zwol** | **Agal** | 0 | 0 | 0 | 0 | 0 | 0 | 0 | 0 | 0 | 0 | 0.0% | 0.0% |
| **Afor_Agal_Zwol** | **Zwol** | 0 | 2.89 | 0.00 | 1.00 | 0.0007 | 0.0022 | -63277 | -56184 | -7093 | 5487 | 100.0% | 0.0% |
| **Afor_Agal_Zcal** | **Afor** | 0 | 0 | 0 | 0 | 0 | 0 | 0 | 0 | 0 | 0 | 0.0% | 0.0% |
| **Afor_Agal_Zcal** | **Agal** | 0 | 0 | 0 | 0 | 0 | 0 | 0 | 0 | 0 | 0 | 0.0% | 0.0% |
| **Afor_Agal_Zcal** | **Zcal** | 0 | 2.89 | 0.00 | 1.00 | 0.0007 | 0.0022 | -63277 | -56184 | -7093 | 5487 | 100.0% | 0.0% |
| **Afor_Agal_Ejub** | **Afor** | 0 | 0 | 0 | 0 | 0 | 0 | 0 | 0 | 0 | 0 | 0.0% | 0.0% |
| **Afor_Agal_Ejub** | **Agal** | 0 | 0 | 0 | 0 | 0 | 0 | 0 | 0 | 0 | 0 | 0.0% | 0.0% |
| **Afor_Agal_Ejub** | **Ejub** | 0 | 2.89 | 0.00 | 1.00 | 0.0007 | 0.0022 | -63277 | -56184 | -7093 | 5487 | 100.0% | 0.0% |
| **Afor_Agal_Curs** | **Afor** | 0 | 0 | 0 | 0 | 0 | 0 | 0 | 0 | 0 | 0 | 0.0% | 0.0% |
| **Afor_Agal_Curs** | **Agal** | 0 | 0 | 0 | 0 | 0 | 0 | 0 | 0 | 0 | 1 | 0.0% | 0.0% |
| **Afor_Agal_Curs** | **Curs** | 0 | 3.80 | 0.00 | 1.00 | 0.0010 | 0.0042 | -58046 | -49043 | -9003 | 5486 | 100.0% | 0.0% |
| **Afor_Agal_Oros** | **Afor** | 0 | 0 | 0 | 0 | 0 | 0 | 0 | 0 | 0 | 0 | 0.0% | 0.0% |
| **Afor_Agal_Oros** | **Agal** | 0 | 0 | 0 | 0 | 0 | 0 | 0 | 0 | 0 | 0 | 0.0% | 0.0% |
| **Afor_Agal_Oros** | **Oros** | 0 | 5.59 | 0.00 | 1.00 | 0.0022 | 0.0133 | -48603 | -36436 | -12167 | 5487 | 100.0% | 0.0% |
| **Afor_Aaus_Phoo** | **Afor** | 0 | 4.60 | 0.93 | 0.07 | 0.0002 | 0.0002 | -651 | -656 | 5 | 45 | 0.0% | 0.0% |
| **Afor_Aaus_Phoo** | **Aaus** | 0 | 7.58 | 0.95 | 0.05 | 0.0003 | 0.0003 | -520 | -516 | -5 | 37 | 0.0% | 0.0% |
| **Afor_Aaus_Phoo** | **Phoo** | 0 | 1.22 | 0.01 | 0.99 | 0.0005 | 0.0008 | -69144 | -66049 | -3095 | 5405 | 97.4% | 0.0% |
| **Afor_Aaus_Zwol** | **Afor** | 0 | 0 | 0 | 0 | 0 | 0 | 0 | 0 | 0 | 0 | 0.0% | 0.0% |
| **Afor_Aaus_Zwol** | **Aaus** | 0 | 0 | 0 | 0 | 0 | 0 | 0 | 0 | 0 | 0 | 0.0% | 0.0% |
| **Afor_Aaus_Zwol** | **Zwol** | 0 | 2.90 | 0.00 | 1.00 | 0.0007 | 0.0022 | -63235 | -56122 | -7113 | 5487 | 100.0% | 0.0% |
| **Afor_Aaus_Zcal** | **Afor** | 0 | 0 | 0 | 0 | 0 | 0 | 0 | 0 | 0 | 0 | 0.0% | 0.0% |
| **Afor_Aaus_Zcal** | **Aaus** | 0 | 0 | 0 | 0 | 0 | 0 | 0 | 0 | 0 | 0 | 0.0% | 0.0% |
| **Afor_Aaus_Zcal** | **Zcal** | 0 | 2.90 | 0.00 | 1.00 | 0.0007 | 0.0022 | -63235 | -56122 | -7113 | 5487 | 100.0% | 0.0% |
| **Afor_Aaus_Ejub** | **Afor** | 0 | 0 | 0 | 0 | 0 | 0 | 0 | 0 | 0 | 0 | 0.0% | 0.0% |

| | | | | | | | | | | | | | |
|---|---|---|---|---|---|---|---|---|---|---|---|---|---|
| **Afor_Aaus_Ejub** | **Aaus** | 0 | 0 | 0 | 0 | 0 | 0 | 0 | 0 | 0 | 0 | 0.0% | 0.0% |
| **Afor_Aaus_Ejub** | **Ejub** | 0 | 2.90 | 0.00 | 1.00 | 0.0007 | 0.0022 | -63235 | -56122 | -7113 | 5487 | 100.0% | 0.0% |
| **Afor_Aaus_Curs** | **Afor** | 0 | 0 | 0 | 0 | 0 | 0 | 0 | 0 | 0 | 0 | 0.0% | 0.0% |
| **Afor_Aaus_Curs** | **Aaus** | 0 | 0 | 0 | 0 | 0 | 0 | 0 | 0 | 0 | 1 | 0.0% | 0.0% |
| **Afor_Aaus_Curs** | **Curs** | 0 | 3.79 | 0.00 | 1.00 | 0.0010 | 0.0042 | -58012 | -49011 | -9001 | 5486 | 100.0% | 0.0% |
| **Afor_Aaus_Oros** | **Afor** | 0 | 0 | 0 | 0 | 0 | 0 | 0 | 0 | 0 | 0 | 0.0% | 0.0% |
| **Afor_Aaus_Oros** | **Aaus** | 0 | 0 | 0 | 0 | 0 | 0 | 0 | 0 | 0 | 0 | 0.0% | 0.0% |
| **Afor_Aaus_Oros** | **Oros** | 0 | 5.57 | 0.00 | 1.00 | 0.0022 | 0.0133 | -48584 | -36426 | -12158 | 5487 | 100.0% | 0.0% |
| **Afor_Phoo_Zwol** | **Afor** | 0 | 1.33 | 0.23 | 0.77 | 0.0005 | 0.0006 | -61 | -62 | 1 | 5 | 0.0% | 0.0% |
| **Afor_Phoo_Zwol** | **Phoo** | 0 | 1.68 | 0.79 | 0.21 | 0.0004 | 0.0004 | -64 | -67 | 3 | 5 | 0.0% | 0.0% |
| **Afor_Phoo_Zwol** | **Zwol** | 0 | 2.49 | 0.00 | 1.00 | 0.0005 | 0.0014 | -66765 | -60935 | -5830 | 5477 | 99.8% | 0.0% |
| **Afor_Phoo_Zcal** | **Afor** | 0 | 1.33 | 0.23 | 0.77 | 0.0005 | 0.0006 | -61 | -62 | 1 | 5 | 0.0% | 0.0% |
| **Afor_Phoo_Zcal** | **Phoo** | 0 | 1.68 | 0.79 | 0.21 | 0.0004 | 0.0004 | -64 | -67 | 3 | 5 | 0.0% | 0.0% |
| **Afor_Phoo_Zcal** | **Zcal** | 0 | 2.49 | 0.00 | 1.00 | 0.0005 | 0.0014 | -66765 | -60935 | -5830 | 5477 | 99.8% | 0.0% |
| **Afor_Phoo_Ejub** | **Afor** | 0 | 1.33 | 0.23 | 0.77 | 0.0005 | 0.0006 | -61 | -62 | 1 | 5 | 0.0% | 0.0% |
| **Afor_Phoo_Ejub** | **Phoo** | 0 | 1.68 | 0.79 | 0.21 | 0.0004 | 0.0004 | -64 | -67 | 3 | 5 | 0.0% | 0.0% |
| **Afor_Phoo_Ejub** | **Ejub** | 0 | 2.49 | 0.00 | 1.00 | 0.0005 | 0.0014 | -66765 | -60935 | -5830 | 5477 | 99.8% | 0.0% |
| **Afor_Phoo_Curs** | **Afor** | 0 | 0 | 0 | 0 | 0 | 0 | 0 | 0 | 0 | 0 | 0.0% | 0.0% |
| **Afor_Phoo_Curs** | **Phoo** | 0 | 0 | 0 | 0 | 0 | 0 | 0 | 0 | 0 | 1 | 0.0% | 0.0% |
| **Afor_Phoo_Curs** | **Curs** | 0 | 3.74 | 0.00 | 1.00 | 0.0009 | 0.0034 | -60024 | -51323 | -8701 | 5486 | 100.0% | 0.0% |
| **Afor_Phoo_Oros** | **Afor** | 0 | 0 | 0 | 0 | 0 | 0 | 0 | 0 | 0 | 0 | 0.0% | 0.0% |
| **Afor_Phoo_Oros** | **Phoo** | 0 | 0 | 0 | 0 | 0 | 0 | 0 | 0 | 0 | 0 | 0.0% | 0.0% |
| **Afor_Phoo_Oros** | **Oros** | 0 | 5.68 | 0.00 | 1.00 | 0.0021 | 0.0125 | -49401 | -37108 | -12293 | 5487 | 100.0% | 0.0% |
| **Afor_Zwol_Zcal** | **Afor** | 0 | 3.97 | 0.00 | 1.00 | 0.0006 | 0.0026 | -63348 | -54297 | -9052 | 5487 | 100.0% | 0.0% |
| **Afor_Zwol_Zcal** | **Zwol** | 0 | 0 | 0 | 0 | 0 | 0 | 0 | 0 | 0 | 0 | 0.0% | 0.0% |
| **Afor_Zwol_Zcal** | **Zcal** | 0 | 0 | 0 | 0 | 0 | 0 | 0 | 0 | 0 | 0 | 0.0% | 0.0% |
| **Afor_Zwol_Ejub** | **Afor** | 0 | 3.98 | 0.00 | 1.00 | 0.0006 | 0.0026 | -63349 | -54308 | -9041 | 5487 | 100.0% | 0.0% |
| **Afor_Zwol_Ejub** | **Zwol** | 0 | 0 | 0 | 0 | 0 | 0 | 0 | 0 | 0 | 0 | 0.0% | 0.0% |
| **Afor_Zwol_Ejub** | **Ejub** | 0 | 0 | 0 | 0 | 0 | 0 | 0 | 0 | 0 | 0 | 0.0% | 0.0% |
| **Afor_Zwol_Curs** | **Afor** | 0 | 0 | 0 | 0 | 0 | 0 | 0 | 0 | 0 | 1 | 0.0% | 0.0% |
| **Afor_Zwol_Curs** | **Zwol** | 0 | 14.97 | 0.67 | 0.33 | 0.0005 | 0.0029 | -33 | -28 | -5 | 3 | 0.0% | 0.0% |
| **Afor_Zwol_Curs** | **Curs** | 0 | 3.13 | 0.00 | 1.00 | 0.0006 | 0.0020 | -64466 | -57101 | -7365 | 5483 | 99.9% | 0.0% |

| | | | | | | | | | | | | | |
|---|---|---|---|---|---|---|---|---|---|---|---|---|---|
| **Afor_Zwol_Oros** | **Afor** | 0 | 0 | 0 | 0 | 0 | 0 | 0 | 0 | 0 | 0 | 0.0% | 0.0% |
| **Afor_Zwol_Oros** | **Zwol** | 0 | 0 | 0 | 0 | 0 | 0 | 0 | 0 | 0 | 0 | 0.0% | 0.0% |
| **Afor_Zwol_Oros** | **Oros** | 0 | 5.89 | 0.00 | 1.00 | 0.0018 | 0.0111 | -51052 | -38420 | -12632 | 5487 | 100.0% | 0.0% |
| **Afor_Zcal_Ejub** | **Afor** | 0 | 3.97 | 0.00 | 1.00 | 0.0006 | 0.0027 | -63116 | -54048 | -9069 | 5487 | 100.0% | 0.0% |
| **Afor_Zcal_Ejub** | **Zcal** | 0 | 0 | 0 | 0 | 0 | 0 | 0 | 0 | 0 | 0 | 0.0% | 0.0% |
| **Afor_Zcal_Ejub** | **Ejub** | 0 | 0 | 0 | 0 | 0 | 0 | 0 | 0 | 0 | 0 | 0.0% | 0.0% |
| **Afor_Zcal_Curs** | **Afor** | 0 | 0 | 0 | 0 | 0 | 0 | 0 | 0 | 0 | 1 | 0.0% | 0.0% |
| **Afor_Zcal_Curs** | **Zcal** | 0 | 14.97 | 0.67 | 0.33 | 0.0005 | 0.0029 | -33 | -28 | -5 | 3 | 0.0% | 0.0% |
| **Afor_Zcal_Curs** | **Curs** | 0 | 3.13 | 0.00 | 1.00 | 0.0006 | 0.0020 | -64466 | -57101 | -7365 | 5483 | 99.9% | 0.0% |
| **Afor_Zcal_Oros** | **Afor** | 0 | 0 | 0 | 0 | 0 | 0 | 0 | 0 | 0 | 0 | 0.0% | 0.0% |
| **Afor_Zcal_Oros** | **Zcal** | 0 | 0 | 0 | 0 | 0 | 0 | 0 | 0 | 0 | 0 | 0.0% | 0.0% |
| **Afor_Zcal_Oros** | **Oros** | 0 | 5.89 | 0.00 | 1.00 | 0.0018 | 0.0111 | -51052 | -38420 | -12632 | 5487 | 100.0% | 0.0% |
| **Afor_Ejub_Curs** | **Afor** | 0 | 0 | 0 | 0 | 0 | 0 | 0 | 0 | 0 | 1 | 0.0% | 0.0% |
| **Afor_Ejub_Curs** | **Ejub** | 0 | 14.97 | 0.67 | 0.33 | 0.0005 | 0.0029 | -33 | -28 | -5 | 3 | 0.0% | 0.0% |
| **Afor_Ejub_Curs** | **Curs** | 0 | 3.13 | 0.00 | 1.00 | 0.0006 | 0.0020 | -64466 | -57101 | -7365 | 5483 | 99.9% | 0.0% |
| **Afor_Ejub_Oros** | **Afor** | 0 | 0 | 0 | 0 | 0 | 0 | 0 | 0 | 0 | 0 | 0.0% | 0.0% |
| **Afor_Ejub_Oros** | **Ejub** | 0 | 0 | 0 | 0 | 0 | 0 | 0 | 0 | 0 | 0 | 0.0% | 0.0% |
| **Afor_Ejub_Oros** | **Oros** | 0 | 5.89 | 0.00 | 1.00 | 0.0018 | 0.0111 | -51052 | -38420 | -12632 | 5487 | 100.0% | 0.0% |
| **Afor_Curs_Oros** | **Afor** | 0 | 0 | 0 | 0 | 0 | 0 | 0 | 0 | 0 | 0 | 0.0% | 0.0% |
| **Afor_Curs_Oros** | **Curs** | 0 | 0 | 0 | 0 | 0 | 0 | 0 | 0 | 0 | 0 | 0.0% | 0.0% |
| **Afor_Curs_Oros** | **Oros** | 0 | 6.17 | 0.00 | 1.00 | 0.0014 | 0.0091 | -53686 | -40615 | -13071 | 5487 | 100.0% | 0.0% |
| **Agal_Aaus_Phoo** | **Agal** | 0 | 12.35 | 0.96 | 0.04 | 0.0004 | 0.0004 | -379 | -370 | -8 | 27 | 0.0% | 0.0% |
| **Agal_Aaus_Phoo** | **Aaus** | 0 | 0.69 | 0.52 | 0.48 | 0.0003 | 0.0003 | -262 | -267 | 5 | 19 | 0.0% | 0.0% |
| **Agal_Aaus_Phoo** | **Phoo** | 0 | 1.91 | 0.01 | 0.99 | 0.0005 | 0.0011 | -67364 | -63129 | -4234 | 5441 | 98.3% | 0.0% |
| **Agal_Aaus_Zwol** | **Agal** | 0 | 0 | 0 | 0 | 0 | 0 | 0 | 0 | 0 | 0 | 0.0% | 0.0% |
| **Agal_Aaus_Zwol** | **Aaus** | 0 | 0 | 0 | 0 | 0 | 0 | 0 | 0 | 0 | 0 | 0.0% | 0.0% |
| **Agal_Aaus_Zwol** | **Zwol** | 0 | 2.94 | 0.00 | 1.00 | 0.0008 | 0.0025 | -61907 | -54733 | -7174 | 5487 | 100.0% | 0.0% |
| **Agal_Aaus_Zcal** | **Agal** | 0 | 0 | 0 | 0 | 0 | 0 | 0 | 0 | 0 | 0 | 0.0% | 0.0% |
| **Agal_Aaus_Zcal** | **Aaus** | 0 | 0 | 0 | 0 | 0 | 0 | 0 | 0 | 0 | 0 | 0.0% | 0.0% |
| **Agal_Aaus_Zcal** | **Zcal** | 0 | 2.94 | 0.00 | 1.00 | 0.0008 | 0.0025 | -61907 | -54733 | -7174 | 5487 | 100.0% | 0.0% |
| **Agal_Aaus_Ejub** | **Agal** | 0 | 0 | 0 | 0 | 0 | 0 | 0 | 0 | 0 | 0 | 0.0% | 0.0% |
| **Agal_Aaus_Ejub** | **Aaus** | 0 | 0 | 0 | 0 | 0 | 0 | 0 | 0 | 0 | 0 | 0.0% | 0.0% |

| | | | | | | | | | | | | | |
|---|---|---|---|---|---|---|---|---|---|---|---|---|---|
| **Agal_Aaus_Ejub** | **Ejub** | 0 | 2.94 | 0.00 | 1.00 | 0.0008 | 0.0025 | -61907 | -54733 | -7174 | 5487 | 100.0% | 0.0% |
| **Agal_Aaus_Curs** | **Agal** | 0 | 0 | 0 | 0 | 0 | 0 | 0 | 0 | 0 | 0 | 0.0% | 0.0% |
| **Agal_Aaus_Curs** | **Aaus** | 0 | 0 | 0 | 0 | 0 | 0 | 0 | 0 | 0 | 0 | 0.0% | 0.0% |
| **Agal_Aaus_Curs** | **Curs** | 0 | 3.74 | 0.00 | 1.00 | 0.0011 | 0.0045 | -57189 | -48272 | -8916 | 5487 | 100.0% | 0.0% |
| **Agal_Aaus_Oros** | **Agal** | 0 | 0 | 0 | 0 | 0 | 0 | 0 | 0 | 0 | 0 | 0.0% | 0.0% |
| **Agal_Aaus_Oros** | **Aaus** | 0 | 0 | 0 | 0 | 0 | 0 | 0 | 0 | 0 | 0 | 0.0% | 0.0% |
| **Agal_Aaus_Oros** | **Oros** | 0 | 5.51 | 0.00 | 1.00 | 0.0023 | 0.0136 | -48235 | -36182 | -12053 | 5487 | 100.0% | 0.0% |
| **Agal_Phoo_Zwol** | **Agal** | 0 | 1.33 | 0.23 | 0.77 | 0.0005 | 0.0006 | -61 | -62 | 1 | 5 | 0.0% | 0.0% |
| **Agal_Phoo_Zwol** | **Phoo** | 0 | 1.68 | 0.79 | 0.21 | 0.0004 | 0.0004 | -64 | -67 | 3 | 5 | 0.0% | 0.0% |
| **Agal_Phoo_Zwol** | **Zwol** | 0 | 2.51 | 0.00 | 1.00 | 0.0005 | 0.0014 | -66785 | -60943 | -5842 | 5477 | 99.8% | 0.0% |
| **Agal_Phoo_Zcal** | **Agal** | 0 | 1.33 | 0.23 | 0.77 | 0.0005 | 0.0006 | -61 | -62 | 1 | 5 | 0.0% | 0.0% |
| **Agal_Phoo_Zcal** | **Phoo** | 0 | 1.68 | 0.79 | 0.21 | 0.0004 | 0.0004 | -64 | -67 | 3 | 5 | 0.0% | 0.0% |
| **Agal_Phoo_Zcal** | **Zcal** | 0 | 2.51 | 0.00 | 1.00 | 0.0005 | 0.0014 | -66785 | -60943 | -5842 | 5477 | 99.8% | 0.0% |
| **Agal_Phoo_Ejub** | **Agal** | 0 | 1.33 | 0.23 | 0.77 | 0.0005 | 0.0006 | -61 | -62 | 1 | 5 | 0.0% | 0.0% |
| **Agal_Phoo_Ejub** | **Phoo** | 0 | 1.68 | 0.79 | 0.21 | 0.0004 | 0.0004 | -64 | -67 | 3 | 5 | 0.0% | 0.0% |
| **Agal_Phoo_Ejub** | **Ejub** | 0 | 2.51 | 0.00 | 1.00 | 0.0005 | 0.0014 | -66785 | -60943 | -5842 | 5477 | 99.8% | 0.0% |
| **Agal_Phoo_Curs** | **Agal** | 0 | 0 | 0 | 0 | 0 | 0 | 0 | 0 | 0 | 1 | 0.0% | 0.0% |
| **Agal_Phoo_Curs** | **Phoo** | 0 | 0 | 0 | 0 | 0 | 0 | 0 | 0 | 0 | 1 | 0.0% | 0.0% |
| **Agal_Phoo_Curs** | **Curs** | 0 | 3.75 | 0.00 | 1.00 | 0.0009 | 0.0034 | -60034 | -51314 | -8720 | 5485 | 100.0% | 0.0% |
| **Agal_Phoo_Oros** | **Agal** | 0 | 0 | 0 | 0 | 0 | 0 | 0 | 0 | 0 | 0 | 0.0% | 0.0% |
| **Agal_Phoo_Oros** | **Phoo** | 0 | 0 | 0 | 0 | 0 | 0 | 0 | 0 | 0 | 0 | 0.0% | 0.0% |
| **Agal_Phoo_Oros** | **Oros** | 0 | 5.69 | 0.00 | 1.00 | 0.0021 | 0.0125 | -49411 | -37109 | -12302 | 5487 | 100.0% | 0.0% |
| **Agal_Zwol_Zcal** | **Agal** | 0 | 3.97 | 0.00 | 1.00 | 0.0006 | 0.0026 | -63348 | -54297 | -9052 | 5487 | 100.0% | 0.0% |
| **Agal_Zwol_Zcal** | **Zwol** | 0 | 0 | 0 | 0 | 0 | 0 | 0 | 0 | 0 | 0 | 0.0% | 0.0% |
| **Agal_Zwol_Zcal** | **Zcal** | 0 | 0 | 0 | 0 | 0 | 0 | 0 | 0 | 0 | 0 | 0.0% | 0.0% |
| **Agal_Zwol_Ejub** | **Agal** | 0 | 3.98 | 0.00 | 1.00 | 0.0006 | 0.0026 | -63349 | -54308 | -9041 | 5487 | 100.0% | 0.0% |
| **Agal_Zwol_Ejub** | **Zwol** | 0 | 0 | 0 | 0 | 0 | 0 | 0 | 0 | 0 | 0 | 0.0% | 0.0% |
| **Agal_Zwol_Ejub** | **Ejub** | 0 | 0 | 0 | 0 | 0 | 0 | 0 | 0 | 0 | 0 | 0.0% | 0.0% |
| **Agal_Zwol_Curs** | **Agal** | 0 | 0 | 0 | 0 | 0 | 0 | 0 | 0 | 0 | 1 | 0.0% | 0.0% |
| **Agal_Zwol_Curs** | **Zwol** | 0 | 12.58 | 0.67 | 0.33 | 0.0005 | 0.0025 | -33 | -29 | -4 | 3 | 0.0% | 0.0% |
| **Agal_Zwol_Curs** | **Curs** | 0 | 3.13 | 0.00 | 1.00 | 0.0006 | 0.0020 | -64466 | -57101 | -7365 | 5483 | 99.9% | 0.0% |
| **Agal_Zwol_Oros** | **Agal** | 0 | 0 | 0 | 0 | 0 | 0 | 0 | 0 | 0 | 0 | 0.0% | 0.0% |

| | | | | | | | | | | | | | |
|---|---|---|---|---|---|---|---|---|---|---|---|---|---|
| **Agal_Zwol_Oros** | **Zwol** | 0 | 0 | 0 | 0 | 0 | 0 | 0 | 0 | 0 | 0 | 0.0% | 0.0% |
| **Agal_Zwol_Oros** | **Oros** | 0 | 5.89 | 0.00 | 1.00 | 0.0018 | 0.0111 | -51052 | -38420 | -12632 | 5487 | 100.0% | 0.0% |
| **Agal_Zcal_Ejub** | **Agal** | 0 | 3.97 | 0.00 | 1.00 | 0.0006 | 0.0027 | -63116 | -54048 | -9069 | 5487 | 100.0% | 0.0% |
| **Agal_Zcal_Ejub** | **Zcal** | 0 | 0 | 0 | 0 | 0 | 0 | 0 | 0 | 0 | 0 | 0.0% | 0.0% |
| **Agal_Zcal_Ejub** | **Ejub** | 0 | 0 | 0 | 0 | 0 | 0 | 0 | 0 | 0 | 0 | 0.0% | 0.0% |
| **Agal_Zcal_Curs** | **Agal** | 0 | 0 | 0 | 0 | 0 | 0 | 0 | 0 | 0 | 1 | 0.0% | 0.0% |
| **Agal_Zcal_Curs** | **Zcal** | 0 | 12.58 | 0.67 | 0.33 | 0.0005 | 0.0025 | -33 | -29 | -4 | 3 | 0.0% | 0.0% |
| **Agal_Zcal_Curs** | **Curs** | 0 | 3.13 | 0.00 | 1.00 | 0.0006 | 0.0020 | -64466 | -57101 | -7365 | 5483 | 99.9% | 0.0% |
| **Agal_Zcal_Oros** | **Agal** | 0 | 0 | 0 | 0 | 0 | 0 | 0 | 0 | 0 | 0 | 0.0% | 0.0% |
| **Agal_Zcal_Oros** | **Zcal** | 0 | 0 | 0 | 0 | 0 | 0 | 0 | 0 | 0 | 0 | 0.0% | 0.0% |
| **Agal_Zcal_Oros** | **Oros** | 0 | 5.89 | 0.00 | 1.00 | 0.0018 | 0.0111 | -51052 | -38420 | -12632 | 5487 | 100.0% | 0.0% |
| **Agal_Ejub_Curs** | **Agal** | 0 | 0 | 0 | 0 | 0 | 0 | 0 | 0 | 0 | 1 | 0.0% | 0.0% |
| **Agal_Ejub_Curs** | **Ejub** | 0 | 12.58 | 0.67 | 0.33 | 0.0005 | 0.0025 | -33 | -29 | -4 | 3 | 0.0% | 0.0% |
| **Agal_Ejub_Curs** | **Curs** | 0 | 3.13 | 0.00 | 1.00 | 0.0006 | 0.0020 | -64466 | -57101 | -7365 | 5483 | 99.9% | 0.0% |
| **Agal_Ejub_Oros** | **Agal** | 0 | 0 | 0 | 0 | 0 | 0 | 0 | 0 | 0 | 0 | 0.0% | 0.0% |
| **Agal_Ejub_Oros** | **Ejub** | 0 | 0 | 0 | 0 | 0 | 0 | 0 | 0 | 0 | 0 | 0.0% | 0.0% |
| **Agal_Ejub_Oros** | **Oros** | 0 | 5.89 | 0.00 | 1.00 | 0.0018 | 0.0111 | -51052 | -38420 | -12632 | 5487 | 100.0% | 0.0% |
| **Agal_Curs_Oros** | **Agal** | 0 | 0 | 0 | 0 | 0 | 0 | 0 | 0 | 0 | 0 | 0.0% | 0.0% |
| **Agal_Curs_Oros** | **Curs** | 0 | 0 | 0 | 0 | 0 | 0 | 0 | 0 | 0 | 0 | 0.0% | 0.0% |
| **Agal_Curs_Oros** | **Oros** | 0 | 6.17 | 0.00 | 1.00 | 0.0014 | 0.0091 | -53688 | -40615 | -13072 | 5487 | 100.0% | 0.0% |
| **Aaus_Phoo_Zwol** | **Aaus** | 0 | 1.33 | 0.23 | 0.77 | 0.0005 | 0.0006 | -61 | -62 | 1 | 5 | 0.0% | 0.0% |
| **Aaus_Phoo_Zwol** | **Phoo** | 0 | 1.68 | 0.79 | 0.21 | 0.0004 | 0.0004 | -64 | -67 | 3 | 5 | 0.0% | 0.0% |
| **Aaus_Phoo_Zwol** | **Zwol** | 0 | 2.50 | 0.00 | 1.00 | 0.0005 | 0.0014 | -66777 | -60937 | -5839 | 5477 | 99.8% | 0.0% |
| **Aaus_Phoo_Zcal** | **Aaus** | 0 | 1.33 | 0.23 | 0.77 | 0.0005 | 0.0006 | -61 | -62 | 1 | 5 | 0.0% | 0.0% |
| **Aaus_Phoo_Zcal** | **Phoo** | 0 | 1.68 | 0.79 | 0.21 | 0.0004 | 0.0004 | -64 | -67 | 3 | 5 | 0.0% | 0.0% |
| **Aaus_Phoo_Zcal** | **Zcal** | 0 | 2.50 | 0.00 | 1.00 | 0.0005 | 0.0014 | -66777 | -60937 | -5839 | 5477 | 99.8% | 0.0% |
| **Aaus_Phoo_Ejub** | **Aaus** | 0 | 1.33 | 0.23 | 0.77 | 0.0005 | 0.0006 | -61 | -62 | 1 | 5 | 0.0% | 0.0% |
| **Aaus_Phoo_Ejub** | **Phoo** | 0 | 1.68 | 0.79 | 0.21 | 0.0004 | 0.0004 | -64 | -67 | 3 | 5 | 0.0% | 0.0% |
| **Aaus_Phoo_Ejub** | **Ejub** | 0 | 2.50 | 0.00 | 1.00 | 0.0005 | 0.0014 | -66777 | -60937 | -5839 | 5477 | 99.8% | 0.0% |
| **Aaus_Phoo_Curs** | **Aaus** | 0 | 0 | 0 | 0 | 0 | 0 | 0 | 0 | 0 | 1 | 0.0% | 0.0% |
| **Aaus_Phoo_Curs** | **Phoo** | 0 | 0 | 0 | 0 | 0 | 0 | 0 | 0 | 0 | 1 | 0.0% | 0.0% |

| | | | | | | | | | | | | | |
|---|---|---|---|---|---|---|---|---|---|---|---|---|---|
| **Aaus_Phoo_Curs** | **Curs** | 0 | 3.74 | 0.00 | 1.00 | 0.0009 | 0.0034 | -60024 | -51312 | -8712 | 5485 | 100.0% | 0.0% |
| **Aaus_Phoo_Oros** | **Aaus** | 0 | 0 | 0 | 0 | 0 | 0 | 0 | 0 | 0 | 0 | 0.0% | 0.0% |
| **Aaus_Phoo_Oros** | **Phoo** | 0 | 0 | 0 | 0 | 0 | 0 | 0 | 0 | 0 | 0 | 0.0% | 0.0% |
| **Aaus_Phoo_Oros** | **Oros** | 0 | 5.69 | 0.00 | 1.00 | 0.0021 | 0.0125 | -49406 | -37108 | - 12297 | 5487 | 100.0% | 0.0% |
| **Aaus_Zwol_Zcal** | **Aaus** | 0 | 3.97 | 0.00 | 1.00 | 0.0006 | 0.0026 | -63348 | -54297 | -9052 | 5487 | 100.0% | 0.0% |
| **Aaus_Zwol_Zcal** | **Zwol** | 0 | 0 | 0 | 0 | 0 | 0 | 0 | 0 | 0 | 0 | 0.0% | 0.0% |
| **Aaus_Zwol_Zcal** | **Zcal** | 0 | 0 | 0 | 0 | 0 | 0 | 0 | 0 | 0 | 0 | 0.0% | 0.0% |
| **Aaus_Zwol_Ejub** | **Aaus** | 0 | 3.98 | 0.00 | 1.00 | 0.0006 | 0.0026 | -63349 | -54308 | -9041 | 5487 | 100.0% | 0.0% |
| **Aaus_Zwol_Ejub** | **Zwol** | 0 | 0 | 0 | 0 | 0 | 0 | 0 | 0 | 0 | 0 | 0.0% | 0.0% |
| **Aaus_Zwol_Ejub** | **Ejub** | 0 | 0 | 0 | 0 | 0 | 0 | 0 | 0 | 0 | 0 | 0.0% | 0.0% |
| **Aaus_Zwol_Curs** | **Aaus** | 0 | 0 | 0 | 0 | 0 | 0 | 0 | 0 | 0 | 1 | 0.0% | 0.0% |
| **Aaus_Zwol_Curs** | **Zwol** | 0 | 12.58 | 0.67 | 0.33 | 0.0005 | 0.0025 | -33 | -29 | -4 | 3 | 0.0% | 0.0% |
| **Aaus_Zwol_Curs** | **Curs** | 0 | 3.13 | 0.00 | 1.00 | 0.0006 | 0.0020 | -64466 | -57101 | -7365 | 5483 | 99.9% | 0.0% |
| **Aaus_Zwol_Oros** | **Aaus** | 0 | 0 | 0 | 0 | 0 | 0 | 0 | 0 | 0 | 0 | 0.0% | 0.0% |
| **Aaus_Zwol_Oros** | **Zwol** | 0 | 0 | 0 | 0 | 0 | 0 | 0 | 0 | 0 | 0 | 0.0% | 0.0% |
| **Aaus_Zwol_Oros** | **Oros** | 0 | 5.89 | 0.00 | 1.00 | 0.0018 | 0.0111 | -51052 | -38420 | - 12632 | 5487 | 100.0% | 0.0% |
| **Aaus_Zcal_Ejub** | **Aaus** | 0 | 3.97 | 0.00 | 1.00 | 0.0006 | 0.0027 | -63116 | -54048 | -9069 | 5487 | 100.0% | 0.0% |
| **Aaus_Zcal_Ejub** | **Zcal** | 0 | 0 | 0 | 0 | 0 | 0 | 0 | 0 | 0 | 0 | 0.0% | 0.0% |
| **Aaus_Zcal_Ejub** | **Ejub** | 0 | 0 | 0 | 0 | 0 | 0 | 0 | 0 | 0 | 0 | 0.0% | 0.0% |
| **Aaus_Zcal_Curs** | **Aaus** | 0 | 0 | 0 | 0 | 0 | 0 | 0 | 0 | 0 | 1 | 0.0% | 0.0% |
| **Aaus_Zcal_Curs** | **Zcal** | 0 | 12.58 | 0.67 | 0.33 | 0.0005 | 0.0025 | -33 | -29 | -4 | 3 | 0.0% | 0.0% |
| **Aaus_Zcal_Curs** | **Curs** | 0 | 3.13 | 0.00 | 1.00 | 0.0006 | 0.0020 | -64466 | -57101 | -7365 | 5483 | 99.9% | 0.0% |
| **Aaus_Zcal_Oros** | **Aaus** | 0 | 0 | 0 | 0 | 0 | 0 | 0 | 0 | 0 | 0 | 0.0% | 0.0% |
| **Aaus_Zcal_Oros** | **Zcal** | 0 | 0 | 0 | 0 | 0 | 0 | 0 | 0 | 0 | 0 | 0.0% | 0.0% |
| **Aaus_Zcal_Oros** | **Oros** | 0 | 5.89 | 0.00 | 1.00 | 0.0018 | 0.0111 | -51052 | -38420 | - 12632 | 5487 | 100.0% | 0.0% |
| **Aaus_Ejub_Curs** | **Aaus** | 0 | 0 | 0 | 0 | 0 | 0 | 0 | 0 | 0 | 1 | 0.0% | 0.0% |
| **Aaus_Ejub_Curs** | **Ejub** | 0 | 12.58 | 0.67 | 0.33 | 0.0005 | 0.0025 | -33 | -29 | -4 | 3 | 0.0% | 0.0% |
| **Aaus_Ejub_Curs** | **Curs** | 0 | 3.13 | 0.00 | 1.00 | 0.0006 | 0.0020 | -64466 | -57101 | -7365 | 5483 | 99.9% | 0.0% |
| **Aaus_Ejub_Oros** | **Aaus** | 0 | 0 | 0 | 0 | 0 | 0 | 0 | 0 | 0 | 0 | 0.0% | 0.0% |
| **Aaus_Ejub_Oros** | **Ejub** | 0 | 0 | 0 | 0 | 0 | 0 | 0 | 0 | 0 | 0 | 0.0% | 0.0% |
| **Aaus_Ejub_Oros** | **Oros** | 0 | 5.89 | 0.00 | 1.00 | 0.0018 | 0.0111 | -51052 | -38420 | - 12632 | 5487 | 100.0% | 0.0% |

| | | | | | | | | | | | | | |
|---|---|---|---|---|---|---|---|---|---|---|---|---|---|
| **Aaus_Curs_Oros** | **Aaus** | 0 | 0 | 0 | 0 | 0 | 0 | 0 | 0 | 0 | 0 | 0.0% | 0.0% |
| **Aaus_Curs_Oros** | **Curs** | 0 | 0 | 0 | 0 | 0 | 0 | 0 | 0 | 0 | 0 | 0.0% | 0.0% |
| **Aaus_Curs_Oros** | **Oros** | 0 | 6.17 | 0.00 | 1.00 | 0.0014 | 0.0091 | -53688 | -40615 | -13072 | 5487 | 100.0% | 0.0% |
| **Phoo_Zwol_Zcal** | **Phoo** | 0 | 3.97 | 0.00 | 1.00 | 0.0006 | 0.0026 | -63346 | -54298 | -9048 | 5487 | 100.0% | 0.0% |
| **Phoo_Zwol_Zcal** | **Zwol** | 0 | 0 | 0 | 0 | 0 | 0 | 0 | 0 | 0 | 0 | 0.0% | 0.0% |
| **Phoo_Zwol_Zcal** | **Zcal** | 0 | 0 | 0 | 0 | 0 | 0 | 0 | 0 | 0 | 0 | 0.0% | 0.0% |
| **Phoo_Zwol_Ejub** | **Phoo** | 0 | 3.98 | 0.00 | 1.00 | 0.0006 | 0.0026 | -63346 | -54309 | -9037 | 5487 | 100.0% | 0.0% |
| **Phoo_Zwol_Ejub** | **Zwol** | 0 | 0 | 0 | 0 | 0 | 0 | 0 | 0 | 0 | 0 | 0.0% | 0.0% |
| **Phoo_Zwol_Ejub** | **Ejub** | 0 | 0 | 0 | 0 | 0 | 0 | 0 | 0 | 0 | 0 | 0.0% | 0.0% |
| **Phoo_Zwol_Curs** | **Phoo** | 0 | 0 | 0 | 0 | 0 | 0 | 0 | 0 | 0 | 1 | 0.0% | 0.0% |
| **Phoo_Zwol_Curs** | **Zwol** | 0 | 14.97 | 0.67 | 0.33 | 0.0005 | 0.0029 | -33 | -28 | -5 | 3 | 0.0% | 0.0% |
| **Phoo_Zwol_Curs** | **Curs** | 0 | 3.13 | 0.00 | 1.00 | 0.0006 | 0.0020 | -64466 | -57099 | -7366 | 5483 | 99.9% | 0.0% |
| **Phoo_Zwol_Oros** | **Phoo** | 0 | 0 | 0 | 0 | 0 | 0 | 0 | 0 | 0 | 0 | 0.0% | 0.0% |
| **Phoo_Zwol_Oros** | **Zwol** | 0 | 0 | 0 | 0 | 0 | 0 | 0 | 0 | 0 | 0 | 0.0% | 0.0% |
| **Phoo_Zwol_Oros** | **Oros** | 0 | 5.89 | 0.00 | 1.00 | 0.0018 | 0.0111 | -51054 | -38420 | -12635 | 5487 | 100.0% | 0.0% |
| **Phoo_Zcal_Ejub** | **Phoo** | 0 | 3.97 | 0.00 | 1.00 | 0.0006 | 0.0027 | -63114 | -54049 | -9065 | 5487 | 100.0% | 0.0% |
| **Phoo_Zcal_Ejub** | **Zcal** | 0 | 0 | 0 | 0 | 0 | 0 | 0 | 0 | 0 | 0 | 0.0% | 0.0% |
| **Phoo_Zcal_Ejub** | **Ejub** | 0 | 0 | 0 | 0 | 0 | 0 | 0 | 0 | 0 | 0 | 0.0% | 0.0% |
| **Phoo_Zcal_Curs** | **Phoo** | 0 | 0 | 0 | 0 | 0 | 0 | 0 | 0 | 0 | 1 | 0.0% | 0.0% |
| **Phoo_Zcal_Curs** | **Zcal** | 0 | 14.97 | 0.67 | 0.33 | 0.0005 | 0.0029 | -33 | -28 | -5 | 3 | 0.0% | 0.0% |
| **Phoo_Zcal_Curs** | **Curs** | 0 | 3.13 | 0.00 | 1.00 | 0.0006 | 0.0020 | -64466 | -57099 | -7366 | 5483 | 99.9% | 0.0% |
| **Phoo_Zcal_Oros** | **Phoo** | 0 | 0 | 0 | 0 | 0 | 0 | 0 | 0 | 0 | 0 | 0.0% | 0.0% |
| **Phoo_Zcal_Oros** | **Zcal** | 0 | 0 | 0 | 0 | 0 | 0 | 0 | 0 | 0 | 0 | 0.0% | 0.0% |
| **Phoo_Zcal_Oros** | **Oros** | 0 | 5.89 | 0.00 | 1.00 | 0.0018 | 0.0111 | -51054 | -38420 | -12635 | 5487 | 100.0% | 0.0% |
| **Phoo_Ejub_Curs** | **Phoo** | 0 | 0 | 0 | 0 | 0 | 0 | 0 | 0 | 0 | 1 | 0.0% | 0.0% |
| **Phoo_Ejub_Curs** | **Ejub** | 0 | 14.97 | 0.67 | 0.33 | 0.0005 | 0.0029 | -33 | -28 | -5 | 3 | 0.0% | 0.0% |
| **Phoo_Ejub_Curs** | **Curs** | 0 | 3.13 | 0.00 | 1.00 | 0.0006 | 0.0020 | -64466 | -57099 | -7366 | 5483 | 99.9% | 0.0% |
| **Phoo_Ejub_Oros** | **Phoo** | 0 | 0 | 0 | 0 | 0 | 0 | 0 | 0 | 0 | 0 | 0.0% | 0.0% |
| **Phoo_Ejub_Oros** | **Ejub** | 0 | 0 | 0 | 0 | 0 | 0 | 0 | 0 | 0 | 0 | 0.0% | 0.0% |
| **Phoo_Ejub_Oros** | **Oros** | 0 | 5.89 | 0.00 | 1.00 | 0.0018 | 0.0111 | -51054 | -38420 | -12635 | 5487 | 100.0% | 0.0% |
| **Phoo_Curs_Oros** | **Phoo** | 0 | 0 | 0 | 0 | 0 | 0 | 0 | 0 | 0 | 0 | 0.0% | 0.0% |
| **Phoo_Curs_Oros** | **Curs** | 0 | 0 | 0 | 0 | 0 | 0 | 0 | 0 | 0 | 0 | 0.0% | 0.0% |

| | | | | | | | | | | | | | |
|---|---|---|---|---|---|---|---|---|---|---|---|---|---|
| **Phoo_Curs_Oros** | **Oros** | 0 | 6.17 | 0.00 | 1.00 | 0.0014 | 0.0091 | -53687 | -40615 | -13072 | 5487 | 100.0% | 0.0% |
| **Zwol_Zcal_Ejub** | **Zwol** | 0 | 0.81 | 0.61 | 0.39 | 0.0002 | 0.0002 | -42248 | -42111 | -137 | 2800 | 19.8% | 0.0% |
| **Zwol_Zcal_Ejub** | **Zcal** | 0 | 0.98 | 0.79 | 0.21 | 0.0002 | 0.0002 | -20088 | -20077 | -11 | 1302 | 4.9% | 0.0% |
| **Zwol_Zcal_Ejub** | **Ejub** | 0 | 0.69 | 0.56 | 0.44 | 0.0002 | 0.0002 | -21415 | -21339 | -76 | 1385 | 11.2% | 0.0% |
| **Zwol_Zcal_Curs** | **Zwol** | 0 | 0 | 0 | 0 | 0 | 0 | 0 | 0 | 0 | 0 | 0.0% | 0.0% |
| **Zwol_Zcal_Curs** | **Zcal** | 0 | 0 | 0 | 0 | 0 | 0 | 0 | 0 | 0 | 0 | 0.0% | 0.0% |
| **Zwol_Zcal_Curs** | **Curs** | 0 | 4.39 | 0.00 | 1.00 | 0.0010 | 0.0046 | -58032 | -48026 | -10005 | 5487 | 100.0% | 0.0% |
| **Zwol_Zcal_Oros** | **Zwol** | 0 | 0 | 0 | 0 | 0 | 0 | 0 | 0 | 0 | 0 | 0.0% | 0.0% |
| **Zwol_Zcal_Oros** | **Zcal** | 0 | 0 | 0 | 0 | 0 | 0 | 0 | 0 | 0 | 0 | 0.0% | 0.0% |
| **Zwol_Zcal_Oros** | **Oros** | 0 | 5.79 | 0.00 | 1.00 | 0.0022 | 0.0137 | -48576 | -36101 | -12476 | 5487 | 100.0% | 0.0% |
| **Zwol_Ejub_Curs** | **Zwol** | 0 | 0 | 0 | 0 | 0 | 0 | 0 | 0 | 0 | 0 | 0.0% | 0.0% |
| **Zwol_Ejub_Curs** | **Ejub** | 0 | 0 | 0 | 0 | 0 | 0 | 0 | 0 | 0 | 0 | 0.0% | 0.0% |
| **Zwol_Ejub_Curs** | **Curs** | 0 | 4.39 | 0.00 | 1.00 | 0.0010 | 0.0046 | -58031 | -48033 | -9998 | 5487 | 100.0% | 0.0% |
| **Zwol_Ejub_Oros** | **Zwol** | 0 | 0 | 0 | 0 | 0 | 0 | 0 | 0 | 0 | 0 | 0.0% | 0.0% |
| **Zwol_Ejub_Oros** | **Ejub** | 0 | 0 | 0 | 0 | 0 | 0 | 0 | 0 | 0 | 0 | 0.0% | 0.0% |
| **Zwol_Ejub_Oros** | **Oros** | 0 | 5.79 | 0.00 | 1.00 | 0.0022 | 0.0137 | -48572 | -36103 | -12469 | 5487 | 100.0% | 0.0% |
| **Zwol_Curs_Oros** | **Zwol** | 0 | 0 | 0 | 0 | 0 | 0 | 0 | 0 | 0 | 0 | 0.0% | 0.0% |
| **Zwol_Curs_Oros** | **Curs** | 0 | 0 | 0 | 0 | 0 | 0 | 0 | 0 | 0 | 0 | 0.0% | 0.0% |
| **Zwol_Curs_Oros** | **Oros** | 0 | 6.17 | 0.00 | 1.00 | 0.0014 | 0.0091 | -53689 | -40616 | -13073 | 5487 | 100.0% | 0.0% |
| **Zcal_Ejub_Curs** | **Zcal** | 0 | 0 | 0 | 0 | 0 | 0 | 0 | 0 | 0 | 0 | 0.0% | 0.0% |
| **Zcal_Ejub_Curs** | **Ejub** | 0 | 0 | 0 | 0 | 0 | 0 | 0 | 0 | 0 | 0 | 0.0% | 0.0% |
| **Zcal_Ejub_Curs** | **Curs** | 0 | 4.39 | 0.00 | 1.00 | 0.0010 | 0.0047 | -57882 | -47885 | -9997 | 5487 | 100.0% | 0.0% |
| **Zcal_Ejub_Oros** | **Zcal** | 0 | 0 | 0 | 0 | 0 | 0 | 0 | 0 | 0 | 0 | 0.0% | 0.0% |
| **Zcal_Ejub_Oros** | **Ejub** | 0 | 0 | 0 | 0 | 0 | 0 | 0 | 0 | 0 | 0 | 0.0% | 0.0% |
| **Zcal_Ejub_Oros** | **Oros** | 0 | 5.78 | 0.00 | 1.00 | 0.0023 | 0.0138 | -48501 | -36053 | -12448 | 5487 | 100.0% | 0.0% |
| **Zcal_Curs_Oros** | **Zcal** | 0 | 0 | 0 | 0 | 0 | 0 | 0 | 0 | 0 | 0 | 0.0% | 0.0% |
| **Zcal_Curs_Oros** | **Curs** | 0 | 0 | 0 | 0 | 0 | 0 | 0 | 0 | 0 | 0 | 0.0% | 0.0% |
| **Zcal_Curs_Oros** | **Oros** | 0 | 6.17 | 0.00 | 1.00 | 0.0014 | 0.0091 | -53689 | -40616 | -13073 | 5487 | 100.0% | 0.0% |
| **Ejub_Curs_Oros** | **Ejub** | 0 | 0 | 0 | 0 | 0 | 0 | 0 | 0 | 0 | 0 | 0.0% | 0.0% |
| **ub_Curs_Oros** | **Curs** | 0 | 0 | 0 | 0 | 0 | 0 | 0 | 0 | 0 | 0 | 0.0% | 0.0% |
| **Ejub_Curs_Oros** | **Oros** | 0 | 6.17 | 0.00 | 1.00 | 0.0014 | 0.0091 | -53689 | -40616 | -13073 | 5487 | 100.0% | 0.0% |

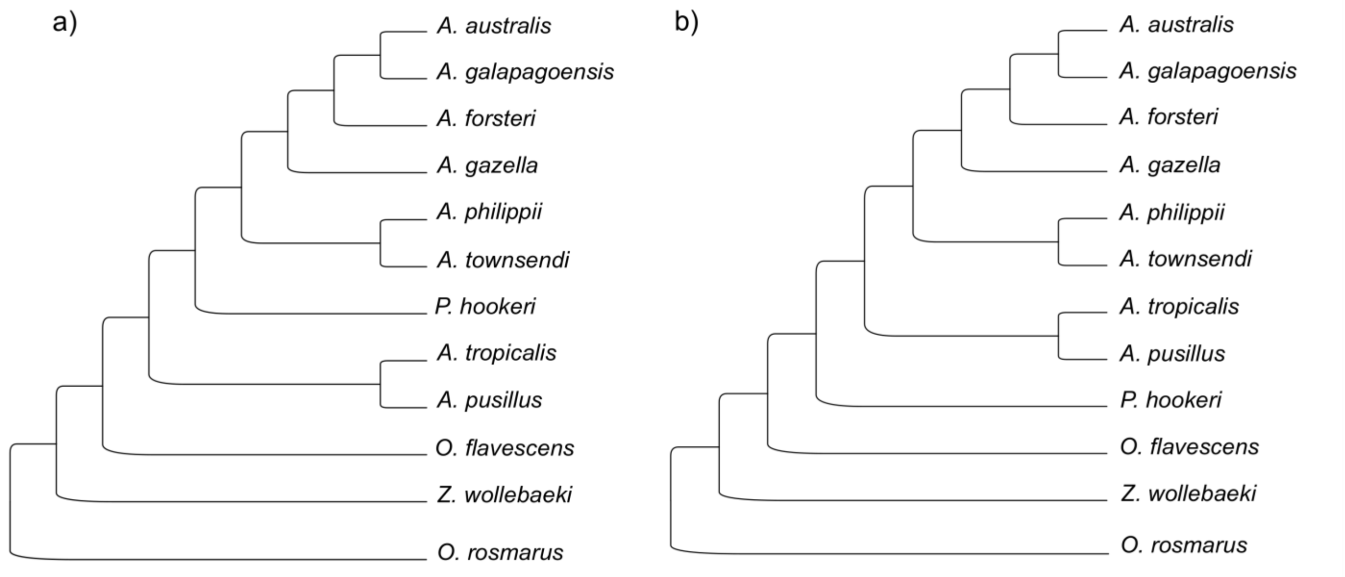


**Figure S1.** Species trees recovered with ASTRAL-III for the different genomic fragment sizes. All branches with support of 1. (A) Species tree recovered for the 10 kb GF. (B) Species tree recovered for the 20, 50, 80, 100, and 200 kb GFs.

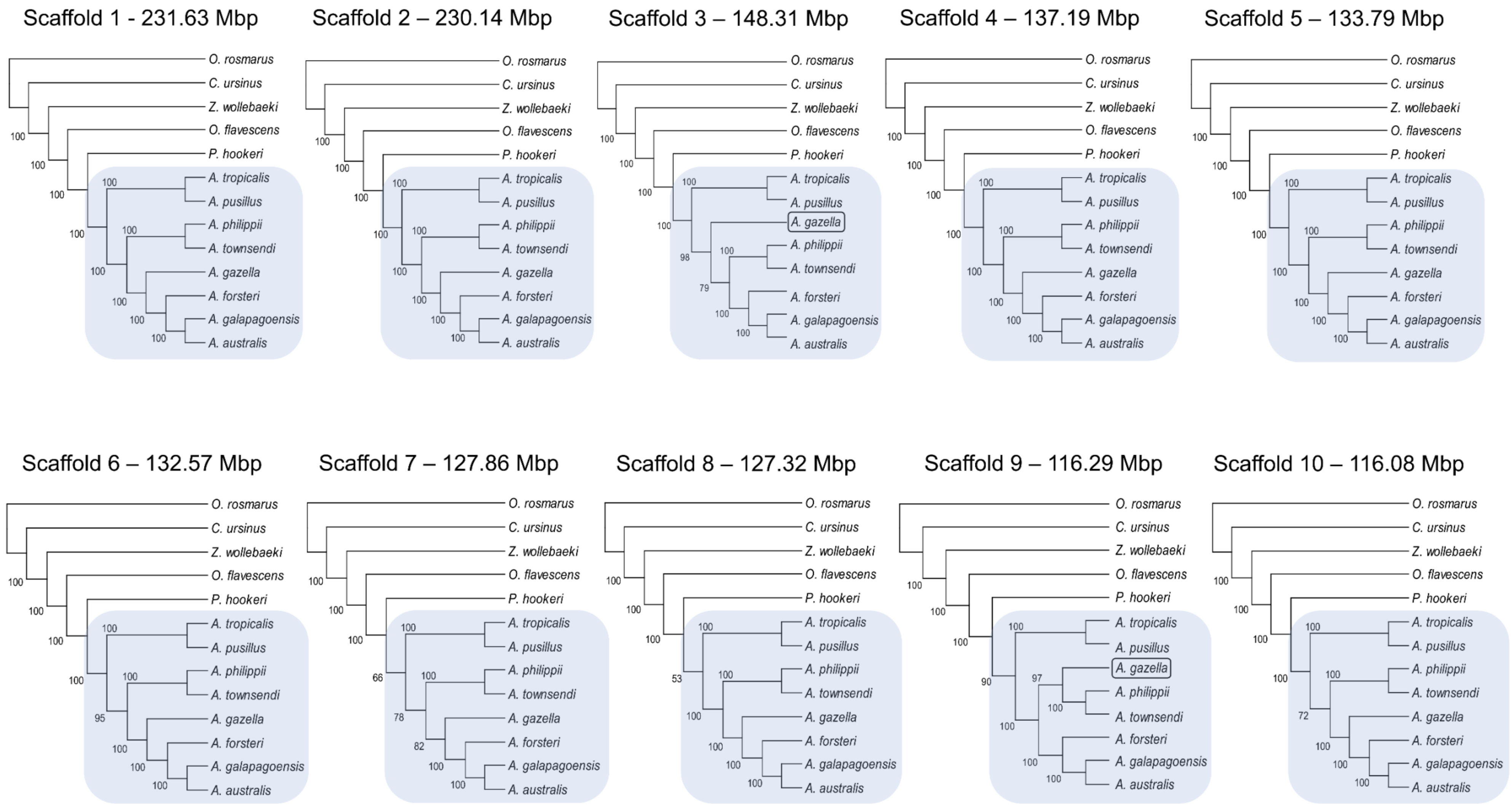


**Figure S2.** Scaffold sizes and their respective ML bootstrapped topology supporting monophyly of *Arctocephalus*.

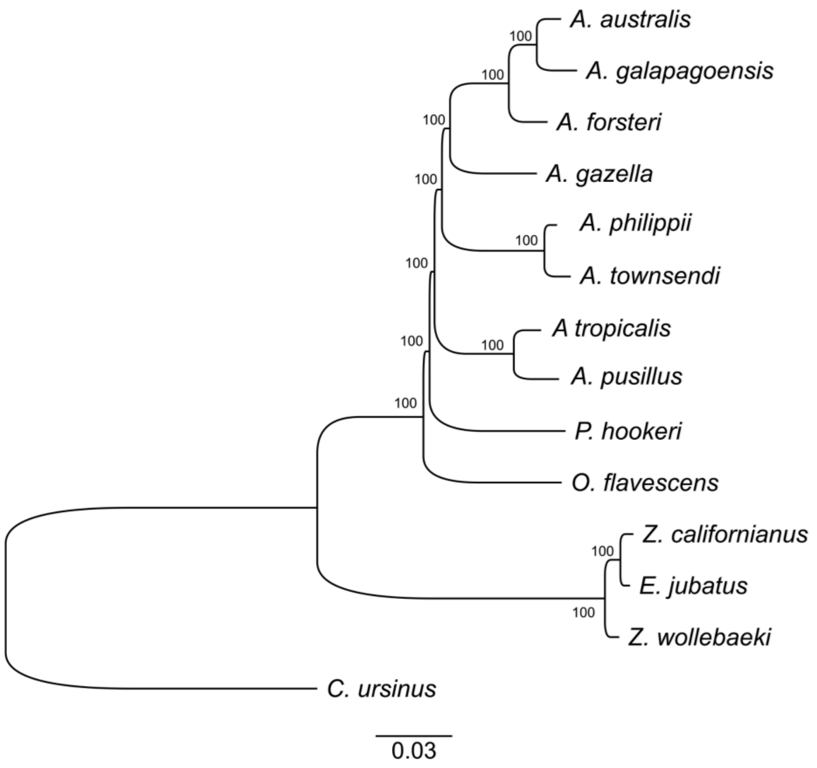


**Figure S3.** Maximum-Likelihood tree of the whole genomes.

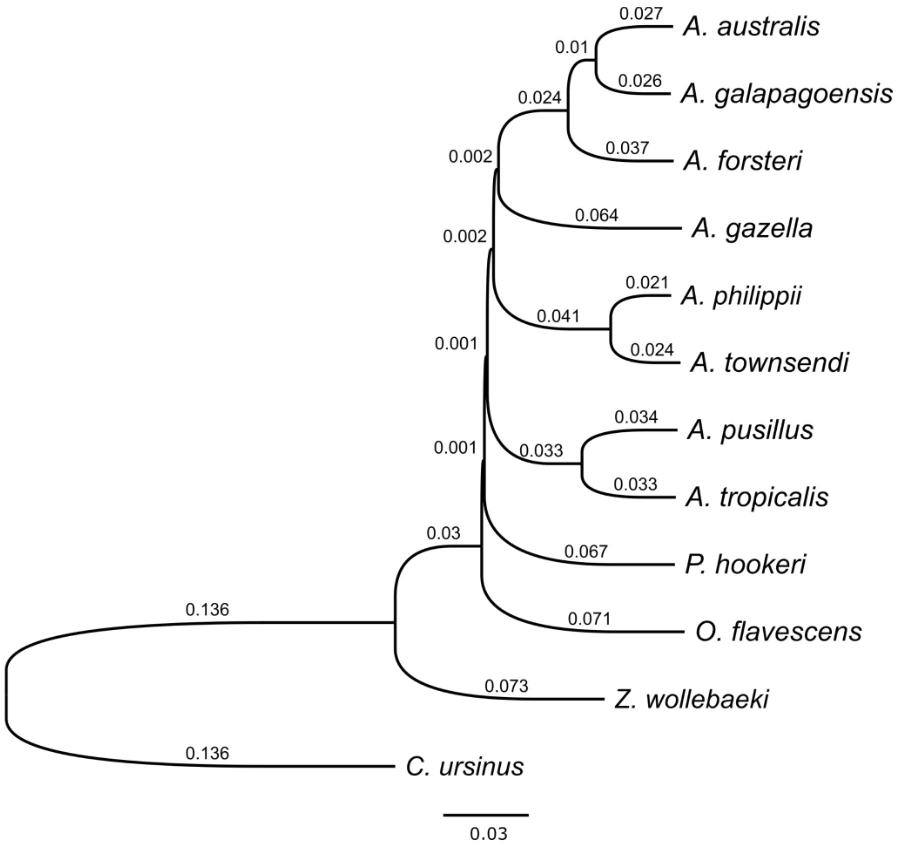


**Figure S4.** Whole-genome genetic distance tree showing the same topology as the species trees recovered in the GFs of 20-200 kb.

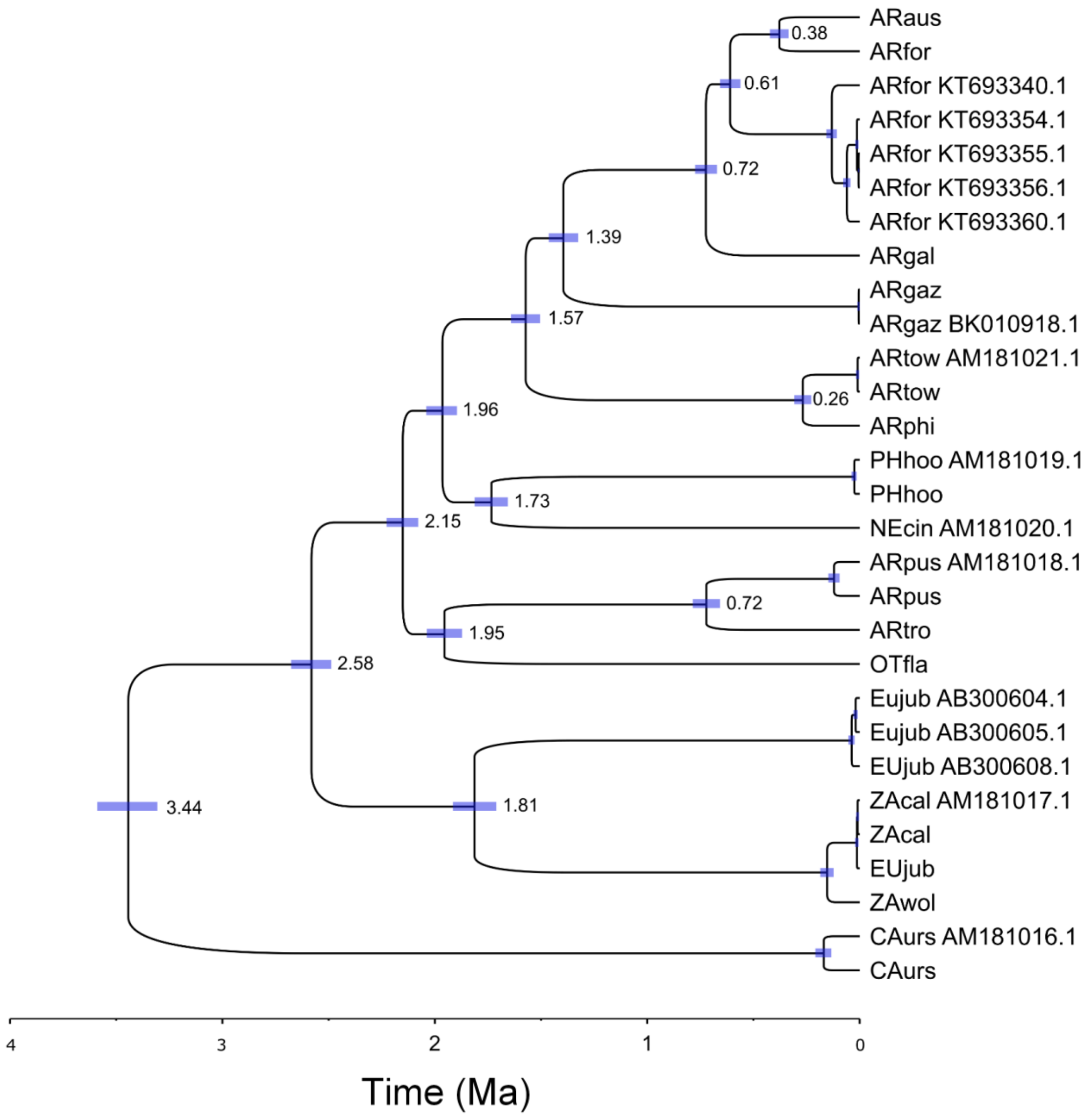


**Figure S5.** Bayesian mitogenome phylogeny and divergence times. Mitogenomes used for species validation have their GenBank Access Number. Nodes with HPD >= 0.99.

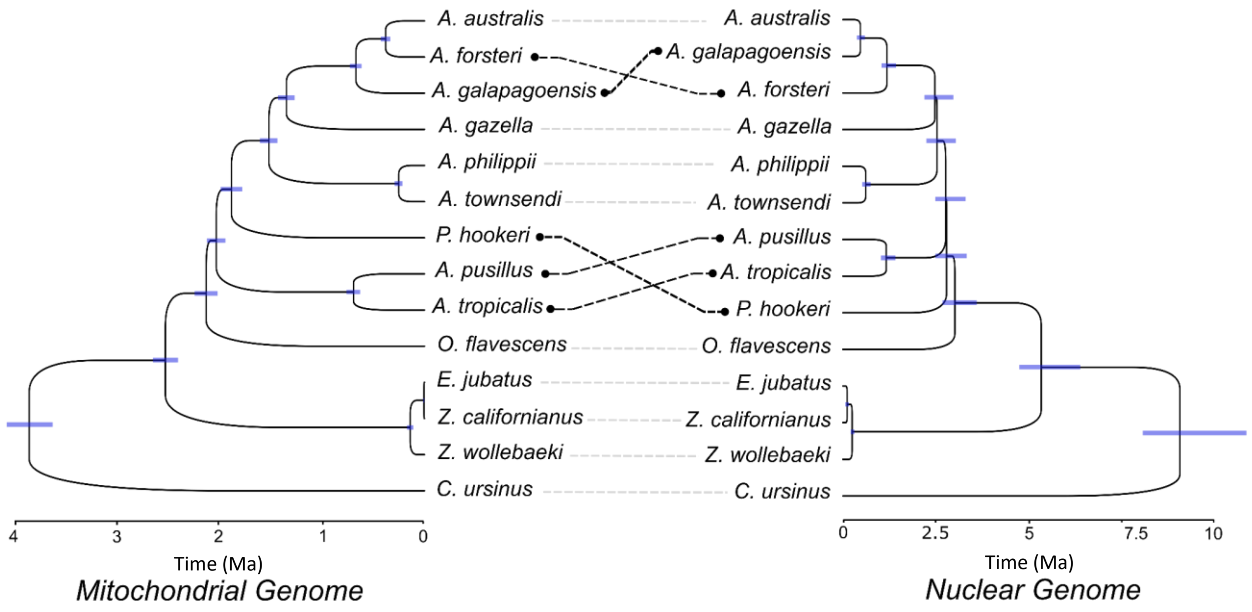


**Figure S6.** Comparison of mitochondrial and nuclear genomes topology of Otariidae.

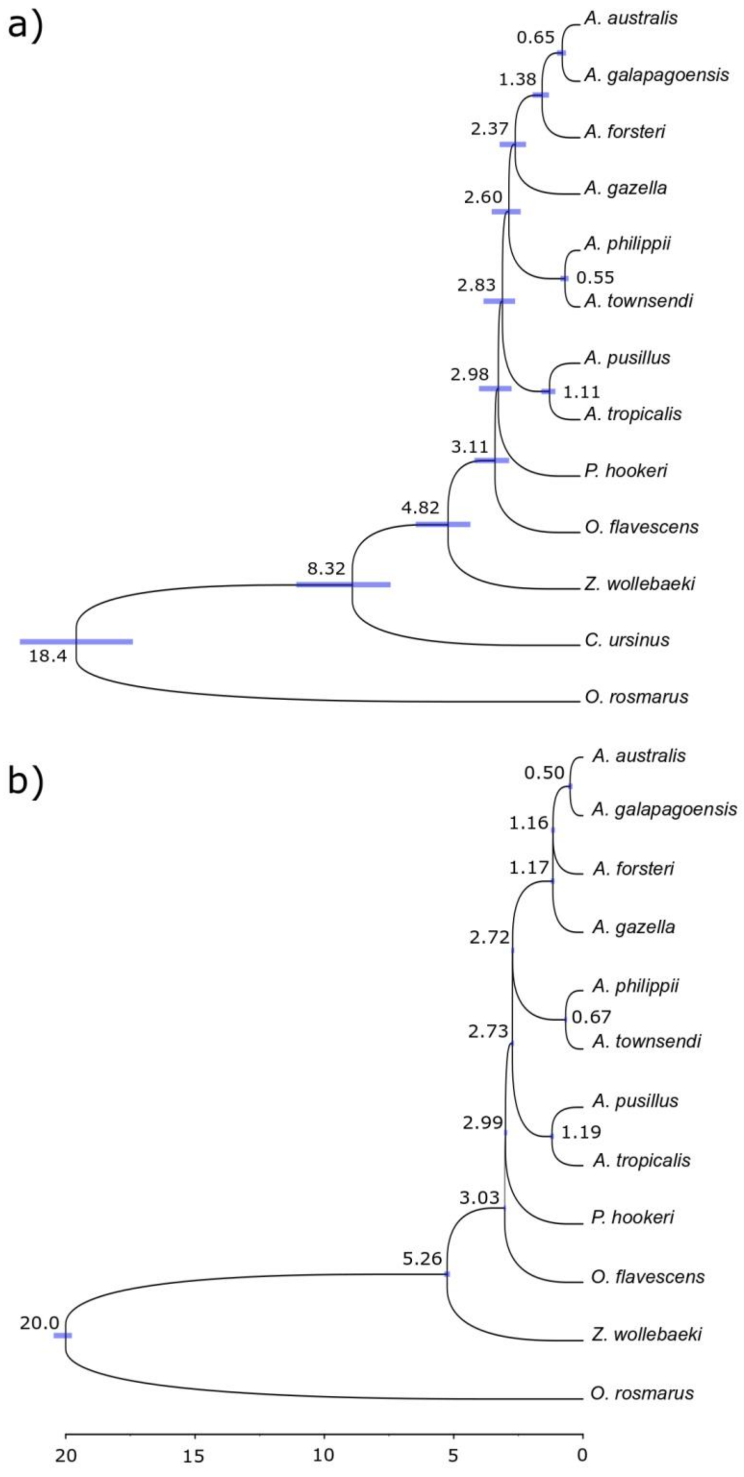


**Figure S7.** Multispecies coalescent trees and divergence times recovered with 300 GF of 50 kb with two different approaches: (a) Calibrated species tree recovered with StarBEAST2. Most nodes with highest posterior density HPD = 1. (b) Divergence times (Mya) estimated with BP&P. Blue bars represent the confidence interval of 95%.

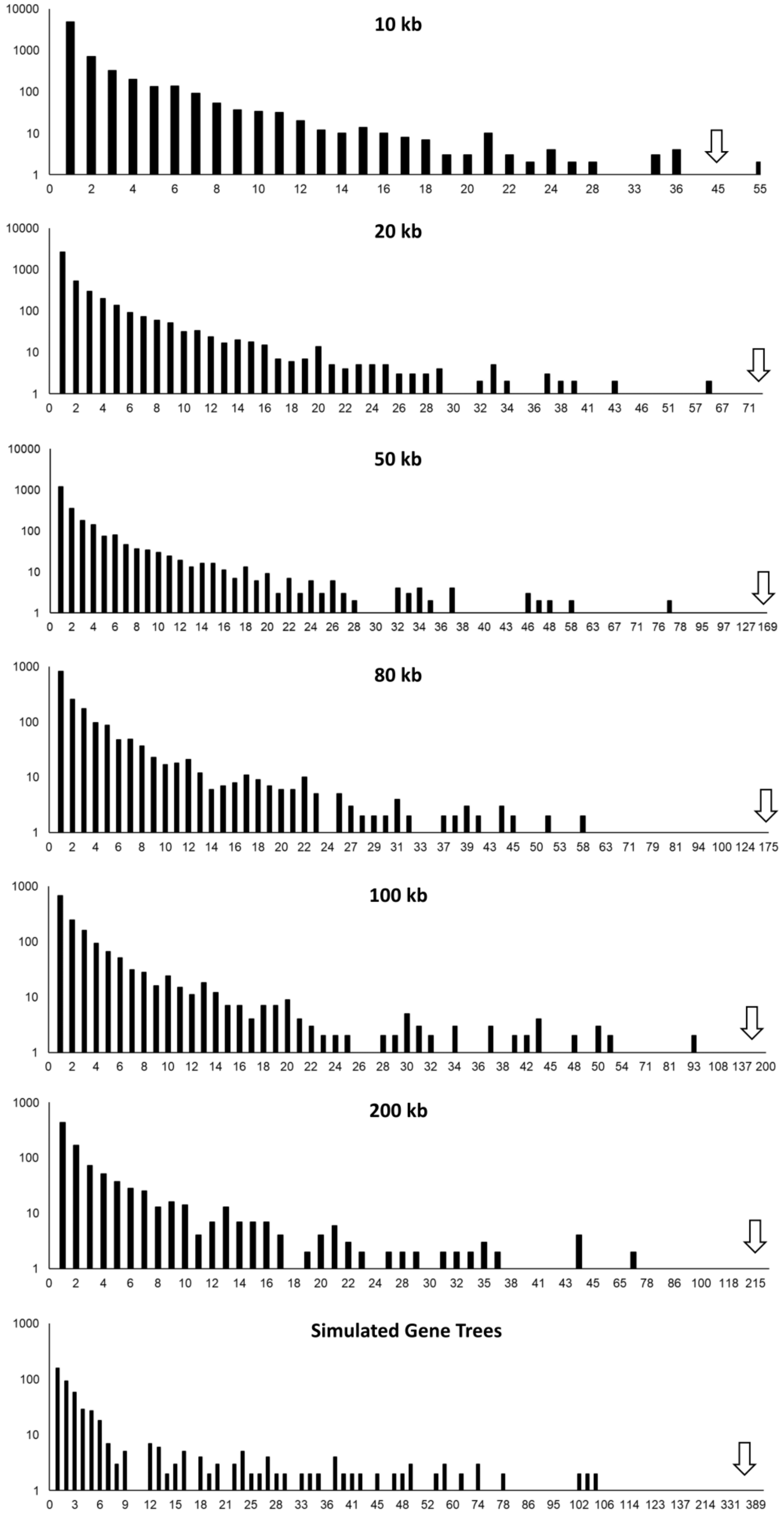


**Figure S8.** The extensive incongruence of topologies in different partitions. The x-axis presents the absolute frequency of a given topology, and the y-axis, cumulative frequencies (log-scale). Arrows indicate the frequency of the species tree.

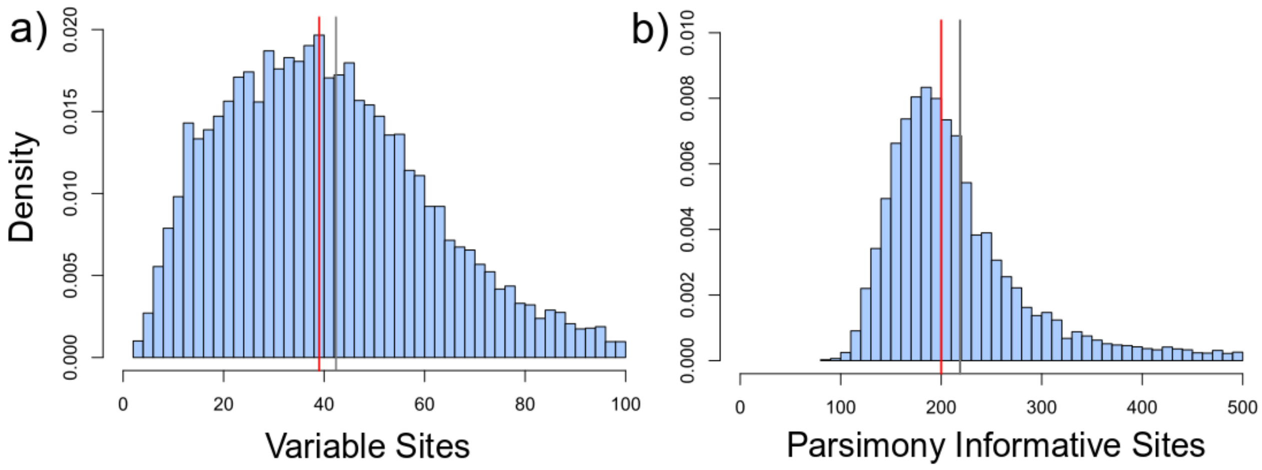


**Figure S9.** Average (red line) and median (grey line) of genetic variation in the GFs of 50 kb. (a) Variable sites between two closest species (*A. australis* and *A. galapagoensis*). (b) Parsimony informative sites in the multispecies GFs.

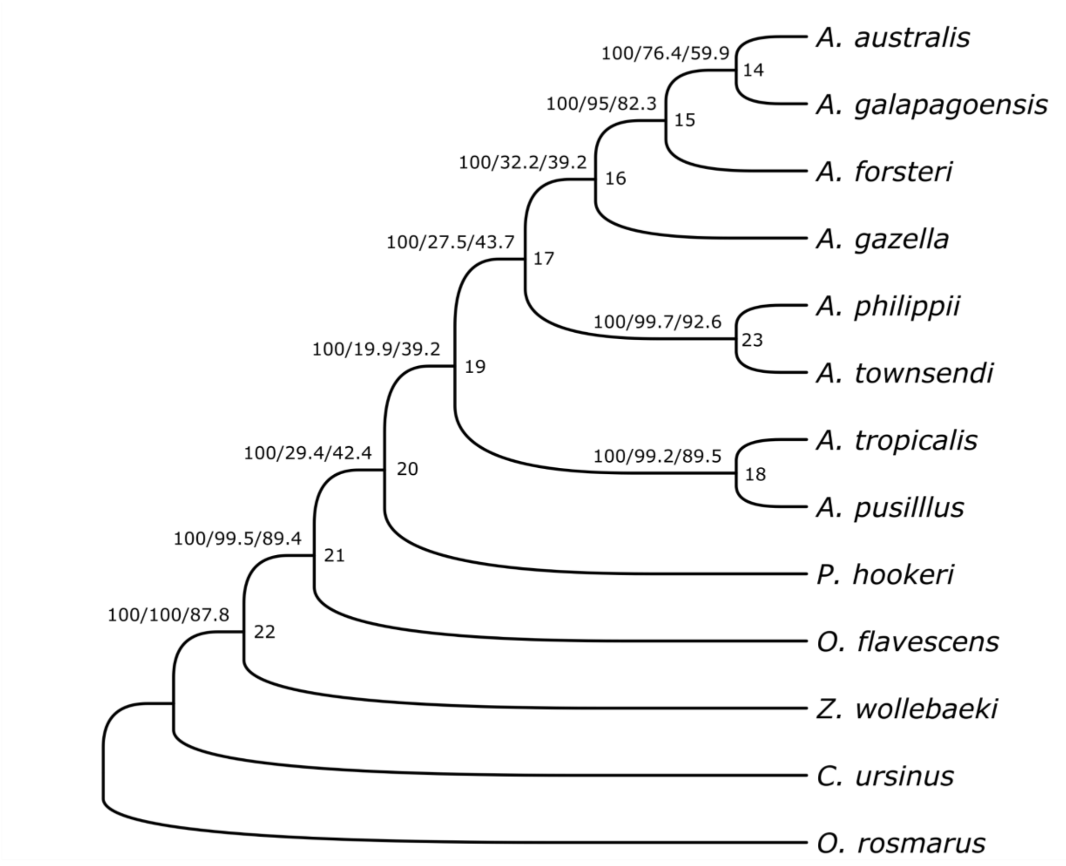


**Figure S10.** Species tree recovered with IQ-TREE. External node values are branch support, gene concordance factor (gCF), and site concordance factor (sCF), respectively. Internal node values are node names. These results show the high discordance of gene trees and the small number of informative sites for the nodes of fast speciation despite the high branch support.

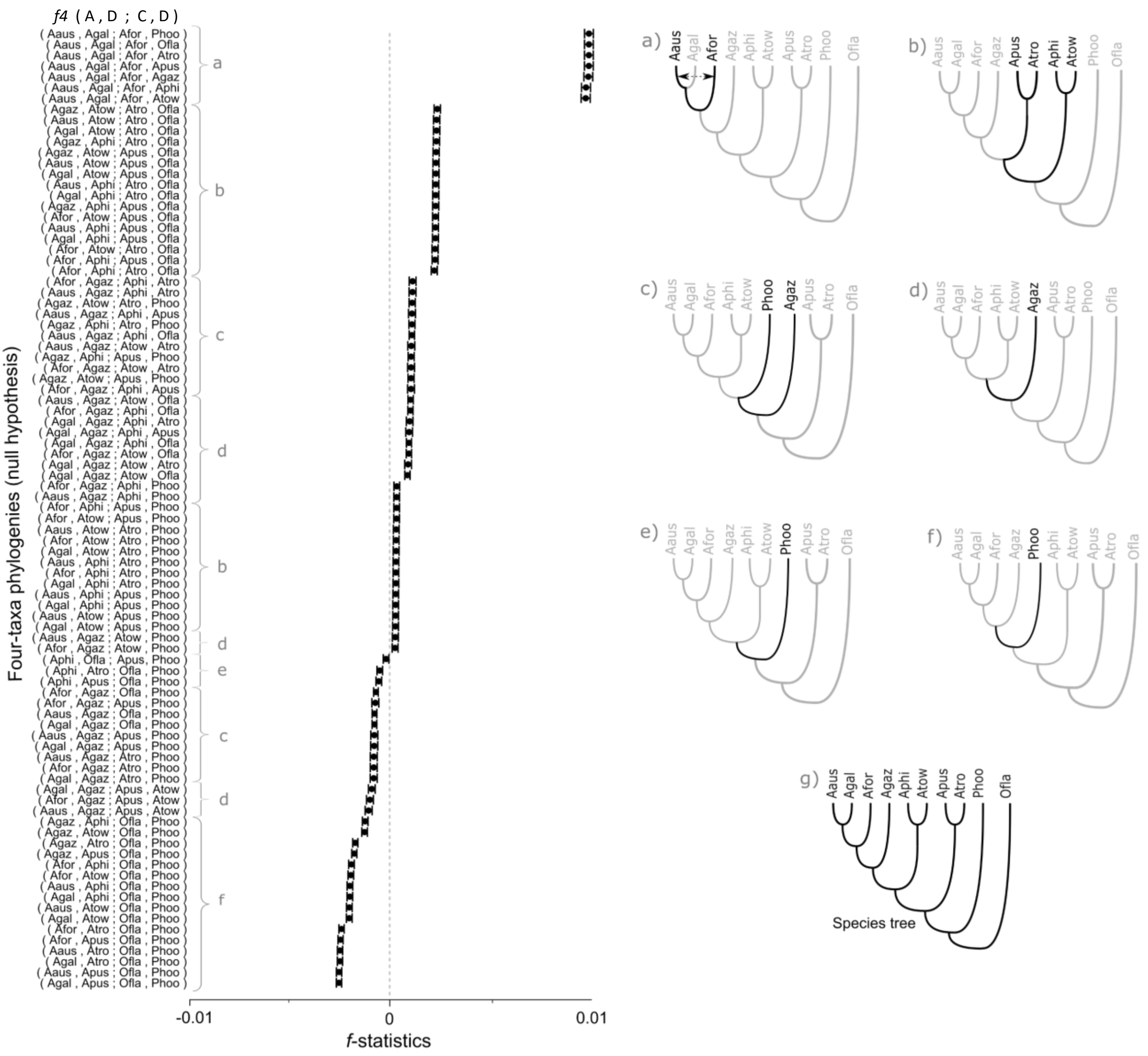


**Figure S11.** Distribution of the signals of $f_4$-statistics (x-axis) for the four-taxa phylogenies combinations (shown on the y-axis) with significant values (|Z-score| > 3). Significantly positive $f_4$ implies gene flow between A and C, or B and D, while significantly negative value implies gene flow between A and D, or B and C. Significant f4 values may also be interpreted as a rejection of the given topology. From *a* to *f*, the alternative branch position (topology) implied by the results. *g* depicts the species tree (null model).

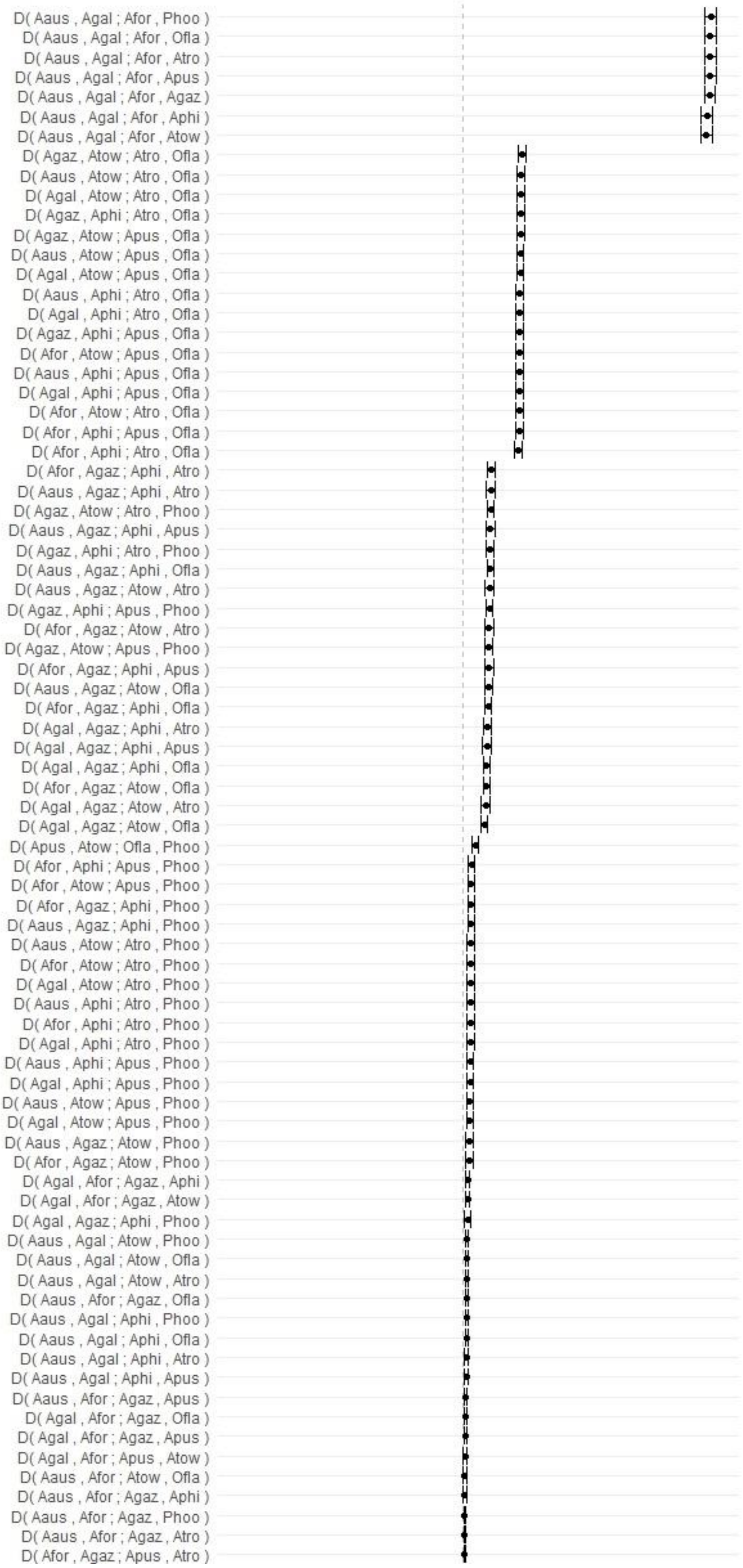


**Figure S12.** $F_4$-statistics distribution (x-axis) for all significant combination of four-taxa phylogenies (y-axis) with significant values (|Z-score| > 3).

**Figure S12.** Cont.

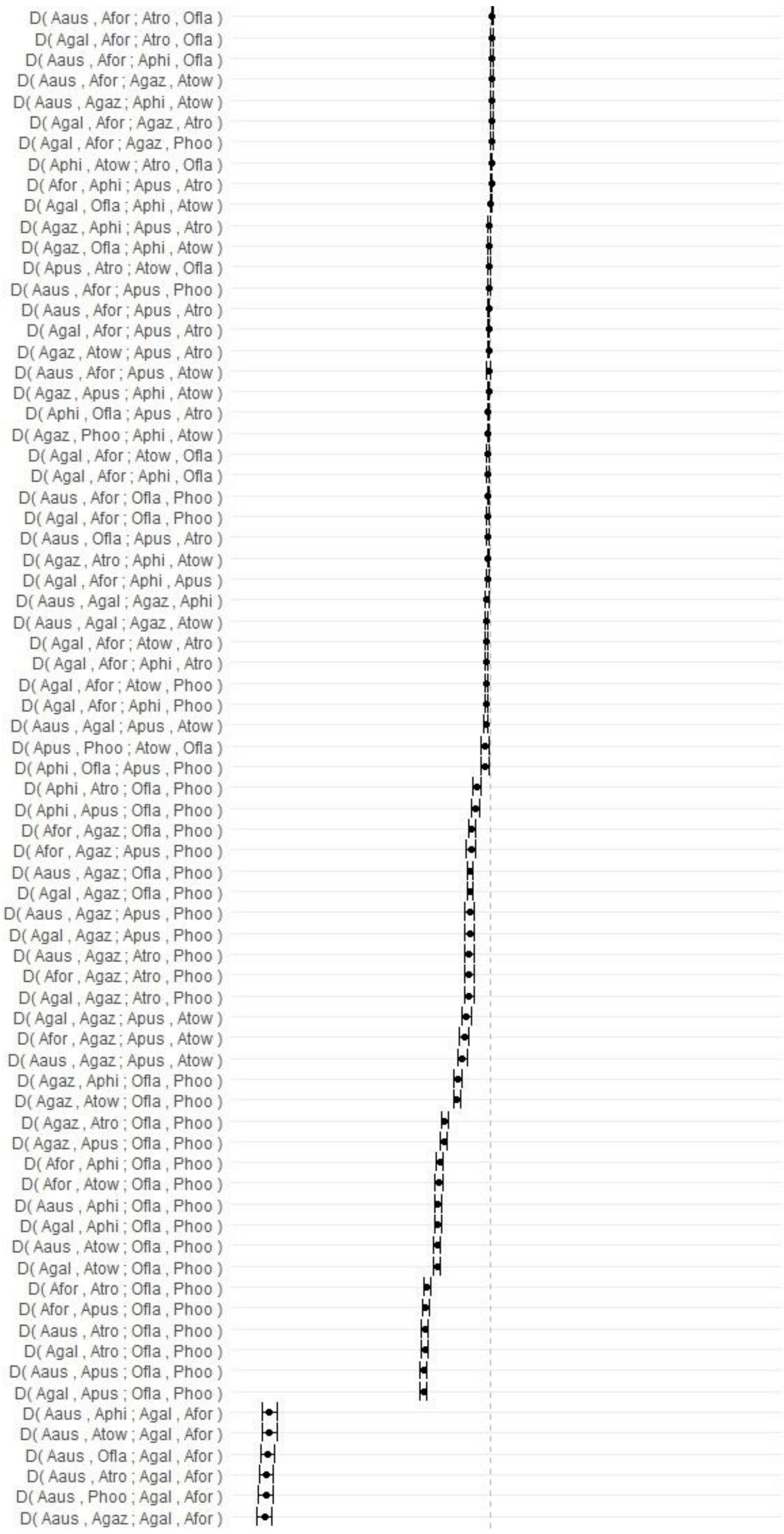

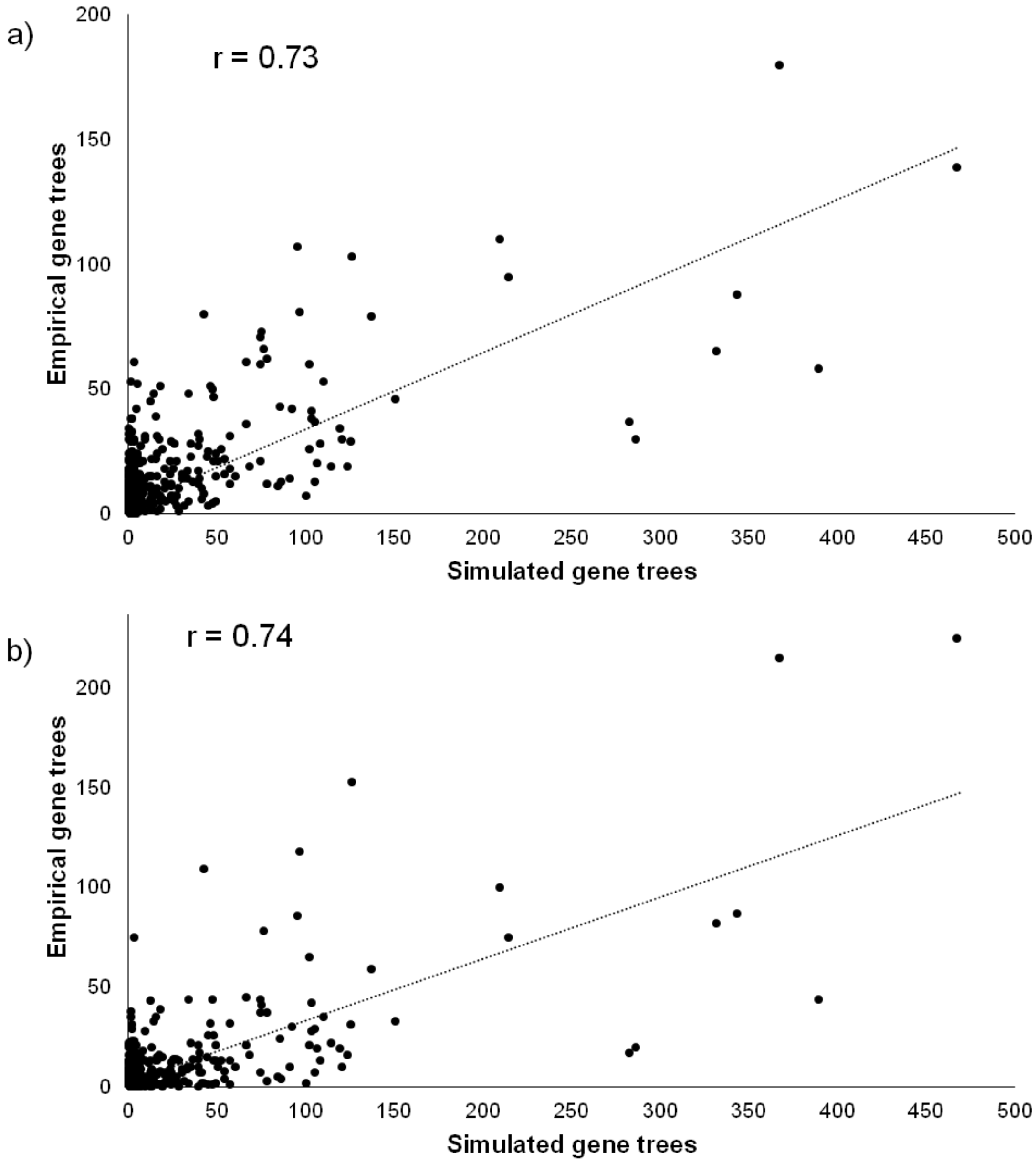


**Figure S13.** Correlation of the number of topologies observed (empirical gene trees) and the number of topologies generated by simulation. (a) 50 kb GFs and (b) 200 kb GFs.
